# Small-molecule modulators of HIPK4 activity and proteostasis

**DOI:** 10.64898/2026.05.12.724395

**Authors:** Zaile Zhuang, Riley K. Togashi, Patrick Kearney, Ian Pass, Steven M. Swick, Fu-Yue Zeng, Andrey A. Bobkov, Lynn M. Fujimoto, Shubhankar Dutta, Athina Zerva, Nicolai D. Raig, Debasmita Saha, Atoosa Emami, Martin P. Schwalm, Bradley K. Moon, Samuel T. Howard, Stefan Knapp, Thomas Hanke, Thomas D.Y. Chung, James K. Chen

## Abstract

Homeodomain-interacting protein kinase 4 (HIPK4) is a dual-specificity kinase that is predominantly expressed in differentiating spermatids, required for sperm development, and a promising target for nonhormonal male contraception. Genetic and functional studies have established an essential role for HIPK4 in spermiogenesis, where it acts at least in part through regulation of the F-actin-scaffolded acroplaxome during spermatid head shaping. The direct molecular targets of HIPK4 and their downstream effectors remain poorly defined, and small-molecule probes would be versatile tools for further investigating HIPK4 functions. Synthetic HIPK4 ligands could also be valuable leads for the development of nonhormonal male contraceptives. Here, we report the discovery of a cyanoquinoline-based series of HIPK4 inhibitors with nanomolar potency. Our lead compounds are selective for HIPK4, both within the HIPK family and across the broader kinome, establishing this scaffold as a useful starting point for probe and lead development. Unexpectedly, we found that a subset of these cyanoquinolines also perturbs HIPK4 proteostasis in a cell type-specific manner. In spermatids, these compounds induce the formation of detergent-insoluble HIPK4 aggregates and promote interactions between this kinase and the autophagy receptor Tax1-binding protein 1 (TAX1BP1). Together, our findings establish cyanoquinoline ligands as a new chemotype for probing HIPK4 biology and advancing male contraceptive discovery.

## INTRODUCTION

Approximately 85 million unintended pregnancies occur globally each year, and new contraceptive methods will be necessary to address this health challenge.^1^ Male contraception remains an underdeveloped aspect of family planning, and there is an urgent need for safe and reversible drugs that disrupt sperm development or function. Since hormonal contraceptives have been associated with mood changes, weight gain, or decreased libido, there is a growing interest in non-hormonal strategies that target testis-specific factors.^2–4^ Non-hormonal agents in development include inhibitors of retinoic acid signaling,^5, 6^ bromodomain testis-specific protein,^7^ soluble adenylyl cyclase,^8^ and serine/threonine kinase 33.^9^ However, the clinical utility of these drugs may be limited by the somatic functions of their targets or closely related isoforms.

Targeting male germ cell-specific factors offers a promising approach for achieving reversible, non-hormonal male contraception. Homeodomain-interacting protein kinase 4 (HIPK4) has been identified as a promising target, as it is selectively expressed in differentiating spermatids and essential for sperm development. Genetic ablation of *Hipk4* in male mice results in infertility without affecting overall development or physiology.^10^ In addition, heterozygous nonsense and missense *HIPK4* mutations have been identified in patients with non-obstructive azoospermia,^11^ and males homozygous for a *HIPK4* variant encoding an N-terminally truncated and destabilized protein were found to be infertile.^12^ These reproductive phenotypes stem from a critical role for HIPK4 in spermiogenesis, the final stage of sperm development that involves the differentiation of round spermatids into spermatozoa. Sperm head shaping is a key morphological transformation during this process, and it requires the coordinated movement of F-actin- and microtubule-based structures that circumscribe opposing ends of the spermatid nucleus: the acrosome-acroplaxome complex and manchette, respectively. Loss of HIPK4 function causes acroplaxome defects that lead to malformed spermatozoa incapable of fertilizing oocytes without assisted reproductive technologies.^10^

The loss-of-function phenotypes described above provide a strong biological rationale for targeting HIPK4 for male contraception. The highly druggable nature of kinases also makes HIPK4 an attractive target for therapeutic development. HIPK4 is a dual-specificity kinase and one of four members of the homeodomain-interacting protein kinase family (**Fig. 1A** and **Fig. S1**). Its classification as a HIPK family member is based solely on sequence homology within the kinase domain, as HIPK4 lacks the homeodomain-interacting region that is conserved in HIPK1-3. Moreover, HIPK4 is a cytoplasmic enzyme, whereas other HIPK isoforms localize to the nucleus where they regulate gene expression, suggesting HIPK4 may function and be regulated through mechanisms distinct from those of its better-studied paralogs.^13–15^

**Figure 1.**
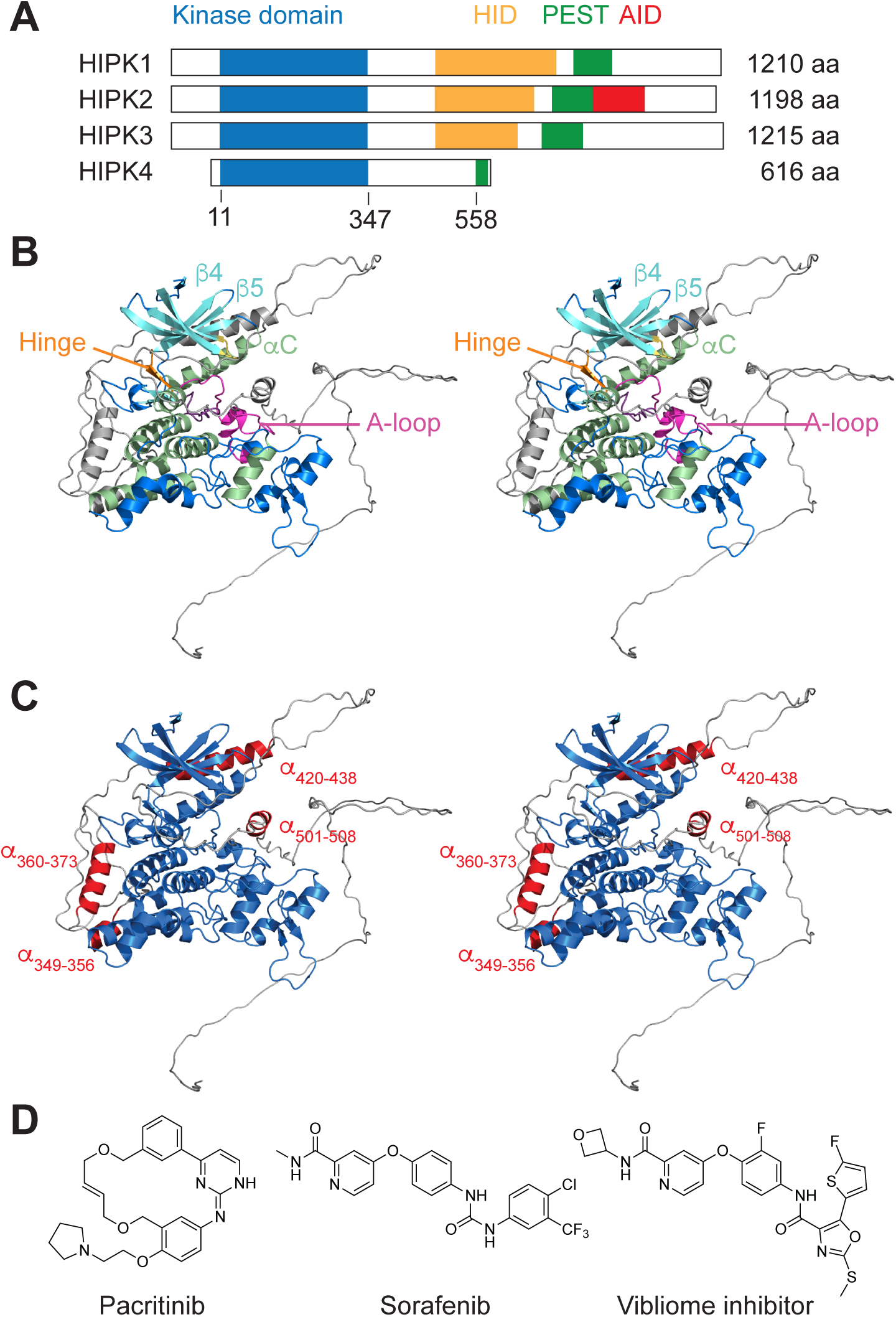
HIPK4 structure and regulation. (A) Domain architecture of the HIPK family members (HIPK1–4). Kinase domain (blue), homeobox-interacting domain (HID, orange), and PEST-like motifs (green), and the autoinhibitory domain in HIPK2 (red) are shown. (B) AlphaFold prediction of the HIPK4 structure (version 4) rendered as ribbon model and shown in wall-eyed stereoview. The canonical kinase fold with N- and C-terminal lobes is depicted in color, and other HIPK4 sequences are shown in grey. Conserved motifs in the kinase domain are color-coded to match the secondary structures and loops highlighted in **Fig. S1** (C) The AlphaFold model also predicts interactions between the N- and C-terminal lobes (shown in blue) with putative helices in C-terminal regions of the kinase (shown in red). The ribbon model is shown in wall-eyed stereoview. (D) Chemical structures of known HIPK4 inhibitors, including the non-selective ligands pacritinib and sorafenib and a HIPK4-selective ligand developed by Vibliome Therapeutics.

Although no three-dimensional structure of HIPK4 has yet been empirically determined, AlphaFold modeling predicts that HIPK4 retains the canonical kinase architecture, with N-terminal and C-terminal lobes forming a hinge-centered ATP-binding cleft (**Fig. 1B**). As in other kinases, activation is expected to involve lobe closure, inward movement of the αC helix to engage the β3 strand and form a conserved salt bridge (K40-E55), and rearrangement of the activation loop into an active DFG-in conformation. In dual-specificity kinases like HIPK4, the latter also coincides with autophosphorylation of a conserved tyrosine (Y175), which then salt-bridges with basic residues (R179 and R182) in the activation loop. Consistent with this model, the HIPK4 Y175F mutant is catalytically inactive.^16^ The AlphaFold model also predicts interactions between the HIPK4 kinase domain and C-terminal regions that most likely form helical structures (residues 349-356, 360-373, 420-438, and 501-508) (**Fig. 1C**). Together, these structural features provide an initial framework for understanding HIPK4 regulation, structure, and function.

Previously described HIPK4 antagonists include pacritinib and sorafenib, ATP-competitive ligands developed to target other kinases,^17, 18^ and a HIPK4-selective scaffold patented by Vibliome Therapeutics (**Fig. 1D**).^19^ The 4-phenoxypyridine and 2-(methylthio)-oxazole scaffold of the latter inhibitor likely has intrinsic physicochemical liabilities such as poor aqueous solubility (MW > 500 Da; cLogP > 4; tPSA > 160 Å^2^). Thus, despite strong biological and clinical interest in HIPK4, the available chemical matter for this kinase remains limited, and new ligands are needed to interrogate HIPK4 biology and to advance HIPK4-directed contraceptive discovery. HIPK4 ligands that act through novel pharmacological modalities could be especially advantageous. Achieving target specificity within the kinome is challenging due to the highly conserved ATP-binding site, and the risk of deleterious off-target side effects is a significant concern for male contraception—an elective therapy administered to healthy populations. Most kinase inhibitors reported to date are occupancy-driven agents that directly target the ATP-binding pocket or allosteric sites within the enzyme. However, ligand binding can also disrupt protein functions via several distinct mechanisms. In particular, proximity-inducing modalities such as proteolysis targeting chimeras (PROTACs) and molecular glues have been used to promote the ubiquitin ligase- and proteasome-dependent degradation of specific clinical targets.

Here we report a new series of HIPK4 ligands based on a cyanoquinoline scaffold. We identified this HIPK4-binding motif by screening a kinase-focused library enriched for chemotypes reported to target allosteric sites. Despite the intended bias toward allosteric inhibitors, the most active hits from our screen proved to be ATP-competitive compounds, including a 4-substituted cyanoquinoline derivative with nanomolar potency and HIPK4 selectivity. Our subsequent structure-activity relationship (SAR) studies have identified key features required for inhibitory potency and support a binding mode in which the C4 and cyano substituents engage the ATP-binding site. In parallel with these studies, we explored PROTAC design as a strategy for improving target specificity. Through these efforts, we unexpectedly discovered that certain cyanoquinoline derivatives can not only inhibit HIPK4 catalytic activity but also perturb HIPK4 proteostasis. In spermatids, the latter pharmacological activity is associated with HIPK4 aggregation and TAX1BP1 binding. Together, these compounds provide a starting point for further medicinal chemistry optimization and a set of chemical tools for probing the mechanisms that govern and mediate HIPK4 function.

## RESULTS

### Discovery of novel HIPK4-selective inhibitors

To identify new HIPK4 inhibitors, we pursued a high-throughput screening campaign. We first developed methods for producing recombinant HIPK4 protein. Our efforts to bacterially express HIPK4 or its kinase domain alone (residues 1-347) yielded only insoluble, inactive proteins, consistent with a previous report.^13^ However, we were able to successfully express full-length HIPK4 and a construct lacking the C-terminal PEST-like motif (HIPK4 ΔPEST; residues 1-557) as soluble proteins in insect (Sf9) and mammalian (Expi293) cells, respectively (**Fig. 2A** and **Fig. S2**). Both forms were active in ADP-Glo^20^ kinase assays. Since we were able to obtain HIPK4 ΔPEST in milligram quantities, we used the recombinant protein to screen a commercial 26,000-compound library designed to target kinases with allosteric pockets (**Fig. 2B** and **Fig. S3**). Our screen utilized both protein thermal shift (PTS) and activity-based (ADP-Glo) assays, with the latter format utilizing a high ATP concentration (100 µM) to favor non-ATP-competitive or potent ATP-competitive inhibitors. Hit validation was then performed with fresh powders, commercial glutathione S-transferase (GST)-tagged full-length HIPK4, and the ADP-Glo assay. These studies uncovered several ATP-competitive inhibitors across several scaffolds and two non-ATP-competitive benzoisoxazoles (**Fig. S3**). Contrary to our initial expectation that non-ATP-competitive ligands would exhibit enhanced HIPK4 selectivity, both benzoisoxazoles inhibited all HIPK family members with comparable potency. Among the ATP-competitive ligands, a cyanoquinoline derivative had the greatest potency and HIPK4 selectivity (**Fig. 2C-D**, compound **1**), and we prioritized this compound for further study.

**Figure 2.**
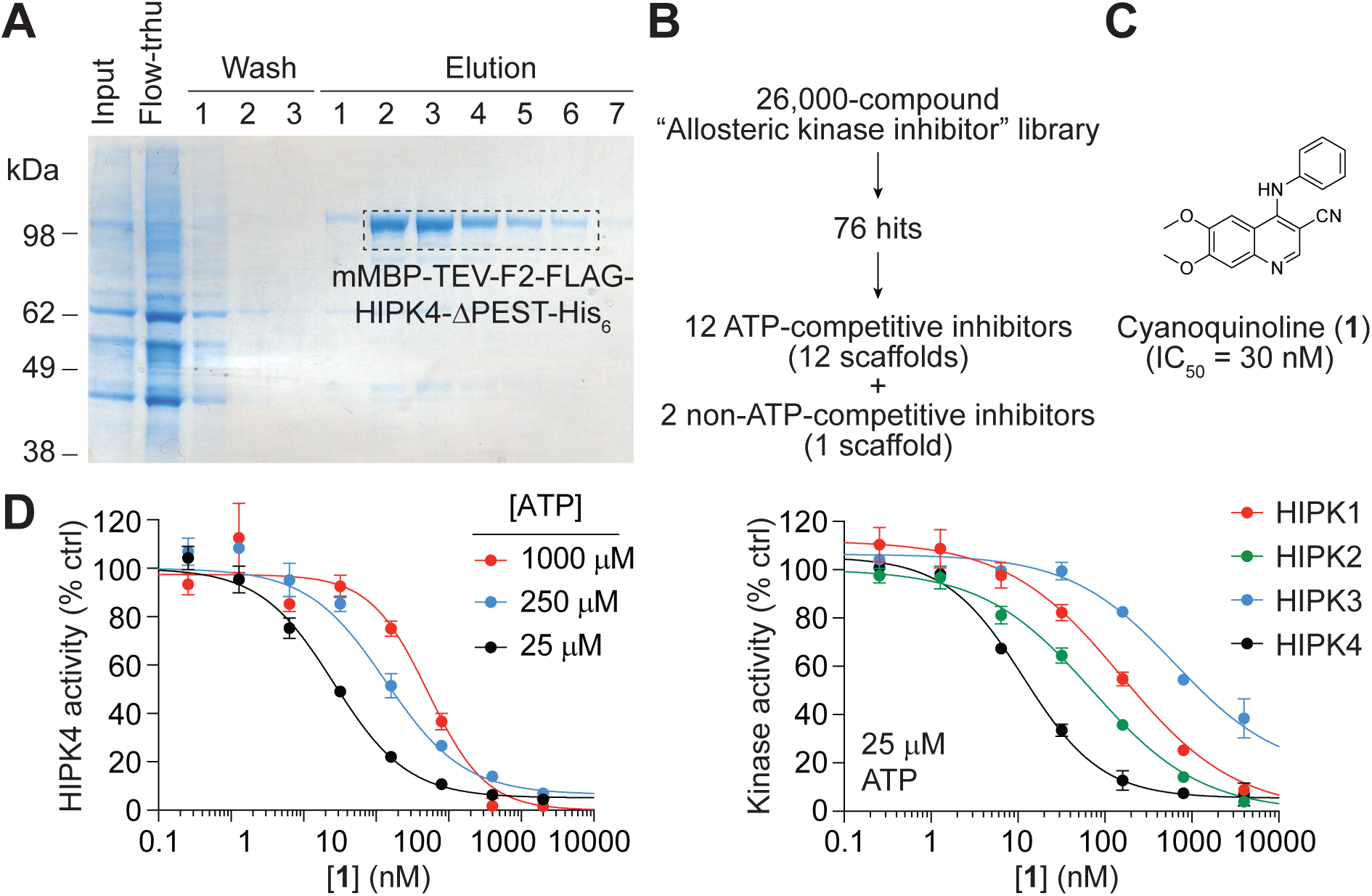
Discovery of an ATP-competitive, HIPK4-selective inhibitor. (A) Expression and purification of a HIPK4 ΔPEST construct in Expi293 cells. HIPK4 ΔPEST was expressed as a proteolytically cleavable maltose binding protein (mMBP) fusion and with N-terminal FLAG and C-terminal His_6_ tags. The final construct used in our high-throughput screen for HIPK4 inhibitors was obtained by thrombin cleavage and His_6_ affinity purification. (B) Summary of the high-throughput screen, which yielded several ATP-competitive HIPK4 ligands and a non-ATP-competitive scaffold. (C) A cyanoquinoline-based inhibitor identified in our screen. (D) ADP-Glo assays using myelin basic protein as a universal substrate, demonstrating the ATP-competitive activity of cyanoquinoline **1** and its selectivity for HIPK4 among the HIPK subfamily. Data are the average of two biological replicates ± s.e.m.

### SAR landscape of cyanoquinoline-mediated HIPK4 inhibition

To lay the groundwork for hit-to-lead optimization, we initiated SAR studies of the cyanoquinoline scaffold. We first tested ten commercially available analogs of the screening hit (**Fig. S4**; compounds **2**-**11**) and found that HIPK4 can tolerate modifications at the C6 and C7 positions of the pharmacophore, whereas *para* substituents on the C4 aniline ring significantly reduce inhibitor activity. This binding mode is consistent with the structures of several kinases bound to quinoline or quinazoline scaffolds, including that of the Abelson tyrosine kinase (ABL)-bosutinib complex (**Fig. 3**).^21–23^

**Figure 3.**
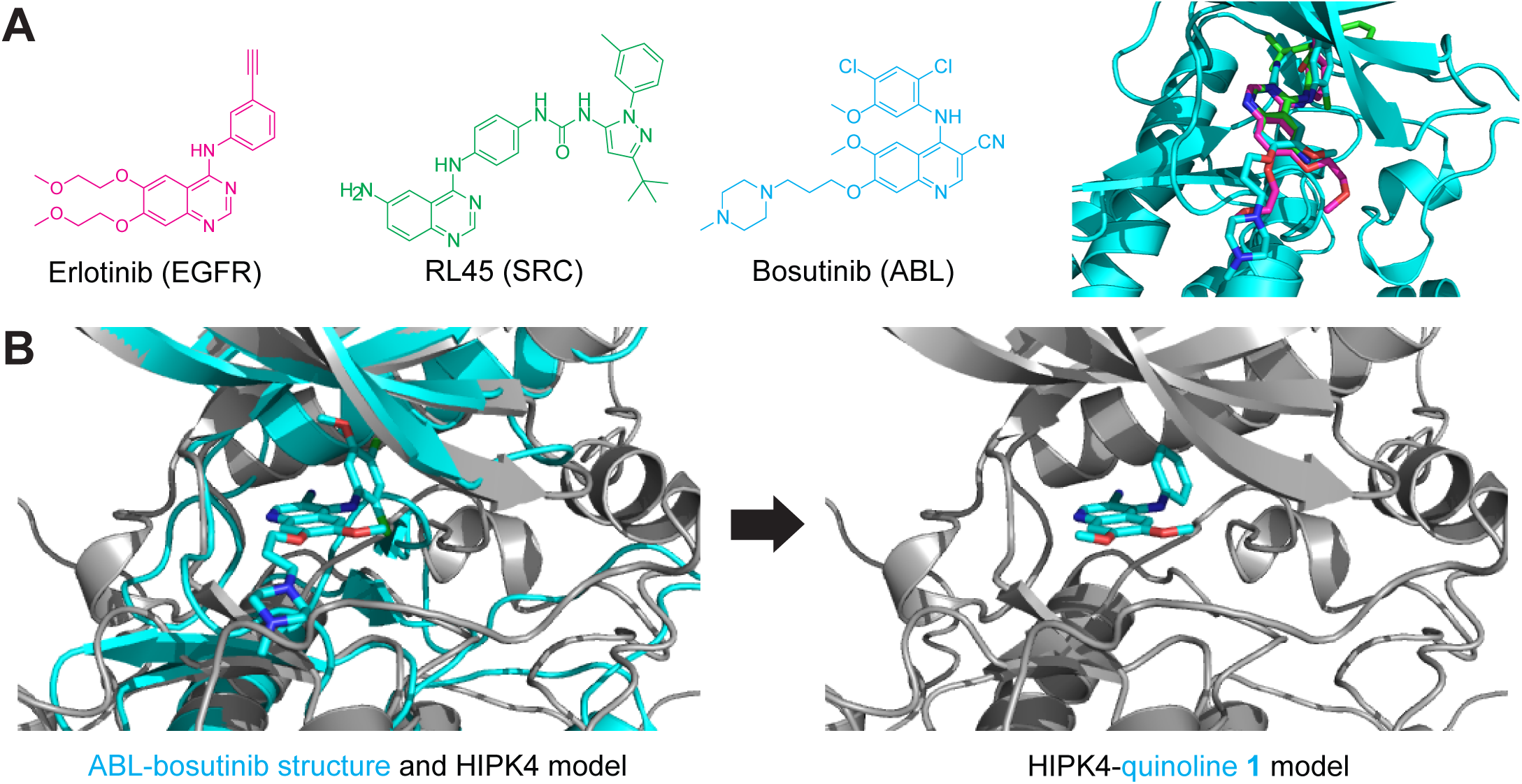
A model for HIPK4-cyanoquinoline binding. (A) Chemical structure of kinase inhibitors with quinoline/quinazoline scaffolds and an overlay of the ABL-bosutinib complex with erlotinib and RL45 in their kinase-binding orientations (PDB IDs: 3UE4, 3F3V, and 1M17). (B) Model of the HIPK4-quinoline **1** complex, which was generated by aligning the AlphaFold model of HIPK4 (version 4) with the ABL-bosutinib structure and replacing bosutinib with **1.**

We next established modular synthetic routes to access noncommercial quinoline analogs (**Table 1**, **Scheme S1**, and **Supplementary File 1**). The chemical series was assembled from chlorinated quinoline intermediates, using a retrosynthetic strategy that leverages late-stage amine installation and phenol functionalization. In the simplest route, direct displacement of the 4-chloro substituent with the corresponding amines (R_3_-NH_2_) in ethanol at 100 °C provided the corresponding 4-aminoquinoline derivatives **23**-**42** and **47**. To access analogs bearing diversified alkoxy substituents, a benzyloxy-protected quinoline intermediate was first subjected to the same amination conditions, after which debenzylation with thioanisole in trifluoroacetic acid (TFA) at 100 °C furnished the corresponding phenol intermediate. Subsequent *O*-alkylation with the desired alkyl bromides (R_1_-Br) in the presence of cesium carbonate (Cs_2_CO_3_) at 100 °C then afforded compounds **12**-**17** and **43**-**44**. A comparable strategy was used to prepare a second subset of analogs from an isopropoxy-protected quinoline precursor. Following amine substitution in ethanol at 100 °C, deprotection with aluminum chloride (AlCl_3_) in dichloromethane (DCM) yielded the corresponding phenol, which was then functionalized with alkyl bromides (R_2_-Br) in the presence of Cs_2_CO_3_ in dimethylformamide (DMF) at 100 °C to afford compounds **18**-**22** and **45**-**46**.

**Table 1.**
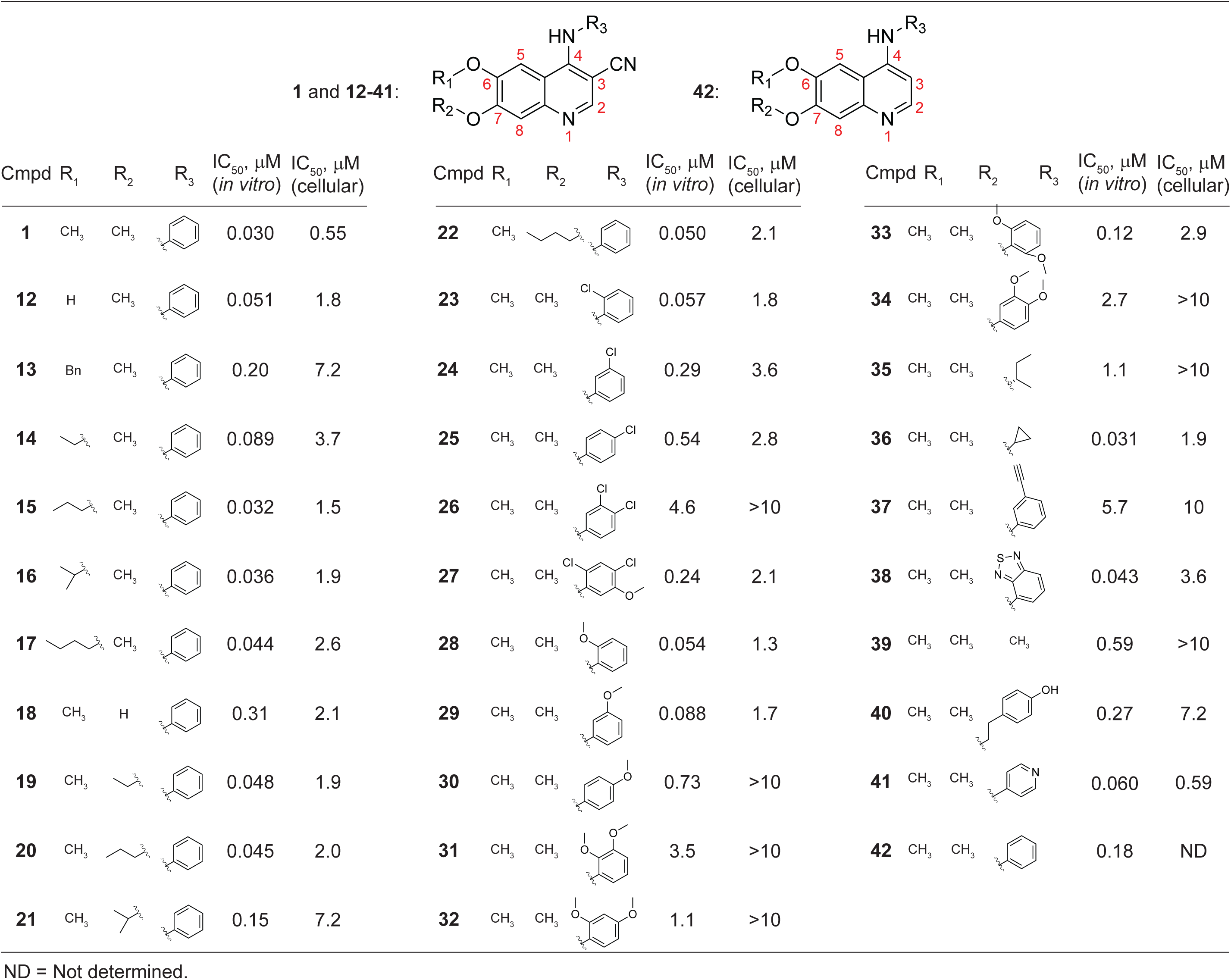
HIPK4 inhibition by cyanoquinolines. *In vitro* and cellular inhibitory activities of cyanoquinolines with varying linker attachment sites and aniline substituents. The numbering scheme for the quinoline scaffold is also shown. Biochemical kinase assays were conducted with 25 µM ATP and quantified using the ADP-Glo system, and cellular HIPK4 engagement was assessed using the NanoBRET platform.

| 1 and 12-41: |  |  |  |  |  | 42: |  |  |  |  |  |  |  |  |  |  |  |
| --- | --- | --- | --- | --- | --- | --- | --- | --- | --- | --- | --- | --- | --- | --- | --- | --- | --- |
| Cmpd | R<sub>1</sub> | R<sub>2</sub> | R<sub>3</sub> | IC<sub>50</sub>, μM<br>(*in vitro*) | IC<sub>50</sub>, μM<br>(cellular) | Cmpd | R<sub>1</sub> | R<sub>2</sub> | R<sub>3</sub> | IC<sub>50</sub>, μM<br>(*in vitro*) | IC<sub>50</sub>, μM<br>(cellular) | Cmpd | R<sub>1</sub> | R<sub>2</sub> | R<sub>3</sub> | IC<sub>50</sub>, μM<br>(*in vitro*) | IC<sub>50</sub>, μM<br>(cellular) |
| 1 | CH<sub>3</sub> | CH<sub>3</sub> |  | 0.030 | 0.55 | 22 | CH<sub>3</sub> |  |  | 0.050 | 2.1 | 33 | CH<sub>3</sub> | CH<sub>3</sub> |  | 0.12 | 2.9 |
| 12 | H | CH<sub>3</sub> |  | 0.051 | 1.8 | 23 | CH<sub>3</sub> | CH<sub>3</sub> |  | 0.057 | 1.8 | 34 | CH<sub>3</sub> | CH<sub>3</sub> |  | 2.7 | >10 |
| 13 | Bn | CH<sub>3</sub> |  | 0.20 | 7.2 | 24 | CH<sub>3</sub> | CH<sub>3</sub> |  | 0.29 | 3.6 | 35 | CH<sub>3</sub> | CH<sub>3</sub> |  | 1.1 | >10 |
| 14 |  | CH<sub>3</sub> |  | 0.089 | 3.7 | 25 | CH<sub>3</sub> | CH<sub>3</sub> |  | 0.54 | 2.8 | 36 | CH<sub>3</sub> | CH<sub>3</sub> |  | 0.031 | 1.9 |
| 15 |  | CH<sub>3</sub> |  | 0.032 | 1.5 | 26 | CH<sub>3</sub> | CH<sub>3</sub> |  | 4.6 | >10 | 37 | CH<sub>3</sub> | CH<sub>3</sub> |  | 5.7 | 10 |
| 16 |  | CH<sub>3</sub> |  | 0.036 | 1.9 | 27 | CH<sub>3</sub> | CH<sub>3</sub> |  | 0.24 | 2.1 | 38 | CH<sub>3</sub> | CH<sub>3</sub> |  | 0.043 | 3.6 |
| 17 |  | CH<sub>3</sub> |  | 0.044 | 2.6 | 28 | CH<sub>3</sub> | CH<sub>3</sub> |  | 0.054 | 1.3 | 39 | CH<sub>3</sub> | CH<sub>3</sub> | CH<sub>3</sub> | 0.59 | >10 |
| 18 | CH<sub>3</sub> | H |  | 0.31 | 2.1 | 29 | CH<sub>3</sub> | CH<sub>3</sub> |  | 0.088 | 1.7 | 40 | CH<sub>3</sub> | CH<sub>3</sub> |  | 0.27 | 7.2 |
| 19 | CH<sub>3</sub> |  |  | 0.048 | 1.9 | 30 | CH<sub>3</sub> | CH<sub>3</sub> |  | 0.73 | >10 | 41 | CH<sub>3</sub> | CH<sub>3</sub> |  | 0.060 | 0.59 |
| 20 | CH<sub>3</sub> |  |  | 0.045 | 2.0 | 31 | CH<sub>3</sub> | CH<sub>3</sub> |  | 3.5 | >10 | 42 | CH<sub>3</sub> | CH<sub>3</sub> |  | 0.18 | ND |
| 21 | CH<sub>3</sub> |  |  | 0.15 | 7.2 | 32 | CH<sub>3</sub> | CH<sub>3</sub> |  | 1.1 | >10 |  |  |  |  |  |  |

ND = Not determined.

We evaluated each of the quinolines for their ability to inhibit commercial GST-HIPK4 in the ADP-Glo assay (**Table 1** and **Supplementary File 1**), and SAR results for this more structurally diverse series broadly corroborated our findings with the commercial derivatives. For example, we compared the activities of several analogs with a C4 aniline substituent and varying groups at the C6 and C7 positions. A wide range of alkyl ethers were tolerated at the latter sites, suggesting that this region in the scaffold is not a major determinant of HIPK4 inhibition. In contrast, modifications of the C4 substituent had substantially greater impact on cyanoquinoline activity, with benzothiadiazole and pyridine analogs (compounds **38** and **41**) exhibiting inhibitory potencies comparable to that of the screening hit. A direct comparison between compounds **1** and **42** revealed loss of the C3-cyano group resulting in an approximately 5-fold reduction in inhibitory activity against HIPK4. The corresponding cyano group in bosutinib participates in a water-mediated hydrogen bond network in the kinase active site,^21^ and our ligands may engage in analogous interactions.

### HIPK4-cyanoquinoline engagement in cells

To directly measure HIPK4-ligand binding in live cells, we implemented a bioluminescence resonance energy transfer (BRET) assay (**Fig. 4A**).^24^ In this protocol, HIPK4 is N-terminally fused to NanoLuc luciferase, and the chimera is overexpressed in HEK293T cells, a human embryonic kidney-derived line. The cells are then treated with a NanoLuc substrate and an ATP-competitive broad-spectrum kinase inhibitor covalently linked to an energy transfer probe. Binding of the cell-permeable probe to HIPK4 generates a BRET signal that can be measured using a luminometer equipped with spectral filters. Competition experiments can then be used to quantify interactions between the HIPK4 active site and compounds of interest.

**Figure 4.**
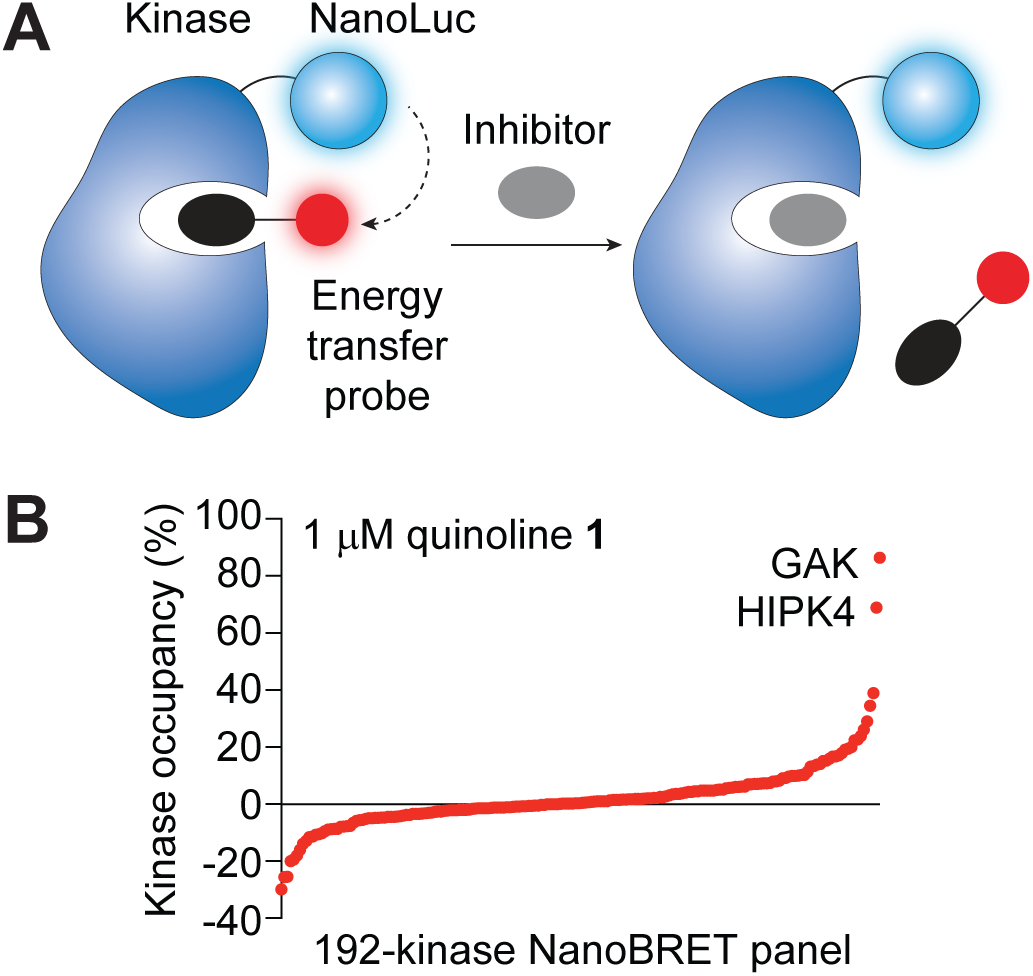
Cellular HIPK4 engagement and target selectivity of a cyanoquinoline ligand. (A) Schematic representation of the NanoBRET assay for kinase-inhibitor binding in live cells. (B) Profiling data for cyanoquinoline **1** in a 192-kinase NanoBRET platform.

Our NanoBRET analysis confirmed that cyanoquinolines effectively engaged HIPK4 in cells, albeit with a reduction in potency that could be due to the millimolar ATP concentrations in cells or limited membrane permeability (**Table 1** and **Supplementary File 1**). For example, the screening hit **1** had an IC_50_ value of 0.55 µM in the NanoBRET assay, which is nearly 20-fold higher than its IC_50_ value in the ADP-Glo assay. Overall, the cyanoquinoline analogs we synthesized and tested had NanoBRET activities that correlated with their *in vitro* efficacy, but none achieved low nanomolar potencies in this assay. Thus, further modifications to the quinoline scaffold will be necessary to optimize cell penetrance.

Nevertheless, the NanoBRET assay also afforded us an opportunity to more comprehensively assess the target specificity of this cyanoquinoline chemotype. We profiled cyanoquinoline **1** in a NanoBRET-based platform that included 192 kinases distributed throughout the kinome (**Fig. 4B**). When tested at a 1-µM concentration, the inhibitor only engaged HIPK4 and cyclin G-associated kinase (GAK) to an appreciable extent. Since GAK is well-known as a kinase binding many diverse inhibitors,^25, 26^ these results indicated that the cyanoquinoline scaffold has high cellular HIPK4 specificity.

### Development of HIPK4 degraders

To enhance the selectivity of our cyanoquinoline-based HIPK4 ligands, which likely act as type I inhibitors, we explored alternative pharmacological modalities. In particular, bifunctional molecules that induce targeted protein degradation through E3 ligase recruitment have broadened the pharmaceutical landscape and unlocked new avenues of target modulation. Small-molecule degraders offer several advantages in comparison to inhibitors, including complete loss of protein function, efficacy at substoichiometric doses, and greater specificity due to the structural requirements for functional ternary complexes.^27, 28^

We initially sought to develop HIPK4 PROTACs and evaluated potential linker exit vectors by installing polyethyleneglycol (PEG)- and alkyl chain-based linkers at multiple exit vectors on the cyanoquinoline scaffold. We tested the resulting compounds in the ADP-Glo assay, and as expected from the predicted cyanoquinoline binding mode and our SAR studies, adding these linkers to the C6 or C7 positions only moderately attenuated inhibitor potency (**Fig. 5A**, compounds **43-46**). In comparison, a derivative containing an alkyl linker-functionalized C4 aniline was 10 – 100-fold less active (compound **47**). We therefore prioritized the C6 and C7 positions as solvent-facing sites that could be utilized for PROTAC design. We also established a chemiluminescence-based HIPK4 degradation assay in live cells using the split-luciferase HiBiT/LgBiT system (**Fig. 5B**).^29^ In this assay format, HIPK4 is fused in-frame to the 11-amino-acid HiBiT tag and the protein is ectopically expressed in an LgBiT-expressing HEK293T reporter cell line, which also has endogenous cereblon (CRBN). Reconstitution of the HiBiT tag with LgBiT generates a functional luciferase in a HIPK4-concentration dependent manner, thereby enabling real-time monitoring of chemically induced protein loss. For our studies, we created an HEK293T line that stably expresses HiBiT-HIPK4 and LgBiT at levels that yield a robust chemiluminescent signal.

**Figure 5.**
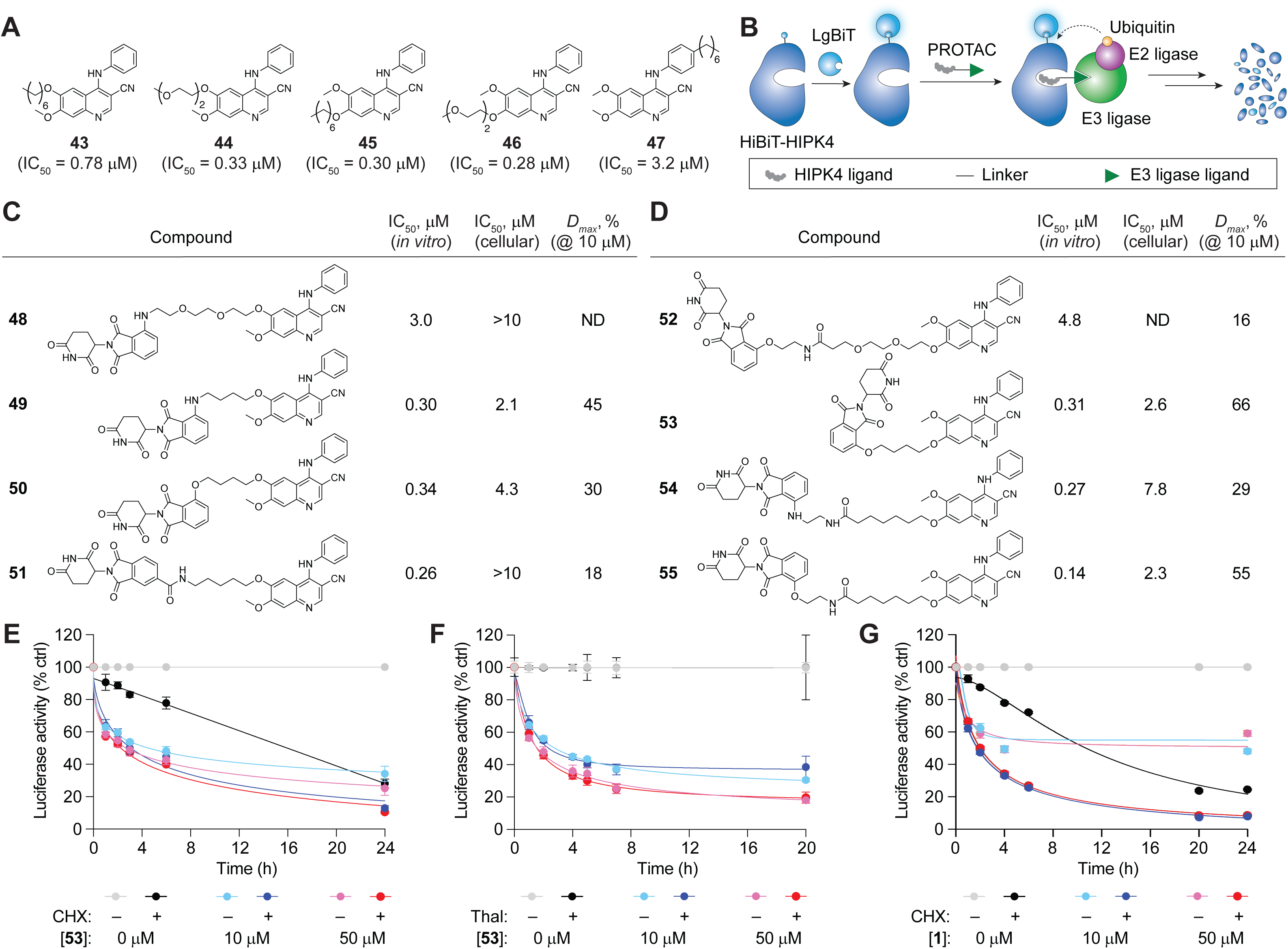
Cyanoquinoline can induce HIPK4 degradation in HEK293T cells through a non-PROTAC mechanism. (A) Inhibitory activities of cyanoquinolines with varying linker attachment sites (ADP-Glo assay with 25 µM ATP). Data are the average of three biological replicates. (B) Schematic representation of PROTAC-mediated degradation of HIPK4 as monitored in real time with the HiBiT/LgBiT assay platform. (C-D) ADP-Glo, NanoBRET, and HiBiT assay activities for cyanoquinoline-based PROTACs with varying CRBN ligand and linker structures, using the C6 (C) and C7 (D) sites as attachment points. ADP-Glo assay data are the average of three biological replicates, and NanoBRET and HiBiT assay data are the average of two biological replicates. (E) Degrader activity profiles of cyanoquinoline **53** in the HiBiT-HIPK4 assay. The cells were cultured in the absence or presence of cycloheximide (CHX; 10 µg/mL) to discern the compounds’ effects on HIPK4 synthesis versus degradation. (F) Degrader activity profiles of **53** in the absence or presence of 20 µM thalidomide (Thal). (G) Degrader activity profiles of cyanoquinoline **1** in the HiBiT-HIPK4 assay, conducted in the absence or presence of CHX. HiBiT-HIPK4 assay data are shown as the average of two biological replicates ± s.e.m.

Having established a strategy for PROTAC design and testing, we prepared a focused set of cyanoquinoline-based PROTACs incorporating CRBN-recruiting thalidomide derivatives at the C6 or C7 positions, using either the PEG- or alkyl chain-based linkers (**Fig. 5C-D** and **Scheme S2**). The compounds were synthesized from common C6- or C7-hydroxylated cyanoquinoline intermediates through *O*-alkylation, followed by deprotection of the terminal linker handle and coupling to appropriately functionalized thalidomide derivatives by nucleophilic substitution or HATU-mediated amide bond formation. We then evaluated the candidate PROTACs in our ADP-Glo and NanoBRET assays (**Supplementary File 1**). While the PEG-linked analogs (**48** and **52**) inhibited HIPK4 *in vitro* with 100-fold less potency and had no detectable HIPK4 engagement in cells using BRET assays, nearly all of the alkyl chain-linked derivatives (**49-51** and **53-55**) engaged with HIPK4 with EC_50_ values that were within 10-fold of the screening hit.

Several of our candidate HIPK4 PROTACs induced HiBiT-HIPK4 degradation in cells, and **53** and **55** emerged as the most potent compounds, achieving maximal degradation (*Dmax*) levels of 66% and 55%, respectively, at 10 µM PROTAC concentration and 24 hours incubation time. We therefore prioritized **53** for further mechanistic characterization. To distinguish between enhanced protein clearance from the suppression of new protein synthesis, we retested **53** in the presence or absence of the protein translation inhibitor cycloheximide (CHX) (**Fig. 5E**). Treating HiBiT-HIPK4- and LgBiT-expressing HEK293T cells with CHX resulted in a gradual loss of luciferase activity, consistent with the intrinsic turnover of the fusion protein. Cyanoquinoline **53** accelerated HiBiT-HIPK4 loss under both CHX treatment conditions, indicating that the decrease in c signal primarily reflects an increase in HiBiT-HIPK4 degradation rather than protein synthesis blockade.

We next sought to confirm that **53** induces HIPK4 degradation through CRBN recruitment. Because the degrader comprises a glutarimide-containing CRBN ligand, we anticipated that excess thalidomide would competitively inhibit CRBN recruitment and attenuate HIPK4 degradation.^30^ However, addition of thalidomide had little effect on the degradation kinetics of HiBiT-HIPK4 induced by **53**, even at concentrations expected to saturate CRBN (**Fig. 5F**). Taken together, these data indicated that although **53** was designed as a CRBN-recruiting PROTAC, it can reduce HIPK4 levels in a CRBN-independent manner.

Consistent with this idea, we made the surprising observation that some cyanoquinolines lacking a CRBN ligand also induced HIPK4 degradation in the HiBiT assay (**Fig. 5G, Fig. S5**, and **Supplementary File 1**). Several analogs achieved *Dmax* values > 90% (compounds **20** and **23**), and as with cyanoquinoline **53**, their degrader activity was additive to the effects of CHX treatment. We also investigated HIPK4 ligands with macrocyclic cyanoquinoline scaffolds that were independently discovered by the Hanke group, represented by compounds **56**-**58**.^31^ All three macrocycles were potent HIPK4 inhibitors and effectively degraded HiBiT-HIPK4. Thus, the cyanoquinoline scaffold can promote HIPK4 degradation through a non-PROTAC mechanism.

### Mechanistic studies of HIPK4 degrader action

The unanticipated CRBN-independent function of our HIPK4 degraders spurred us to further explore their mechanisms of action. We first compared the compounds’ potencies in the *in vitro* HIPK4 ADP-Glo assays with their *Dmax* values in the HEK293T cell-based HiBiT-HIPK4 platform, finding no correlation between the two activities (**Fig. 6A** and **Supplementary File 1**). For instance, compound **33** was a potent HIPK4 inhibitor with negligible HIPK4 degradation in the HiBiT system. These results indicated that binding affinity alone was insufficient to predict degradative activity, mirroring the SAR landscapes previously defined for other small-molecule degraders.^32^ Rather, it appears that certain quinoline substituents more effectively converted HIPK4 to a degradation-prone state.

**Figure 6.**
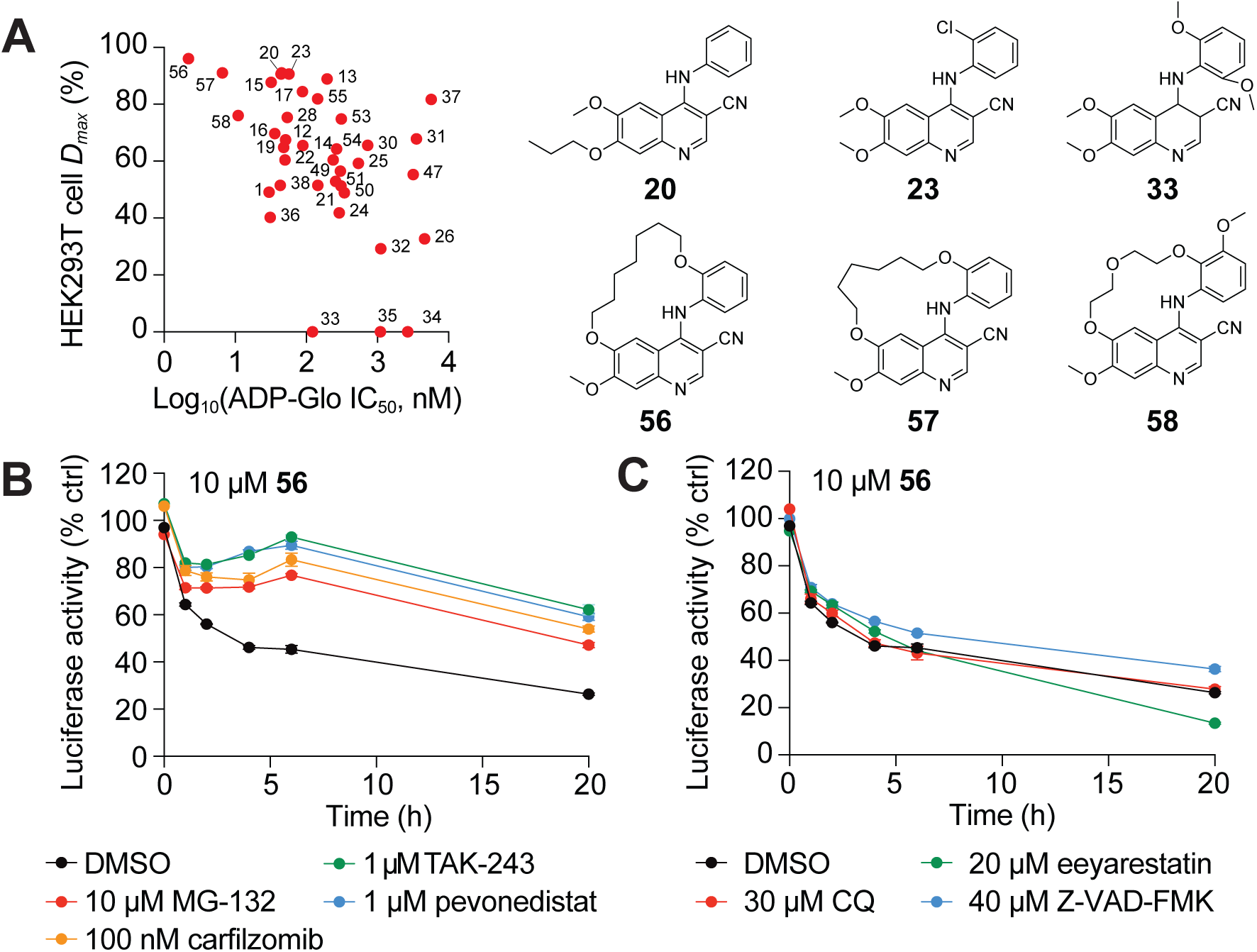
Cyanoquinoline-induced HIPK4 degradation requires the ubiquitin-proteasome system. (A) Cyanoquinoline-mediated HIPK4 inhibition (ADP-Glo assay IC_50_ values) and degradation (HiBiT-HIPK4 assay *Dmax* values; 50 µM compound) are not correlated. (B-C) Cyanoquinoline-mediated HIPK4 degradation in HEK293T cells is suppressed by UPS inhibitors (B) but not antagonists of the autophagy-lysosomal pathway (chloroquine, CQ), ERAD (eeyarestatin), or non-apoptotic caspases (Z-VAD-FEMK) (C).

To better understand the molecular and cellular bases for HIPK4 degrader function, we pharmacologically interrogated key proteostasis mechanisms, including ubiquitin-proteasome system (UPS), endoplasmic reticulum-associated degradation (ERAD), caspase-dependent proteolysis, and the autophagy-lysosomal pathway. Using cyanoquinoline **56** as a representative degrader in the HiBiT-HIPK4 assay, we found that its effects on the HiBiT signal were strongly suppressed by inhibitors of the proteasome (MG-132 and carfilzomib), ubiquitin-activating enzyme E1 (TAK-243), and NEDD8 (neural precursor cell expressed, developmental down-regulated protein 8)-activating enzyme (pevonedistat) (**Fig. 6B**). In contrast, perturbations of lysosomal function (chloroquine), ERAD (eeyarestatin), or non-apoptotic caspase activity (Z-VAD-FMK) produced little or no rescue (**Fig. 6C**). These data indicated that in HEK293T cells, cyanoquinoline-induced HIPK4 degradation proceeds primarily through a UPS-dependent process that likely involves a cullin E3 ligase due its neddylation dependence.

### Cyanoquinolines can disrupt HIPK4 proteostasis in spermatids

We next examined whether the degradative effects of cyanoquinolines we observed in HEK293T cells extended to spermatids, the physiologically relevant cell type for this kinase. Immunoblot analyses of detergent lysates obtained from dissociated mouse testes showed a clear concentration-dependent reduction in endogenous HIPK4 levels upon treatment with compounds **17**, **25**, **53**, and **56** for 16 hours (**Fig. 7A** and **Fig. S5**), with EC_50_ values ranging from 1.5 to 11 µM. Cyanoquinoline **53** contains a CRBN ligand, and corroborating our results in HEK293T cells, we confirmed that it acts through a non-PROTAC mechanism in spermatids (**Fig. S6**) However, in contrast to our HiBiT assay results, cyanoquinoline-induced HIPK4 loss in spermatids was not rescued by inhibitors of the UPS system (**Fig. 7B**), indicating the cyanoquinolines alter HIPK4 proteostasis in HEK293T cells and the germ cells through distinct mechanisms. We subsequently noted that the lower levels of HIPK4 in lysates obtained from cyanoquinoline-treated spermatids coincided with a redistribution of the kinase to the detergent-insoluble fraction (**Fig. 7C**). To better understand the molecular and cellular basis for this effect, we next used immunofluorescence microscopy to follow the subcellular distributions of HIPK4 in compound-treated spermatids. HIPK4 protein is normally distributed through the spermatid cytoplasm with a few isolated puncta, and treatment with **25** markedly increased the number of HIPK4-positive puncta after 5-hour incubation (**Fig. 7D**). Collectively, these observations indicated that certain cyanoquinoline ligands for HIPK4 drive the partitioning of this kinase into detergent-insoluble aggregates.

**Figure 7.**
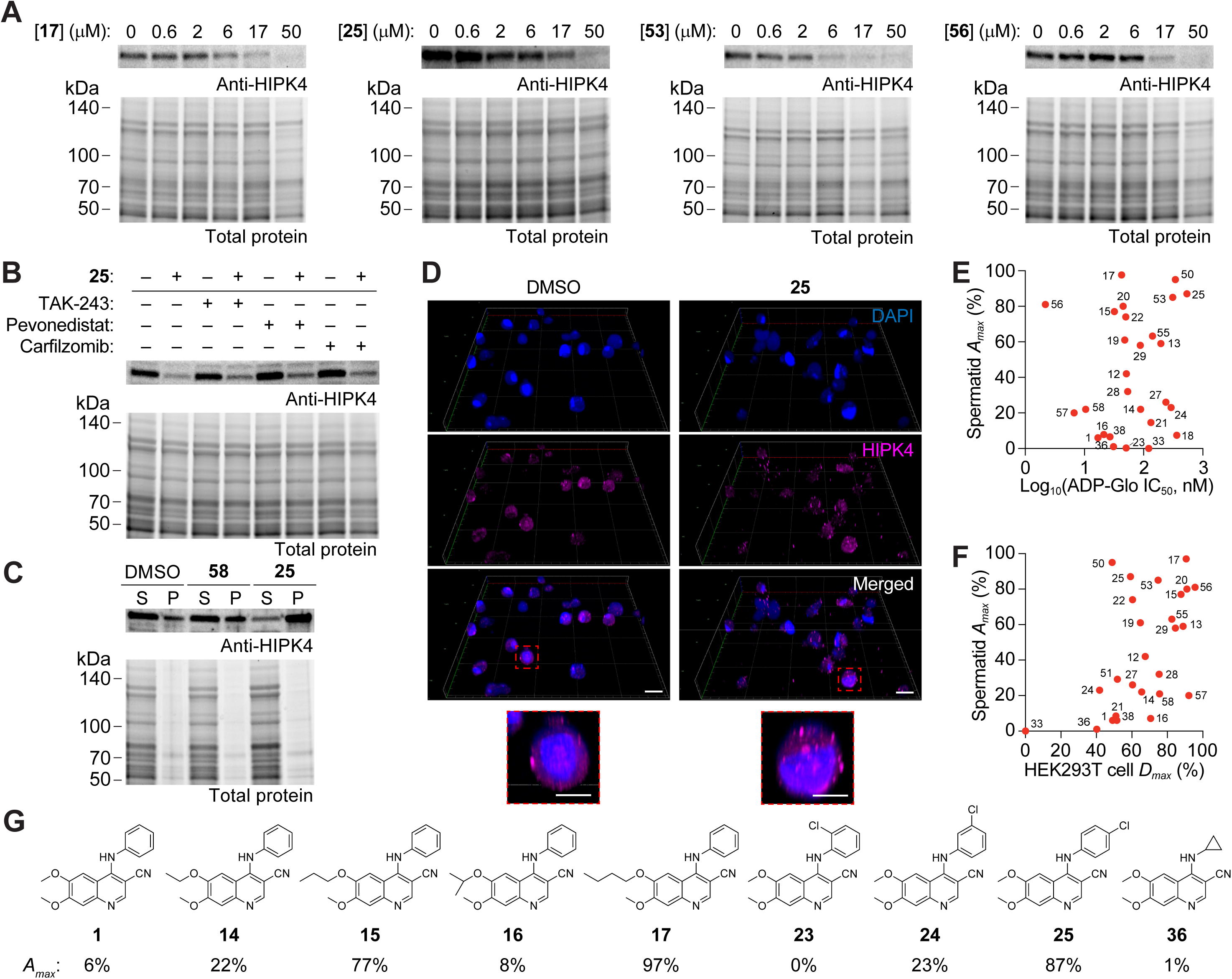
Cyanoquinolines can induce HIPK4 aggregation in spermatids. (**A**) Cyanoquinoline-induced loss of HIPK4 in spermatids, as determined by western blot analyses of detergent-solubilized cell lysates. Compounds **17**, **25**, **53** and **56** have EC_50_ values of 9.1, 4.5, 1.5, and 11.4 µM, respectively. (B) Cyanoquinoline **25**-induced HIPK4 loss in spermatids cannot be rescued by UPS inhibitors. (C) Spermatid treatment with cyanoquinoline **25** but not **58** induces the conversion of HIPK4 to a detergent-insoluble form, as determined by western blot analyses of the spermatid lysate supernatant (S) and centrifugal pellet (P). (D) Compound **25** also induces the formation of HIPK4-positive puncta in spermatids within 5 h of treatment (25 μM dose). Scale bars: 10 µm (full image) and 5 µm (inset). (E) Cyanoquinoline-mediated HIPK4 inhibition (ADP-Glo assay IC_50_ values) and aggregation (*Amax* values in spermatid immunoblotting experiments; 50 µM compound) are not correlated. (F) Cyanoquinoline-induced HIPK4 degradation in HEK293T cells (*Dmax* values; 50 µM compound) and HIPK4 aggregation in spermatids (*Amax* values; 50 µM compound) are not correlated. Data are the average of two biological replicates. (G) Representative cyanoquinolines that illustrate how substituents at the C4, C6, and C7 positions influence *Amax* values.

To quantify how cyanoquinolines disrupted HIPK4 proteostasis in spermatids, we defined the term *Amax* as the percentage of the targeted protein that is converted into detergent-insoluble aggregates at a given time point. Similar to *Dmax* values assessed by our HiBiT assays that did not correlate with *in vitro* IC_50_ values, we observed no correlation between *Amax* values in spermatids and inhibitory potency (**Fig. 7E** and **Supplementary File 1**). Moreover, *Dmax* values in the HEK293T-based HiBiT assay were not reliably predictive of *Amax* values in spermatids (**Fig. 7F** and **Supplementary File 1**). For instance, only one of the macrocyclic cyanoquinolines we tested induced HIPK4 aggregation, even though all three compounds tested were potent degraders of HIPK4 when the kinase was heterologously expressed in HEK293T cells. This divergence provided further evidence that the specific effects of cyanoquinolines on HIPK4 proteostasis depend on the cellular environment in which the kinase resides.

Nevertheless, there are shared structural features among the cyanoquinolines which induced the greatest HIPK4 aggregation in spermatids. Compounds with linear alkoxy groups at either the C6 or C7 position of the quinoline scaffold generally exhibited high *Amax* values, including analogs bearing *n*-propoxy or *n*-butoxy substituents (**Fig. 7G** and **Supplementary File 1**). In contrast, the branched isopropoxy analog **16** was markedly less active. Aniline ring substituents also had a strong effect, with *para*-substitution, particularly 4-chloro, favoring HIPK4 aggregation and *ortho*-substitutions often suppressing this activity. Moreover, the cyanoquinolines in our series with the most sterically compact C4 groups, such as **1** and the cyclopropyl derivative **36**, exhibited the lowest *Amax* values despite having the greatest *in vitro* inhibitory activities. Collectively, these results suggest that cyanoquinolines can disrupt HIPK4 proteostasis in spermatids by stabilizing a protein conformation that is prone to aggregation.

### Cyanoquinoline-induced HIPK4 aggregation requires other spermatid proteins

We concluded our studies by investigating the mechanisms by which cyanoquinolines can induce HIPK4 aggregation in spermatids. To assess whether HIPK4-cyanoquinoline complexes can self-associate, we used mass photometry to monitor the oligomerization state of recombinant His_6_-tagged full-length HIPK4. The full-length protein exhibited light scattering properties consistent with a predominantly monomeric state, and incubating the kinase with 5 µM **25** did not alter its apparent size (**Fig. 8A**). These observations indicated that HIPK4-cyanoquinoline complexes cannot self-associate in isolation but rather require cellular cofactors to aggregate.

**Figure 8.**
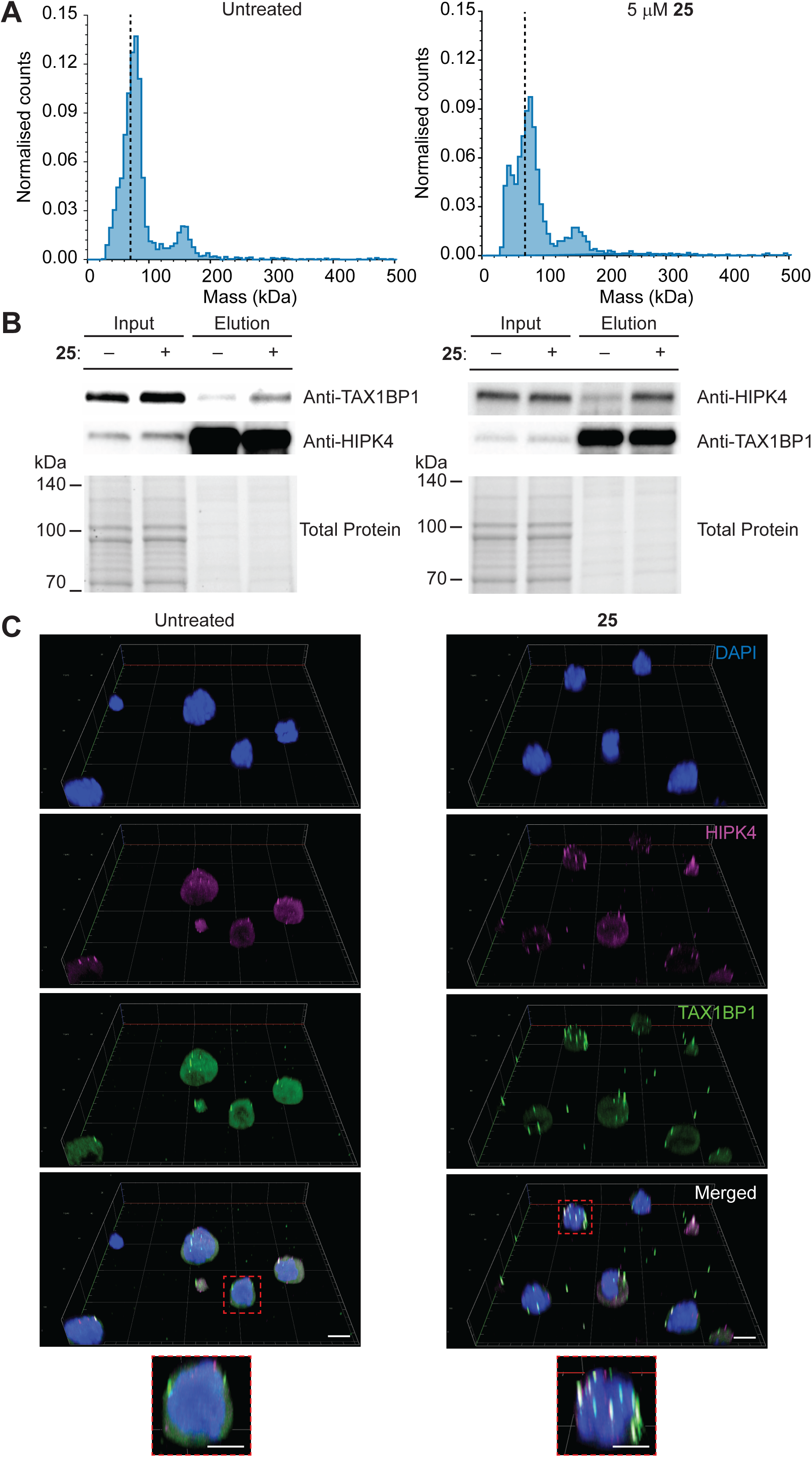
Cyanoquinolines can promote HIPK4-TAX1BP1 binding. (A) Mass photometry histograms of recombinant, full-length His_6_-HIPK4 protein treated with 5 µM cyanoquinoline **25**. Normalized landing counts are plotted as a function of molecular mass, and the dashed vertical lines indicate the predicted mass of monomeric HIPK4 (69.5 kDa). (B) Spermatid treatment with **25** promotes the co-immunoprecipitation of HIPK4 and TAX1BP1. The spermatids were treated with 50 µM **25** or DMSO alone for 5 h prior to lysis and pulldown experiments with anti-HIPK4 or anti-TAX1BP1 antibodies. (C) Representative 3D confocal fluorescence images of spermatids treated with **25** or DMSO alone for 5 h. The germ cells were fixed and then stained with antibodies against HIPK4 (green) and TAX1BP1 (magenta) and DAPI (blue). Cyanoquinoline **25** induces the formation of HIPK4- and TAX1BP1-positive puncta. Scale bars: 10 µm (full image) and 5 µm (inset).

We then investigated plausible drivers of HIPK4 aggregation in spermatids. Autophagic regulation of proteostasis plays a major role in spermatid differentiation,^33, 34^ and this process is predominantly mediated by three autophagy receptors: SQSTM1 (sequestome 1)/p62, NBR1 (Neighbor of *BRCA1* gene 1), and TAX1BP1 (Tax1-binding protein 1). We therefore conducted co-immunoprecipitation experiments to examine whether HIPK4 can associate with one or more of these receptors in a cyanoquinoline-sensitive manner. We cultured dissociated testis cells in the absence or presence of compound **25** for 5 hours, lysed these spermatid-containing suspensions, and then immunoprecipitated either HIPK4 or individual autophagy receptors. We observed that cyanoquinoline treatment markedly increased the co-immunoprecipitation of TAX1BP1 and HIPK4, using either anti-HIPK4 or anti-TAX1BP1 as the pulldown baits, respectively (**Fig. 8B**). Notably, HIPK4 interactions with NBR1 and p62 were not enriched upon treatment of compound **25** (**Fig. S7**).

Building on these findings, we compared the subcellular localizations of HIPK4 and TAX1BP1 in spermatids by immunofluorescence microscopy. Like HIPK4, TAX1BP1 is distributed through the spermatid cytoplasm and forms a limited number of puncta, which are spatially distinct from the sporadic HIPK4-positive foci in untreated germ cells (**Fig. 8C**). When we cultured the spermatids with compound **25** for 5 hours, the number of TAX1BP1-positive puncta increased, and a substantial fraction of these foci were also HIPK4-positive. Together, these results supported a model in which cyanoquinoline ligands induced a HIPK4 conformation that recruited TAX1BP1 and perhaps other cellular proteins to form an aggregated or oligomerized state.

## DISCUSSION

Our study identifies cyanoquinolines as a versatile chemotype for HIPK4 ligand development. Several cyanoquinolines selectively inhibit HIPK4 with nanomolar potency, and our SAR studies support a binding mode in which the C4 substituent and C3 cyano group engage the ATP-binding site, mirroring the molecular interactions observed in other kinase/quinoline and kinase/quinazoline complexes.^23^ The parental scaffold exhibits an intrinsic bias toward HIPK4 binding, as evidenced by the activity profile of cyanoquinoline **1** in the 192-kinase NanoBRET platform. This selectivity for HIPK4 may reflect unique spatial constraints within the ATP-binding pocket, as more sterically demanding C4 substituents generally decrease inhibitory potency.^35^ Consistent with this idea, the macrocyclic quinolines developed by the Hanke group also have relatively compact hinge-facing C4 substituents. The locked conformations of these macrocycles also provide a means for eliminating GAK inhibition,^31^ and further optimization of the cyanoquinoline scaffold could yield improvements in cellular potency.

In addition to developing a new series of HIPK4 inhibitors, we fortuitously discovered that certain cyanoquinoline-based ligands can disrupt HIPK4 proteostasis. This activity does not correlate with inhibitor potency, indicating it is not merely a consequence of blocking HIPK4 kinase activity. Our findings align with a recent report that a considerable fraction of kinase inhibitors, including clinical compounds, act through additional mechanisms to induce target degradation or depletion.^36^ Moreover, the cyanoquinoline-dependent effects on HIPK4 homeostasis are dependent on cellular context. In HEK293T cells, the synthetic ligands induce HIPK4 degradation through a UPS-dependent mechanism, whereas the compounds promote HIPK4 aggregation in spermatids, the physiologically relevant cell type for HIPK4 expression and function. The SAR profiles for cyanoquinoline-altered HIPK4 proteostasis also do not completely overlap between the HEK293T cells and spermatids.

Although the structural basis for cyanoquinoline-dependent HIPK4 degradation or aggregation remains to be determined, the HIPK4 primary sequence and AlphaFold model of its three-dimensional structure provide some clues. The HIPK4 kinase domain shares only 50% sequence identity with the other HIPK family members and residues within the αC helix and flanking β4-β5 sheet that are directed away from the ATP-binding site are especially divergent (**Fig. 1** and **Fig. S1**). Many of these HIPK4-specific residues are hydrophobic or positively charged. In the AlphaFold model, this weakly conserved region within the N-terminal lobe engages one of the predicted helical sequences C-terminal to the kinase domain (residues 420-438) (**Fig. 9A**). The other putative helix (residues 501-508) engages the opposite face of the αC helix. If the AlphaFold model is correct, it is likely that ligand-induced conformational shifts in the HIPK4 kinase domain translate into greater changes in tertiary structure for the full-length protein (**Fig. 9B**). This could involve interactions between the αC helix “in” and “out” states and the predicted C-terminal helices, and more distal sites in HIPK4 could also participate in these regulatory mechanisms. For example, HIPK4 could contain C-terminal sequences that are functionally analogous to the autoinhibitory domain in HIPK2.^37^

**Figure 9.**
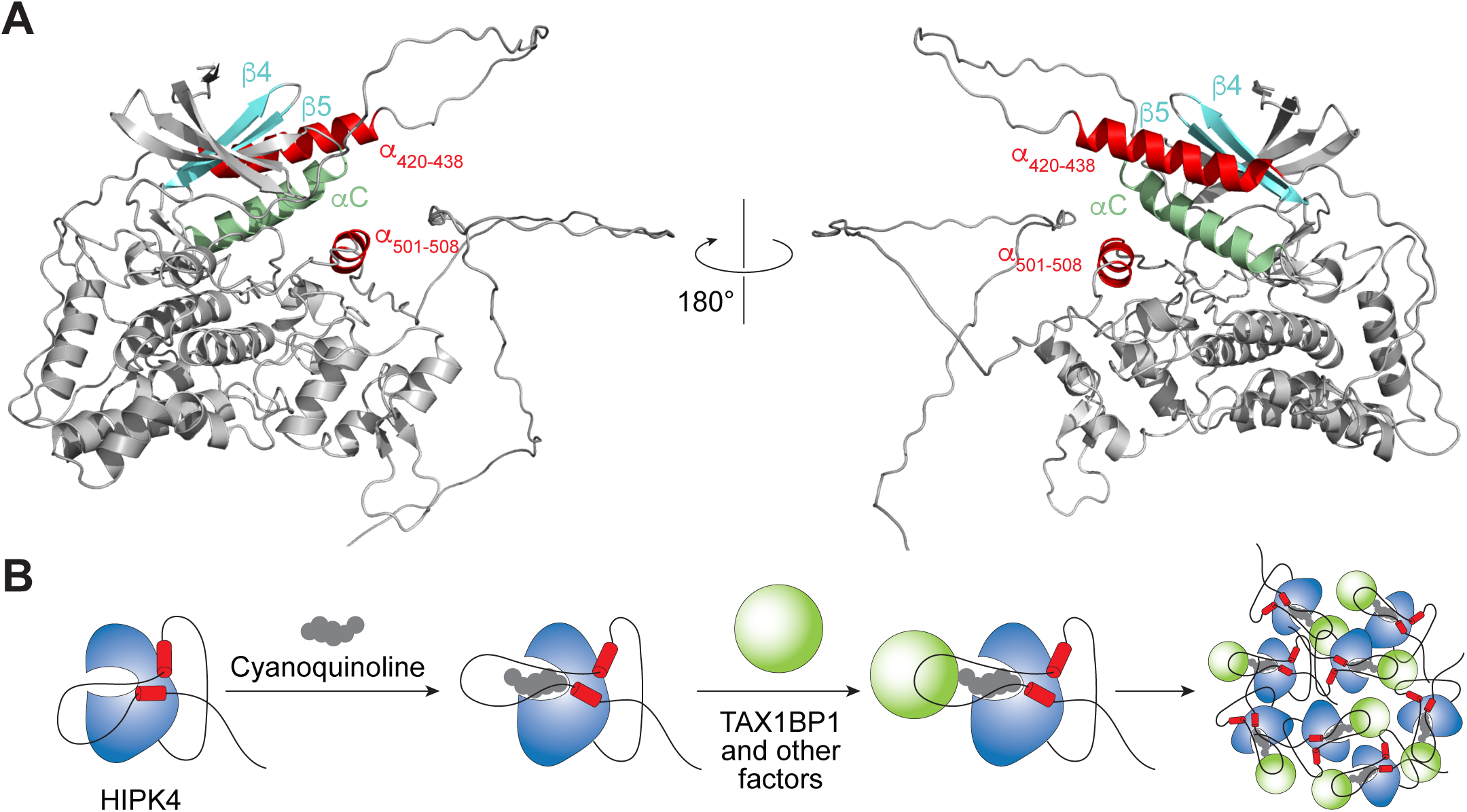
A model for cyanoquinoline action on HIPK4 proteostasis. (A) AlphaFold model of HIPK4 highlighting the predicted interactions between the αC helix, β4 and β5 strands, and putative helical structures (residues 420-438 and 501-508) in the C-terminal region. (B) These interactions could enable cyanoquinoline-induced conformational changes to actuate HIPK4 aggregation and TAX1BP1 binding.

Consistent with this model, we observed that the HIPK4-aggregating activity of cyanoquinolines correlates with scaffold modifications that are most likely to modulate HIPK4 conformation. These include *para* substituents on the C4 aniline ring that could directly engage the αC helix, and sterically bulky groups at the C6 and C7 sites that might prevent the kinase domain from adopting a “closed, αC helix-in” state. Notably, some of these modifications increased *Amax* values in spermatids at the expense of inhibitory potency, perhaps reflecting an energetic cost of achieving an aggregation-prone HIPK4 conformer. Our SAR studies further demonstrated that subtle changes in ligand structure can result in substantial differences in HIPK4-aggregating activity. For example, *para-*substituted chloroaniline **25** efficiently promoted HIPK4 aggregation but the corresponding *ortho*- or *meta*-substituted analogs (**23** and **24**, respectively) did not. Similarly, macrocyclic quinolines that differed in only one linker carbon exhibited highly divergent *Amax* values (compounds **56** vs **57**). These observations provide further evidence that ligand-dependent HIPK4 aggregation requires specific protein conformations. We hypothesize that in spermatids this process involves the binding of TAX1BP1 to a HIPK4 epitope that becomes more accessible upon cyanoquinoline binding (**Fig. 9B**).

Our findings have important implications for the development of HIPK4-targeting male contraceptives. Whether the cyanoquinoline-induced HIPK4 aggregates in spermatids are eventually cleared through autophagy remains unknown; current spermatid culture protocols maintain cell viability for only one day, precluding our ability to follow HIPK4 proteostasis at later time points. Nevertheless, it is possible that HIPK4 aggregation is an irreversible process, as dissociation of the protein assemblies could be kinetically unfavorable. If so, many of the pharmacological advantages of small-molecule degraders would be applicable to our compounds. The structural requirements for HIPK4 aggregation extend beyond ligand binding, providing an additional layer of specificity. The cyanoquinoline ligands could also act in a substoichiometric manner, inducing a HIPK4 conformational state that catalytically seeds an aggregation cascade. These mechanistic modalities could be especially valuable for male contraceptives, since blood-testis barrier penetrance and limited drug access to germ cells remain significant hurdles for clinical development.

Finally, the ability of cyanoquinolines to promote the formation of HIPK4- and TAX1BP1-containing aggregates could reflect physiological mechanisms of HIPK4 regulation and function. We could observe a limited number of HIPK4-positive puncta in untreated male germ cells, suggesting that at least some fraction of the kinase could normally exist in an oligomeric state. TAX1BP1 engagement with protein cargos often triggers the recruitment of machinery required for autophagosome formation and aggregate clearance,^38^ and cyanoquinolines could shift HIPK4 proteostasis toward TAX1BP1-dependent aggrephagy. Alternatively, the developed inhibitors could lock HIPK4-TAX1BP1 complexes in a conformational state that is incompetent for autophagy. Our ability to distinguish between these two modes of action is limited by the *ex vivo* viability of spermatids, and cyanoquinoline derivatives suitable for *in vivo* studies will be valuable tools for investigating these mechanistic possibilities.

## MATERIALS AND METHODS

### Plasmid constructs

For HIPK4 ΔPEST expression, cDNA corresponding to 2-557 amino acid sequence of the human HIPK4 (Uniprot Q8NE63) and codon-optimized for mammalian expression was synthesized by Genscript and subcloned into a pcDNA3.1 vector (Thermo Fisher Scientific, V79020). The final construct encoded HIPK4 ΔPEST with an N-terminal mammalianized maltose binding protein fused through a TEV and thrombin-cleavable linker, an N-terminal FLAG tag, and a C-terminal hexahistidine epitope (mMBP-TEV-F2-FLAG-HIPK4 ΔPEST-His_6_) under control of a CMV promoter. For HiBiT reporter assays, full-length human HIPK4 was cloned into a pFN38K HiBiT CMV-neo vector to generate cDNA encoding an N-terminal HiBiT-HIPK4 fusion. For NanoBRET studies, full-length human HIPK4 was cloned into a p15A vector (Promega, NV3241) to generate cDNA encoding an N-terminal NanoLuc-HIPK4 fusion protein. Both reporter constructs contained short linker sequences between HIPK4 and the HiBiT or NanoLuc tag.

### Antibodies

A custom rabbit polyclonal anti-HIPK4 antibody was generated by FabGennix against the peptide PAGSKSDSNFSNLIRLSQVSPEED. The following commercial primary antibodies were used: anti-HiBiT (Promega, N7200), anti-LgBiT (Promega, N7100), anti-His tag (Proteintech, 66005-1-Ig), anti-BRD4 (Thermo Fisher Scientific, BL-149-2H5, A700-004), anti-TAX1BP1 (Proteintech, 14424-1-AP), anti-NBR1 (Proteintech, 16004-1-AP), and anti-p62/SQSTM1 (Proteintech, 66184-1-Ig).

### HIPK4 ΔPEST expression

Expi293F cells (Thermo Fisher Scientific) were maintained in Expi293 Expression Medium in vented, non-baffled Erlenmeyer shaker flasks at 37 °C in a humidified incubator (8% CO_2_) with orbital shaking at 125 rpm. Cells were cultured in suspension and passaged every 3 to 4 days by dilution into fresh, prewarmed medium to a final density of 0.3 × 10^6^ to 0.5 × 10^6^ viable cells/mL. Routine maintenance cultures were kept in early log phase and not allowed to exceed 5 × 10^6^ cells/mL; cells were typically split when they reached 3 × 10^6^ to 5 × 10^6^ cells/mL. The Expi293F cells were transfected the pcDNA3.1 mMBP-TEV-F2-HIPK4 ΔPEST-His_6_ construct using ExpiFectamine 293 according to the manufacturer’s protocols and cultured for 4 days. The cells were harvested by centrifugation and resuspended in the lysis buffer (1X PBS, pH 7.4, 2 mM MgCl_2_, 0.2 mM ATP, 10 mM β-mercaptoethanol, 5% glycerol) supplemented with DNase I and protease inhibitor cocktail.

The cells were homogenized by three passes through an EmulsiFlex-C3 homogenizer (Avestin). The lysate was clarified by centrifugation and loaded on gravity flow column packed with Co-NTA resin. After washes with the lysis buffer and wash buffer (1X PBS, pH 7.4, 1 M NaCl, 2 mM MgCl_2_, 0.2 mM ATP, 10 mM β-mercaptoethanol, 5% glycerol), the protein was eluted with the CoNTA elution buffer (1X PBS, pH 7.4, 300 mM imidazole, 2 mM MgCl_2_, 0.2mM ATP, 10 mM β-mercaptoethanol, 10% glycerol) and dialyzed into the lysis buffer. After dialysis, the protein was loaded on an amylose gravity flow column, washed with the lysis buffer and eluted, with the MBP elution buffer (1X PBS, pH7.4, 10 mM maltose, 2 mM MgCl_2_, 0.2 mM ATP, 10 mM - mercaptoethanol, 10% glycerol). Eluted protein was dialyzed into the lysis buffer and loaded on Superdex-200 column. Superdex-200 elution fractions were analyzed by SDS-PAGE. Purified mMBP-TEV-F2-FLAG-HIPK4-ΔPEST-His_6_ fractions were pooled, concentrated and frozen in liquid nitrogen.

To remove the mMBP, a portion of the prep was dialyzed into cleavage buffer (20 mM Tris-HCl, pH 8.4, 150 mM NaCl, 2.5 mM CaCl_2_) and incubated with 30 units of thrombin per 1 mg of protein for 2 h at room temperature. After the cleavage the mixture was loaded on a Co-NTA column, washed with the lysis buffer, and eluted with imidazole. Elution fractions were analyzed by SDS-PAGE, and fractions containing purified FLAG-HIPK4-ΔPEST-His_6_ were pulled, concentrated, and frozen in liquid nitrogen.

### High-throughput screen

Differential scanning fluorimetry (also known as protein thermal shift (PTS)) measurements of FLAG-HIPK4 ΔPEST-His_6_ was performed using optimized methods and conditions in accordance with those previously described.^39^ The PTS assays were prepared in a 384-well plate format by combining FLAG-HIPK4 ΔPEST-His_6_ with individual compounds (final 25 µM in assay), thermal shift dye (SYPRO Orange) and buffer to a final assay volume of 10 μL. Test compounds (25 nL of 10 mM DMSO stocks) were dry spotted into MicroAmp 384-well real-time PCR plates (Applied Biosystems, 4483285) using an Echo 555 liquid handler (Beckman). Then 5 μL of the FLAG-HIPK4 ΔPEST-His_6_ working solution (various concentrations for assay development and 2.0 µg protein for screen in PBS) were added to each well using a Multidrop Combi reagent dispenser (Thermo Fisher Scientific). 5 μL of 5X SYPRO Orange (Invitrogen/Thermo Fisher Scientific) dissolved in PBS was equally dispensed into the PCR plates diluting the enzyme solution 1:2 (0.25% DMSO final concentration). The plates were then sealed with MicroAmp optical adhesive film (Applied Biosystems) and centrifuged briefly (1 min at 1000 g) to collect the assay mix at the bottom of the plate. The plates were developed and analyzed using a ViiA 7 real-time PCR instrument (Applied Biosystems) and an 8-min temperature gradient with a temperature increase of 0.1125 °C/s. The melting temperature (T_m_D) and thermogram (fluorescence vs temperature) were determined as described previously using Protein Thermal Shift software (version 1.2; Applied Biosystems).^39^ Hits were defined as those compounds that shifted the melting temperature relative to DMSO controls by 1°C or more (ΔT_m_D ≥ 1°C). T_m_D values for 7,500 compounds in the ChemDiv Allosteric Kinase Inhibitors collection at 25 µM against DMSO controls (0.25% DMSO final), and “hits” were selected at ΔT_m_D ≥ 1°C. These compounds were “cherry-picked’ for confirmation assays by dry spotting them in triplicate wells of a 384-well PCR plate, which was then processed as described above.

The remaining 18,500 compounds in the ChemDiv Allosteric Kinase Inhibitors collection were screened using the ADP-Glo assay platform (Promega, V6930). ADP-Glo is a universal bioluminescent, homogeneous assay for monitoring ADP produced by kinases of many classes phosphorylating a wide variety of substrates in the presence of ATP. Positive control pacritinib (at Col 1-2, final 5 μM) and DMSO at Col 3-4 (10 nL) and test compounds at Col 5-44 (10 μM final; 10 nL of 2 mM DMSO stocks) were dry spotted into wells of 1,536-well microtiter plates (Corning, 3725) using an Echo 555 liquid handler (Beckman). Then 1 μL of an mMBP-TEV-F2-HIPK4 ΔPEST-His_6_ protein solution (2 µg/mL) in 50 mM Tris-HCl, pH 8.0, 5 mM MgCl_2_, 0.1% Triton X-100 was added to each well using a Multidrop Combi reagent dispenser (Thermo Fisher Scientific), and kinase reaction was started by adding 1 μL of myelin basic protein/ATP (MBP/ATP) working solution [(1.34 mg/mL MBP; Sigma-Aldrich, M1891) + 200 μM ATP (Sigma-Aldrich, 797189)] in the same buffer to each well (final DMSO concentration of 0.5%) and incubation for 60 min at room temperature. Final assay conditions were as follows: 1 μg/mL mMBP-TEV-F2-HIPK4 ΔPEST-His_6_, 0.67 mg/mL MBP substrate, 100 μM ATP in 50 mM Tris-HCl, pH 8.0, 5 mM MgCl_2_, 0.1% Triton X-100; 10 μM test compounds, 5 μM pacritinib control and DMSO control in final 0.5% DMSO.

The kinase reaction was then quenched and developed in two steps. Unused ATP was depleted by the addition of 2 μL ADP-Glo reagent 1 (Promega ADP-Glo Kit, V9102) to each well of the 1536-well plate with Multidrop Combi, mixing each plate by vortexing, centrifuging briefly to collect volume into well, then incubating at room temperature in darkened room for 40 min. Second, ADP from the kinase reaction was detected by adding 4 μL of ADP-Glo reagent 2 to each well with another Multidrop Combi, mixing each plate by vortexing, centrifuging briefly to collect volume into well, then incubating at room temperature in darkened room for 30-40 min.

Luminescence is then read out on each plate by an Envision (Perkin Elmer) or Pherastar FSX (BMG Labtech). Hits are defined usually as ≥50% Inhibition relative to controls.

To confirm hits for the ADP-Glo assay-based screen, the kinase reaction was conducted as described above, except that the Echo 555 is used to “cherry pick” by dry spotting 10 nL of 2 mM compound stock solutions in DMSO from each hit well from Echo-certified compound plates. Three independent compound transfers were arranged in adjacent blocks, with individual replicates spatially separated. This format minimizes any variability that may be caused by dispensing issues or positional plate effects and achieves technical triplicates for each initial hit. Hit confirmation is obtained when any one of the triplicates achieves ≥50% inhibition relative to controls.

Reproducible hits from the PTS and ADP-Glo screens were then purchased as dry powders and re-tested in the ADP-Glo assay. These validation studies identified 2 non-ATP-competitive and 12 ATP-competitive HIPK4 ligands, with the cyanoquinoline hit exhibiting the greatest potency and selectivity.

### ADP-Glo assays

Recombinant human HIPK1, HIPK2, HIPK3, and HIPK4 proteins were purchased as N-terminal GST fusions from Sino Biological (HIPK1, H03-11G-SCB; HIPK2, H04-11BG-SCB; HIPK3, H05-13H-SCB; HIPK4, H06-10G-SCB). Inhibitory activities of individual compounds against the HIPK family members were measured using the ADP-Glo Kinase Assay kit, according to the manufacturer’s instructions with minor modifications. Reactions were performed in 96-well plates in kinase reaction buffer (40 mM Tris-HCl, pH 7.5, 20 mM MgCl_2_, 0.1 mg/mL BSA). Each 25-µL reaction mixture contained 3 nM HIPK4, 2 µM myelin basic protein as substrate, and 25 µM ATP. Test compounds were prepared as serial dilutions in DMSO and added to give a final DMSO concentration of 1% (v/v). After preincubation of enzyme with compounds, reactions were initiated by addition of the substrate/ATP mixture and incubated for 30 min at room temperature. Kinase reactions were terminated by addition of 25 µL ADP-Glo Reagent followed by a 40 min incubation at room temperature to deplete remaining ATP. Subsequently, 50 µL of the Kinase Detection Reagent was added, and plates were incubated for an additional 30 min before luminescence measurement using a Promega GloMax Navigator microplate reader. Percent inhibition was calculated relative to DMSO-treated controls, and IC_50_ values were determined by nonlinear regression analysis using a four-parameter logistic equation in GraphPad Prism. Reported values represent the mean of at least two independent experiments performed in triplicate.

### NanoBRET assays

Freshly passaged HEK293 cells were resuspended in assay medium at 2 × 10^5^ cells/mL and transiently transfected with the p15A-HIPK4-NanoLuc plasmid using FuGENE HD transfection reagent (Promega, E2311). Briefly, the carrier DNA (Promega, E4881) and HIPK4-NanoLuc plasmid were prepared at a 9:1 ratio (w/w) and total concentration of 10 µg/mL in Opti-MEM I Reduced Serum Medium without phenol red, followed by addition of the FuGENE HD reagent at 30 µL/mL and incubation for 20 min at room temperature. The transfection mixture was added to cells at a 1:20 (v/v) ratio, and 100 µL/well was plated into white 96-well plates, corresponding to approximately 20,000 cells/well. After 20 to 30 h, NanoBRET measurements were performed in live cells using the NanoBRET Nano-Glo Detection System (Promega). Donor and acceptor emissions were measured on BMG Labtech CLARIOstar Plus luminometer equipped with 450-nm BP (donor) and 610-nm LP (acceptor) filters, and NanoBRET values were calculated as acceptor/donor ratios.

### NanoBRET assay-based kinome profiling

To assess kinase inhibitor selectivity, the K192 Kinase Selectivity System (Promega, NP4050) was performed as previously described.^40^ In brief, for plate preparation, the transfection mix was prepared in white 384-well small-volume plates (Greiner, 784075) by pre-plating 3 µL of 20 µL/mL FuGENE HD, diluted in optiMEM medium (Gibco, 11058-021). 1 µL DNA/kinase from both vector source plates of the K192 kit was added using an Echo acoustic dispenser (Beckman Coulter). The mix was incubated for 30 min and 6 µL of HEK293T cells in optiMEM medium were added. The proteins were allowed to express for 20 h. After expression, Tracer K10 was added using the concentrations recommended in the K192 technical manual, and 1 µM inhibitor was added to every second well. After 2 h of equilibration, detection was carried out using substrate solution comprising optiMEM with a 1:166 dilution of NanoBRET Nano-Glo Substrate and a 1:500 dilution of the Extracellular NanoLuc Inhibitor. 5 µL of substrate solution was added to every well and filtered luminescence was measured on a PHERAstar plate reader (BMG Labtech) equipped with a luminescence filter pair (450 nm BP filter (donor) and 610 nm LP filter (acceptor)). For every kinase, occupancy was calculated and plotted using GraphPad Prism 10.

### HiBiT-HIPK4 degradation assays

LgBiT-HEK293 cells (Promega, N2672) were transfected with the HiBiT-HIPK4 expression construct using FuGENE 4K transfection reagent (Promega, E5911) according to the manufacturer’s protocol. After 48 h, transfected cells were subjected to three rounds of selection with G418 to enrich for stable integrants. The selected population was then single-cell sorted by fluorescence-activated cell sorting (FACS), and individual clones were expanded in culture. Expression of HiBiT-HIPK4 and LgBiT in each clone was evaluated by western blot analysis. A clone with robust luminescence signals was selected for real-time degradation assays.

The HiBiT-HIPK4- and LgBiT-expressing HEK293T cell line was maintained in Dulbecco’s modified Eagle’s medium (DMEM; Gibco, 11965118) supplemented with 10% fetal bovine serum (FBS; Sigma-Aldrich, F0392) and 1% penicillin-streptomycin (Thermo Fisher Scientific, 15140122). HiBiT degradation assays were carried out by seeding the cells into white 96-well plates (Corning 3917) at 2 × 10^4^ cells/well in 100 μL of medium and cultured overnight. To reduce edge effects, only the inner 60 wells were used for analysis; outer wells were filled with medium or PBS, and PBS was added to the spaces between wells. The next day, growth medium was replaced with 90 μL of growth medium containing Endurazine (Promega, N2571) diluted 1:100, and the plates were incubated at 37 °C for 2 h. Test compounds were prepared as 10X stocks in DMEM and added at 10 μL/well to achieve the indicated final concentrations. Luminescence was recorded at the indicated time points on the Promega GloMax Navigator microplate reader.

For cycloheximide experiments, the protein translation inhibitor was added to a final concentration of 10 μg/mL 1 h before degrader addition, with DMSO used as the vehicle control. For mechanistic studies, the HiBiT-HIPK4- and LgBiT-expressing HEK293T cells were preincubated with individual proteolysis inhibitors (MedChemExpress) for 2 h prior to treatment with 10 µM degrader. The inhibitors were used at the following concentrations: MG-132 (10 μM), carfilzomib (0.1 μM), TAK-243 (1 μM), pevonedistat (1 μM), chloroquine (30 μM), eeyarestatin I (20 μM), and Z-VAD-FMK (40 μM).

The effects of individual compounds on the cell viability were also assessed using the CellTiter-Glo 3D Cell Viability Assay (Promega, G9683) following the manufacturer’s instructions. Briefly, after compound treatment, the 96-well plate cultures of HiBiT-HIPK4- and LgBiT-expressing HEK293T cells were equilibrated to room temperature and an equal volume of CellTiter-Glo 3D reagent was added to each well. The plates were shaken for 5 min, incubated at room temperature for 25 min, and luminescence was then measured using the Promega GloMax Navigator microplate reader.

### Spermatid assays of HIPK4 proteostasis

All animal experiments were performed in accordance with protocols approved by the Institutional Animal Care and Use Committee of Stanford (IACUC protocol 29999). Male C57BL/6J mice (4 to 6 weeks old; The Jackson Laboratory, stock no. 000664) were maintained at Stanford University under approved housing conditions and used for testis collection. Mice were euthanized by regulated carbon dioxide inhalation followed by cervical dislocation, in accordance with the recommendations of the 2020 American Veterinary Medical Association Panel on Euthanasia. Testes were collected immediately after euthanasia, transferred into PBS, cleared of adherent fat and epididymal tissue, and decapsulated by removal of the tunica albuginea.

Testicular cells were prepared by sequential enzymatic dissociation adapted from published STA-PUT and BSA sedimentation protocols. Briefly, decapsulated testes were incubated in digestion medium I consisting of Krebs buffer (120 mM NaCl, 25 mM NaHCO_3_, 11 mM dextrose, 4.8 mM KCl, 1.3 mM CaCl_2_, 1.2 mM KH_2_PO_4_, and 1.2 mM MgSO_4_) supplemented with collagenase A (0.5 mg/mL; Worthington Biochem, LS004196) at 37 °C for 15 min, with gentle trituration after the first 5 min and again at the end of the incubation to release seminiferous tubules while minimizing tubule fragmentation. The seminiferous tubule-enriched fraction was then transferred to digestion medium II consisting of Krebs buffer supplemented with trypsin (0.05%; Worthington Biochem, LS02123) and DNase I (0.2 mg/mL; Sigma-Aldrich, 11284932001) and incubated at 37 °C for 15 min with gentle pipetting until a single-cell suspension was obtained. The suspension was passed sequentially through 40-μm cell strainers, and cells were collected by centrifugation at 500 g for 5 min at 4 °C. Single cell suspension can be readily used for experiments or further enrichment for spermatids.

To isolate round spermatids from the dissociated testes, the single-cell suspension was subjected to unit-gravity sedimentation through a BSA gradient prepared in Krebs buffer. A sedimentation column was layered sequentially with 4% BSA, 2% BSA, and 0.5% BSA, and the testicular cell suspension in 0.5% BSA was gently loaded onto the top of the column. Cells were allowed to sediment for 150 min, after which fractions were collected at a flow rate of 2 mL/min in 4-mL aliquots. Fractions were pelleted at 1000 g for 10 min at 4 °C, resuspended in Krebs buffer, and examined microscopically. Spermatid-enriched fractions were identified on the basis of sedimentation profile and cell morphology and were pooled for subsequent experiments. This procedure was adapted from a spermatid isolation method that uses a STA-PUT BSA sedimentation workflow.^41^

Cells from dissociated testes or isolated spermatids were cultured in Advanced DMEM/F-12 (Dulbecco’s Modified Eagle Medium/Ham’s F-12; Thermo Fisher Scientific, 11320033) supplemented with 1% FBS and 1% penicillin-streptomycin. To assess the effects of individual compounds on the endogenous HIPK4 expressed in spermatids, the germ cells were cultured with the compounds for 16 h, after which they were collected by centrifugation and lysed in RIPA buffer [(50 mM Tris-HCl, pH 7.4, 150 mM NaCl, 1% NP-40, 0.25% deoxycholic acid, and 1 mM EDTA prepared from a 10X stock; Millipore, 20-188), supplemented with 1× cOmplete protease inhibitor (Roche Cat. No. 4693132001), 1X PhosSTOP (Roche, 4906845001), and 1 mM PMSF]. The detergent-solubilized lysates were next centrifuged at 13 000 g for 15 min at 4 °C to separate RIPA-soluble and RIPA-insoluble fractions. Both fractions were resuspended in 1X SDS sample buffer, and boiled for 10 min, and subsequently analyzed by SDS-PAGE and western blot.

For western blot studies, proteins were separated on 4–15% Criterion TGX Stain-Free gels (Bio-Rad, 5678084) and transferred onto 0.2-μm PVDF membranes using a Trans-Blot Turbo Transfer System (Bio-Rad, 1704150) with Midi PVDF transfer packs (Bio-Rad, 1704157). The membranes were blocked with 4% BSA in TBST [1X Tris Buffered Saline (BioRad, 1706435) with 0.1% Triton X-100] for 1 h at room temperature, followed by incubation with primary antibodies overnight at 4 °C. Primary antibodies are described above and were used at the following dilutions: anti-HIPK4, 1:300; anti-His tag, 1:1000; all other primary antibodies, 1:500. After washing with TBST, the membranes were incubated with horseradish peroxidase (HRP)-labeled secondary antibodies (Cytiva Amersham ECL; anti-rabbit, NA934-1ML; anti-mouse, NA931-1ML) as appropriate) at a 1:10,000 dilution for 1 h at room temperature. Following additional washes with TBST, the chemiluminescent signal was obtained using the SuperFemto ECL Chemiluminescence Kit (Vazyme, E423) and recorded with a Bio-Rad Gel Doc imaging system.

For mechanistic studies in isolated spermatids, the germ cells were preincubated with the ubiquitin/proteasome system inhibitors described above for 2 h, followed by treatment with degrader at the indicated concentration. The inhibitors were used at the following concentrations: TAK-243 (1 μM), pevonedistat (1 μM), and carfilzomib (0.1 µM). After 16 h, the spermatids were harvested, treated with trypsin, and pelleted by centrifugation at 500 g for 5 min. The cell pellets were lysed in RIPA buffer for 15 min and centrifuged at 13,000 g for 15 min at 4 °C to separate the soluble and insoluble fractions for subsequent analyses by SDS-PAGE and western blotting.

### Mass photometry

Solution-phase interferometric scattering measurements were performed using a TwoMP mass photometer (Refeyn). A buffer blank (100 mM Tris, pH 7.4, 200 mM NaCl, and 2 mM MgCl₂) was acquired immediately prior to sample measurement. Mass calibration was performed using P1 standard (Refeyn, MP-CON-41033), prepared by diluting 2 μL of the calibrant into 498 μL of buffer and subsequently diluted 1:1 with the blank buffer on the sample carrier to generate a calibration curve. Full-length His_6_-HIPK4 protein (100 nM; Sino Biological, H06-10H) was incubated with 5 µM cyanoquinoline **25** or DMSO vehicle alone for 30 min at room temperature prior to measurement. The samples were diluted 1:1 on the sample carrier to a final protein concentration of 50 nM. Mass photometry videos for both the calibrant and samples were recorded for 60 s and analyzed using DiscoverMP software (Refeyn).

### Immunofluorescence microscopy

Testes from ∼6-week-old mice were dissected, decapsulated, and dissociated into a single-cell suspension as described above. The dissociated testis cells were washed with PBS and incubated with either degrader (25 μM) or DMSO vehicle control for 5 h. Following treatment, the cells were collected, trypsinized, and pelleted by centrifugation at 500 g for 5 min. The cell pellet was washed once with PBS and resuspended in 500 µL PBS. An 50-µL aliquot of the cell suspension was then applied to a microscope slide (Sigma-Aldrich, MTCM710045) and allowed to settle for 25 min at room temperature. Cells were next fixed with 4% (w/v) paraformaldehyde and 0.1 M sucrose in PBS for 20 min, washed with PBS, and permeabilized with 1% (v/v) Triton X-100 in PBS for 10 min. Following an additional PBS wash, cells were in blocking buffer [5% (v/v) donkey serum, 1% (w/v) bovine serum albumin, and 0.1% (v/v) Triton X-100] for 1 h at room temperature.

Cells were then incubated with primary antibody against HIPK4 (1:100 dilution in blocking buffer) overnight at 4 °C. After washing with PBS, cells were incubated with Alexa Fluor 594-conjugated secondary antibody (Thermo Fisher Scientific, A-11012; 1:400 dilution in blocking buffer) for 1 h at room temperature. Cells were washed with PBS and mounted using mounting medium containing DAPI (Vector Laboratories, H-1800-2). For colocalization experiments, primary antibodies against HIPK4 and TAX1BP1 were labeled with fluorophores (Proteintech, KFA503 and KFA509) and used as described above. Fluorescence imaging was performed on a Zeiss Axio Imager Z2 confocal microscope equipped with a 63X oil-immersion objective.

### Co-immunoprecipitation studies

Testes from a ∼6-week-old mouse were dissected, decapsulated, and dissociated into a single-cell suspension as described above. Cells were washed with PBS and incubated with degrader (25 μM) or DMSO vehicle control for 5 h. Cells were then collected, trypsinized, and pelleted by centrifugation at 500 g for 5 min. The cell pellet was washed with PBS and lysed on ice for 15 min in lysis buffer (25 mM Tris-HCl, pH 7.4, 150 mM NaCl, 1% NP-40, 5% glycerol, and 2 mM MgCl₂) supplemented with protease and phosphatase inhibitors. Lysates were clarified by centrifugation (15,000 g,15 min, 4 °C), and 400 µL of the soluble fraction was incubated with the pull-down antibody (8 μg) overnight at 4 °C with rotation. Protein A/G-conjugated magnetic beads (50 μL; Thermo Fisher Scientific, 80104G) were then added, and the mixture was incubated for 30 min at room temperature with mixing. Beads were collected using a magnetic stand and washed with PBS. Bound proteins were released by boiling the beads in 1X SDS-PAGE loading buffer and then analyzed by SDS-PAGE and western blotting.

## AUTHOR INFORMATION

### Author contributions

Z.Z., R.K.T., and J.K.C. designed the project; Z.Z., P.K., R.K.T, S.D., D.S., B.K.M., and S.H. synthesized the compounds; A.B., L.M.F., and S.M.S. performed HIPK4 overexpression and purification; I.P., F.Z, S.W.S., and A.E. conducted HTS and hits validation; Z.Z. and R.K.T. performed ADP-Glo assays, NanoBRET assays, HiBiT assays; A.Z. and N.D.R. provided compounds **56-58**; M.P.S., A.Z., and N.D.R. performed K192-screens; R.K.T. and Z.Z. performed immunofluorescence, mass photometry and co-immunoprecipitations studies; J.K.C, T.D.C., and T.H. supervised the research. The manuscript was written by Z.Z., R.K.T., and J.K.C. with contributions from all authors.

## Supporting information

Compound SAR Summary

## ACKNOWLEDGEMENTS

This work was supported by the National Institutes of Health (R33 HD09972005 to J.K.C. and T.D.Y.C; R35 GM127030 to J.K.C.; T32 GM136631 training grant support to R.K.T.; and a P50 HD106793 subaward to Z.Z.), the Male Contraceptive Initiative (Agility Grant to J.K.C.; MCI Postdoctoral Fellowship to Z.Z.; MCI Predoctoral Fellowship to R.K.T.). S.K., M.P.S., A.Z. N.R. and T.H. are grateful for support by the Structural Genomics Consortium (SGC), a registered charity (no. 1097737) that receives funds from Bayer AG, Boehringer Ingelheim, Bristol Myers Squibb, Genome Canada through the Ontario Genomics Institute, EU/EFPIA/OICR/McGill/KTH/Diamond Innovative Medicines Initiative 2 Joint Undertaking [EUbOPEN grant 875510], Pfizer, and Takeda. S.K. also received funding from the German Cancer Research Center (DKTK), the German Cancer Aid (Krebsilfe) network TACTIC, the “Clusters4Future” network funded by the Federal Ministry of Education and Research (BMBF, 03ZU1109FA) and the BMFTR grant PREVENT (01GR2501A). Compound characterization was conducted using high-field NMR instrumentation supported by NIH S10 OD0288697, and mass photometry studies were conducted using a MassFluidix Mass Photometer supported by NIH S10 OD038355.

## REFERENCES

(1) Sedgh, G.; Singh, S.; Hussain, R. Intended and unintended pregnancies worldwide in 2012 and recent trends. Stud. Fam. Plann. 2014, 45 (3), 301–314. DOI: 10.1111/j.1728-4465.2014.00393.x.

(2) Basaria, S.; Coviello, A. D.; Travison, T. G.; Storer, T. W.; Farwell, W. R.; Jette, A. M.; Eder, R.; Tennstedt, S.; Ulloor, J.; Zhang, A.;, et al. Adverse events associated with testosterone administration. N. Engl. J. Med. 2010, 363 (2), 109–122. DOI: 10.1056/NEJMoa1000485.

(3) Ilani, N.; Swerdloff, R. S.; Wang, C. Male hormonal contraception: potential risks and benefits. Rev. Endocr. Metab. Disord. 2011, 12 (2), 107–117. DOI: 10.1007/s11154-011-9183-3.

(4) Finkle, W. D.; Greenland, S.; Ridgeway, G. K.; Adams, J. L.; Frasco, M. A.; Cook, M. B.; Fraumeni, J. F., Jr.; Hoover, R. N. Increased risk of non-fatal myocardial infarction following testosterone therapy prescription in men. PLoS One 2014, 9 (1), e85805. DOI: 10.1371/journal.pone.0085805.

(5) Chung, S. S.; Wang, X.; Roberts, S. S.; Griffey, S. M.; Reczek, P. R.; Wolgemuth, D. J. Oral administration of a retinoic Acid receptor antagonist reversibly inhibits spermatogenesis in mice. Endocrinology 2011, 152 (6), 2492–2502. DOI: 10.1210/en.2010-0941.

(6) Amory, J. K.; Muller, C. H.; Shimshoni, J. A.; Isoherranen, N.; Paik, J.; Moreb, J. S.; Amory, D. W., Sr.; Evanoff, R.; Goldstein, A. S.; Griswold, M. D. Suppression of spermatogenesis by bisdichloroacetyldiamines is mediated by inhibition of testicular retinoic acid biosynthesis. J. Androl. 2011, 32 (1), 111–119. DOI: 10.2164/jandrol.110.010751.

(7) Matzuk, M. M.; McKeown, M. R.; Filippakopoulos, P.; Li, Q.; Ma, L.; Agno, J. E.; Lemieux, M. E.; Picaud, S.; Yu, R. N.; Qi, J.;, et al. Small-molecule inhibition of BRDT for male contraception. Cell 2012, 150 (4), 673–684. DOI: 10.1016/j.cell.2012.06.045.

(8) Balbach, M.; Rossetti, T.; Ferreira, J.; Ghanem, L.; Ritagliati, C.; Myers, R. W.; Huggins, D. J.; Steegborn, C.; Miranda, I. C.; Meinke, P. T.;, et al. On-demand male contraception via acute inhibition of soluble adenylyl cyclase. Nat. Commun. 2023, 14 (1), 637. DOI: 10.1038/s41467-023-36119-6.

(9) Ku, A. F.; Sharma, K. L.; Ta, H. M.; Sutton, C. M.; Bohren, K. M.; Wang, Y.; Chamakuri, S.; Chen, R.; Hakenjos, J. M.; Jimmidi, R.;, et al. Reversible male contraception by targeted inhibition of serine/threonine kinase 33. Science 2024, 384 (6698), 885–890. DOI: 10.1126/science.adl2688.

(10) Crapster, J. A.; Rack, P. G.; Hellmann, Z. J.; Le, A. D.; Adams, C. M.; Leib, R. D.; Elias, J. E.; Perrino, J.; Behr, B.; Li, Y.;, et al. HIPK4 is essential for murine spermiogenesis. Elife 2020, 9. DOI: 10.7554/eLife.50209.

(11) Liu, X.; Zang, C.; Wu, Y.; Meng, R.; Chen, Y.; Jiang, T.; Wang, C.; Yang, X.; Guo, Y.; Situ, C.;, et al. Homeodomain-interacting protein kinase HIPK4 regulates phosphorylation of manchette protein RIMBP3 during spermiogenesis. J. Biol. Chem. 2022, 298 (9), 102327. DOI: 10.1016/j.jbc.2022.102327.

(12) Koser, S. A.; Rieck, C.; Aprea, I.; Krallmann, C.; Gaikwad, A. S.; Wallmeier, J.; Tenardi-Wenge, R.; Persio, S. D.; Neuhaus, N.; Raidt, J.;, et al. HIPK4 is a novel gene associated with teratozoospermia and male infertility. In medRxiv, 2026.

(13) Arai, S.; Matsushita, A.; Du, K.; Yagi, K.; Okazaki, Y.; Kurokawa, R. Novel homeodomain-interacting protein kinase family member, HIPK4, phosphorylates human p53 at serine 9. FEBS Lett. 2007, 581 (29), 5649–5657. DOI: 10.1016/j.febslet.2007.11.022.

(14) Kim, Y. H.; Choi, C. Y.; Lee, S. J.; Conti, M. A.; Kim, Y. Homeodomain-interacting protein kinases, a novel family of co-repressors for homeodomain transcription factors. J. Biol. Chem. 1998, 273 (40), 25875–25879. DOI: 10.1074/jbc.273.40.25875.

(15) Moilanen, A. M.; Karvonen, U.; Poukka, H.; Janne, O. A.; Palvimo, J. J. Activation of androgen receptor function by a novel nuclear protein kinase. Mol. Biol. Cell. 1998, 9 (9), 2527–2543. DOI: 10.1091/mbc.9.9.2527.

(16) van der Laden, J.; Soppa, U.; Becker, W. Effect of tyrosine autophosphorylation on catalytic activity and subcellular localisation of homeodomain-interacting protein kinases (HIPK). Cell Commun. Signal. 2015, 13, 3. DOI: 10.1186/s12964-014-0082-6.

(17) Singer, J. W.; Al-Fayoumi, S.; Ma, H.; Komrokji, R. S.; Mesa, R.; Verstovsek, S. Comprehensive kinase profile of pacritinib, a nonmyelosuppressive Janus kinase 2 inhibitor. J. Exp. Pharmacol. 2016, 8, 11–19. DOI: 10.2147/JEP.S110702.

(18) Davis, M. I.; Hunt, J. P.; Herrgard, S.; Ciceri, P.; Wodicka, L. M.; Pallares, G.; Hocker, M.; Treiber, D. K.; Zarrinkar, P. P. Comprehensive analysis of kinase inhibitor selectivity. Nat. Biotechnol. 2011, 29 (11), 1046–1051. DOI: 10.1038/nbt.1990.

(19) Flynn, G. A.; Bajji, A.; Huynh, K. HIPK4 inhibitors and uses thereof. US 11,629,143 B2, 2023.

(20) Zegzouti, H.; Zdanovskaia, M.; Hsiao, K.; Goueli, S. A. ADP-Glo: A Bioluminescent and homogeneous ADP monitoring assay for kinases. Assay Drug Dev. Technol. 2009, 7 (6), 560–572. DOI: 10.1089/adt.2009.0222.

(21) Levinson, N. M.; Boxer, S. G. A conserved water-mediated hydrogen bond network defines bosutinib’s kinase selectivity. Nat. Chem. Biol. 2014, 10 (2), 127–132. DOI: 10.1038/nchembio.1404.

(22) Getlik, M.; Grutter, C.; Simard, J. R.; Kluter, S.; Rabiller, M.; Rode, H. B.; Robubi, A.; Rauh, D. Hybrid compound design to overcome the gatekeeper T338M mutation in cSrc. J. Med. Chem. 2009, 52 (13), 3915–3926. DOI: 10.1021/jm9002928.

(23) Stamos, J.; Sliwkowski, M. X.; Eigenbrot, C. Structure of the epidermal growth factor receptor kinase domain alone and in complex with a 4-anilinoquinazoline inhibitor. J. Biol. Chem. 2002, 277 (48), 46265–46272. DOI: 10.1074/jbc.M207135200.

(24) Vasta, J. D.; Corona, C. R.; Wilkinson, J.; Zimprich, C. A.; Hartnett, J. R.; Ingold, M. R.; Zimmerman, K.; Machleidt, T.; Kirkland, T. A.; Huwiler, K. G.;, et al. Quantitative, wide-spectrum kinase profiling in live cells for assessing the effect of cellular ATP on target engagement. Cell Chem. Biol. 2018, 25 (2), 206–214 e211. DOI: 10.1016/j.chembiol.2017.10.010.

(25) Gerninghaus, J.; Zhubi, R.; Kramer, A.; Karim, M.; Tran, D. H. N.; Joerger, A. C.; Schreiber, C.; Berger, L. M.; Berger, B. T.; Ehret, T. A. L.;, et al. Back-pocket optimization of 2-aminopyrimidine-based macrocycles leads to potent EPHA2/GAK kinase inhibitors. J. Med. Chem. 2024, 67 (15), 12534–12552. DOI: 10.1021/acs.jmedchem.4c00411.

(26) Mensing, T. E.; Kurz, C. G.; Amrhein, J. A.; Ehret, T. A. L.; Preuss, F.; Mathea, S.; Karim, M.; Tran, D. H. N.; Kadlecova, Z.; Tolvanen, T. A.;, et al. Development of pyrazolo[1,5-a]pyrimidine based macrocyclic kinase inhibitors targeting AAK1. Eur. J. Med. Chem. 2025, 299, 118076. DOI: 10.1016/j.ejmech.2025.118076.

(27) Li, K.; Crews, C. M. PROTACs: past, present and future. Chem. Soc. Rev. 2022, 51 (12), 5214–5236. DOI: 10.1039/d2cs00193d.

(28) Saraswat, A. L.; Vartak, R.; Hegazy, R.; Patel, A.; Patel, K. Drug delivery challenges and formulation aspects of proteolysis targeting chimera (PROTACs). Drug Discov. Today 2023, 28 (1), 103387. DOI: 10.1016/j.drudis.2022.103387.

(29) Riching, K. M.; Mahan, S.; Corona, C. R.; McDougall, M.; Vasta, J. D.; Robers, M. B.; Urh, M.; Daniels, D. L. Quantitative Live-Cell Kinetic Degradation and Mechanistic Profiling of PROTAC Mode of Action. ACS Chem. Biol. 2018, 13 (9), 2758–2770. DOI: 10.1021/acschembio.8b00692.

(30) Mori, T.; Ito, T.; Liu, S.; Ando, H.; Sakamoto, S.; Yamaguchi, Y.; Tokunaga, E.; Shibata, N.; Handa, H.; Hakoshima, T. Structural basis of thalidomide enantiomer binding to cereblon. Sci. Rep. 2018, 8 (1), 1294. DOI: 10.1038/s41598-018-19202-7.

(31) Zerva, A.; Raig, N. D.; Zhuang, Z.; Krämer, A.; Dopfer, J.; Togashi, R.; Schwalm, M. P.; Elson, L.; Frischkorn, J.; Berger, B.-T.;, et al. Macrocyclization of Broad-Spectrum Kinase Inhibitor Bosutinib Leads to Potent and Selective Quinoline-based HIPK4 Inhibitor AZ137. In bioRxiv, 2026.

(32) Donovan, K. A.; Ferguson, F. M.; Bushman, J. W.; Eleuteri, N. A.; Bhunia, D.; Ryu, S.; Tan, L.; Shi, K.; Yue, H.; Liu, X.;, et al. Mapping the Degradable Kinome Provides a Resource for Expedited Degrader Development. Cell 2020, 183 (6), 1714–1731 e1710. DOI: 10.1016/j.cell.2020.10.038.

(33) Huang, Q.; Liu, Y.; Zhang, S.; Yap, Y. T.; Li, W.; Zhang, D.; Gardner, A.; Zhang, L.; Song, S.; Hess, R. A.;, et al. Autophagy core protein ATG5 is required for elongating spermatid development, sperm individualization and normal fertility in male mice. Autophagy 2021, 17 (7), 1753–1767. DOI: 10.1080/15548627.2020.1783822.

(34) Shang, Y.; Wang, H.; Jia, P.; Zhao, H.; Liu, C.; Liu, W.; Song, Z.; Xu, Z.; Yang, L.; Wang, Y.;, et al. Autophagy regulates spermatid differentiation via degradation of PDLIM1. Autophagy 2016, 12 (9), 1575–1592. DOI: 10.1080/15548627.2016.1192750.

(35) Denny, W. A.; Rewcastle, G. W.; Bridges, A. J.; Fry, D. W.; Kraker, A. J. Structure-activity relationships for 4-anilinoquinazolines as potent inhibitors at the ATP binding site of the epidermal growth factor receptor in vitro. Clin. Exp. Pharmacol. Physiol. 1996, 23 (5), 424–427. DOI: 10.1111/j.1440-1681.1996.tb02752.x.

(36) Scholes, N. S.; Bertoni, M.; Comajuncosa-Creus, A.; Kladnik, K.; Guo, X.; Frommelt, F.; Hinterndorfer, M.; Razumkov, H.; Prokofeva, P.; Schwalm, M. P.;, et al. Inhibitors supercharge kinase turnover through native proteolytic circuits. Nature 2026, 649 (8098), 1032–1041. DOI: 10.1038/s41586-025-09763-9.

(37) Gresko, E.; Roscic, A.; Ritterhoff, S.; Vichalkovski, A.; del Sal, G.; Schmitz, M. L. Autoregulatory control of the p53 response by caspase-mediated processing of HIPK2. EMBO J. 2006, 25 (9), 1883–1894. DOI: 10.1038/sj.emboj.7601077.

(38) Turco, E.; Savova, A.; Gere, F.; Ferrari, L.; Romanov, J.; Schuschnig, M.; Martens, S. Reconstitution defines the roles of p62, NBR1 and TAX1BP1 in ubiquitin condensate formation and autophagy initiation. Nat. Commun. 2021, 12 (1), 5212. DOI: 10.1038/s41467-021-25572-w.

(39) Niesen, F. H.; Berglund, H.; Vedadi, M. The use of differential scanning fluorimetry to detect ligand interactions that promote protein stability. Nat. Protoc. 2007, 2 (9), 2212–2221. DOI: 10.1038/nprot.2007.321.

(40) Schwalm, M. P.; Knapp, S. Single-plate kinome screening in live-cells to enable highly cost-efficient kinase inhibitor profiling. SLAS Discov. 2025, 31, 100214. DOI: 10.1016/j.slasd.2025.100214.

(41) Bryant, J. M.; Meyer-Ficca, M. L.; Dang, V. M.; Berger, S. L.; Meyer, R. G. Separation of spermatogenic cell types using STA-PUT velocity sedimentation. J. Vis. Exp. 2013, (80). DOI: 10.3791/50648.

