## Supplementary Information for "Small-molecule modulators of HIPK4 activity and proteostasis"

Synthetic Methods .......... 12 – 46

Compound Spectra .......... 47 – 178

```
                        β1     G-loop  β2            β3              αC
HIPK1  180 HEILCSMTNSYEVLEFLGRGTFGQVAKCWKRSTKEIVAIKILKNHPSYARQGQIEVSILS 239
HIPK2  189 HEVLCSMTNTYEVLEFLGRGTFGQVVKCWKRGTNEIVAIKILKNHPSYARQGQIEVSILA 248
HIPK3  187 HEVLCSMKNTYEVLDFLGRGTFGQVVKCWKRGTNEIVAIKILKNHPSYARQGQIEVSILA 246
HIPK4    1 MSTIQSETDCYDIIEVLGKGTFGEVAKGWRRSTGEMVAIKILKNDAYRNRIIKNELKLLH  60
          . *     .*....  **.****.*  *  *.*  *  *.********      *    .  *.  .*

                                β4      β5     Hinge    αD              αE
HIPK1  240 RLSSENADEYNFVRSYECFQHKNHTCLVFEMLEQNLYDFLKQNKFSPLPLKYIRPILQQV 299
HIPK2  249 RLSTESADDYNFVRAYECFQHKNHTCLVFEMLEQNLYDFLKQNKFSPLPLKYIRPVLQQV 308
HIPK3  247 RLSTENADEYNFVRAYECFQHRNHTCLVFEMLEQNLYDFLKQNKFSPLPLKVIRPILQQV 306
HIPK4   61 CMRGLDPEEAHVIRFLEFFHDALKFYLVFELLEQNLFEFQKENNFAPLPARHIRTVTLQV 120
            .        ..  .  .*   *  *.     .     ****.*****..*  *.*  *.***   .  **    .   **

                     Catalytic loop β6         β7        Activation loop
HIPK1  300 ATALMKLKSLGLIHADLKPENIMLVDPVRQPYRVKVIDFGSASHVSKA--VCSTYLQSRY 357
HIPK2  309 ATALMKLKSLGLIHADLKPENIMLVDPSRQPYRVKVIDFGSASHVSKA--VCSTYLQSRY 366
HIPK3  307 ATALKKLKSLGLIHADLKPENIMLVDPVRQPYRVKVIDFGSASHVSKT--VCSTYLQSRY 364
HIPK4  121 LTALARLKELAIIHADLKPENIMLVDQTRCPFRVKVIDFGSASIFSEVRYVKEPYIQSRF 180
             ***  .**  *  .**************    *  *.***********    *      *    *.***.

                              αF                       αG
HIPK1  358 YRAPEIILGLPFCEAIDMWSLGCVIAELFLGWPLYPGASEYDQIRYISQTQGLPAEYLLS 417
HIPK2  367 YRAPEIILGLPFCEAIDMWSLGCVIAELFLGWPLYPGASEYDQIRYISQTQGLPAEYLLS 426
HIPK3  365 YRAPEIILGLPFCEAIDMWSLGCVIAELFLGWPLYPGALEYDQIRYISQTQGLPGEQLLN 424
HIPK4  181 YRAPEILLGLPFCEKVDVWSLGCVMAELHLGWPLYPGNNEYDQVRYICETQGLPKPHLLH 240
           ******.*******   .*.******.***  ********   ****.***..*****    **

HIPK1  418 AGTKTTRFFNRDPNLGYP-LWRLKTPEEHELETGIKSKEARKYIFNCLDDMAQVN----- 471
HIPK2  427 AGTKTTRFFNRDTDSPYP-LWRLKTPDDHEAETGIKSKEARKYIFNCLDDMAQVN----- 480
HIPK3  425 VGTKSTRFFCKETDMSHS-GWRLKTLEEHEAETGMKSKEARKYIFNSLDDVAHVN----- 478
HIPK4  241 AACKAHHFFKRNPHPDAANPWQLKSSADYLAETKVRPLERRKYMLKSLDQIETVNGGSVA 300
             .*.  .**  .          *.**.    .     **   ..    *  ***.    .**   .    **

                                 αH                   αI
HIPK1  472 MSTDLEGTDMLAEKADRREYIDLLKKMLTIDADKRITPLKTLNHQFV 518
HIPK2  481 MTTDLEGSDMLVEKADRREFIDLLKKMLTIDADKRITPIETLNHPFV 527
HIPK3  479 TVMDLEGSDLLAEKADRREFVSLLKKMLLIDADLRITPAETLNHPFV 525
HIPK4  301 SRLTFPDREALAEHADLKSMVELIKRMLTWESHERISPSAALRHPFV 347
                  .  *  *.**   .   .  *.*.**    ..    **.*    .*  *  **
```

**Figure S1. Sequence alignment of HIPK family kinase domains.** Primary sequences of the kinase domains in HIPK1-4, highlighting canonical structural elements. Asterisks and periods indicate fully conserved and similar residues, respectively. Contiguous HIPK4-specific sequences primarily reside in the αC helix, β4 and β5 strands, and distal regions in the C-terminal lobe (residues 250-260 and 296-308).

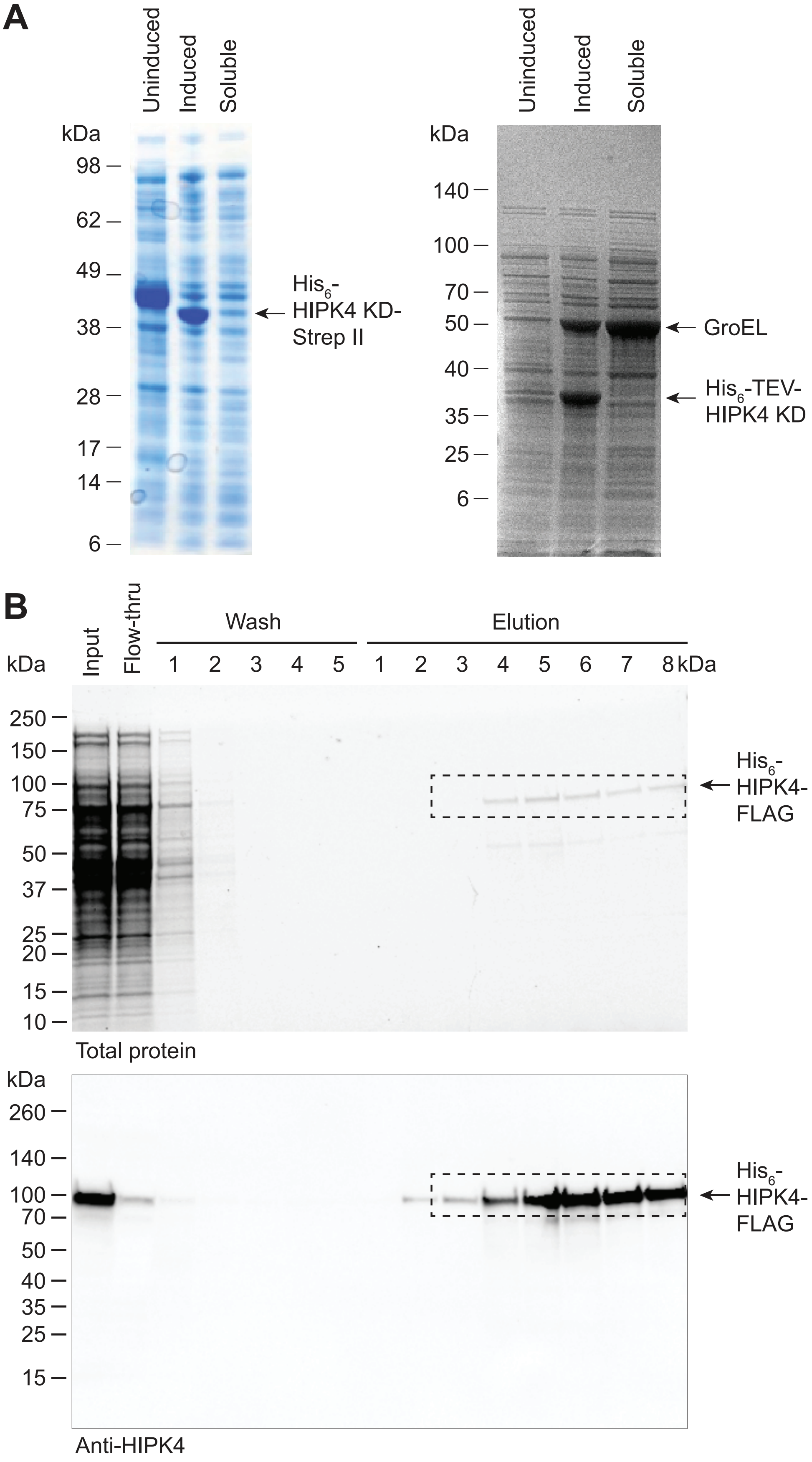

**Figure S2. Expression of HIPK4 constructs in bacteria and insect cells.** (A) SDS-PAGE analyses of *E. coli* expressing the HIPK4 kinase domain, demonstrating the insolubility of this recombinant protein even when co-expressed with chaperones such as GroEL/ES. (B) SDS-PAGE and western blot analyses of Sf9 cell expression and two-step His$_6$//FLAG tag purification of full-length HIPK4 (dashed boxes) using transient transfection. Fractions associated with the final FLAG affinity purification and elution with FLAG peptide are shown.

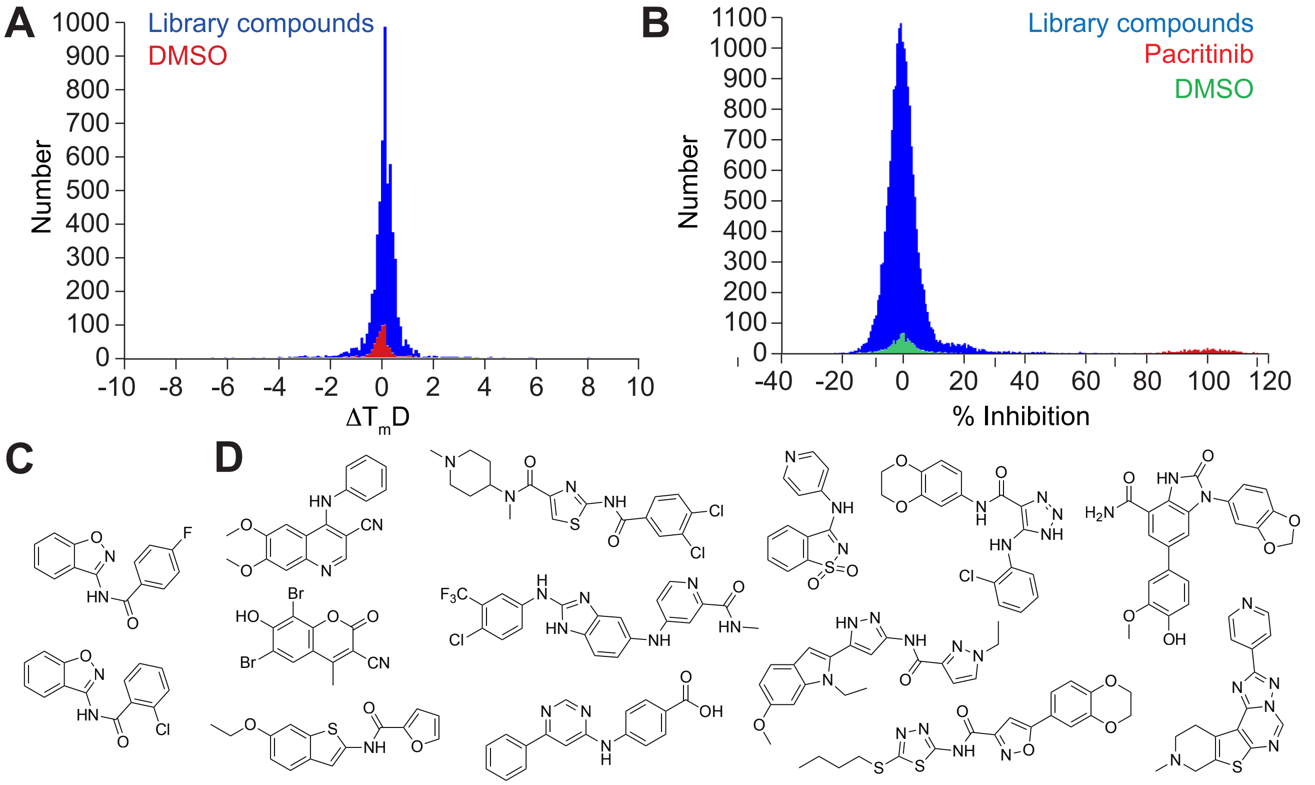

**Figure S3. A high-throughput screen for HIPK4 inhibitors.** (A) Representative screening results using a protein thermal shift (PTS) assay. HIPK4 ΔPEST was used as the the protein target, and the distribution of $\Delta T_m D$ values ($\Delta T_m$ determined from the first derivative of the melt curve) for 2,240 compounds and DMSO controls are shown. The complete PTS-based screen surveyed 7,500 compounds. (B) Screening results using the ADP-Glo assay. The distribution of inhibitory activities for 18,500 compounds against mMBP-HIPK4 ΔPEST fusion protein are shown, using 5 μM pacritinib and DMSO alone as positive and negative controls, respectively. (C) 2 compounds of the same chemotype were validated as non ATP-competitive HIPK4 inhibitors. (D) 12 compounds with distinct scaffolds were are validated as ATP-competitive HIPK4 inhibitors, with the cyanoquinoline emerging as the most potent hit.

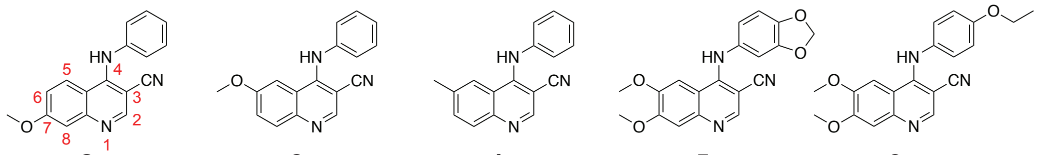

**2** **3** **4** **5** **6**

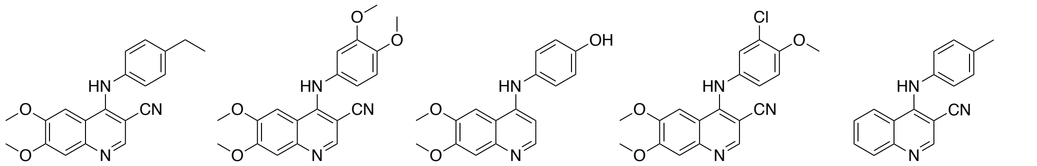

**7** **8** **9** **10** **11**

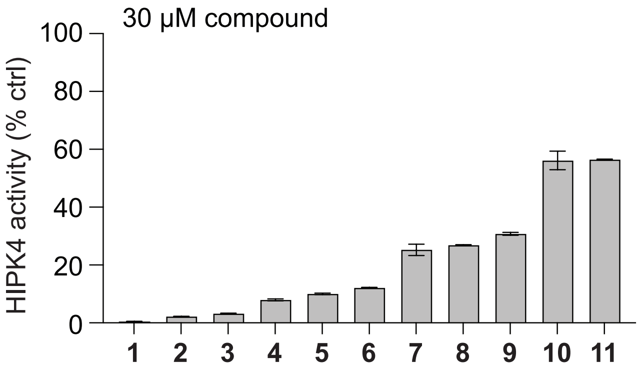

**Figure S4. Activities of commercially available quinoline derivatives against recombinant full-length HIPK4.** Individual compounds were evaluated at a 30-µM concentration in the ADP-Glo assay using 10 µM ATP and GST-HIPK4. The numbering scheme for the quinoline scaffold is also shown. Data are the average of two biological replicates ± s.e.m.

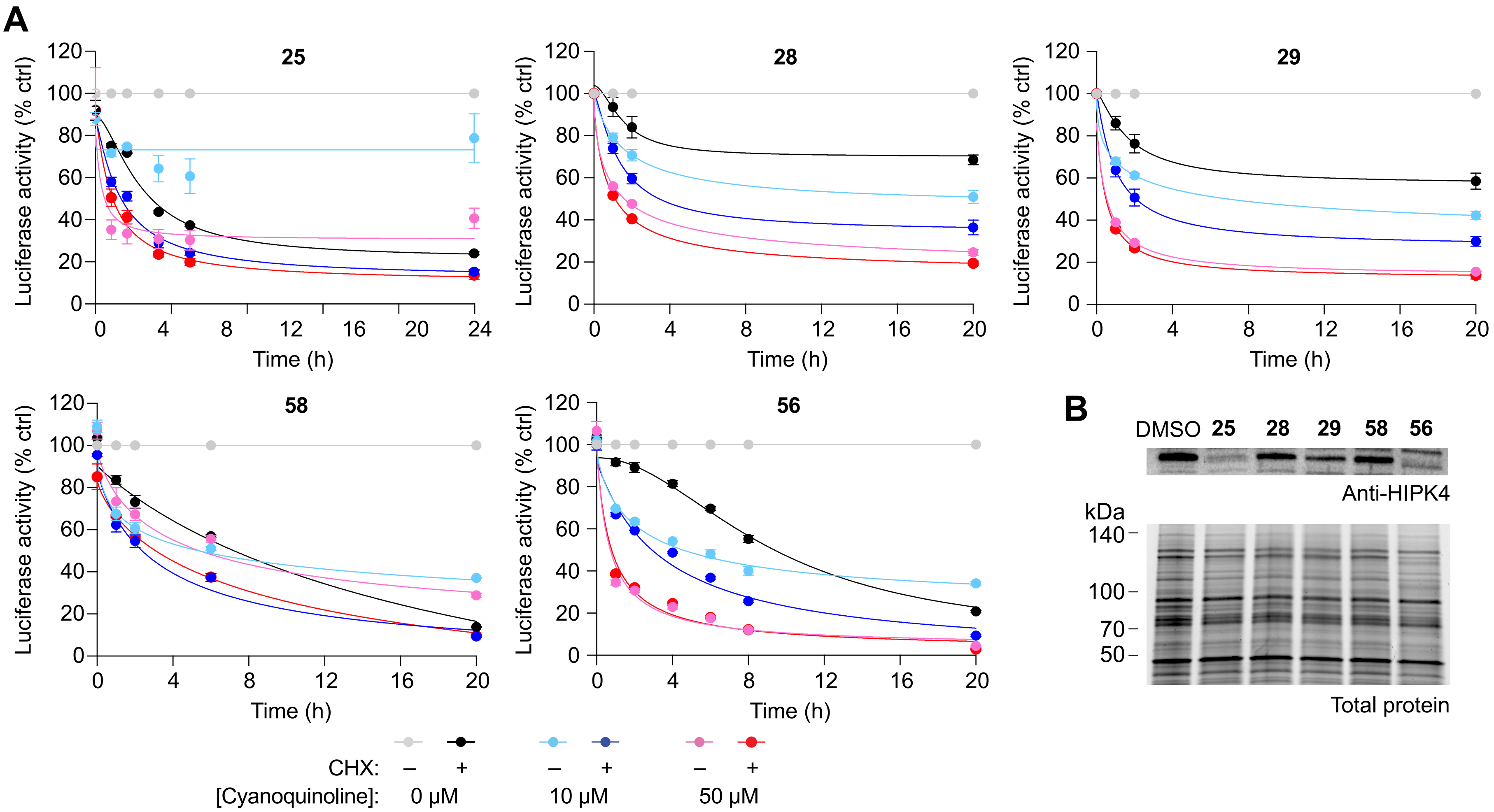

**Figure S5. Activities of repensentative cyanoquinoline analogs in the HiBiT-HIPK4 assay and in spermatids.** (A) HiBiT-HIPK4- and LgBiT-expressing HEK293T cells were cultured in the absence or presence of cycloheximide (CHX; 10 µg/mL) to discern the effects of individual cyanoquinolines on HIPK4 synthesis and degradation. Data are the average of two biological replicates ± s.e.m. (B) Immunoblots of detergent-solubilized spermatid lysates after culturing the germ cells with representative cyanoquinolines (50 µM) for 16 h.

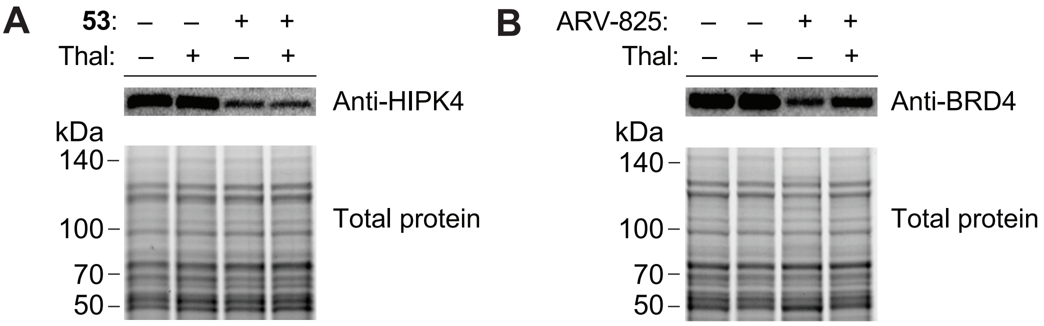

**Figure S6. Thalidomide competition assays in spermatids.** (A) Western blot analyses of HIPK4 levels in spermatids treated with candidate HIPK4 PROTAC **53** in the absence or presence of 20 μM thalidomide (Thal). (B) Western blot analyses of BRD4 levels in spermatids treated with the BRD4 PROTAC ARV-825 in the absence or presence of 20 μM thalidomide.

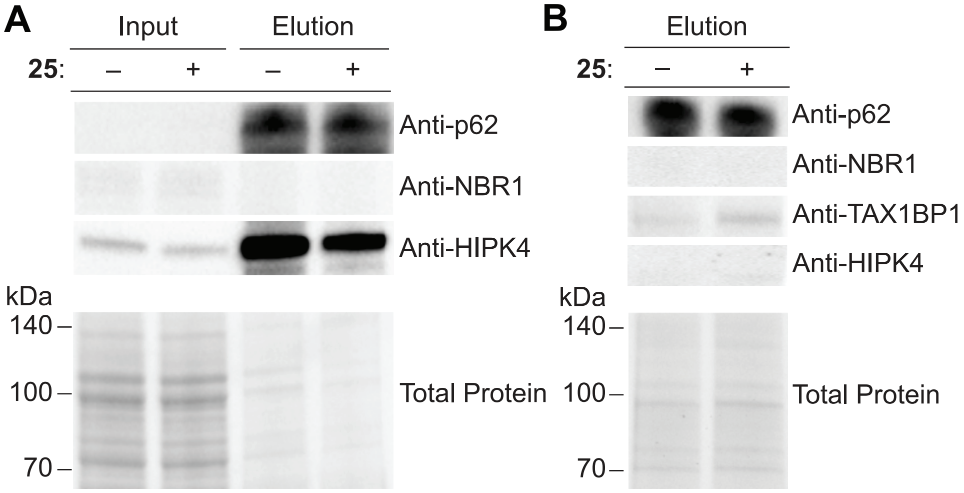

**Figure S7. Co-immunoprecipitation studies of HIPK4, NBR1, and SQSTM1/p62.** (A) Spermatids treated with cyanoquinoline **25** or DMSO alone for 5 h were lysed, and HIPK4 was immunoprecipitated. (B) Immunoprecipitation experiments with rabbit IgG were conducted in parallel as comparison controls. Input and elution fractions for the two immunoprecipitation conditions were immunoblotted with anti-p62, anti-NBR1, anti-TAX1BP1, or anti-HIPK4 antibod-

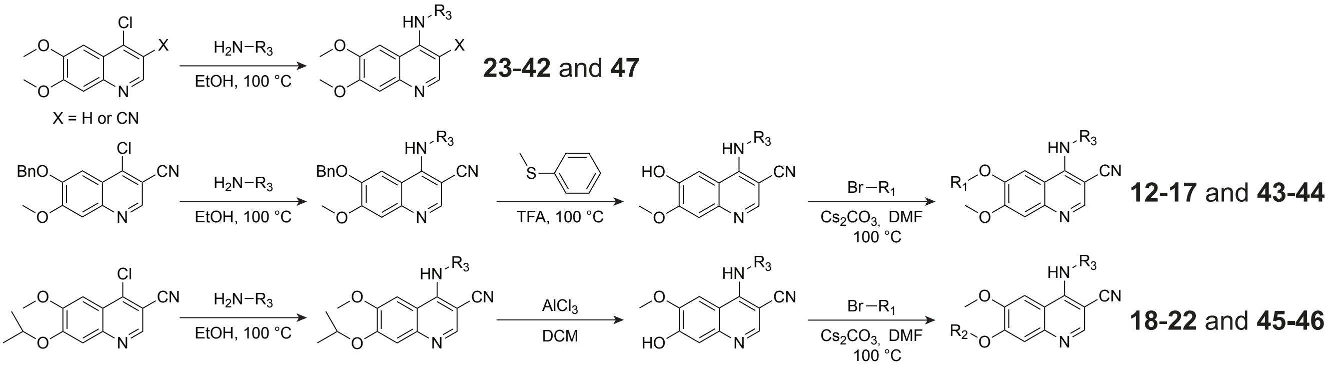

**Scheme S1. General synthetic schemes for quinoline scaffolds.** Compound **42** was prepared using a des-cyano 4-chloroquinoline, and all other analogs were generated with C3-cyano-functionalized starting materials.

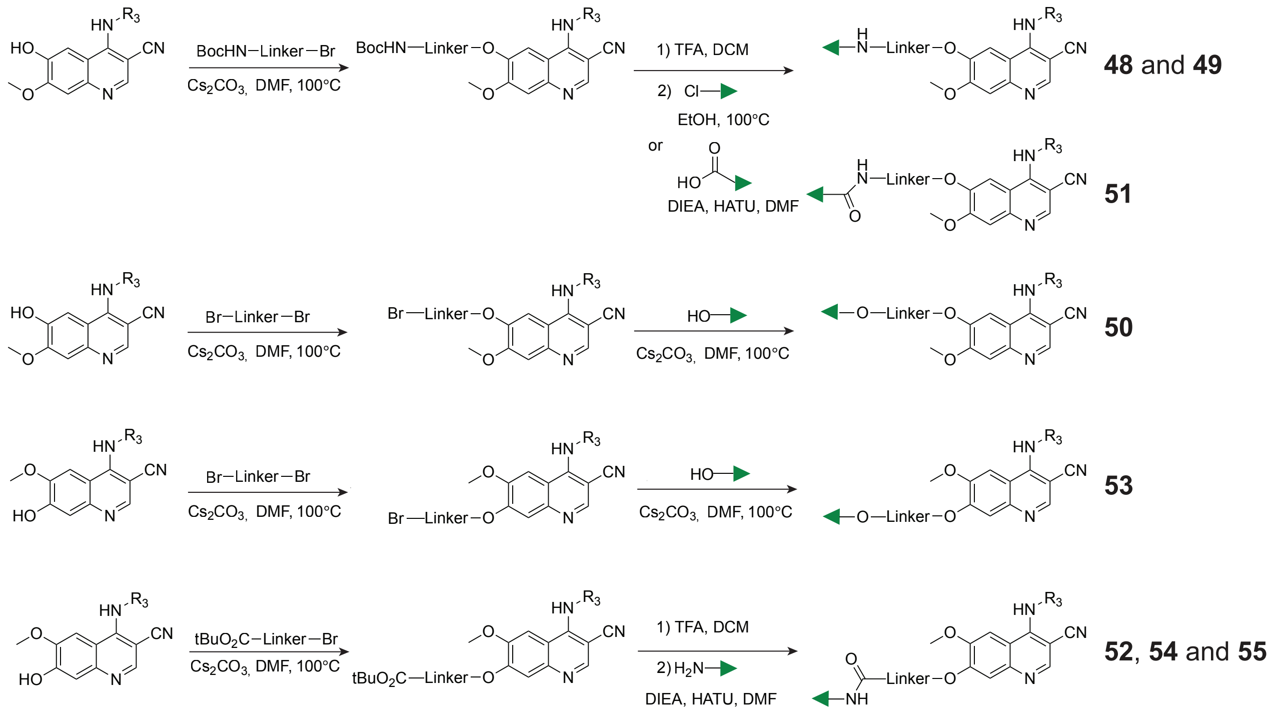

**Scheme S2. General synthetic schemes for candidate HIPK4 PROTACs.** All analogs were generated with C6 or C7 hydroxyl cyanoquinolines. The E3 ligase receptor ligand is depicted by the green triangle.

#### SYNTHETIC METHODS

##### 6-hydroxy-7-methoxy-4-(phenylamino)quinoline-3-carbonitrile (12)

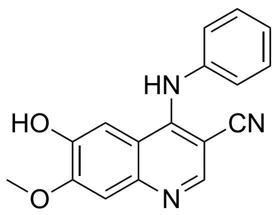

A mixture of 6-(benzyloxy)-4-chloro-7-methoxyquinoline-3-carbonitrile (200 mg, 0.62 mmol, 1.0 equiv) and aniline (63 mg, 62 μL, 0.68 mmol, 1.1 equiv) in EtOH (5 mL) was heated at 100 ºC for 18 h. After completion of the reaction, the mixture was concentrated under reduced pressure, and the residue was purified by flash column chromatography (EtOAc/hexanes, then 1–5% MeOH in EtOAc) to afford 6-(benzyloxy)-7-methoxy-4-(phenylamino)quinoline-3-carbonitrile as the product (195 mg) as a solid followed by treatment of thioanisole (770 mg, 0.73 mL, 6.20 mmol, 10 equiv) and trifluoroacetic acid (TFA) (700 mg, 0.48 mL, 6.20 mmol, 10 equiv). The mixture was refluxed at 80 ºC for 2 h. The solvent was removed and the residue was stirred with ice water and $NH_4OH$ was slowly added until pH reached 10. The reaction mixture was filtered and the solids were washes with EtOAc to give the compound **12** (120 mg, 67% yield) as a yellow solid. $^1$H NMR (500 MHz, DMSO-$d_6$) δ 9.32 (br s, 1H), 8.45 (s, 1H), 7.60 (s, 1H), 7.36 – 7.32 (m, 3H), 7.16 – 7.10 (m, 3H), 3.98 (s, 3H) ppm. $^{13}$C NMR (126 MHz, DMSO-$d_6$) δ 153.8, 150.3, 149.3, 147.8, 146.1, 141.3, 129.4, 124.4, 122.4, 117.8, 115.5, 109.2, 105.9, 89.9, 56.4 ppm. HRMS (ESI) m/z calcd for $C_{17}H_{13}N_3O_2$ [M + H]$^+$, 292.1081; found, 292.1075.

**6-(benzyloxy)-7-methoxy-4-(phenylamino)quinoline-3-carbonitrile (13)**

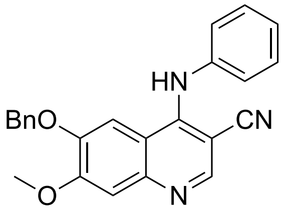

A mixture of 6-(benzyloxy)-4-chloro-7-methoxyquinoline-3-carbonitrile (200 mg, 0.62 mmol, 1.0 equiv) and aniline (63 mg, 62 μL, 0.68 mmol, 1.1 equiv) in EtOH (5 mL) was heated at 100 ºC for 18 h. After completion of the reaction, the mixture was concentrated under reduced pressure, and the residue was purified by flash column chromatography (EtOAc/hexanes, then 1–5% MeOH in EtOAc) to afford compound **13** as the product (195 mg, 83% yield) as a yellow solid. $^1$H NMR (500 MHz, $CDCl_3$) δ 8.64 (s, 1H), 7.40 – 7.36 (m, 3H), 7.34 – 7.28 (m, 3H), 7.07 (d, *J* = 7.7 Hz, 2H), 6.90 (s, 1H), 7.40 – 7.35 (m, 2H), 7.28 (s, 1H), 6.76 (s, 1H), 4.78 (s, 2H), 4.03 (s, 3H) ppm. $^{13}$C NMR (126 MHz, $CDCl_3$) δ 148.0, 142.9, 135.7, 135.7, 129.5, 129.5, 128.7, 128.7, 128.6, 128.2, 128.2, 127.2, 127.2, 125.3, 125.2, 122.4, 122.4, 104.9, 70.7, 56.3 ppm. HRMS (ESI) m/z calcd for $C_{24}H_{19}N_3O_2$ $[M + H]^+$, 382.1550; found, 382.1547.

**6-ethoxy-7-methoxy-4-(phenylamino)quinoline-3-carbonitrile (14)**

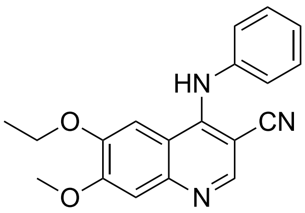

A mixture of compound **12** (40 mg, 0.14 mmol, 1 equiv) and $Cs_2CO_3$ (45 mg, 0.14 mmol, 1 equiv) in DMF (5 mL) was stirred at room temperature for 30 min, after which bromoethane (22 mg, 0.21 mmol, 1.5 equiv) was added. The mixture was then heated at 100 °C for 2 h. After cooling to room temperature, the reaction was quenched with water (5 mL) and extracted with EtOAc (3 × 5 mL). The combined organic extracts were concentrated under reduced pressure, and the residue was purified by flash column chromatography on silica gel (50% EtOAc in Hexane) to afford compound **14** as the product (35 mg, 80% yield) as a white solid. $^1$H NMR (500 MHz, DMSO-$d_6$) δ 9.46 (s, 1H), 8.19 (s, 1H), 7.72 (s, 1H), 7.38 – 6.92 (m, 6H), 4.09 (q, $J$ = 7.0 Hz, 2H), 3.90 (s, 3H), 1.38 (t, $J$ = 6.9 Hz, 3H). $^{13}$C NMR (126 MHz, DMSO-$d_6$) δ 154.0, 151.4, 149.9, 149.0, 146.5, 129.5, 125.4, 124.1, 117.8, 114.3, 109.2, 103.0, 88.4, 70.6, 56.4, 22.4, 11.0 ppm. HRMS (ESI) m/z calcd for $C_{19}H_{17}N_3O_2$ [M + H]$^+$, 320.1394; found, 320.1390.

**7-methoxy-4-(phenylamino)-6-propoxyquinoline-3-carbonitrile (15)**

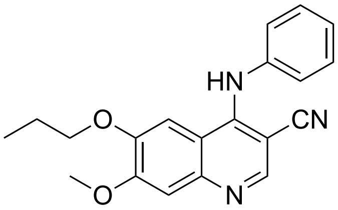

A mixture of compound **12** (40 mg, 0.14 mmol, 1 equiv) and $Cs_2CO_3$ (45 mg, 0.14 mmol, 1 equiv) in DMF (5 mL) was stirred at room temperature for 30 min, after which bromopropane (25 mg, 0.21 mmol, 1.5 equiv) was added. The mixture was then heated at 100 °C for 2 h. After cooling to room temperature, the reaction was quenched with water (5 mL) and extracted with EtOAc (3 × 5 mL). The combined organic extracts were concentrated under reduced pressure, and the residue was purified by flash column chromatography on silica gel (50% EtOAc in Hexane) to afford compound **15** as the product (33 mg, 72% yield) as a white solid. $^1$H NMR (500 MHz, $CDCl_3$) δ 8.64 (s, 1H), 8.43 (br s, 1H), 7.39 – 7.35 (m, 3H), 7.22 (t, $J$ = 7.5 Hz, 1H), 7.11 (d, $J$ = 7.1 Hz, 2H), 4.01 (s, 3H), 3.63 (t, $J$ = 6.9 Hz, 2H), 1.71 – 1.65 (m, 2H), 0.92 (t, $J$ = 7.4 Hz, 3H) ppm. $^{13}$C NMR (126 MHz, $CDCl_3$) δ 154.3, 149.5, 149.4, 148.5, 147.6, 141.1, 129.5, 125.4, 122.8, 117.1, 113.6, 109.0, 103.6, 92.3, 70.2, 56.2, 21.8, 10.2 ppm. HRMS (ESI) m/z calcd for $C_{20}H_{19}N_3O_2$ [M + H]$^+$, 334.1550; found, 334.1545.

**6-isopropoxy-7-methoxy-4-(phenylamino)quinoline-3-carbonitrile (16)**

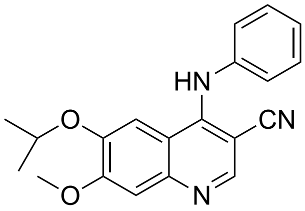

A mixture of compound **12** (40 mg, 0.14 mmol, 1 equiv) and $Cs_2CO_3$ (45 mg, 0.14 mmol, 1 equiv) in DMF (5 mL) was stirred at room temperature for 30 min, after which 2-bromopropane (25 mg, 0.21 mmol, 1.5 equiv) was added. The mixture was then heated at 100 °C for 2 h. After cooling to room temperature, the reaction was quenched with water (5 mL) and extracted with EtOAc (3 × 5 mL). The combined organic extracts were concentrated under reduced pressure, and the residue was purified by flash column chromatography on silica gel (50% EtOAc in Hexane) to afford compound **16** as the product (36 mg, 79% yield) as a white solid. $^1$H NMR (500 MHz, $CDCl_3$) δ 8.64 (s, 1H), 7.37 – 7.33 (m, 3H), 7.22 (t, *J* = 7.5 Hz, 1H), 7.09 (d, *J* = 7.1 Hz, 2H), 6.84 (s, 1H), 6.79 (s, 1H), 4.13 – 4.09 (m, 1H), 4.00 (s, 3H), 1.17 (d, *J* = 6.0 Hz, 6H) ppm. $^{13}$C NMR (126 MHz, $CDCl_3$) δ 154.9, 149.6, 149.3, 147.6, 147.3, 141.2, 129.7, 125.3, 122.6, 117.1, 109.2, 105.4, 92.5, 71.4, 56.2, 21.6 ppm. HRMS (ESI) m/z calcd for $C_{20}H_{19}N_3O_2$ $[M + H]^+$, 334.1550; found, 334.1546.

**6-butoxy-7-methoxy-4-(phenylamino)quinoline-3-carbonitrile (17)**

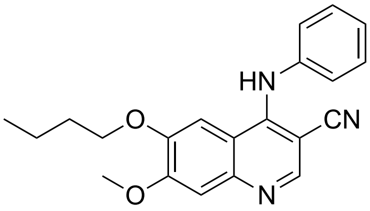

A mixture of compound **12** (40 mg, 0.14 mmol, 1 equiv) and $Cs_2CO_3$ (45 mg, 0.14 mmol, 1 equiv) in DMF (5 mL) was stirred at room temperature for 30 min, after which 2-bromobutane (28 mg, 0.21 mmol, 1.5 equiv) was added. The mixture was then heated at 100 °C for 2 h. After cooling to room temperature, the reaction was quenched with water (5 mL) and extracted with EtOAc

(3 × 5 mL). The combined organic extracts were concentrated under reduced pressure, and the residue was purified by flash column chromatography on silica gel (50% EtOAc in Hexane) to afford compound **17** as the product (39 mg, 82% yield) as a white solid. $^1$H NMR (500 MHz, $CDCl_3$) δ 8.64 (s, 1H), 7.38 – 7.35 (m, 3H), 7.22 (t, *J* = 7.5 Hz, 1H), 7.11 – 7.09 (m, 2H), 6.83 – 6.81 (m, 2H), 4.01 (s, 3H), 3.66 (t, *J* = 6.9 Hz, 2H), 1.69 – 1.63 (m, 2H), 1.39 – 1.32 (m, 2H), 0.92 (t, *J* = 7.4 Hz, 3H) ppm. $^{13}$C NMR (126 MHz, $CDCl_3$) δ 154.3, 149.5, 149.4, 148.5, 147.6, 141.1, 129.5, 125.3, 122.8, 117.1, 113.6, 109.0, 103.6, 92.3, 76.8, 68.5, 56.2, 30.5, 19.0, 13.7 ppm. HRMS (ESI) m/z calcd for $C_{21}H_{21}N_3O_2$ $[M + H]^+$, 348.1707; found, 348.1705.

**7-hydroxy-6-methoxy-4-(phenylamino)quinoline-3-carbonitrile (18)**

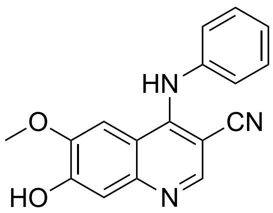

A mixture of 4-chloro-7-isopropoxy-6-methoxyquinoline-3-carbonitrile (200 mg, 0.72 mmol, 1.0 equiv) and aniline (74 mg, 72 µL, 0.79 mmol, 1.1 equiv) in EtOH (5 mL) was heated at 100 ºC for 18 h. After completion of the reaction, the mixture was concentrated under reduced pressure, and the residue was purified by flash column chromatography (EtOAc/hexanes, then 1–5% MeOH in EtOAc) to afford 7-isopropoxy-6-methoxy-4-(phenylamino)quinoline-3-carbonitrile as the product (180 mg) as a white solid followed by treatment of $AlCl_3$ (145 mg, 1.10 mmol, 1.5 equiv) in anhydrous DCM (10 mL). The mixture was refluxed at room temperature for 2 h. The reaction mixture was filtered and the solids were washes with 0.1 N HCl and $H_2O$ to give the compound **18** (130 mg, 62% yield) as a yellow solid. $^1$H NMR (500 MHz, DMSO-$d_6$) δ 10.53 (br s, 1H), 9.56 (br s, 1H), 8.43 (s, 1H), 7.78 (s, 1H), 7.42 – 7.39 (m 2H), 7.23 (dd, *J* = 16.4, 7.4 Hz, 4H), 3.92 (s, 3H) ppm. $^{13}$C NMR (126 MHz, DMSO-$d_6$) δ 152.7, 151.1, 150.0, 149.5, 140.7, 129.5, 125.4, 124.0,

117.8, 113.5, 102.8, 88.0, 56.5, 22.0 ppm. HRMS (ESI) m/z calcd for $C_{17}H_{13}N_3O_2$ $[M + H]^+$, 292.1081; found, 292.1078.

**7-ethoxy-6-methoxy-4-(phenylamino)quinoline-3-carbonitrile (19)**

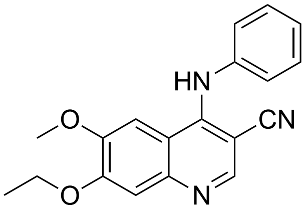

A mixture of compound **18** (40 mg, 0.14 mmol, 1 equiv) and $Cs_2CO_3$ (45 mg, 0.14 mmol, 1 equiv) in DMF (5 mL) was stirred at room temperature for 30 min, after which bromoethane (22 mg, 0.21 mmol, 1.5 equiv) was added. The mixture was then heated at 100 °C for 2 h. After cooling to room temperature, the reaction was quenched with water (5 mL) and extracted with EtOAc (3 × 5 mL). The combined organic extracts were concentrated under reduced pressure, and the residue was purified by flash column chromatography on silica gel (50% EtOAc in Hexane) to afford compound **19** as the product (37 mg, 84% yield) as a white solid. $^1$H NMR (500 MHz, DMSO-$d_6$) δ 9.48 (s, 1H), 8.43 (br s, 1H), 7.74 (s, 1H), 7.41 (t, $J$ = 7.7 Hz, 2H), 7.32 (s, 1H), 7.24 – 7.20 (m, 3H), 4.14 (q, $J$ = 6.9, 7.0 Hz, 2H), 3.91 (s, 3H), 1.44 (t, $J$ = 6.9 Hz, 3H) ppm. $^{13}$C NMR (126 MHz, DMSO-$d_6$) δ 153.2, 151.3, 149.8, 149.7, 146.6, 132.3, 129.5, 125.3, 124.1, 117.7, 114.0, 109.7, 102.3, 88.5, 64.6, 56.5, 14.9 ppm. HRMS (ESI) m/z calcd for $C_{19}H_{17}N_3O_2$ $[M + H]^+$, 320.1394; found, 320.1388.

**6-methoxy-4-(phenylamino)-7-propoxyquinoline-3-carbonitrile (20)**

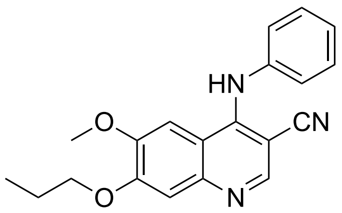

A mixture of compound **18** (40 mg, 0.14 mmol, 1 equiv) and $Cs_2CO_3$ (45 mg, 0.14 mmol, 1 equiv) in DMF (5 mL) was stirred at room temperature for 30 min, after which 2-bromopropane (25 mg, 0.21 mmol, 1.5 equiv) was added. The mixture was then heated at 100 °C for 2 h. After cooling to room temperature, the reaction was quenched with water (5 mL) and extracted with EtOAc (3 × 5 mL). The combined organic extracts were concentrated under reduced pressure, and the residue was purified by flash column chromatography on silica gel (50% EtOAc in Hexane) to afford compound **20** as the product (41 mg, 90% yield) as a white solid. $^{1}$H NMR (500 MHz, DMSO-$d_6$) δ 8.47 (s, 1H), 8.45 (s, 1H), 7.74 (s, 1H), 7.42 – 7.39(m, 2H), 7.33 (s, 1H), 7.26 – 7.19 (m, 3H), 6.53 (s, 1H), 4.14 (t, *J* = 6.6 Hz, 2H), 3.91 (s, 3H), 1.86 – 1.79 (m, 2H),1.04 (t, *J* = 1.1 Hz, 3H) ppm. $^{13}$C NMR (126 MHz, DMSO-$d_6$) δ 153.4, 151.3, 149.8, 146.6, 140.6, 129.5, 125.5, 124.0, 114.0, 112.5, 112.4, 112.3, 109.7, 102.4, 70.3, 61.6, 56.5, 22.3, 10.9 ppm. HRMS (ESI) m/z calcd for $C_{20}H_{19}N_3O_2$ [M + H]$^+$, 334.1550; found, 334.1547.

**7-isopropoxy-6-methoxy-4-(phenylamino)quinoline-3-carbonitrile (21)**

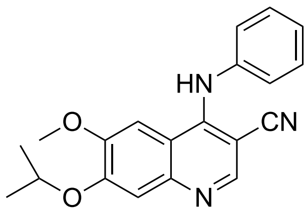

A mixture of 4-chloro-7-isopropoxy-6-methoxyquinoline-3-carbonitrile (200 mg, 0.72 mmol, 1.0 equiv) and aniline (74 mg, 72 µL, 0.79 mmol, 1.1 equiv) in EtOH (5 mL) was heated at 100 ºC for 18 h. After completion of the reaction, the mixture was concentrated under reduced pressure, and the residue was purified by flash column chromatography (EtOAc/hexanes, then 1–5% MeOH in EtOAc) to afford compound **21** as the product (180 mg, 75%) as a white solid. $^1$H NMR (500 MHz, DMSO-$d_6$) δ 10.95 (br s, 1H), 8.94 (s, 1H), 8.08 (s, 1H), 7.53 – 7.50 (m, 2H), 7.48 (s, 1H), 7.46-7.41 (m, 3H), 4.85 – 4.81 (m, 1H), 3.07 (s, 3H), 1.42 (d, $J$ = 6.0 Hz, 6H) ppm. $^{13}$C NMR (126 MHz, DMSO-$d_6$) δ 153.9, 152.8, 152.8, 151.1, 148.1, 148.1, 138.1, 129.8, 128.4, 126.7, 112.8, 103.9, 86.6, 72.2, 57.1, 21.8 ppm. HRMS (ESI) m/z calcd for $C_{20}H_{19}N_3O_2$ [M + H]$^+$, 334.1550; found, 334.1544.

**7-butoxy-6-methoxy-4-(phenylamino)quinoline-3-carbonitrile (22)**

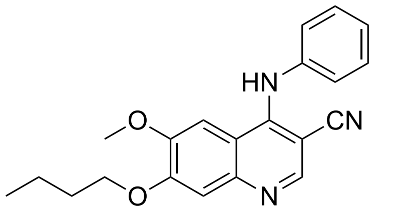

A mixture of compound **18** (40 mg, 0.14 mmol, 1 equiv) and $Cs_2CO_3$ (45 mg, 0.14 mmol, 1 equiv) in DMF (5 mL) was stirred at room temperature for 30 min, after which 2-bromobutane (28 mg, 0.21 mmol, 1.5 equiv) was added. The mixture was then heated at 100 °C for 2 h. After cooling to room temperature, the reaction was quenched with water (5 mL) and extracted with EtOAc (3 × 5 mL). The combined organic extracts were concentrated under reduced pressure, and the residue was purified by flash column chromatography on silica gel (50% EtOAc in Hexane) to afford compound **22** as the product (35 mg, 73% yield) as a white solid. $^1$H NMR (500 MHz, DMSO-$d_6$) δ 9.47 (s, 1H), 8.45 (s, 1H), 7.74 (s, 1H), 7.42 (t, $J$ = 7.7 Hz, 2H), 7.34 (s, 1H), 7.26 – 7.20 (m, 3H), 4.18 (t, $J$ = 6.5 Hz, 2H), 3.91 (s, 3H), 1.82 – 1.77 (m, 2H), 1.53 – 1.47 (m, 2H), 0.99 (t, $J$ = 7.4 Hz, 3H) ppm. $^{13}$C NMR (126 MHz, $CDCl_3$) δ 153.5, 151.9, 150.6, 149.4, 145.7, 143.8, 128.8, 127.7, 126.7, 119.8, 111.7, 109.7, 98.7, 84.1, 56.3, 56.2, 49.2, 44.1, 36.1 ppm. HRMS (ESI) m/z calcd for $C_{21}H_{21}N_3O_2$ [M + H]$^+$, 348.1707; found, 348.1703.

**4-((2-chlorophenyl)amino)-6,7-dimethoxyquinoline-3-carbonitrile (23)**

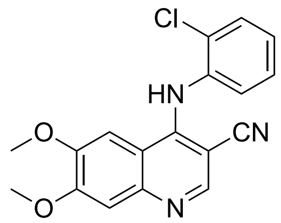

A mixture of 4-chloro-6,7-dimethoxyquinoline-3-carbonitrile (200 mg, 0.80 mmol, 1.0 equiv) and 2-chloroaniline (113 mg, 93 µL, 0.89 mmol, 1.1 equiv) in EtOH (15 mL) was heated at 100 ºC for 18 h. After completion of the reaction, the mixture was concentrated under reduced pressure, and the residue was purified by flash column chromatography (EtOAc/hexanes, then 1–5% MeOH in EtOAc) to afford compound **23** as the product (255 mg, 93% yield) as a yellow solid. $^{1}H$ NMR (500 MHz, DMSO-$d_6$) δ 9.93 (br s, 1H), 8.50 (s, 1H), 7.85 (s, 1H), 7.58 – 7.54 (m, 1H), 7.48 – 7.45 (m, 1H), 7.40 – 7.35 (m, 2H), 7.28 (s, 1H), 3.90 (s, 3H), 3.88 (s, 3H) ppm. $^{13}C$ NMR (126 MHz, DMSO-$d_6$) δ 174.5, 154.6, 151.3, 150.1, 136.1, 132.3, 130.4, 130.3, 130.0, 128.6, 116.2, 112.8, 106.7, 102.7, 86.4, 56.9, 56.6 ppm. HRMS (ESI) m/z calcd for $C_{18}H_{14}ClN_3O_2$ [M + H]$^+$, 340.0847; found, 340.0843.

**4-((3-chlorophenyl)amino)-6,7-dimethoxyquinoline-3-carbonitrile (24)**

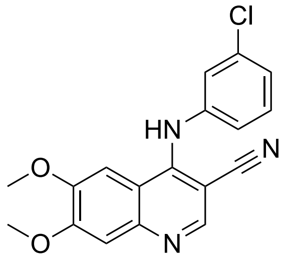

A mixture of 4-chloro-6,7-dimethoxyquinoline-3-carbonitrile (200 mg, 0.80 mmol, 1.0 equiv) and 3-chloroaniline (113 mg, 94 µL, 0.89 mmol, 1.1 equiv) in EtOH (15 mL) was heated at 100 ºC for 18 h. After completion of the reaction, the mixture was concentrated under reduced pressure, and the residue was purified by flash column chromatography (EtOAc/hexanes, then 1–5% MeOH in

EtOAc) to afford compound **24** as the product (260 mg, 95% yield) as a yellow solid. $^{1}H$ NMR (500 MHz, $CDCl_3$) δ 8.69 (s, 1H), 7.40 (s, 1H), 7.30 (t, $J$ = 7.2 HZ, 1H), 7.16 – 7.14 (m, 1H), 7.06 (t, $J$ = 2.1 Hz, 1H), 6.95 – 6.92 (m, 1H), 6.84 (s, 1H), 6.78 (s, 1H), 4.04 (s, 3H), 3.66 (s, 3H) ppm. $^{13}C$ NMR (126 MHz, $CDCl_3$): δ 154.3, 149.5, 149.5, 148.5, 147.9, 142.3, 135.2, 130.4, 124.8, 121.6, 119.6, 116.7, 114.2, 109.1, 102.3, 93.8, 56.4, 55.9 ppm. HRMS (ESI) m/z calcd for $C_{18}H_{14}ClN_3O_2$ $[M + H]^+$, 340.0847; found, 340.0847.

**4-((4-chlorophenyl)amino)-6,7-dimethoxyquinoline-3-carbonitrile (25)**

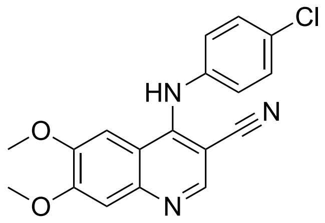

A mixture of 4-chloro-6,7-dimethoxyquinoline-3-carbonitrile (200 mg, 0.80 mmol, 1.0 equiv) and 4-chloroaniline (113 mg, 0.89 mmol, 1.1 equiv) in EtOH (15 mL) was heated at 100 ºC for 18 h. After completion of the reaction, the mixture was concentrated under reduced pressure, and the residue was purified by flash column chromatography (EtOAc/hexanes, then 1–5% MeOH in EtOAc) to afford compound **25** as the product (257 mg, 94% yield) as a yellow solid. $^{1}H$ NMR (500 MHz, DMSO-$d_6$) δ 10.82 (br s, 1H), 8.92 (s, 1H), 8.06 (s, 1H), 7.57 (d, $J$ = 8.7 HZ, 2H), 7.47 (d, $J$ = 7.5 Hz, 2H), 7.43 (s, 1H), 4.00 (s, 3H), 3.99 (s, 3H) ppm. $^{13}C$ NMR (126 MHz, DMSO-$d_6$) δ 155.8, 152.9, 150.6, 148.0, 137.3, 132.2, 129.6, 129.4, 128.3, 115.0, 113.5, 104.1, 102.5, 87.0, 57.3, 56.9 ppm. HRMS (ESI) m/z calcd for $C_{18}H_{14}ClN_3O_2$ $[M + H]^+$, 340.0847; found, 340.0842.

**4-((3,4-Dichlorophenyl)amino)-6,7-dimethoxyquinoline-3-carbonitrile (26)**

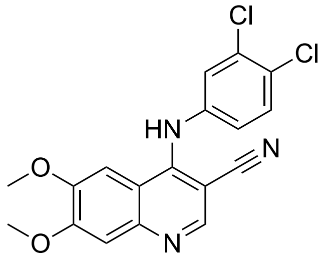

A mixture of 4-chloro-6,7-dimethoxyquinoline-3-carbonitrile (200 mg, 0.80 mmol, 1.0 equiv) and 3,4-dichloroaniline (143 mg, 0.89 mmol, 1.1 equiv) in EtOH (15 mL) was heated at 100 ºC for 18 h. After completion of the reaction, the mixture was concentrated under reduced pressure, and the residue was purified by flash column chromatography (EtOAc/hexanes, then 1–5% MeOH in EtOAc) to afford compound **26** (268 mg, 89% yield) as a yellow solid. $^{1}H$ NMR (500 MHz, DMSO-$d_6$) δ 9.82 (br s, 1H), 8.57 (s, 1H), 7.78 (s, 1H), 7.61 (d, $J$ = 8.7 Hz, 1H), 7.59 (d, $J$ = 2.6 HZ, 1H), 7.38 (s, 1H), 7.23 (dd, $J$ = 2.6, 8.7 Hz, 1H), 3.97 (s, 3H), 3.94 (s, 3H) ppm. 13C NMR (126 MHz, DMSO-$d_6$) δ 154.3, 151.0, 150.0, 149.0, 147.0, 141.5, 131.7, 131.1, 125.9, 123.9, 122.5, 117.7, 115.0, 109.1, 102.5, 90.5, 56.6, 56.4 ppm. HRMS (ESI) m/z calcd for $C_{18}H_{13}Cl_2N_3O_2$ $[M + H]^+$, 374.0458; found, 374.0455.

**4-((2,4-Dichloro-5-methoxyphenyl)amino)-6,7-dimethoxyquinoline-3-carbonitrile (27)**

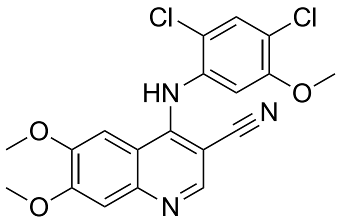

A mixture of 4-chloro-6,7-dimethoxyquinoline-3-carbonitrile (200 mg, 0.80 mmol, 1.0 equiv) and 2,4-dichloro-5-methoxyaniline (170 mg, 0.89 mmol, 1.1 equiv) in EtOH (15 mL) was heated at 100 ºC for 18 h. After completion of the reaction, the mixture was concentrated under reduced pressure, and the residue was purified by flash column chromatography (EtOAc/hexanes, then

1–5% MeOH in EtOAc) to afford compound **27** (280 mg, 93% yield) as a yellow solid. $^{1}$H NMR (500 MHz, DMSO-$d_6$) δ 9.72 (s, 1H), 8.35 (s, 1H), 7.83 (s, 1H), 7.69 (s, 1H), 7.29 (s, 1H), 7.24 (s, 1H), 3.94 (s, 3H), 3.93 (s, 3H), 3.85 (s, 3H) ppm. $^{13}$C NMR (126 MHz, DMSO-$d_6$) δ 154.3, 153.5, 151.7, 150.6, 149.2, 145.7, 130.0, 124.7, 122.6, 118.6, 112.8, 108.9, 104.0, 102.9, 102.8, 85.9, 57.1, 56.5, 56.3 ppm. HRMS (ESI) m/z calcd for $C_{19}H_{15}Cl_2N_3O_3$ [M + H]$^+$, 404.0563; found, 404.0559.

**4-((2-Methoxyphenyl)amino)-6,7-dimethoxyquinoline-3-carbonitrile (28)**

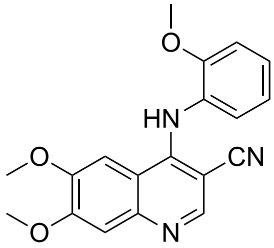

A mixture of 4-chloro-6,7-dimethoxyquinoline-3-carbonitrile (200 mg, 0.80 mmol, 1.0 equiv) and 2-methoxyaniline (98 mg, 92 µL, 0.89 mmol, 1.1 equiv) in EtOH (15 mL) was heated at 100 ºC for 18 h. After completion of the reaction, the mixture was concentrated under reduced pressure, and the residue was purified by flash column chromatography (EtOAc/hexanes, then 1–5% MeOH in EtOAc) to afford compound **28** (230 mg, 90% yield) as a white solid. $^{1}$H NMR (500 MHz, $CDCl_3$) δ 8.64 (s, 1H), 7.38 (s, 1H), 7.17 – 7.13 (m, 1H), 7.01 (dd, $J$ = 1.9, 7.0 Hz, 1H), 6.97 (s, 1H), 6.95 – 6.89 (m, 1H), 6.84 (s, 1H), 4.04 (s, 3H), 3.92 (s, 3H), 3.69 (s, 3H) ppm. $^{13}$C NMR (126 MHz, $CDCl_3$) δ 154.0, 151.2, 150.0, 149.3, 149.2, 149.1, 129.5, 125.4, 121.5, 120.4, 117.0, 114.2, 111.2, 109.0, 102.2, 92.7, 56.3, 55.8, 55.7 ppm. HRMS (ESI) m/z calcd for $C_{19}H_{17}N_3O_3$ [M + H]$^+$, 336.1343; found, 336.1337.

**4-((3-Methoxyphenyl)amino)-6,7-dimethoxyquinoline-3-carbonitrile (29)**

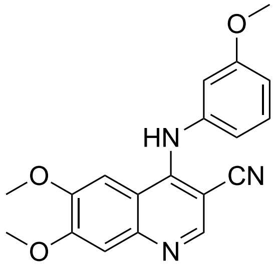

A mixture of 4-chloro-6,7-dimethoxyquinoline-3-carbonitrile (200 mg, 0.80 mmol, 1.0 equiv) and 3-methoxyaniline (109 mg, 98 µL, 0.89 mmol, 1.1 equiv) in EtOH (15 mL) was heated at 100 ºC for 18 h. After completion of the reaction, the mixture was concentrated under reduced pressure, and the residue was purified by flash column chromatography (EtOAc/hexanes, then 1–5% MeOH in EtOAc) to afford compound **29** (230 mg, 90% yield) as a white solid. $^1$H NMR (500 MHz, $CDCl_3$) δ 8.66 (s, 1H), 7.36 (s, 1H), 6.88 (s, 1H), 6.79 (s, 1H), 6.75 (dd, *J* = 2.4, 8.2 Hz, 1H), 6.68 (dd, *J* = 2.4, 7.0 Hz, 1H), 6.64 (t, *J* = 7.0 Hz, 1H), 4.03 (s, 3H), 3.77 (s, 3H), 3.61 (s, 3H) ppm. $^{13}$C NMR (126 MHz, $CDCl_3$) δ 160.6, 154.0, 149.6, 149.5, 149.4, 149.0, 147.8, 142.2, 130.3, 117.0, 114.8, 110.7, 109.0, 108.4, 102.8, 92.7, 56.3, 55.7, 55.4 ppm. HRMS (ESI) m/z calcd for $C_{19}H_{17}N_3O_3$ $[M + H]^+$, 336.1343; found, 336.1339.

**4-((4-Methoxyphenyl)amino)-6,7-dimethoxyquinoline-3-carbonitrile (30)**

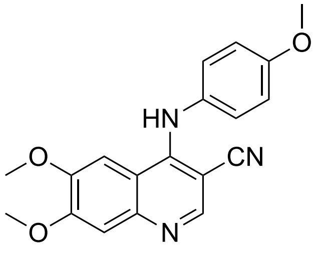

A mixture of 4-chloro-6,7-dimethoxyquinoline-3-carbonitrile (100 mg, 0.40 mmol, 1.0 equiv) and 4-methoxyaniline (55 mg, 0.45 mmol, 1.1 equiv) in EtOH (5 mL) was heated at 100 ºC for 18 h. After completion of the reaction, the mixture was concentrated under reduced pressure, and the residue was purified by flash column chromatography (EtOAc/hexanes, then 1–5% MeOH in

EtOAc) to afford compound **30** (122 mg, 90% yield) as a white solid. $^1$H NMR (500 MHz, DMSO-$d_6$) δ 10.05 (s, 1H), 8.48 (s, 1H), 7.91 (s, 1H), 7.30 (s, 1H), 7.23 (d, *J* = 8.7 Hz, 2H), 6.94 (d, *J* = 8.8 Hz, 2H), 3.88 (s, 3H), 3.88 (s, 3H), 3.73 (s, 3H) ppm. $^{13}$C NMR (126 MHz, DMSO-$d_6$) δ 156.5, 152.3, 149.8, 148.1, 147.7, 140.3, 129.8, 126.1, 114.5, 112.6, 110.8, 104.3, 100.9, 83.8, 54.8, 54.4, 53.7 ppm. HRMS (ESI) m/z calcd for $C_{19}H_{17}N_3O_3$ $[M + H]^+$, 336.1343; found, 336.1338.

**4-((2,3-Dimethoxyphenyl)amino)-6,7-dimethoxyquinoline-3-carbonitrile (31)**

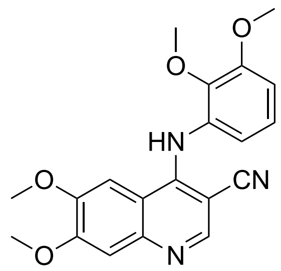

A mixture of 4-chloro-6,7-dimethoxyquinoline-3-carbonitrile (100 mg, 0.40 mmol, 1.0 equiv) and 2,3-dimethoxyaniline (68 mg, 61 μL, 0.44 mmol, 1.1 equiv) in EtOH (3 mL) was heated at 100 ºC for 18 h. After completion of the reaction, the mixture was concentrated under reduced pressure, and the residue was purified by flash column chromatography (EtOAc/hexanes, then 1–5% MeOH in EtOAc) to afford compound **31** (137 mg, 93% yield) as a white solid. $^1$H NMR (500 MHz, $CDCl_3$) δ 8.68 (s, 1H), 7.40 (s, 1H), 7.03 (s, 1H), 6.99 (t, *J* = 8.2 Hz, 1H), 6.91 (s, 1H), 6.73 (dd, *J* = 1.3, 8.3 Hz, 1H), 6.55 (dd, *J* = 1.3, 8.1 Hz, 1H), 4.05 (s, 3H), 3.93 (s, 3H), 3.92 (s, 3H), 3.76 (s, 3H) ppm. $^{13}$C NMR (126 MHz, DMSO-$d_6$) δ 153.8, 153.4, 151.4, 150.8, 149.6, 146.2, 144.9, 124.2, 119.8, 117.6, 113.5, 111.8, 109.2, 102.4, 87.5, 60.3, 56.6, 56.4, 56.3 ppm. HRMS (ESI) m/z calcd for $C_{20}H_{19}N_3O_4$ $[M + H]^+$, 366.1448; found, 366.1445.

**4-((2,4-Dimethoxyphenyl)amino)-6,7-dimethoxyquinoline-3-carbonitrile (32)**

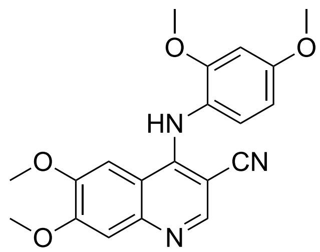

A mixture of 4-chloro-6,7-dimethoxyquinoline-3-carbonitrile (100 mg, 0.40 mmol, 1.0 equiv) and 2,4-dimethoxyaniline (68 mg, 60 µL, 0.44 mmol, 1.1 equiv) in EtOH (3 mL) was heated at 100 ºC for 18 h. After completion of the reaction, the mixture was concentrated under reduced pressure, and the residue was purified by flash column chromatography (EtOAc/hexanes, then 1–5% MeOH in EtOAc) to afford compound **32** (134 mg, 91% yield) as a white solid. $^{1}$H NMR (500 MHz, $CDCl_3$) δ 10.14 (br s, 1H), 8.61 (s, 1H), 7.97 (s, 1H), 7.32 (s, 1H), 7.23 (d, *J* = 8.6 Hz, 1H), 6.64 (d, *J* = 2.6 Hz, 1H), 6.55 (dd, *J* = 2.6, 8.6 Hz, 1H), 3.90 (s, 3H), 3.89 (s, 3H), 3.76 (s, 3H), 3.70 (s, 3H) ppm. $^{13}$C NMR (126 MHz, DMSO-$d_6$) δ 175.4, 174.6, 161.2, 157.3, 154.9, 152.7, 150.1, 149.1, 130.1, 118.8, 114.3, 112.2, 105.2, 103.2, 99.2, 85.2, 57.0, 56.6, 56.2, 56.0 ppm. HRMS (ESI) m/z calcd for $C_{20}H_{19}N_3O_4$ $[M + H]^+$, 366.1448; found, 366.1443.

**4-((2,6-Dimethoxyphenyl)amino)-6,7-dimethoxyquinoline-3-carbonitrile (33)**

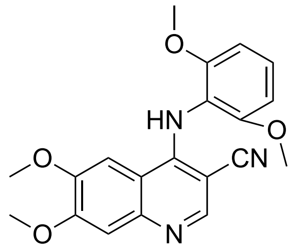

A mixture of 4-chloro-6,7-dimethoxyquinoline-3-carbonitrile (100 mg, 0.40 mmol, 1.0 equiv) and 2,6-dimethoxyaniline (68 mg, 0.44 mmol, 1.1 equiv) in EtOH (3 mL) was heated at 100 ºC for 18 h. After completion of the reaction, the mixture was concentrated under reduced pressure, and the residue was purified by flash column chromatography (EtOAc/hexanes, then 1–5% MeOH in

EtOAc) to afford compound **33** (132 mg, 90% yield) as a white solid. 1H NMR (500 MHz, DMSO-*d*$_6$) δ 8.92 (s, 1H), 8.25 (s, 1H), 7.87 (s, 1H), 7.35 (t, *J* = 8.5 Hz 1H), 7.26 (s, 1H), 6.76 (s, 1H), 6.74 (s, 1H), 3.92 (s, 3H), 3.92 (s, 3H), 3.75(s, 6H) ppm. $^{13}$C NMR (126 MHz, DMSO-*d*$_6$) δ 157.5, 153.4, 151.8, 150.8, 149.3, 145.8, 129.5, 117.8, 115.0, 112.5, 109.2, 104.6, 102.5, 84.4, 56.7, 56.2, 56.2 ppm. HRMS (ESI) m/z calcd for $C_{20}H_{19}N_3O_4$ [M + H]$^+$, 366.1448; found, 366.1442.

**4-((3,4-Dimethoxyphenyl)amino)-6,7-dimethoxyquinoline-3-carbonitrile (34)**

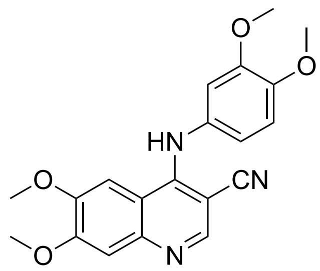

A mixture of 4-chloro-6,7-dimethoxyquinoline-3-carbonitrile (100 mg, 0.40 mmol, 1.0 equiv) and 3,4-dimethoxyaniline (68 mg, 0.44 mmol, 1.1 equiv) in EtOH (3 mL) was heated at 100 ºC for 18\ h. After completion of the reaction, the mixture was concentrated under reduced pressure, and the residue was purified by flash column chromatography (EtOAc/hexanes, then 1–5% MeOH in EtOAc) to afford compound **34** (132 mg, 90% yield) as a white solid. $^1$H NMR (500 MHz, DMSO-*d*$_6$) δ 10.88 (br s, 1H), 8.82 (s, 1H), 8.03 (s, 1H), 7.35 (s, 1H), 7.03 (d, *J* = 7.0 Hz, 1H), 7.00 – 6.98 (m, 1H), 6.94 (dd, *J* = 2.4, 8.5 Hz, 1H), 3.93 (s, 3H), 3.91 (s, 3H), 3.75 (s, 3H), 3.70 (s, 3H) ppm. $^{13}$C NMR (126 MHz, DMSO-*d*$_6$) δ 155.6, 153.4, 150.5, 149.5, 149.4, 148.1, 130.4, 119.8, 114.8, 112.6, 112.0, 111.7, 103.6, 102.3, 85.9, 57.2, 56.9, 56.2, 56.1 ppm. HRMS (ESI) m/z calcd for $C_{20}H_{19}N_3O_4$ [M + H]$^+$, 366.1448; found, 366.1445.

**(*S*)-4-(sec-butylamino)-6,7-dimethoxyquinoline-3-carbonitrile (35)**

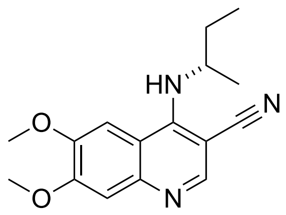

A mixture of 4-chloro-6,7-dimethoxyquinoline-3-carbonitrile (50 mg, 0.20 mmol, 1.0 equiv) and (1R)-(-)-1-methylpropylamine (16 mg, 22 μL, 0.22 mmol, 1.1 equiv) in EtOH (3 mL) was heated at 100 ºC for 18 h. After completion of the reaction, the mixture was concentrated under reduced pressure, and the residue was purified by flash column chromatography (EtOAc/hexanes, then 1–5% MeOH in EtOAc) to afford compound **35** (49.4 mg, 85% yield) as a white solid. $^{1}$H NMR (500 MHz, $CDCl_3$) δ 8.49 (s, 1H), 7.33 (s, 1H), 6.89 (s, 1H), 4.92 (d, *J* = 8.9 HZ, 1H), 4.57 – 4.51 (m, 1H), 4.03 (s, 3H), 4.02 (s, 3H), 1.85 – 1.71 (m, 2H), 1.41 (d, *J* = 6.3 HZ, 3H), 1.07 (t, *J* = 7.4 Hz, 3H) ppm. $^{13}$C NMR (126 MHz, $CDCl_3$) δ 153.5, 151.9, 150.4, 149.5, 145.9, 119.8, 112.0, 109.8, 98.8, 84.7, 56.3, 56.3, 51.6, 30.8, 21.5, 10.0 ppm. HRMS (ESI) m/z calcd for $C_{16}H_{19}N_3O_2$ $[M + H]^+$, 286.1550; found, 286.1546.

**4-(Cyclopropylamino)-6,7-dimethoxyquinoline-3-carbonitrile (36)**

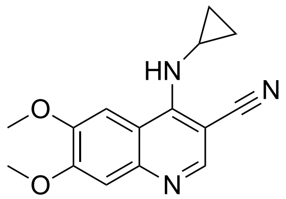

A mixture of 4-chloro-6,7-dimethoxyquinoline-3-carbonitrile (200 mg, 0.80 mmol, 1.0 equiv) and cyclopropanamine (50 mg, 61 μL, 0.89 mmol, 1.1 equiv) was heated at 100 ºC for 18 h. After completion of the reaction, the mixture was concentrated under reduced pressure, and the residue was purified by flash column chromatography (EtOAc/hexanes, then 1–5% MeOH in EtOAc) to afford compound **36** (195 mg, 90% yield) as a white solid. $^{1}$H NMR (500 MHz, $CDCl_3$) δ 8.54 (s,

1H), 7.33 (s, 1H), 7.01 (s, 1H), 5.54 (br s, 1H), 4.02 (s, 3H), 4.01 (s, 3H), 3.28 – 3.23 (m, 1H), 1.15 – 1.11 (m, 2H), 0.92 – 0.89 (m, 2H) ppm. $^{13}$C NMR (126 MHz, $CDCl_3$) δ 158.1, 157.3, 156.8, 154.0, 150.0, 125.6, 116.8, 114.0, 106.8, 88.3, 61.5, 60.9, 31.5, 16.0 ppm. HRMS (ESI) m/z calcd for $C_{15}H_{15}N_3O_2$ $[M + H]^+$, 270.1237; found, 270.1233.

**4-((4-Ethynylphenyl)amino)-6,7-dimethoxyquinoline-3-carbonitrile (37)**

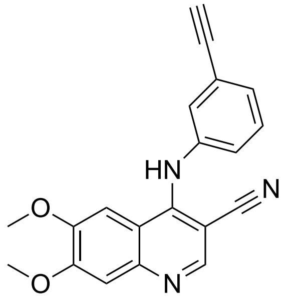

A mixture of 4-chloro-6,7-dimethoxyquinoline-3-carbonitrile (200 mg, 0.80 mmol, 1.0 equiv) and 4-ethynylaniline (104 mg, 0.89 mmol, 1.1 equiv) in EtOH (15 mL) was heated at 100 ºC for 18 h. After completion of the reaction, the mixture was concentrated under reduced pressure, and the residue was purified by flash column chromatography (EtOAc/hexanes, then 1–5% MeOH in EtOAc) to afford compound **37** (244 mg, 92% yield) as a white solid. $^1$H NMR (500 MHz, DMSO-$d_6$) δ 10.47 (br s, 1H), 8.73 (s, 1H), 7.94 (s, 1H), 7.41 – 7.33 (m, 5H), 4.20 (s, 1H), 3.92 (s, 3H), 3.90 (s, 3H) ppm. $^{13}$C NMR (126 MHz, DMSO-$d_6$) δ 153.1, 149.4, 148.2, 146.9, 138.5, 137.4, 128.0, 127.9, 125.9, 123.7, 120.8, 113.8, 111.8, 102.7, 101.4, 85.9, 81.3, 79.8, 54.9, 54.6 ppm. HRMS (ESI) m/z calcd for $C_{20}H_{15}N_3O_2$ $[M + H]^+$, 330.1237; found, 330.1232.

**4-((Benzo[c][1,2,5]thiadiazol-4-yl)amino)-6,7-dimethoxyquinoline-3-carbonitrile (38)**

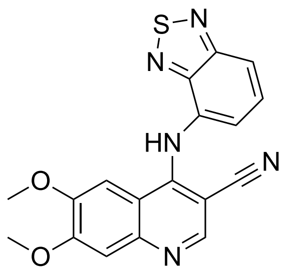

A mixture of 4-chloro-6,7-dimethoxyquinoline-3-carbonitrile (200 mg, 0.80 mmol, 1.0 equiv) and benzo[c][1,2,5]thiadiazol-4-amine (134 mg, 0.89 mmol, 1.1 equiv) in EtOH (15 mL) was heated at 100 ºC for 18 h. After completion of the reaction, the mixture was concentrated under reduced pressure, and the residue was purified by flash column chromatography (EtOAc/hexanes, then 1–5% MeOH in EtOAc) to afford compound **38** (252 mg, 87% yield) as a yellow solid. $^{1}$H NMR (500 MHz, DMSO-$d_6$) δ 10.90 (br s, 1H), 8.61 (s, 1H), 8.01 – 7.99 (m, 2H), 7.76 (dd, $J$ = 7.2, 8.8 Hz, 1H), 7.63 (d, $J$ = 7.1 Hz, 1H), 7.37 (s, 1H), 3.94 (s, 3H), 3.91 (s, 3H) ppm. $^{13}$C NMR (126 MHz, DMSO-$d_6$) δ 155.6, 155.1, 151.4, 151.0, 150.4, 149.2, 149.2, 130.7, 124.5, 124.5, 119.9, 116.5, 113.9, 103.2, 88.9, 57.0, 56.7 ppm. HRMS (ESI) m/z calcd for $C_{18}H_{13}N_5O_2S$ $[M + H]^+$, 364.0863; found, 364.0859.

**4-(Methylamino)-6,7-dimethoxyquinoline-3-carbonitrile (39)**

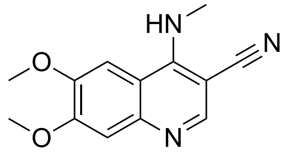

A mixture of 4-chloro-6,7-dimethoxyquinoline-3-carbonitrile (50 mg, 0.20 mmol, 1.0 equiv) and methanamine (17 mg, 19 µL, 0.22 mmol, 1.1 equiv, 40 wt % in H2O) in EtOH (3 mL) was heated at 100 ºC for 18 h. After completion of the reaction, the mixture was concentrated under reduced pressure, and the residue was purified by flash column chromatography (EtOAc/hexanes, then 1–5% MeOH in EtOAc) to afford compound **39** (47 mg, 95% yield) as a white solid. $^{1}$H NMR

(500 MHz, DMSO-$d_6$) δ 8.31 (s, 1H), 7.97 (br s, 1H), 7.57 (s, 1H), 7.16 (s, 1H), 3.84 (m, 6H), 3.28 (d, *J* = 7.0 Hz, 3H) ppm. $^{13}$C NMR (126 MHz, DMSO-$d_6$) δ 153.5, 152.4, 151.7, 149.3, 120.9, 112.1, 108.8, 102.0, 82.0, 56.7, 56.2, 31.8, 24.9 ppm. HRMS (ESI) m/z calcd for $C_{13}H_{13}N_3O_2$ $[M + H]^+$, 244.1081; found, 244.1079.

**4-((4-Hydroxyphenethyl)amino)-6,7-dimethoxyquinoline-3-carbonitrile (40)**

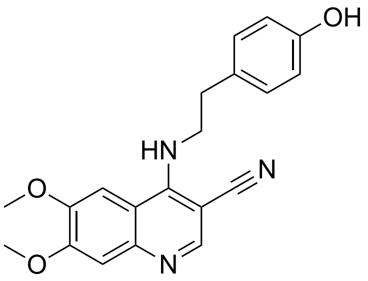

A mixture of 4-chloro-6,7-dimethoxyquinoline-3-carbonitrile (50 mg, 0.20 mmol, 1.0 equiv) and tyramine (30 mg, 0.22 mmol, 1.1 equiv) in EtOH (3 mL) was heated at 100 ºC for 18 h. After completion of the reaction, the mixture was concentrated under reduced pressure, and the residue was purified by flash column chromatography (EtOAc/hexanes, then 1–5% MeOH in EtOAc) to afford compound **40** (63 mg, 90% yield) as a white solid. $^1$H NMR (500 MHz, DMSO-$d_6$) δ 9.17 (s, 1H), 8.28 (s, 1H), 7.83 (t, *J* = 6.3 Hz, 1H), 7.52 (s, 1H), 7.16 (s, 1H), 7.06 (d, *J* = 8.4 Hz, 2H), 6.63 (d, *J* = 8.1 Hz, 2H), 3.84 (s, 3H), 3.83 (s, 3H),3.25 – 3.23 (m, 2H), 2.86 (t, *J* = 7.0 Hz, 2H) ppm. $^{13}$C NMR (126 MHz, DMSO-$d_6$) δ 156.4, 153.4, 152.0, 151.1, 149.3, 145.4, 130.2, 129.0, 121.2, 115.7, 112.3, 109.4, 109.0, 102.3, 101.9, 81.9, 56.6, 56.2, 46.0, 35.7 ppm. HRMS (ESI) m/z calcd for $C_{20}H_{19}N_3O_3$ $[M + H]^+$, 350.1499; found, 350.1495.

**6,7-Dimethoxy-N-phenylquinolin-4-amine (41)**

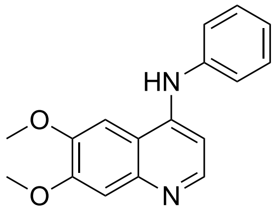

A mixture of 4-chloro-6,7-dimethoxyquinoline (100 mg, 0.45 mmol, 1.0 equiv) and aniline (46 mg, 45 µL, 0.49 mmol, 1.1 equiv) in EtOH (7.5 mL) was heated at 100 ºC for 18 h. After completion of the reaction, the mixture was concentrated under reduced pressure, and the residue was purified by flash column chromatography (EtOAc/hexanes, then 1–5% MeOH in EtOAc) to afford compound **41** (112 mg, 90% yield) as a white solid. $^1$H NMR (500 MHz, DMSO-$d_6$) δ 8.34 (d, *J* = 6.7 Hz, 1H), 8.10 (d, *J* = 3.9 Hz, 1H), 7.57 (t, *J* = 7.8 Hz, 2H), 7.49 – 7.45 (m, 2H), 7.45 – 7.35 (m, 2H), 6.73 (d, *J* = 6.7 Hz, 1H), 4.00 (s, 3H), 3.98 (s, 3H) ppm. $^{13}$C NMR (126 MHz, DMSO-$d_6$) δ 154.7, 149.8, 141.5, 138.4, 130.3, 127.1, 125.5, 112.4, 102.9, 101.5, 99.8, 57.0, 56.6, 52.5, 40.6, 40.5, 40.4, 40.3, 40.2, 40.2, 40.1, 40.0, 39.9, 39.8, 39.7, 39.5 ppm. HRMS (ESI) m/z calcd for $C_{17}H_{16}N_2O_2$ [M + H]$^+$, 281.1285; found, 281.1282.

**4-((Pyridin-3-yl)amino)-6,7-dimethoxyquinoline-3-carbonitrile (42)**

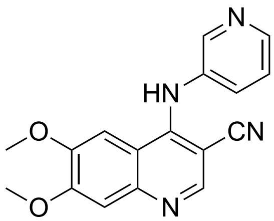

A mixture of 4-chloro-6,7-dimethoxyquinoline-3-carbonitrile (200 mg, 0.80 mmol, 1.0 equiv) and pyridin-3-amine (83 mg, 0.89 mmol, 1.1 equiv) in EtOH (15 mL) was heated at 100 ºC for 18 h. After completion of the reaction, the mixture was concentrated under reduced pressure, and the residue was purified by flash column chromatography (EtOAc/hexanes, then 1–5% MeOH in EtOAc) to afford compound **42** (55 mg, 21% yield) as a solid. $^1$H NMR (500 MHz, DMSO-$d_6$) δ

8.57 – 8.50 (m, 2H), 8.40 (d, J = 4.7 Hz, 1H), 7.76 (s, 1H), 7.68 – 7.66 (m, 1H), 7.45 (s, 1H), 7.36 (s, 1H), 7.11 (s, 1H), 3.88 (s, 3H), 3.86 (s, 3H) ppm. $^{13}$C NMR (126 MHz, DMSO-$d_6$) δ 173.7, 154.2, 148.4, 145.9, 145.3, 138.3, 135.2, 128.6, 125.9, 125.9, 119.6, 117.7, 104.6, 100.6, 93.2, 56.4, 56.1 ppm. HRMS (ESI) m/z calcd for $C_{17}H_{14}N_4O_2$ $[M + H]^+$, 307.1190; found, 307.1188.

**6-(heptyloxy)-7-methoxy-4-(phenylamino)quinoline-3-carbonitrile (43)**

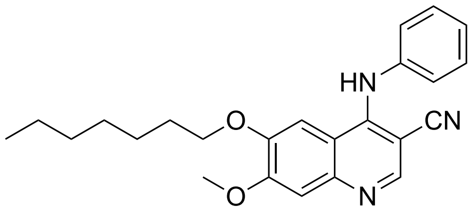

A mixture of compound **12** (40 mg, 0.14 mmol, 1 equiv) and $Cs_2CO_3$ (45 mg, 0.14 mmol, 1 equiv) in DMF (5 mL) was stirred at room temperature for 30 min, after which bromoheptane (38 mg, 0.21 mmol, 1.5 equiv) was added. The mixture was then heated at 100 °C for 2 h. After cooling to room temperature, the reaction was quenched with water (5 mL) and extracted with EtOAc (3 × 5 mL). The combined organic extracts were concentrated under reduced pressure, and the residue was purified by flash column chromatography on silica gel (30% EtOAc in Hexane) to afford compound **43** as the product (35 mg, 65% yield) as a white solid. $^1$H NMR (500 MHz, $CDCl_3$) δ 8.63 (s, 1H), 7.40 – 7.33 (m, 3H), 7.19 (d, *J* = 7.5 Hz, 1H), 7.10 (d, *J* = 7.8 Hz, 2H), 6.85 (s, 1H), 6.83 (s, 1H), 4.00 (s, 3H), 3.63 (t, *J* = 6.9 Hz, 2H), 1.68–1.65 (m, 2H), 1.36 – 1.23 (m, 8H), 0.90 (t, *J* = 6.9 Hz, 3H) ppm. $^{13}$C NMR (126 MHz, $CDCl_3$) δ 154.3, 149.5, 149.4, 148.5, 147.6, 141.1, 129.5, 125.3, 122.8, 117.1, 113.6, 109.0, 103.6, 92.2, 68.8, 56.2, 31.7, 28.9, 28.5, 25.7, 22.6, 14.1 ppm. HRMS (ESI) m/z calcd for $C_{24}H_{27}N_3O_2$ $[M + H]^+$, 390.2176; found, 390.2174.

**7-methoxy-6-(2-(2-methoxyethoxy)ethoxy)-4-(phenylamino)quinoline-3-carbonitrile (44)**

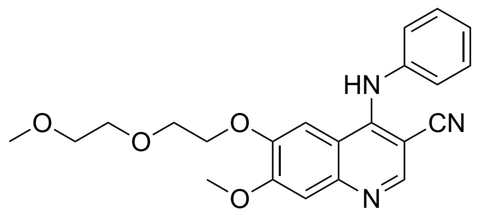

A mixture of compound **12** (40 mg, 0.14 mmol, 1 equiv) and $Cs_2CO_3$ (45 mg, 0.14 mmol, 1 equiv) in DMF (5 mL) was stirred at room temperature for 30 min, after which 1-bromo-2-(2-methoxyethoxy)ethane (38 mg, 0.21 mmol, 1.5 equiv) was added. The mixture was then heated at 100 °C for 2 h. After cooling to room temperature, the reaction was quenched with water (5 mL) and extracted with EtOAc (3 × 5 mL). The combined organic extracts were concentrated under reduced pressure, and the residue was purified by flash column chromatography on silica gel (50% EtOAc in Hexane) to afford compound **44** as the product (30 mg, 56% yield) as a white solid. $^1$H NMR (500 MHz, $CDCl_3$) δ 8.62 (s, 1H), 7.42 – 7.34 (m, 3H), 7.22 (t, $J$ = 7.5 Hz, 1H), 7.13 (d, $J$ = 7.4 Hz, 2H), 7.09 (s, 1H), 7.07 (s, 1H), 4.00 (s, 3H), 3.97 – 3.94 (m, 2H), 3.81 – 3.76 (m, 2H), 3.70 – 3.65 (m, 2H), 3.58 – 3.54 (m, 2H), 3.34 (s, 3H) ppm. $^{13}$C NMR (126 MHz, $CDCl_3$) δ 154.4, 150.1, 149.7, 148.5, 140.7, 129.5, 125.5, 122.9, 117.0, 113.7, 109.1, 104.4, 91.6, 71.8, 70.5, 69.4, 68.6, 58.9, 56.2 ppm. HRMS (ESI) m/z calcd for $C_{22}H_{23}N_3O_4$ $[M + H]^+$, 394.1761; found, 394.1759.

**7-(heptyloxy)-6-methoxy-4-(phenylamino)quinoline-3-carbonitrile (45)**

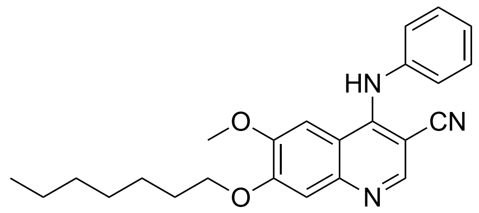

A mixture of compound **18** (40 mg, 0.14 mmol, 1 equiv) and $Cs_2CO_3$ (45 mg, 0.14 mmol, 1 equiv) in DMF (5 mL) was stirred at room temperature for 30 min, after which bromoheptane (38 mg,

0.21 mmol, 1.5 equiv) was added. The mixture was then heated at 100 °C for 2 h. After cooling to room temperature, the reaction was quenched with water (5 mL) and extracted with EtOAc (3 × 5 mL). The combined organic extracts were concentrated under reduced pressure, and the residue was purified by flash column chromatography on silica gel (30% EtOAc in Hexane) to afford compound **45** as the product (32 mg, 59% yield) as a white solid. $^1$H NMR (500 MHz, $CDCl_3$) δ 8.63 (s, 1H), 7.40 – 7.32 (m, 3H), 7.20 (t, *J* = 7.5 Hz, 1H), 7.11 (d, *J* = 7.6 Hz, 2H), 6.85 (s, 1H), 6.83 (s, 1H), 4.16 (t, *J* = 6.9 Hz, 2H), 3.55 (s, 3H), 1.95 – 1.87 (m, 2H), 1.51 – 1.44 (m, 2H), 1.40 – 1.35 (m, 2H), 1.35 – 1.28 (m, 4H), 0.91 – 0.86 (m, 3H) ppm. $^{13}$C NMR (126 MHz, $CDCl_3$) δ 153.6, 149.5, 149.2, 147.8, 141.0, 129.5, 125.4, 122.8, 117.1, 113.3, 109.6, 102.8, 92.1, 69.3, 55.6, 31.7, 29.0, 28.7, 25.9, 22.6, 14.1 ppm. HRMS (ESI) m/z calcd for $C_{24}H_{27}N_3O_2$ $[M + H]^+$, 390.2176; found, 390.2174.

**6-methoxy-7-(2-(2-methoxyethoxy)ethoxy)-4-(phenylamino)quinoline-3-carbonitrile (46)**

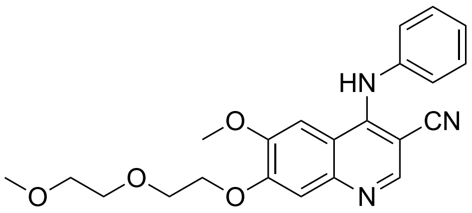

A mixture of compound **18** (40 mg, 0.14 mmol, 1 equiv) and $Cs_2CO_3$ (45 mg, 0.14 mmol, 1 equiv) in DMF (5 mL) was stirred at room temperature for 30 min, after which 1-bromo-2-(2-methoxyethoxy)ethane (38 mg, 0.21 mmol, 1.5 equiv) was added. The mixture was then heated at 100 °C for 2 h. After cooling to room temperature, the reaction was quenched with water (5 mL) and extracted with EtOAc (3 × 5 mL). The combined organic extracts were concentrated under reduced pressure, and the residue was purified by flash column chromatography on silica gel (50% EtOAc in Hexane) to afford compound **46** as the product (33 mg, 62% yield) as a white solid. $^1$H NMR (500 MHz, DMSO-$d_6$) δ 9.45 (s, 1H), 8.37 (s, 1H), 7.69 (d, *J* = 2.1 Hz, 1H), 7.35 – 7.30 (m, 2H), 7.28 (d, *J* = 4.4 Hz, 1H), 7.19 – 7.15 (m, 2H), 7.13 (t, *J* = 7.4 Hz, 1H), 4.24 – 4.19

(m, 2H), 3.84 (s, 3H), 3.79 – 3.74 (m, 2H), 3.59 – 3.53 (m, 2H), 3.45 – 3.38 (m, 2H), 3.19 (s, 3H) ppm. $^{13}$C NMR (126 MHz, DMSO-$d_6$) δ 165.8, 153.1, 151.4, 149.9, 149.7, 146.5, 129.5, 125.5, 124.1, 117.8, 114.3, 110.0, 109.9, 102.5, 88.5, 71.8, 71.2, 70.2, 69.1, 69.0, 68.6, 58.5, 56.5, 56.5, 44.0 ppm. HRMS (ESI) m/z calcd for $C_{22}H_{23}N_3O_4$ [M + H]$^+$, 394.1761; found, 394.1759.

**4-((4-heptylphenyl)amino)-6,7-dimethoxyquinoline-3-carbonitrile (47)**

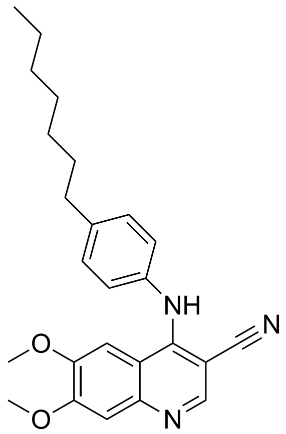

A mixture of 4-chloro-6,7-dimethoxyquinoline-3-carbonitrile (50 mg, 0.20 mmol, 1.0 equiv) and heptamine (25 mg, 0.22 mmol, 1.1 equiv) in EtOH (3 mL) was heated at 100 ºC for 18 h. After completion of the reaction, the mixture was concentrated under reduced pressure, and the residue was purified by flash column chromatography (50% EtOAc in Hexane) to afford compound **47** (62 mg, 78% yield) as a white solid. $^1$H NMR (500 MHz, DMSO-$d_6$) δ 10.68 (s, 1H), 8.76 (s, 1H), 7.99 (s, 1H), 7.36 (s, 1H), 7.24 – 7.22 (m, 4H), 3.92 (s, 3H), 3.90 (s, 3H), 2.57 (t, $J$ = 7.5 Hz, 2H), 1.55 – 1.50 (m, 2H), 1.25 – 1.14 (m, 8H), 0.79 (t, $J$ = 6.8 Hz, 3H). $^{13}$C NMR (126 MHz, DMSO-$d_6$) δ 155.4, 150.4, 148.7, 129.5, 126.6, 113.1, 103.6, 86.6, 57.1, 56.8, 35.2, 31.7, 31.4, 29.0, 28.7, 22.5, 14.4 ppm. HRMS (ESI) m/z calcd for $C_{25}H_{29}N_3O_2$ [M + H]$^+$, 404.2333; found, 404.2330.

**6-(2-(2-(2-((2-(2,6-dioxopiperidin-3-yl)-1,3-dioxoisoindolin4yl)amino)ethoxy)ethox )ethoxy)-7-methoxy-4-(phenylamino)quinoline-3-carbonitrile (48)**

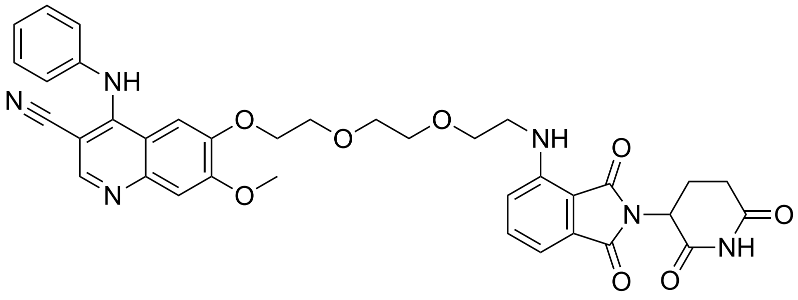

A mixture of compound **12** (100 mg, 0.34 mmol, 1 equiv) and $Cs_2CO_3$ (112 mg, 0.34 mmol, 1 equiv) in DMF (10 mL) was stirred at room temperature for 30 min, after which tert-butyl (2-(2-(2-bromoethoxy)ethoxy)ethyl)carbamate (161 mg, 0.52 mmol, 1.5 equiv) was added. The mixture was then heated at 100 °C for 2 h. After cooling to room temperature, the reaction was quenched with water (15 mL) and extracted with EtOAc (3 × 15 mL). The combined organic extracts were concentrated under reduced pressure, and the residue was redissolved in DCM (5 mL). After addition of TFA (196 mg, 0.13 mL, 1.72 mmol, 5 equiv), the reaction mixture was stirred at room temperature for 4 h. The solvent was removed under reduced pressure, and the residue was redissolved in EtOH (5 mL), followed by addition of 4-chloro-2-(2,6-dioxopiperidin-3-yl)isoindoline-1,3-dione (201 mg, 0.69 mmol, 2 equiv). The reaction mixture was then heated at 100 °C for 2 h. After cooling to room temperature, the reaction was quenched with water (15 mL) and extracted with EtOAc (3 × 15 mL). The residue was purified by flash column chromatography on silica gel (5% MeOH in DCM) to afford compound **48** as the product (56 mg, 24% yield) as a white solid. $^1$H NMR (500 MHz, DMSO-$d_6$) δ 11.08 (s, 1H), 10.20 (s, 1H), 8.74 (s, 1H), 7.90 (s, 1H), 7.59 – 7.53 (m, 1H), 7.47 (dd, $J$ = 8.4, 7.2 Hz, 2H), 7.36 (d, $J$ = 8.6 Hz, 4H), 7.12 (d, $J$ = 8.6 Hz, 1H), 7.01 (d, $J$ = 7.0 Hz, 1H), 6.60 (t, $J$ = 5.8 Hz, 1H), 5.03 (dd, $J$ = 12.9, 5.4 Hz, 1H), 4.26 (t, $J$ = 5.8 Hz, 2H), 3.98 (s, 3H), 3.86 (t, $J$ = 5.8 Hz, 2H), 3.68 – 3.60 (m, 8H), 2.85 – 2.82 (m, 2H), 2.55 – 1.96 (m, 2H). $^{13}$C NMR (126 MHz, DMSO-$d_6$) δ 178.3, 173.2, 171.1, 170.8, 170.5, 169.4, 167.7, 166.4, 161.3, 149.4, 146.9, 136.7, 134.2, 132.5, 129.7, 128.1, 125.6, 117.9, 113.4, 111.1, 109.7,

103.7, 87.4, 70.5, 70.3, 69.4, 69.0, 68.9, 56.7, 49.0, 42.2, 31.4, 22.6 ppm. HRMS (ESI) m/z calcd for $C_{36}H_{34}N_6O_8$ $[M + H]^+$, 679.2511; found, 679.2509.

**6-(4-((2-(2,6-dioxopiperidin-3-yl)-1,3-dioxoisoindolin-4-yl)amino)butoxy)-7-methoxy-4-(phenylamino)quinoline-3-carbonitrile (49)**

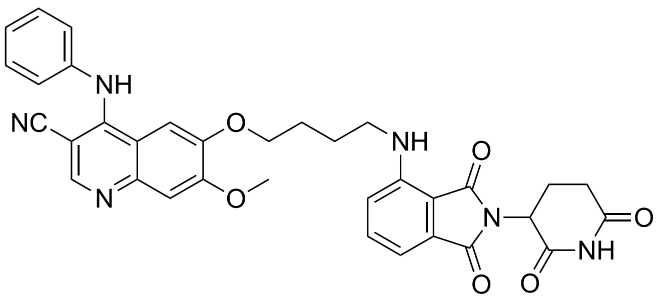

A mixture of compound **12** (100 mg, 0.34 mmol, 1 equiv) and $Cs_2CO_3$ (112 mg, 0.34 mmol, 1 equiv) in DMF (10 mL) was stirred at room temperature for 30 min, after which tert-butyl (4-bromobutyl)carbamate (130 mg, 0.52 mmol, 1.5 equiv) was added. The mixture was then heated at 100 °C for 2 h. After cooling to room temperature, the reaction was quenched with water (15 mL) and extracted with EtOAc (3 × 15 mL). The combined organic extracts were concentrated under reduced pressure, and the residue was redissolved in DCM (5 mL). After addition of TFA (196 mg, 0.13 mL, 1.72 mmol, 5 equiv), the reaction mixture was stirred at room temperature for 4 h. The solvent was removed under reduced pressure, and the residue was redissolved in EtOH (5 mL), followed by addition of 4-chloro-2-(2,6-dioxopiperidin-3-yl)isoindoline-1,3-dione (201 mg, 0.69 mmol, 2 equiv). The reaction mixture was then heated at 100 °C for 2 h. After cooling to room temperature, the reaction was quenched with water (15 mL) and extracted with EtOAc (3 × 15 mL). The residue was purified by flash column chromatography on silica gel (5% MeOH in DCM) to afford compound **49** as the product (50 mg, 24% yield) as a white solid. $^1H$ NMR (500 MHz, DMSO-$d_6$) δ 11.02 (s, 1H), 8.42 (s, 1H), 7.70 (s, 1H), 7.50 (dd, $J$ = 8.6, 7.1 Hz, 1H), 7.34 (t, $J$ = 7.7 Hz, 2H), 7.27 (s, 1H), 7.22 – 7.14 (m, 3H), 7.07 (d, $J$ = 8.6 Hz, 1H), 6.95 (d, $J$ = 7.0 Hz, 1H), 6.56 (t, $J$ = 6.0 Hz, 1H), 4.97 (dd, $J$ = 12.8, 5.4 Hz, 1H), 4.08 (t, $J$ = 6.3 Hz, 2H), 3.34 (d, $J$ =

6.6 Hz, 2H), 2.57 – 2.56 (m, 1H), 2.51 – 2.46 (m, 2H), 2.30 – 2.29 (m, 1H), 1.84 – 1.79 (m, 2H), 1.73 – 1.68 (m, 2H). $^{13}$C NMR (126 MHz, DMSO-$d_6$) δ 173.3, 170.6, 169.4, 167.8, 161.3, 157.6, 155.1, 154.4, 149.1, 148.1, 146.8, 136.7, 132.7, 129.6, 124.6, 117.7, 113.9, 110.9, 109.6, 103.3, 79.4, 69.0, 56.5, 49.0, 41.9, 34.8, 34.3, 31.4, 26.2, 25.9, 22.6 ppm. HRMS (ESI) m/z calcd for $C_{34}H_{30}N_6O_6$ $[M + H]^+$, 619.2300; found, 619.2297.

**6-(4-((2-(2,6-dioxopiperidin-3-yl)-1,3-dioxoisoindolin-4-yl)oxy)butoxy)-7-methoxy 4(phenyl-amino)quinoline-3-carbonitrile (50)**

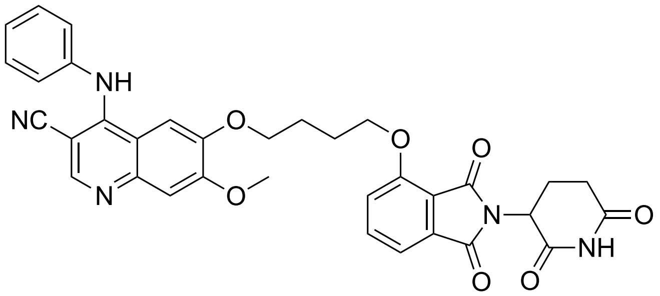

A mixture of compound **12** (100 mg, 0.34 mmol, 1 equiv) and $Cs_2CO_3$ (112 mg, 0.34 mmol, 1 equiv) in DMF (10 mL) was stirred at room temperature for 30 min, after which 1,4-dibromobutane (371 mg, 1.72 mmol, 5 equiv) was added. The mixture was then heated at 100 °C for 2 h. After cooling to room temperature, the reaction was quenched with water (15 mL) and extracted with EtOAc (3 × 15 mL). The combined organic extracts were concentrated under reduced pressure, and the residue was purified by flash column chromatography on silica gel (50% EtOAc in Hexane) to afford linker-modified quinoline. Then 2-(2,6-dioxopiperidin-3-yl)-4-hydroxyisoindoline -1,3-dione (94 mg, 0.34 mmol, 1 equiv) and $Cs_2CO_3$ (112 mg, 0.34 mmol, 1 equiv) in DMF (10 mL) was stirred at room temperature for 30 min, after which purified linker-modified quinoline was added. The mixture was then heated at 100 °C for 2 h. After cooling to room temperature, the reaction was quenched with water (15 mL) and extracted with EtOAc (3 × 15 mL). The combined organic extracts were concentrated under reduced pressure, and the residue was purified by flash column chromatography on silica gel (5% MeOH in DCM) to afford compound **50** as the product (33 mg, 16% yield) as a white solid. $^1$H NMR (500 MHz, DMSO-$d_6$) δ 9.38 (s, 1H), 8.37 (s, 1H), 7.67 (s, 1H), 7.57 (dd, $J$ = 8.4, 7.1 Hz, 1H), 7.35 – 7.30 (m, 3H), 7.26 (s, 1H), 7.22 (d, $J$ = 7.1 Hz, 1H), 7.19 – 7.10 (m, 5H), 5.10 (dd, $J$ = 13.0, 5.5 Hz, 1H), 4.05 – 4.00 (m, 2H), 3.88 (s, 3H), 3.74 – 3.63 (m, 2H), 2.95 – 2.91 (m, 2H), 2.76 – 2.56 (m, 2H), 1.74 – 1.68 (m, 2H), 1.59 – 1.53 (m, 2H) ppm. $^{13}$C NMR (126 MHz, DMSO-$d_6$) δ 172.2, 170.2, 167.6, 166.5, 166.5, 162.6, 157.0, 154.1, 151.3, 149.9, 149.0, 146.5, 140.6, 136.6, 133.7, 129.5, 125.6, 124.2, 117.7, 114.1, 112.4, 109.3, 103.1, 88.4, 68.9, 56.4, 49.6, 40.6, 40.5, 40.4, 40.3, 40.2, 40.1, 40.0, 39.8, 39.6, 39.5, 31.6, 29.5, 26.5, 24.7, 21.9 ppm. HRMS (ESI) m/z calcd for $C_{34}H_{29}N_5O_7$ $[M + H]^+$, 620.2140; found, 620.2139.

**N-(5-((3-cyano-7-methoxy-4-(phenylamino)quinolin-6-yl)oxy)pentyl)-2-(2,6-dioxopiperidin-3-yl)-1,3-dioxoisoindoline-5-carboxamide (51)**

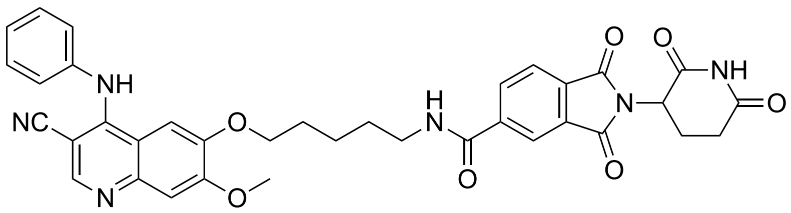

A mixture of compound **12** (100 mg, 0.34 mmol, 1 equiv) and $Cs_2CO_3$ (112 mg, 0.34 mmol, 1 equiv) in DMF (10 mL) was stirred at room temperature for 30 min, after which tert-butyl (5-bromopentyl)carbamate (137 mg, 0.52 mmol, 1.5 equiv) was added. The mixture was then heated at 100 °C for 2 h. After cooling to room temperature, the reaction was quenched with water (15 mL) and extracted with EtOAc (3 × 15 mL). The combined organic extracts were concentrated under reduced pressure, and the residue was redissolved in DCM (5 mL). After addition of TFA (196 mg, 0.13 mL, 1.72 mmol, 5 equiv), the reaction mixture was stirred at room temperature for 4 h. The solvent was removed under reduced pressure, and the residue was redissolved in DMF (5 mL), followed by addition of 2-(3H-[1,2,3]triazolo[4,5-b]pyridin-3-yl)-1,1,3,3-tetramethy-lisouronium hexafluorophosphate(V) (HATU) (196 mg, 0.52 mmol, 1.5 equiv), N-ethyl-N-isopropylpropan-2-amine (DIPEA) (67 mg, 90 μL, 0.52 mmol, 1.5 equiv) and 2-(2,6-dioxopiperidin-3-yl)-1,3-dioxoisoindoline-4-carboxylic acid (104 mg, 0.34 mmol, 1 equiv). The reaction mixture was stirred at room temperature for 8 h, after which the reaction was quenched with water (15 mL) and extracted with EtOAc (3 × 15 mL). The residue was purified by flash column chromatography on silica gel (5% MeOH in DCM) to afford compound **51** as the product (39 mg, 17% yield) as a white solid. $^{1}H$ NMR (500 MHz, DMSO-$d_6$) δ 11.16 (s, 1H), 8.90 (t, *J* = 5.6 Hz, 1H), 8.68 (s, 1H), 8.36 – 8.28 (m, 2H), 8.02 (d, *J* = 7.7 Hz, 1H), 7.85 (s, 1H), 7.46 (t, *J* = 7.7 Hz, 2H), 7.34 (d, *J* = 7.3 Hz, 4H), 5.20 (dd, *J* = 12.9, 5.4 Hz, 1H), 4.13 (t, *J* = 6.5 Hz, 2H), 3.96 (s, 3H), 2.94 – 2.87 (m, 1H), 2.12 – 2.07 (m, 1H), 1.89 – 1.84 (m, 2H), 1.69 – 1.63 (m, 2H), 1.55 – 1.49 (m, 2H) ppm. $^{13}C$ NMR (126 MHz, DMSO-$d_6$) δ 173.3, 173.1, 170.4, 167.3, 165.7, 158.4, 156.2, 154.9, 152.4, 150.6, 148.7, 138.4, 137.5, 133.7, 129.8, 128.1, 126.4, 120.6, 116.9, 116.0, 113.1, 103.3, 86.9, 69.4, 67.9, 57.0, 49.2, 35.6, 31.4, 28.8, 28.5, 25.7, 25.5, 22.5 ppm. HRMS (ESI) m/z calcd for $C_{36}H_{32}N_6O_7$ $[M + H]^+$, 661.2405; found, 661.2407.

**3-(2-(2-((3-cyano-6-methoxy-4-(phenylamino)quinolin-7-yl)oxy)ethox)ethoxy)-N-(2-((2-(2,6-dioxopiperidin-3-yl)-1,3-dioxoisoindolin-4-yl)oxy)ethyl)propanamide (52)**

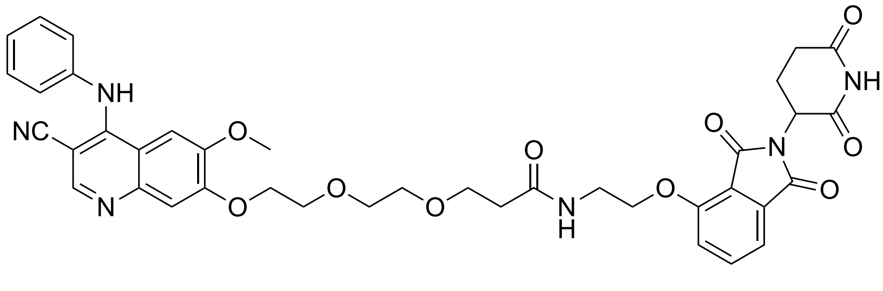

A mixture of compound **18** (100 mg, 0.34 mmol, 1 equiv) and $Cs_2CO_3$ (112 mg, 0.34 mmol, 1 equiv) in DMF (10 mL) was stirred at room temperature for 30 min, after which tert-butyl 3-(2-(2-bromoethoxy)ethoxy)propanoate (153 mg, 0.52 mmol, 1.5 equiv) was added. The mixture was then heated at 100 °C for 2 h. After cooling to room temperature, the reaction was quenched with water (15 mL) and extracted with EtOAc (3 × 15 mL). The combined organic extracts were concentrated under reduced pressure, and the residue was redissolved in DCM (5 mL). After addition of TFA (196 mg, 0.13 mL, 1.72 mmol, 5 equiv), the reaction mixture was stirred at room temperature for 4 h. The solvent was removed under reduced pressure, and the residue was redissolved in DMF (5 mL), followed by addition of 2-(3H-[1,2,3]triazolo[4,5-b]pyridin-3-yl)-1,1,3,3-tetramethy-lisouronium hexafluorophosphate(V) (HATU) (196 mg, 0.52 mmol, 1.5 equiv), N-ethyl-N-isopropylpropan-2-amine (DIPEA) (67 mg, 90 µL, 0.52 mmol, 1.5 equiv) and 4-(2-aminoethoxy)-2-(2,6-dioxopiperidin-3-yl)isoindoline-1,3-dione (163 mg, 0.52 mmol, 1.5 equiv). The reaction mixture was stirred at room temperature for 8 h, after which the reaction was quenched with water (15 mL) and extracted with EtOAc (3 × 15 mL). The residue was purified by flash column chromatography on silica gel (5% MeOH in DCM) to afford compound **52** as the product (52 mg, 20% yield) as a white solid. $^{1}$H NMR (500 MHz, $CDCl_3$) δ 9.82 (s, 1H), 8.66 (s, 1H), 7.63 (dd, *J* = 8.5, 7.3 Hz, 1H), 7.53 (d, *J* = 5.2 Hz, 1H), 7.44 (d, *J* = 7.2 Hz, 1H), 7.40 – 7.32 (m, 3H), 7.24 – 7.17 (m, 2H), 7.14 – 7.10 (m, 2H), 7.09 – 7.04 (m, 1H), 6.86 (s, 1H), 4.96 (dd, *J* = 12.5, 5.4 Hz, 1H), 4.34 (t, *J* = 4.8 Hz, 2H), 4.20 (t, *J* = 5.2 Hz, 2H), 3.91 (dd, *J* = 6.0, 3.9 Hz, 2H), 3.77 – 3.69 (m, 3H), 3.67-3.64 (m, 3H), 3.61 (t, *J* = 4.3 Hz, 2H), 3.56 (s, 3H), 2.94 – 2.67 (m, 3H), 2.50 (t, *J* = 5.8 Hz, 2H), 2.20 – 1.93 (m, 2H) ppm. $^{13}$C NMR (126 MHz, $CDCl_3$) δ 172.2, 171.1, 169.2, 166.9, 165.9, 156.1, 136.7, 133.5, 131.1, 129.6, 119.8, 117.2, 116.4, 110.7, 102.5, 89.4, 70.5, 70.2, 69.1, 68.4, 67.1, 62.8, 55.8, 53.5, 49.3, 41.9, 38.5, 36.9, 31.4, 22.6, 18.6, 17.5, 11.8 ppm. HRMS (ESI) m/z calcd for $C_{39}H_{38}N_6O_{10}$ $[M + H]^+$, 751.2722; found, 751.2725.

**7-(4-((2-(2,6-dioxopiperidin-3-yl)-1,3-dioxoisoindolin-4-yl)oxy)butoxy)-6-methoxy-4(phenyl -amino)quinoline-3-carbonitrile (53)**

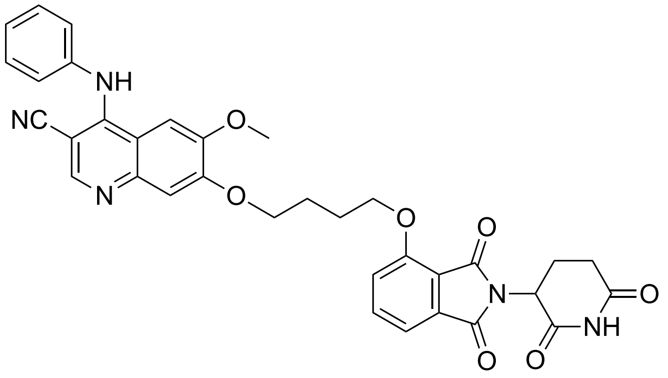

A mixture of compound **18** (100 mg, 0.34 mmol, 1 equiv) and $Cs_2CO_3$ (112 mg, 0.34 mmol, 1 equiv) in DMF (10 mL) was stirred at room temperature for 30 min, after which 1,4-dibromobutane (371 mg, 1.72 mmol, 5 equiv) was added. The mixture was then heated at 100 °C for 2 h. After cooling to room temperature, the reaction was quenched with water (15 mL) and extracted with EtOAc (3 × 15 mL). The combined organic extracts were concentrated under reduced pressure, and the residue was purified by flash column chromatography on silica gel (50% EtOAc in Hexane) to afford linker-modified quinoline. Then 2-(2,6-dioxopiperidin-3-yl)-4-hydroxyisoindoline -1,3-dione (94 mg, 0.34 mmol, 1 equiv) and $Cs_2CO_3$ (112 mg, 0.34 mmol, 1 equiv) in DMF (10 mL) was stirred at room temperature for 30 min, after which purified linker-modified quinoline was added. The mixture was then heated at 100 °C for 2 h. After cooling to room temperature, the reaction was quenched with water (15 mL) and extracted with EtOAc (3 × 15 mL). The combined organic extracts were concentrated under reduced pressure, and the residue was purified by flash column chromatography on silica gel (5% MeOH in DCM) to afford compound **53** as the product (61 mg, 30% yield) as a white solid. $^1$H NMR (500 MHz, DMSO-$d_6$) δ 11.10 (s, 1H), 8.58 (br s, 1H), 7.89 – 7.77 (m, 2H), 7.54 (d, $J$ = 8.6 Hz, 1H), 7.47 – 7.41 (m, 4H), 7.34 (s, 1H), 7.32 – 7.27 (m, 3H), 5.09 – 5.04 (m, 1H), 4.32 (t, $J$ = 6.0 Hz, 2H), 4.23 (t, $J$ = 6.0 Hz, 2H), 3.95 (s, 3H), 2.99 – 2.79 (m, 2H), 2.60 – 2.53 (m, 2H), 2.03 – 2.01 (m, 2H), 2.01 – 1.97 (m, 2H) ppm. $^{13}$C NMR (126 MHz, DMSO-$d_6$) δ 172.2, 170.2, 167.5, 166.4, 164.9, 160.2, 154.1, 151.3,

149.9, 149.0, 146.5, 140.6, 136.7, 133.7, 129.5, 125.6, 124.2, 117.7, 114.7, 114.1, 109.3, 103.1, 88.4, 69.7, 68.9, 56.4, 49.6, 31.6, 26.5, 24.7, 21.8 ppm. HRMS (ESI) m/z calcd for $C_{34}H_{29}N_5O_7$ $[M + H]^+$, 620.2140; found, 620.2140.

**7-((3-cyano-6-methoxy-4-(phenylamino)quinolin-7-yl)oxy)-N-(2-((2-(2,6-dioxopiperidin-3-yl)-1,3-dioxoisoindolin-4-yl)amino)ethyl)heptanamide (54)**

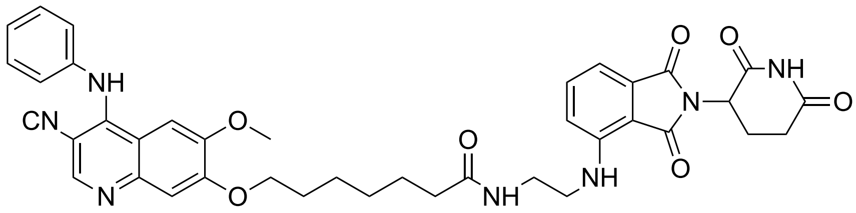

A mixture of compound **18** (100 mg, 0.34 mmol, 1 equiv) and $Cs_2CO_3$ (112 mg, 0.34 mmol, 1 equiv) in DMF (10 mL) was stirred at room temperature for 30 min, after which tert-butyl (5-bromopentyl)carbamate (137 mg, 0.52 mmol, 1.5 equiv) was added. The mixture was then heated at 100 °C for 2 h. After cooling to room temperature, the reaction was quenched with water (15 mL) and extracted with EtOAc (3 × 15 mL). The combined organic extracts were concentrated under reduced pressure, and the residue was redissolved in DCM (5 mL). After addition of TFA (196 mg, 0.13 mL, 1.72 mmol, 5 equiv), the reaction mixture was stirred at room temperature for 4 h. The solvent was removed under reduced pressure, and the residue was redissolved in DMF (5 mL), followed by addition of 2-(3H-[1,2,3]triazolo[4,5-b]pyridin-3-yl)-1,1,3,3-tetramethy-lisouronium hexafluorophosphate(V) (HATU) (196 mg, 0.52 mmol, 1.5 equiv), N-ethyl-N-isopropylpropan-2-amine (DIPEA) (67 mg, 90 μL, 0.52 mmol, 1.5 equiv) and 4-((2-aminoethyl)amino)-2-(2,6-dioxopiperidin-3-yl)isoindoline-1,3-dione (109 mg, 0.34 mmol, 1 equiv). The reaction mixture was stirred at room temperature for 8 h, after which the reaction was quenched with water (15 mL) and extracted with EtOAc (3 × 15 mL). The residue was purified by flash column chromatography on silica gel (5% MeOH in DCM) to afford compound **54** as the product (42 mg, 17% yield) as a white solid. $^1$H NMR (500 MHz, DMSO-$d_6$) δ 11.11 (s, 1H), 8.69

(s, 1H), 8.03 (t, $J$ = 5.7 Hz, 1H), 7.86 (s, 1H), 7.57 (dd, $J$ = 8.6, 7.1 Hz, 1H), 7.47 (t, $J$ = 7.8 Hz, 2H), 7.42 – 7.29 (m, 4H), 7.17 (d, $J$ = 8.7 Hz, 1H), 7.02 (d, $J$ = 7.0 Hz, 1H), 6.72 (t, $J$ = 6.2 Hz, 1H), 5.06 (dd, $J$ = 12.8, 5.4 Hz, 1H), 4.15 (t, $J$ = 6.5 Hz, 2H), 3.94 (s, 3H), 3.25 (q, $J$ = 6.2 Hz, 4H), 2.92–2.85 (m, 1H), 2.67 – 2.54 (m, 2H), 2.08 (t, $J$ = 7.4 Hz, 2H), 2.05 – 2.00 (m, 1H), 1.82–1.75 (m, 2H), 1.56–1.50 (m, 2H), 1.47–1.41 (m, 2H), 1.35 –1.29 (m, 2H). $^{13}$C NMR (126 MHz, DMSO-$d_6$) δ 173.3, 173.2, 170.6, 169.2, 167.8, 158.5, 158.3, 154.3, 151.4, 150.3, 149.6, 146.8, 139.2, 136.7, 132.6, 129.7, 127.1, 125.5, 117.6, 113.4, 111.0, 109.7, 102.9, 87.5, 69.2, 56.8, 49.0, 41.9, 38.5, 35.8, 31.4, 28.8, 28.6, 25.7, 25.6, 22.6 ppm. HRMS (ESI) m/z calcd for $C_{39}H_{39}N_7O_7$ $[M + H]^+$, 718.2984; found, 718.2981.

**7-((3-cyano-6-methoxy-4-(phenylamino)quinolin-7-yl)oxy)-N-(2-((2-(2,6-dioxopiperidin-3-yl)-1,3-dioxoisoindolin-4-yl)oxy)ethyl)heptanamide (55)**

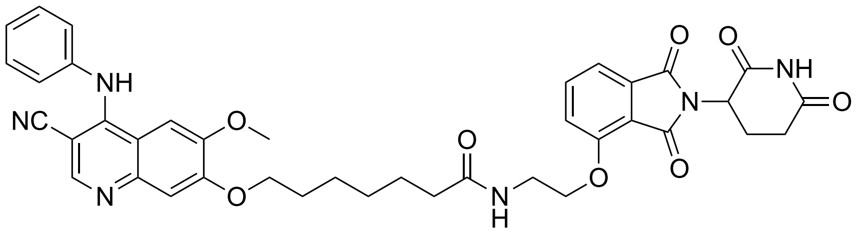

A mixture of compound **18** (100 mg, 0.34 mmol, 1 equiv) and $Cs_2CO_3$ (112 mg, 0.34 mmol, 1 equiv) in DMF (10 mL) was stirred at room temperature for 30 min, after which tert-butyl (5-bromopentyl)carbamate (137 mg, 0.52 mmol, 1.5 equiv) was added. The mixture was then heated at 100 °C for 2 h. After cooling to room temperature, the reaction was quenched with water (15 mL) and extracted with EtOAc (3 × 15 mL). The combined organic extracts were concentrated under reduced pressure, and the residue was redissolved in DCM (5 mL). After addition of TFA (196 mg, 0.13 mL, 1.72 mmol, 5 equiv), the reaction mixture was stirred at room temperature for 4 h. The solvent was removed under reduced pressure, and the residue was redissolved in DMF (5 mL), followed by addition of 2-(3H-[1,2,3]triazolo[4,5-b]pyridin-3-yl)-1,1,3,3-tetramethy-lisouronium hexafluorophosphate(V) (HATU) (196 mg, 0.52 mmol, 1.5 equiv), N-ethyl-N-isopropylpropan-2-amine (DIPEA) (67 mg, 90 µL, 0.52 mmol, 1.5 equiv) and 4-(2-aminoethoxy)-2-(2,6-dioxopiperidin-3-yl)isoindoline-1,3-dione (109 mg, 0.34 mmol, 1 equiv). The reaction mixture was stirred at room temperature for 8 h, after which the reaction was quenched with water (15 mL) and extracted with EtOAc (3 × 15 mL). The residue was purified by flash column chromatography on silica gel (5% MeOH in DCM) to afford compound **55** as the product (60 mg, 24% yield) as a white solid. $^1$H NMR (500 MHz, DMSO-$d_6$) δ 11.11 (s, 1H), 8.81 (s, 1H), 8.04 (t, *J* = 5.6 Hz, 1H), 7.92 (s, 1H), 7.80 (dd, *J* = 8.5, 7.3 Hz, 1H), 7.55 (d, *J* = 8.5 Hz, 1H), 7.53 – 7.44 (m, 3H), 7.39 (t, *J* = 5.8 Hz, 3H), 7.33 (s, 1H), 5.09 (dd, *J* = 12.7, 5.5 Hz, 1H), 4.25 (t, *J* = 5.9 Hz, 2H), 4.16 (t, *J* = 6.5 Hz, 2H), 3.95 (s, 3H), 2.98 – 2.79 (m, 2H), 2.66 – 2.53 (m, 2H), 2.12 (t, *J* = 7.3 Hz, 2H), 2.05 – 2.00 (m, 1H), 1.82 – 1.77 (m, 2H), 1.57 – 1.51 (m, 2H), 1.47 – 1.41 (m, 2H), 1.36 – 1.29 (m, 2H) ppm. $^{13}$C NMR (126 MHz, DMSO-$d_6$) δ 173.3, 173.1, 170.4, 167.3, 165.7, 158.6, 156.2, 154.8, 150.6, 148.8, 138.4, 137.5, 133.7, 129.8, 128.0, 126.3, 120.6, 116.9, 116.0, 113.1, 103.3, 86.9, 79.7, 79.4, 69.4, 67.9, 57.0, 49.2, 40.6, 40.5, 40.4, 40.3, 40.2, 40.1, 40.0, 39.8, 39.6, 39.5, 38.4, 35.6, 31.4, 28.8, 28.5, 25.7, 25.5, 22.5 ppm. HRMS (ESI) m/z calcd for $C_{39}H_{38}N_6O_8$ $[M + H]^+$, 719.2824; found, 719.2820.

#### Compound 12

LCMS

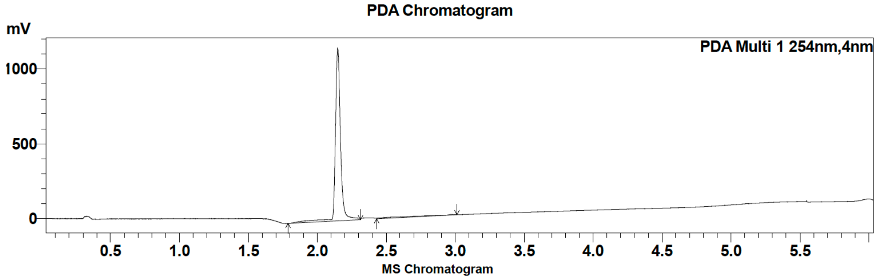

| Peak # | Ret. time | Area | Area % |
|---|---|---|---|
| 1 | 2.1 | 3072227 | 95.97 |
| 2 | 3.0 | 129038 | 4.03 |

HRMS

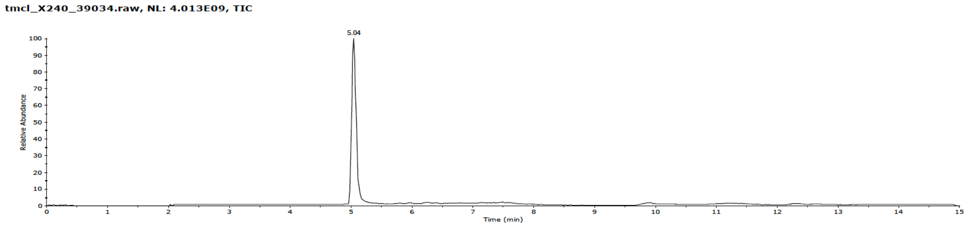

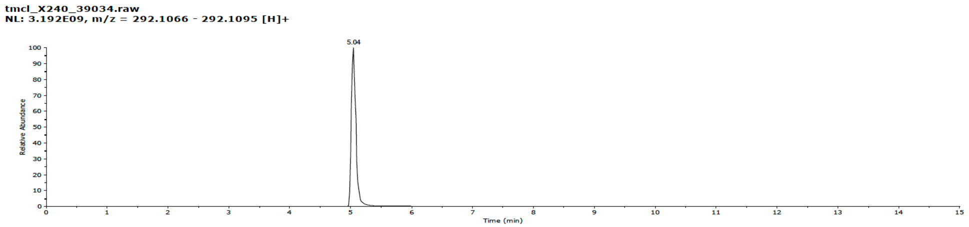

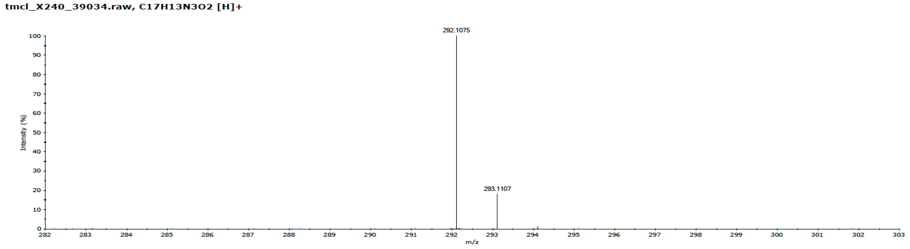

**Compound 12**

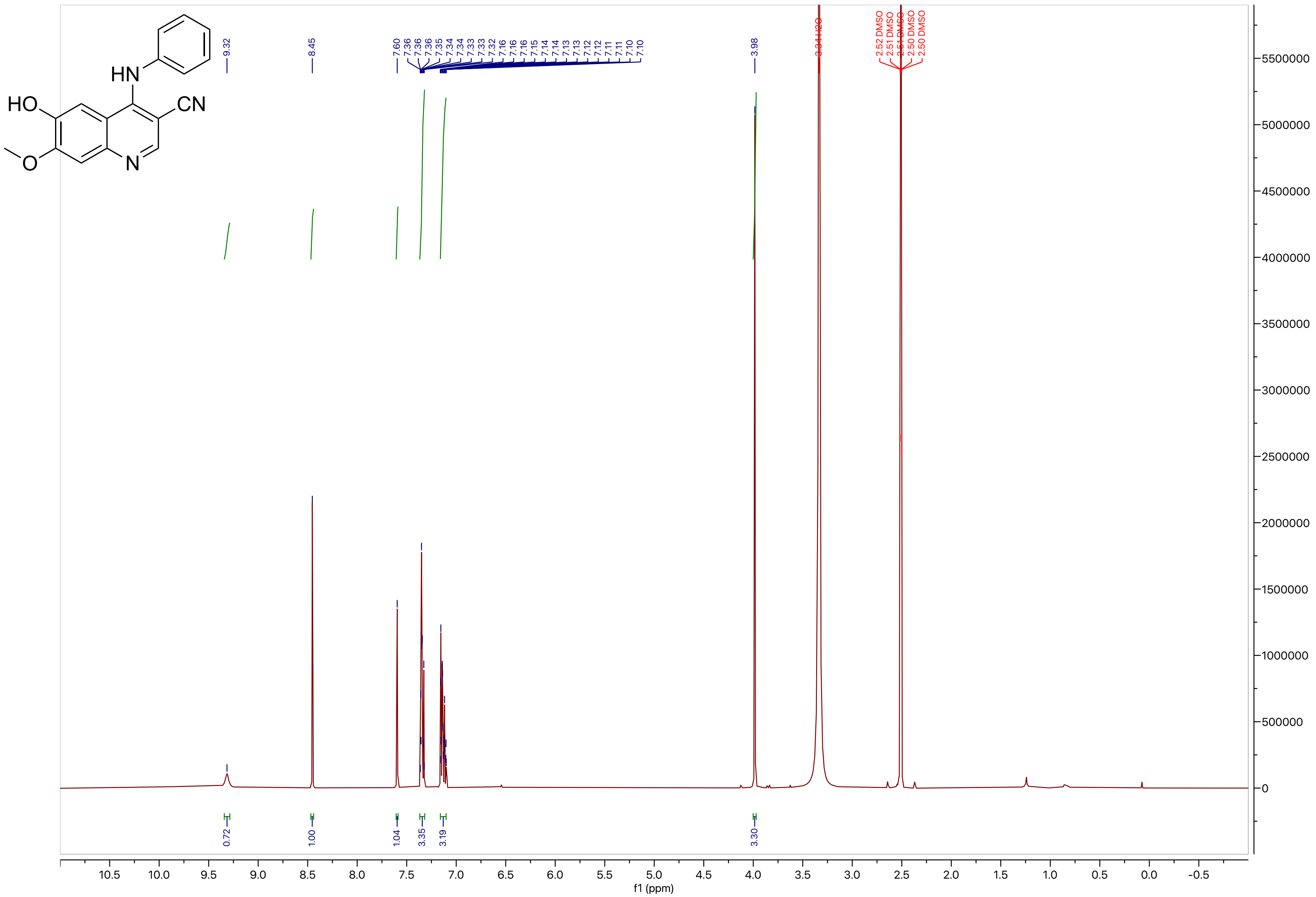

**Compound 12**

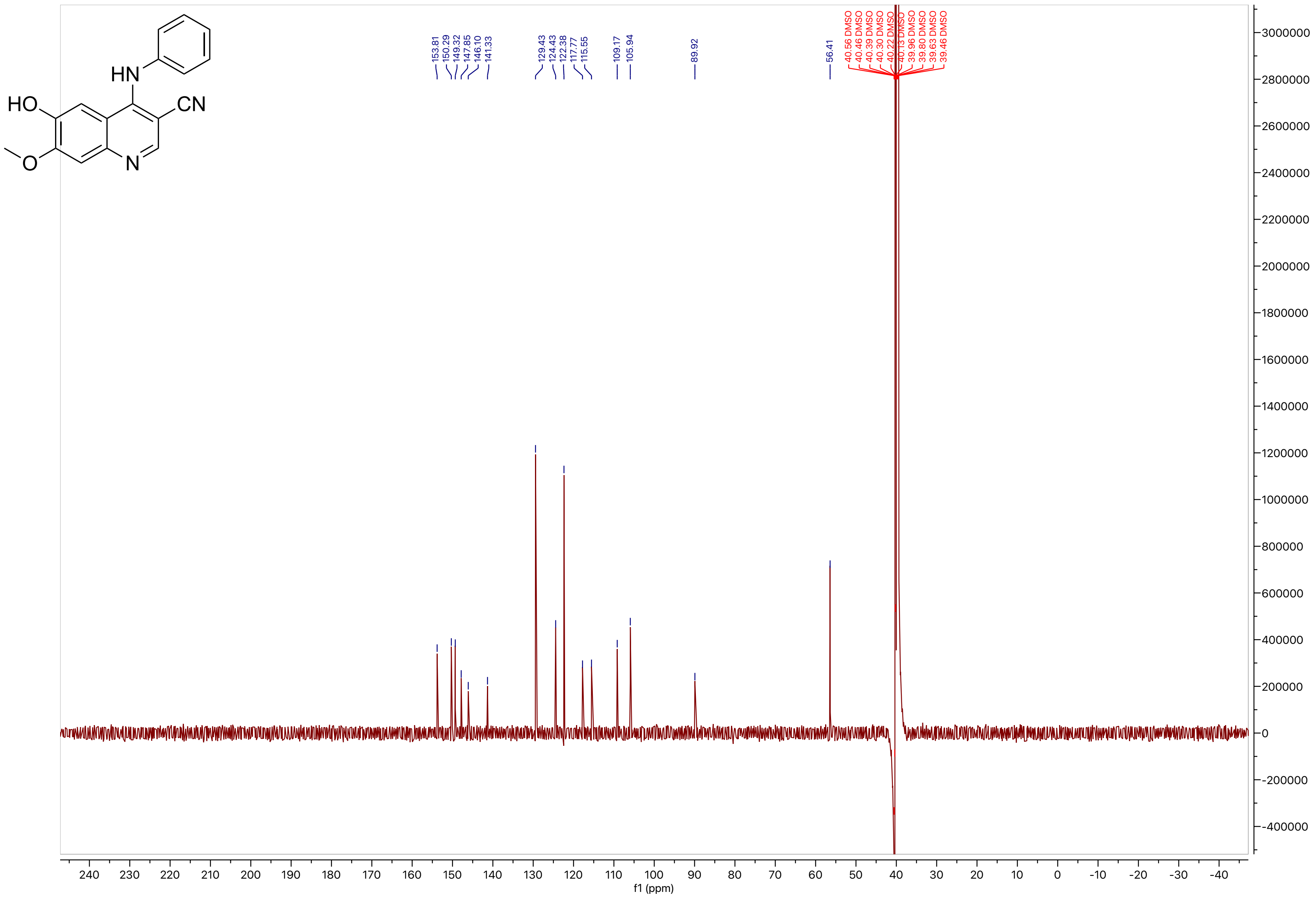
HN
HO
CN
O
N
153.81
150.29
149.32
147.85
146.10
141.33
129.43
124.43
122.38
117.77
115.55
109.17
105.94
89.92
56.41
40.56 DMSO
40.46 DMSO
40.39 DMSO
40.30 DMSO
40.22 DMSO
40.13 DMSO
39.96 DMSO
39.80 DMSO
39.63 DMSO
39.46 DMSO
f1 (ppm)

**Compound 13**

LCMS

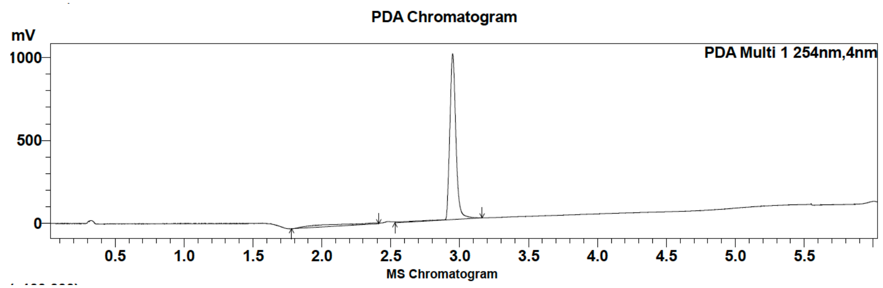

| Peak # | Ret. time | Area | Area % |
|---|---|---|---|
| 1 | 2.9 | 3124839 | 100.00 |

HRMS

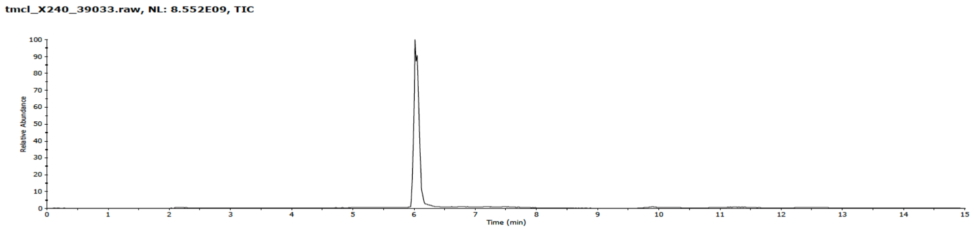

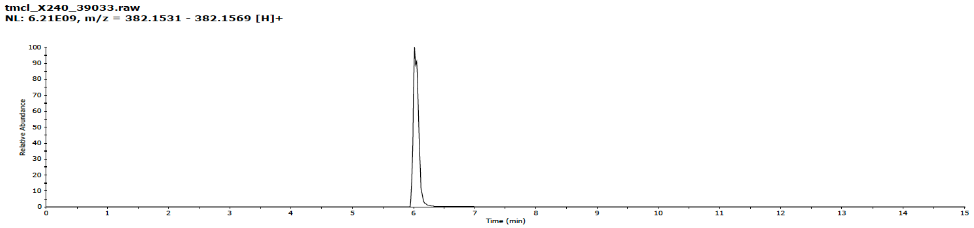

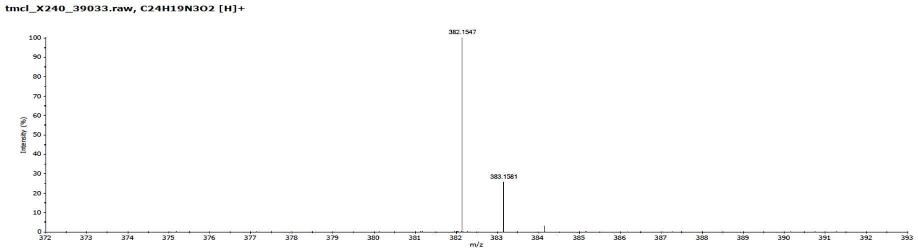

### Compound 13

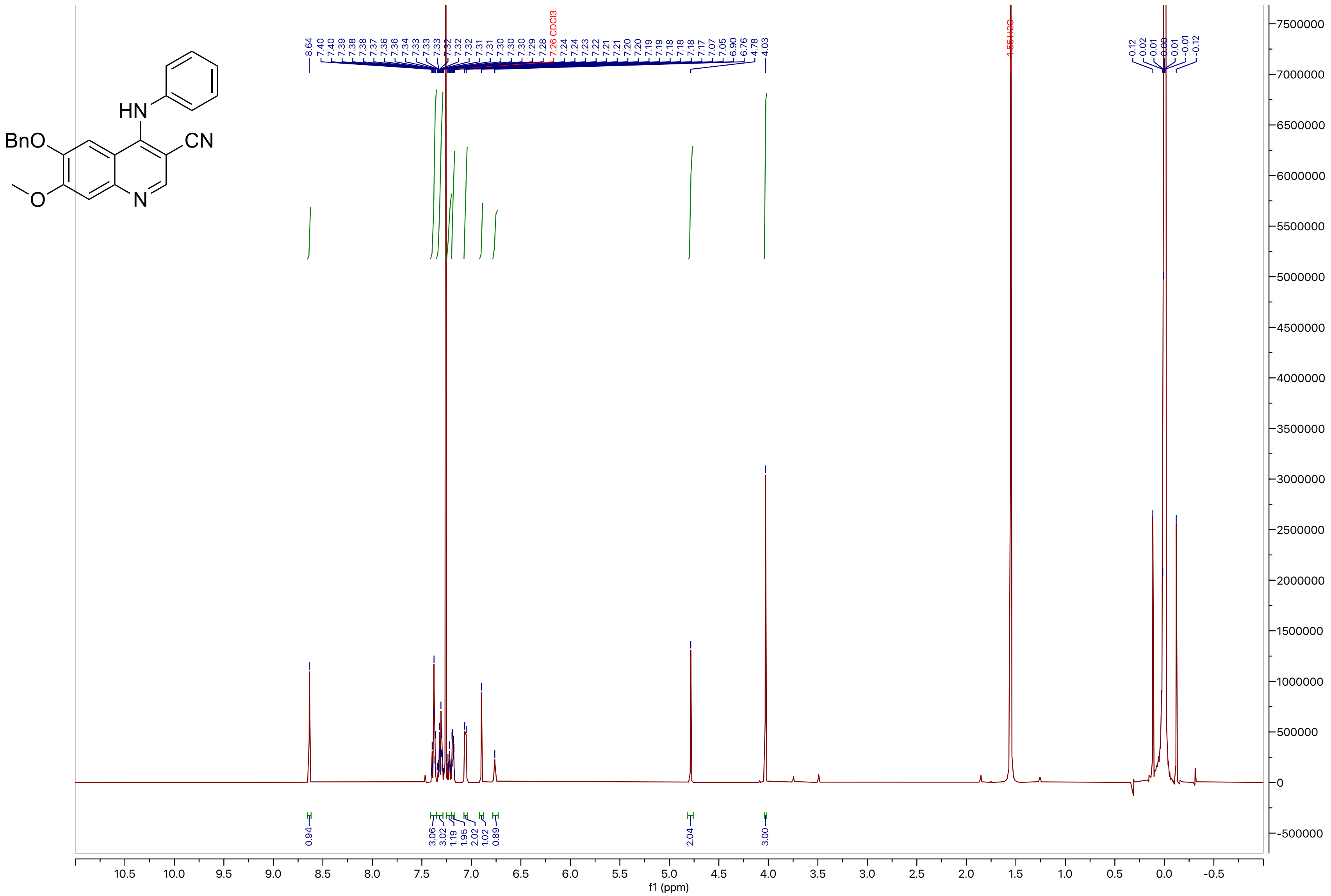

**Compound 13**

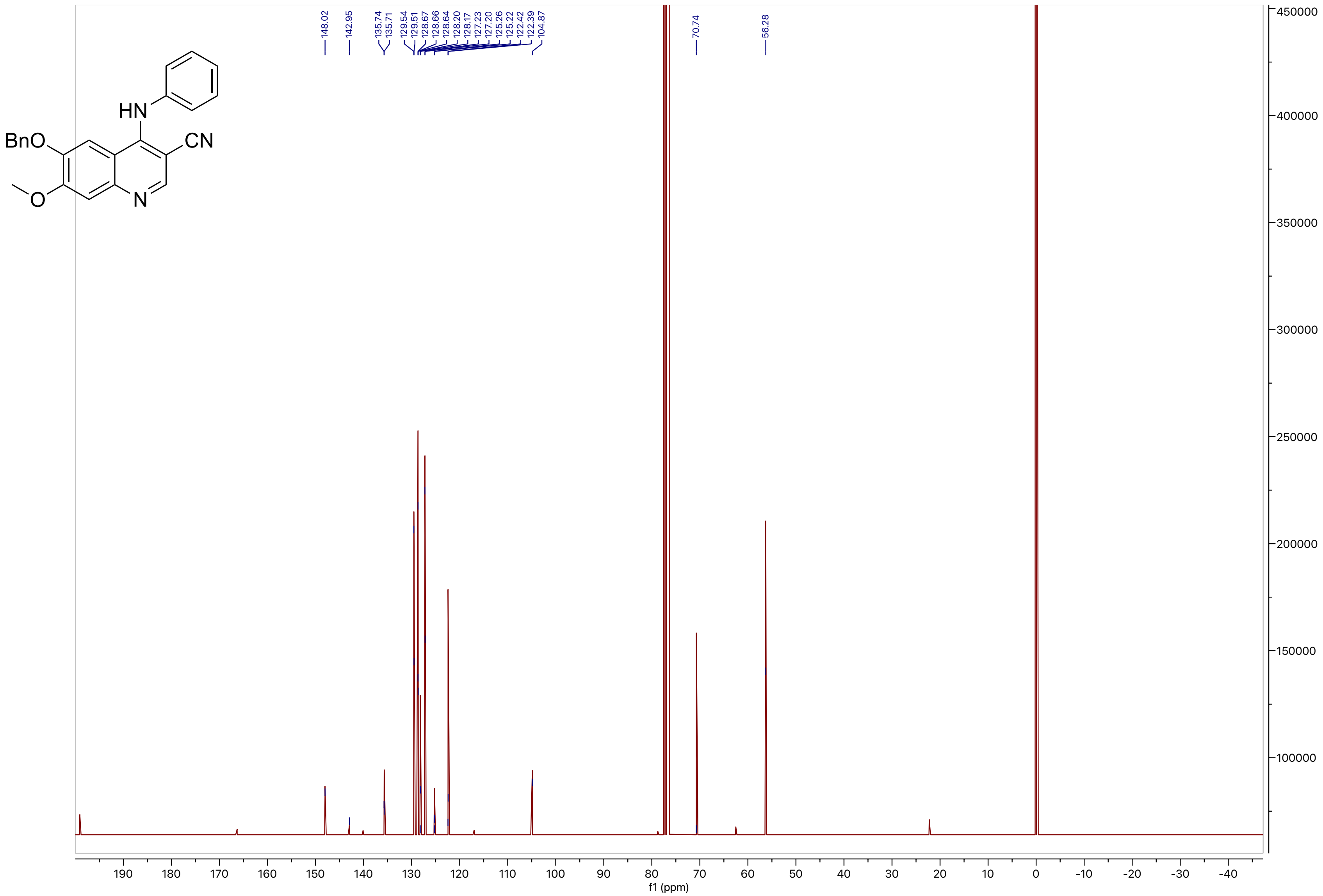
148.02
142.95
135.74
135.71
129.54
129.51
128.67
128.66
128.64
128.20
128.17
127.23
127.20
125.26
125.22
122.42
122.39
104.87
70.74
56.28
f1 (ppm)

**Compound 14**

LCMS

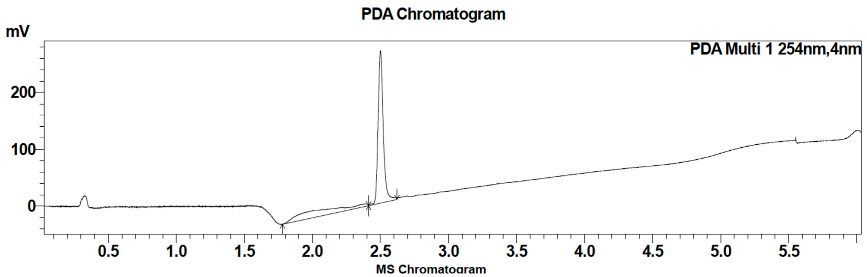

| Peak # | Ret. time | Area | Area % |
| --- | --- | --- | --- |
| 1 | 2.5 | 710734 | 100.00 |

HRMS

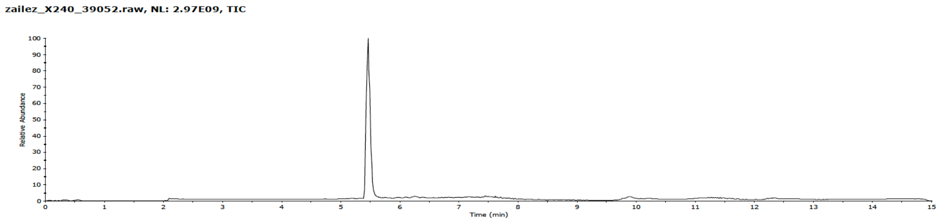

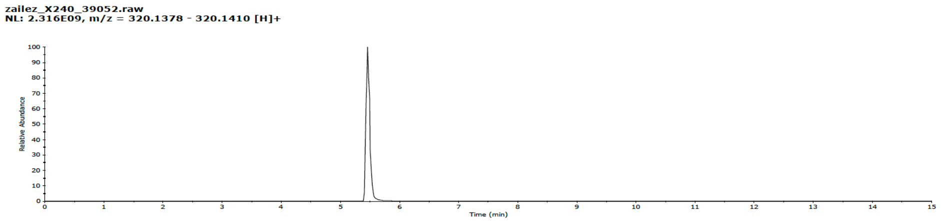

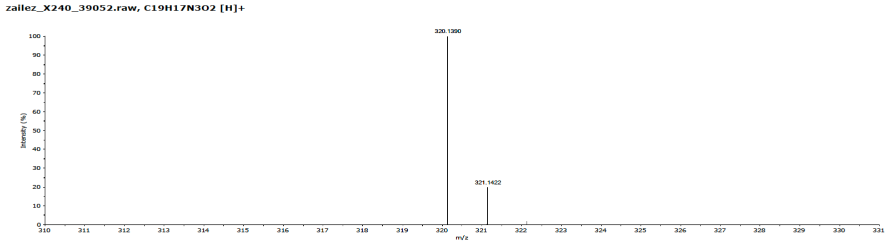

**Compound 14**

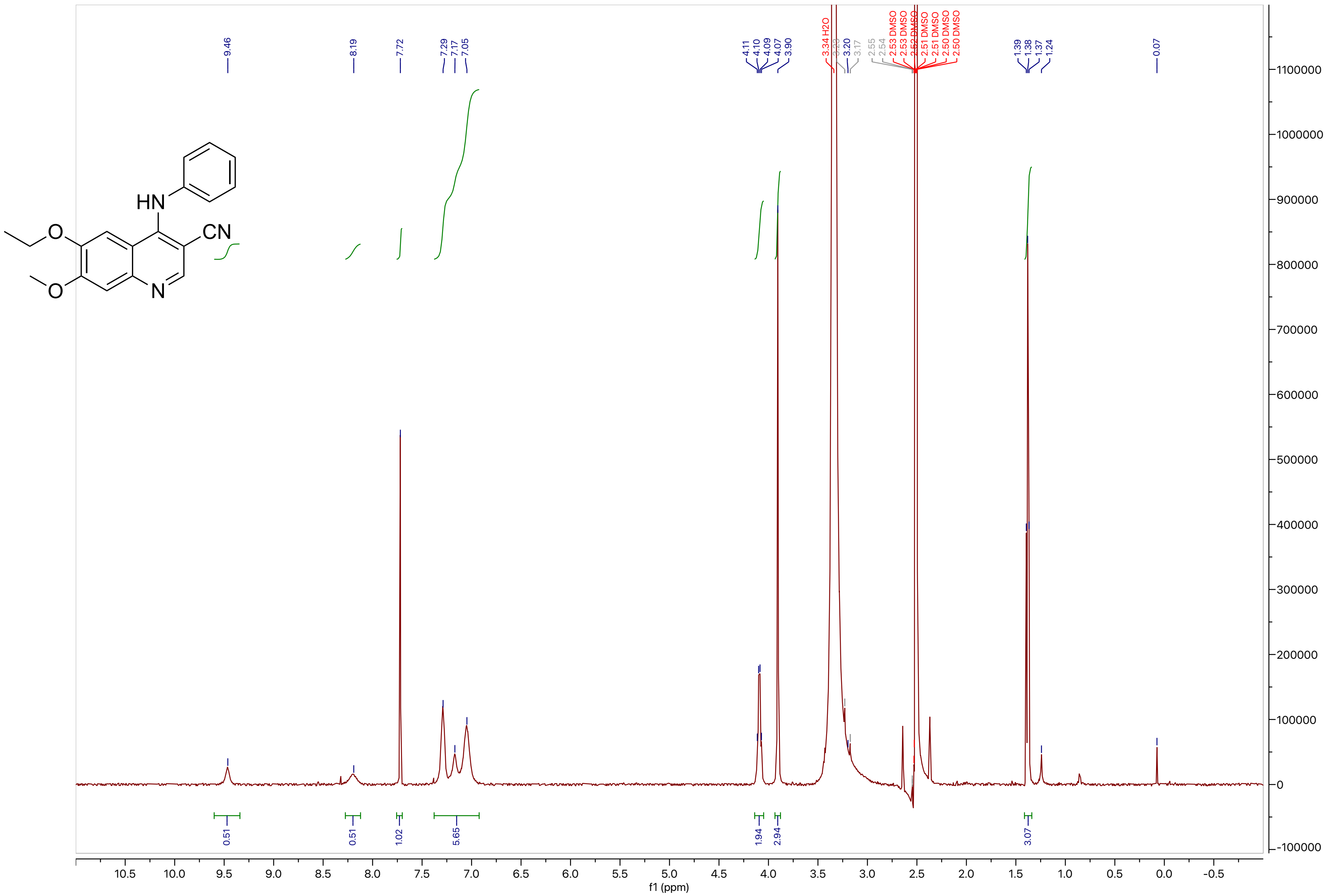

HN
CN
O
O
N
9.46
8.19
7.72
7.29
7.17
7.05
4.11
4.10
4.09
4.07
3.90
3.34 H2O
3.20
3.17
2.55
2.54
2.53 DMSO
2.53 DMSO
2.52 DMSO
2.51 DMSO
2.51 DMSO
2.50 DMSO
2.50 DMSO
1.39
1.38
1.37
1.24
0.07
0.51
0.51
1.02
5.65
1.94
2.94
3.07
f1 (ppm)

**Compound 14**

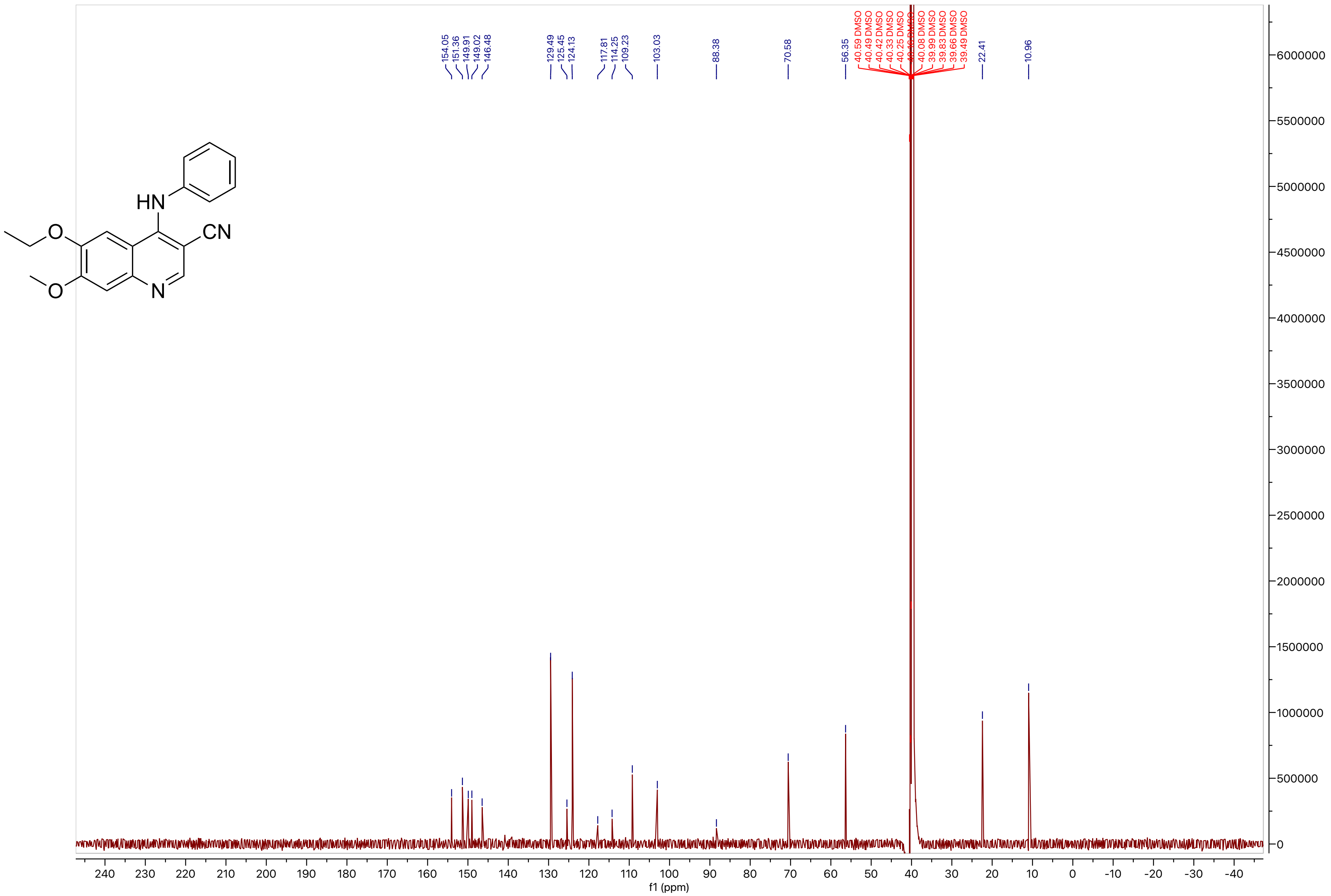
HN
O
CN
O
N
154.05
151.36
149.91
149.02
146.48
129.49
125.45
124.13
117.81
114.25
109.23
103.03
88.38
70.58
56.35
40.59 DMSO
40.49 DMSO
40.42 DMSO
40.33 DMSO
40.25 DMSO
40.08 DMSO
39.99 DMSO
39.83 DMSO
39.66 DMSO
39.49 DMSO
22.41
10.96
f1 (ppm)

#### Compound 15

LCMS

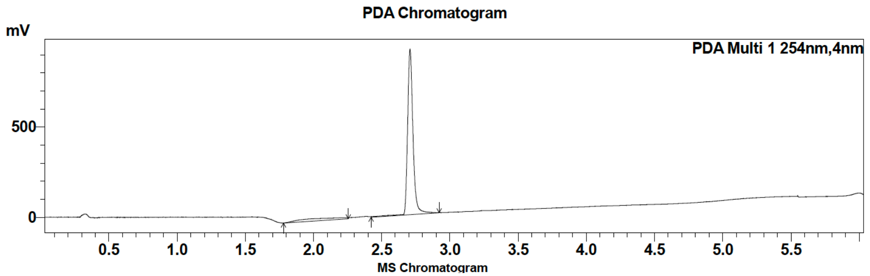

| Peak # | Ret. time | Area | Area % |
|---|---|---|---|
| 1 | 2.2 | 109543 | 4.15 |
| 2 | 2.7 | 2528823 | 95.85 |

HRMS

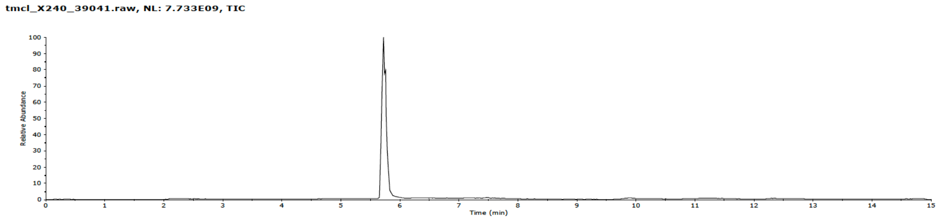

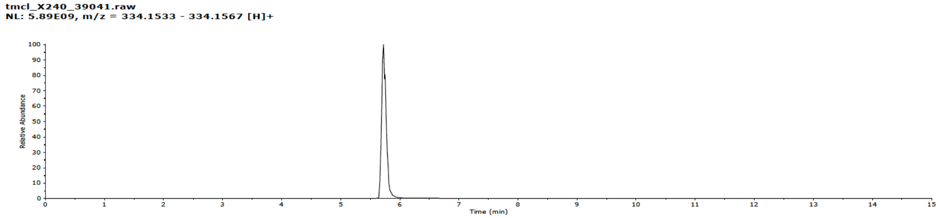

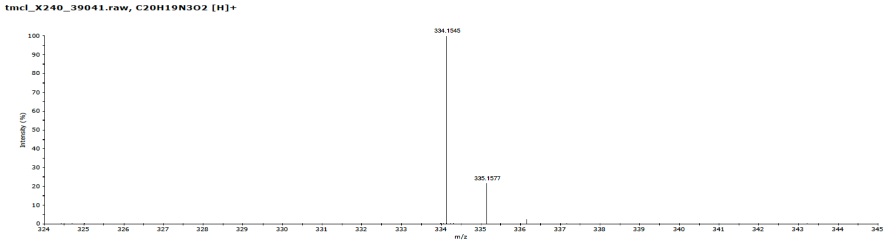

**Compound 15**

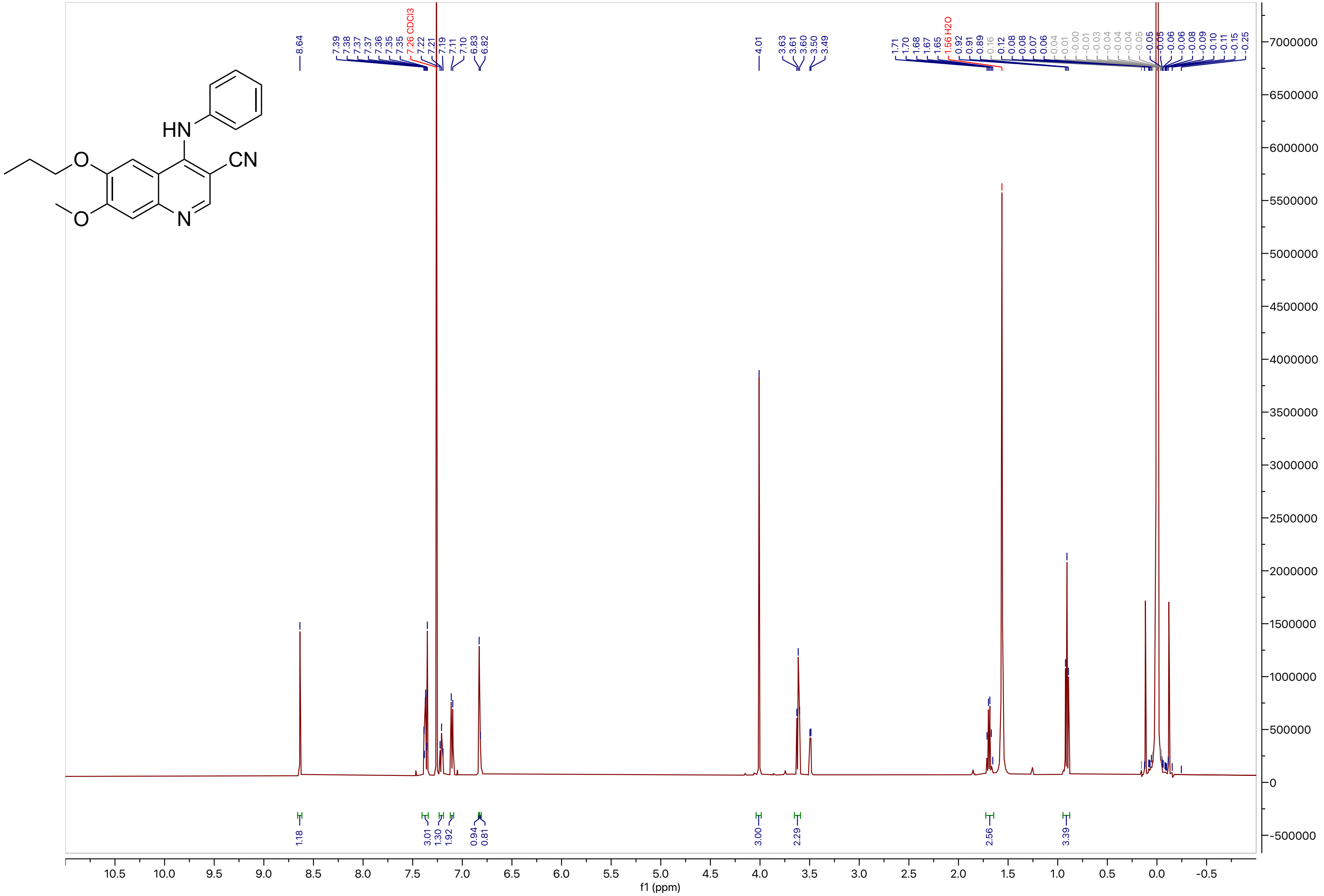

**Compound 15**

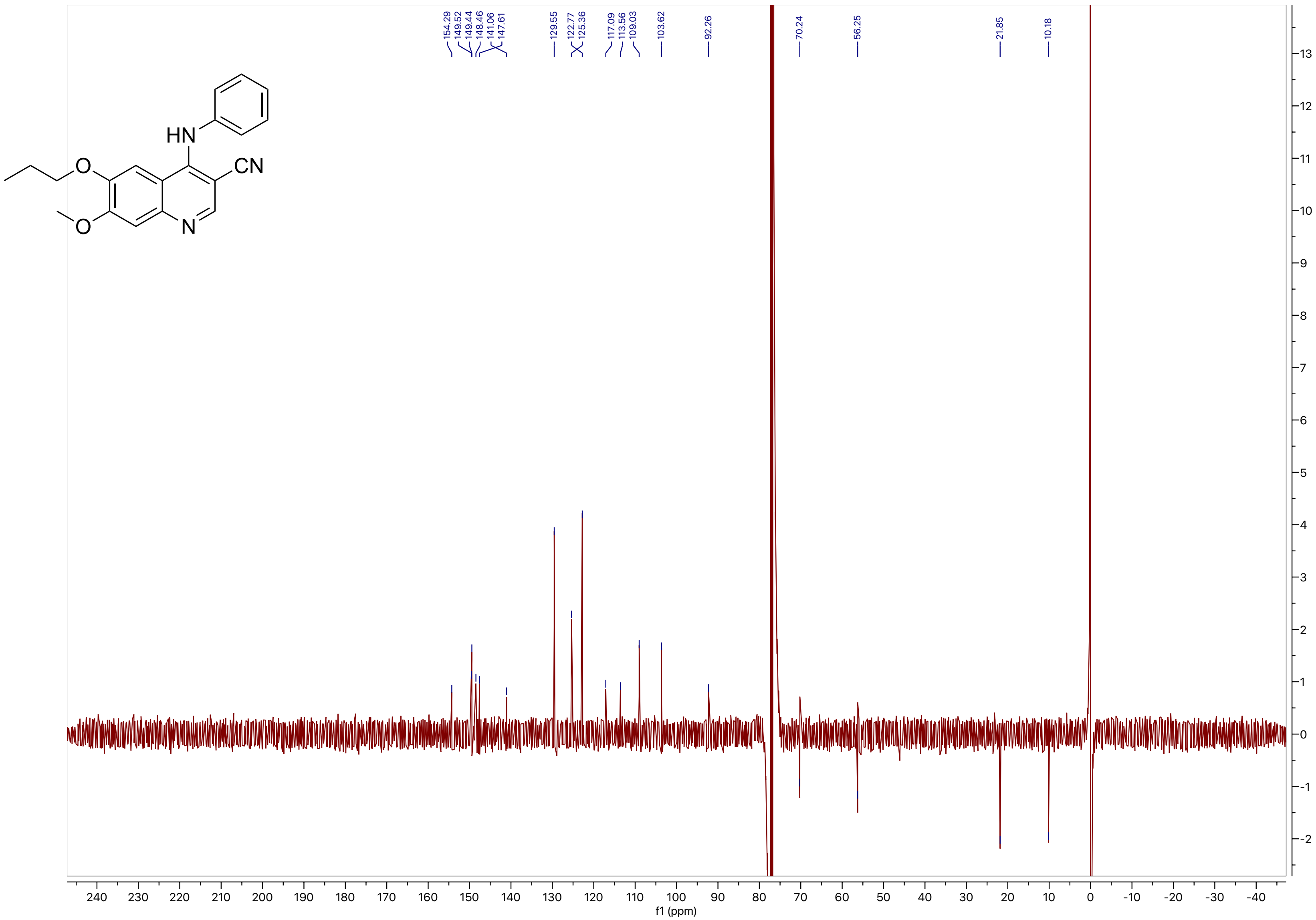
HN
O
CN
O
N
154.29
149.52
149.44
148.46
141.06
147.61
129.55
122.77
125.36
117.09
113.56
109.03
103.62
92.26
70.24
56.25
21.85
10.18
f1 (ppm)

**Compound 16**

LCMS

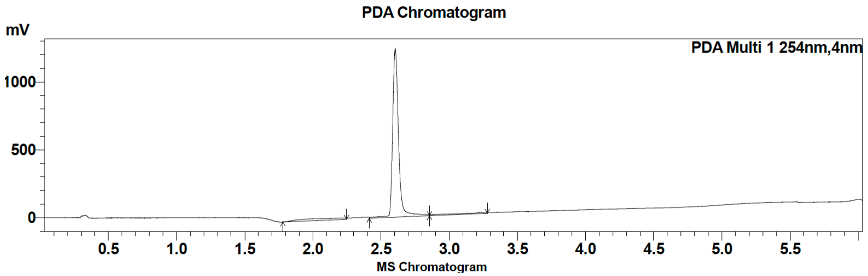

| Peak # | Ret. time | Area | Area % |
|---|---|---|---|
| 1 | 2.6 | 3754637 | 96.07 |
| 2 | 3.2 | 153574 | 3.93 |

HRMS

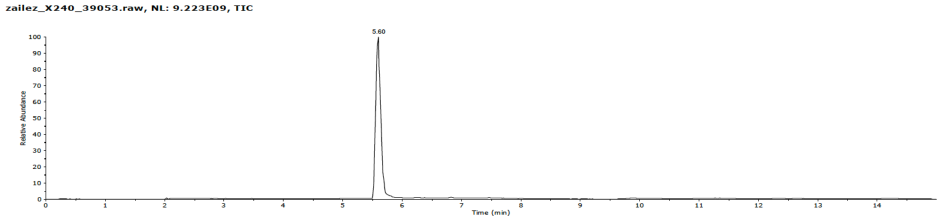

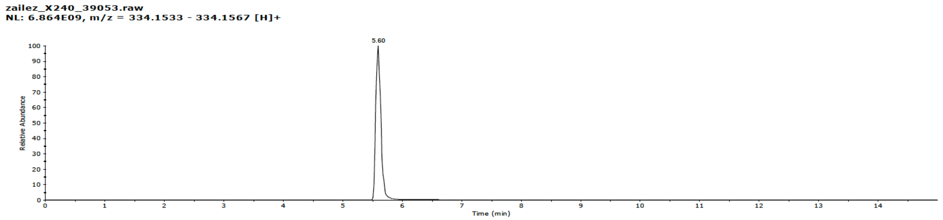

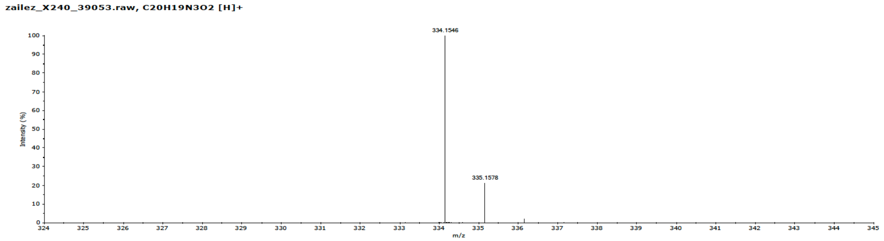

**Compound 16**

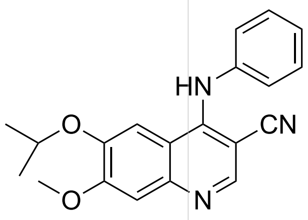

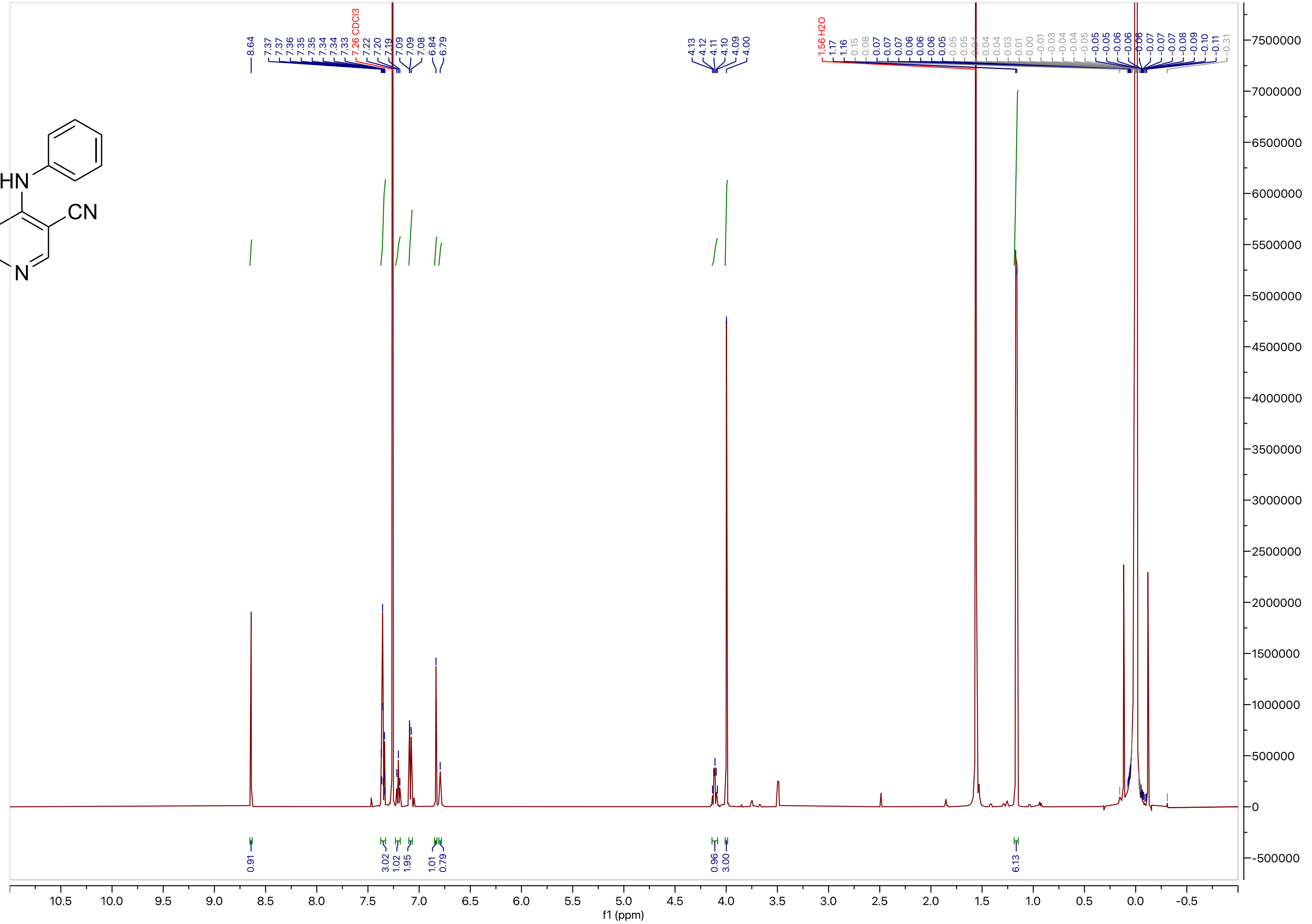

**Compound 16**

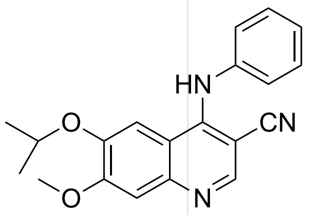
HN
O
CN
O
N

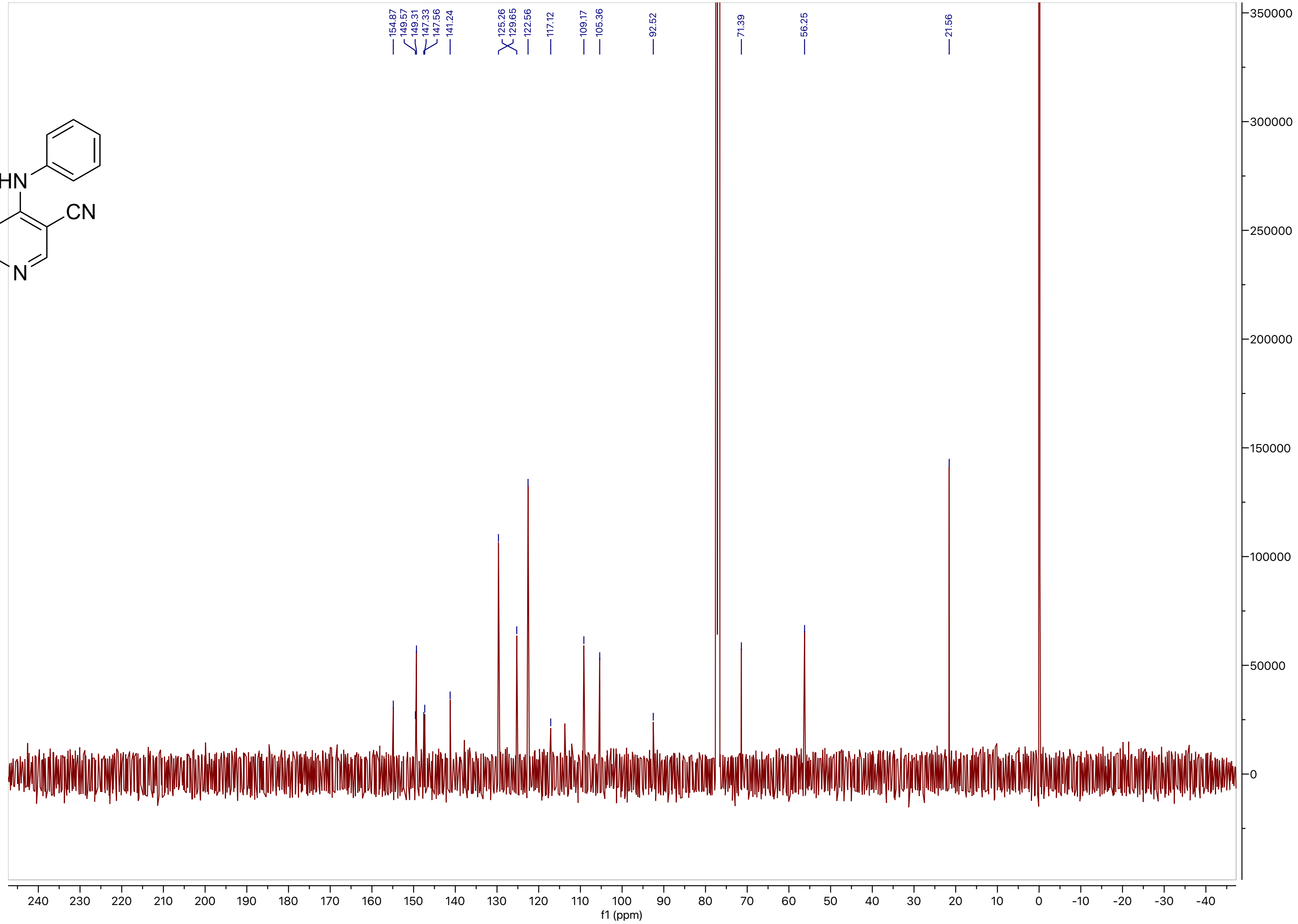
154.87
149.57
149.31
147.33
147.56
141.24
125.26
129.65
122.56
117.12
109.17
105.36
92.52
71.39
56.25
21.56
f1 (ppm)

**Compound 17**

LCMS

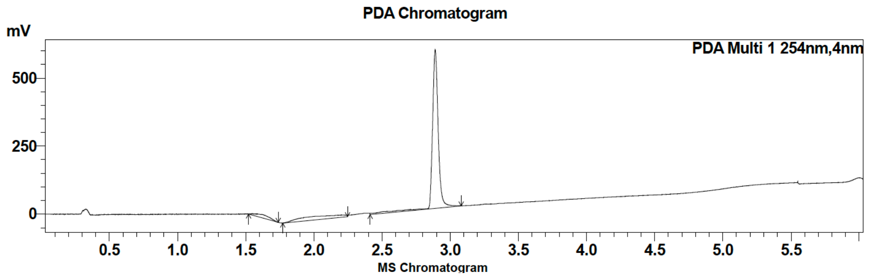

| Peak # | Ret. time | Area | Area % |
|---|---|---|---|
| 1 | 1.5 | 88220 | 4.80 |
| 2 | 2.9 | 1749491 | 95.20 |

HRMS

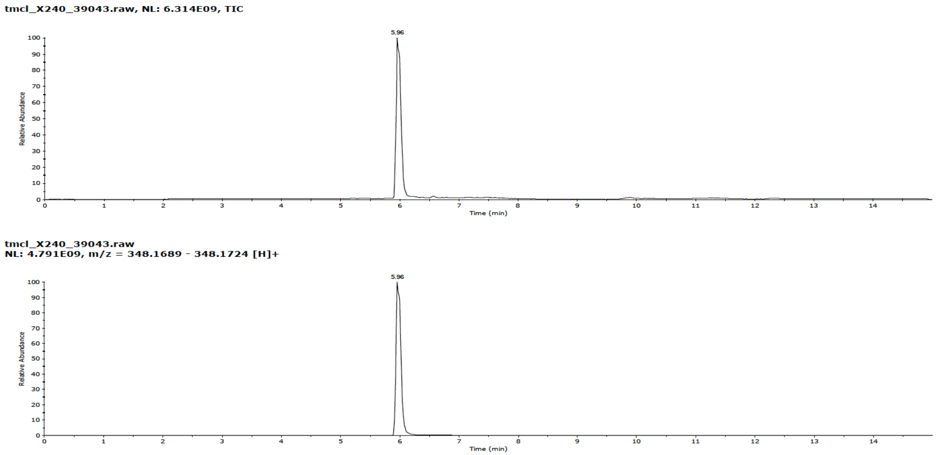

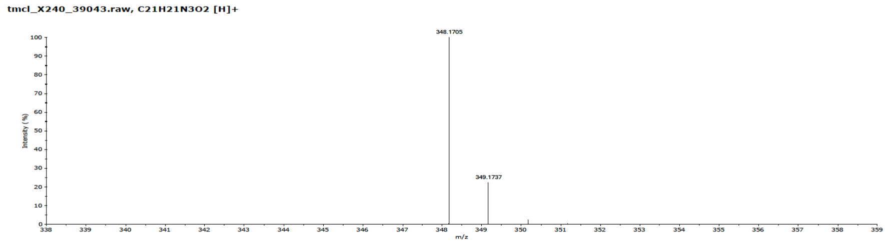

**Compound 17**

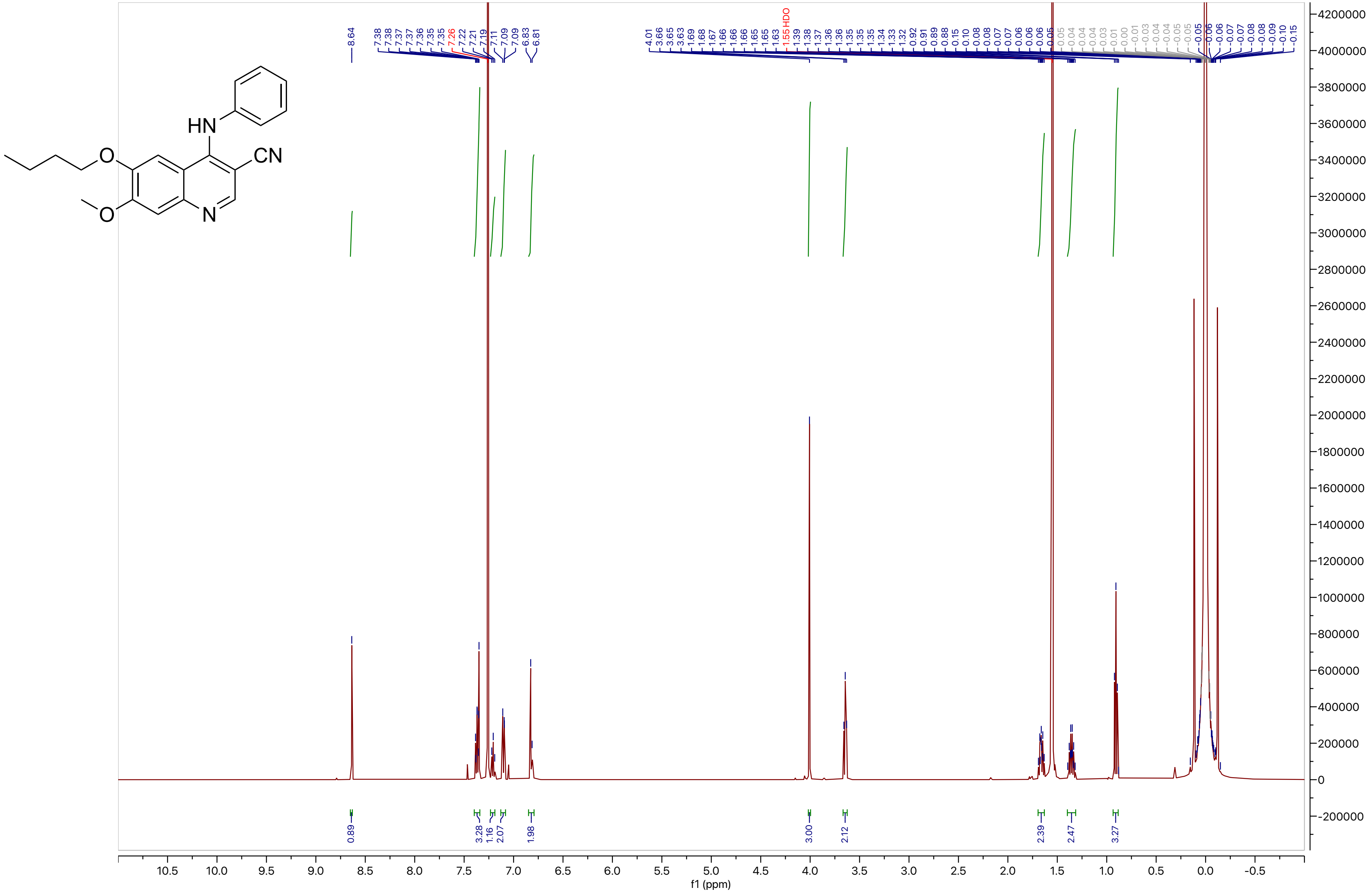

**Compound 17**

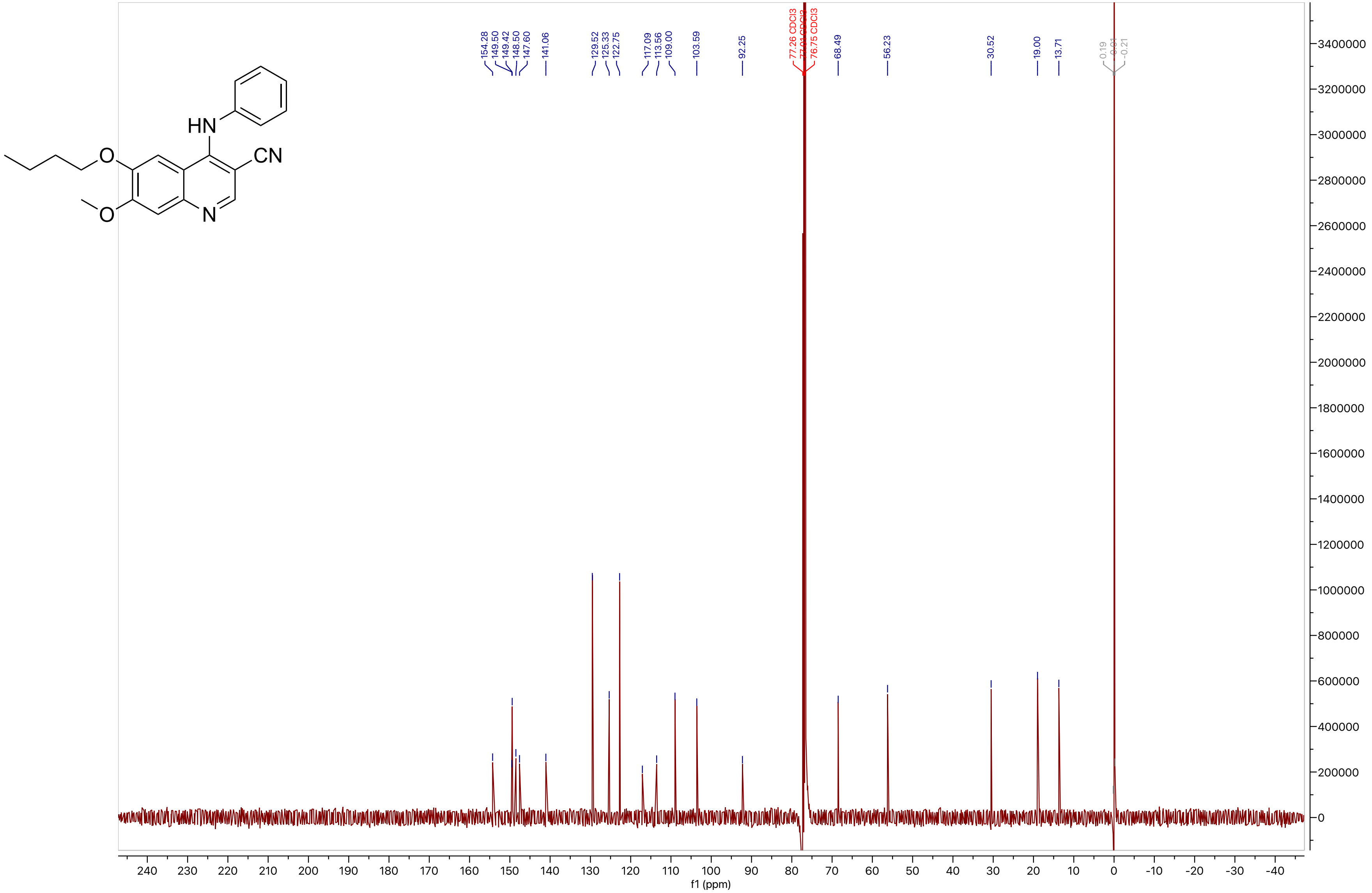
HN
CN
O
O
N
154.28
149.50
149.42
148.50
147.60
141.06
129.52
125.33
122.75
117.09
113.56
109.00
103.59
92.25
77.26 CDCl3
76.75 CDCl3
68.49
56.23
30.52
19.00
13.71
0.19
-0.21
f1 (ppm)

**Compound 18**

LCMS

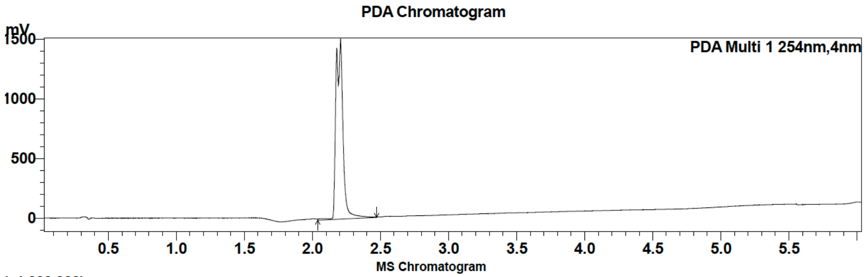

| Peak # | Ret. time | Area | Area % |
|---|---|---|---|
| 1 | 2.2 | 5399472 | 100.00 |

HRMS

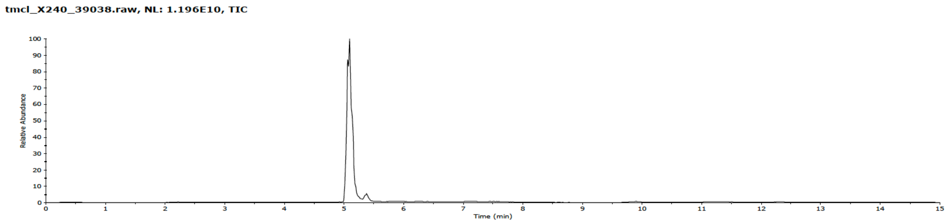

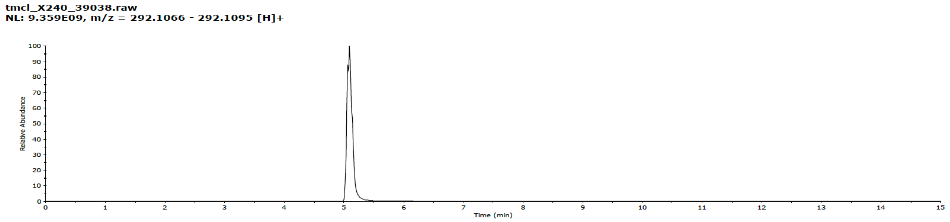

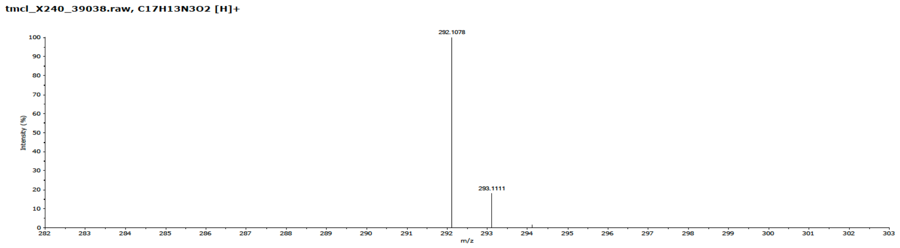

**Compound 18**

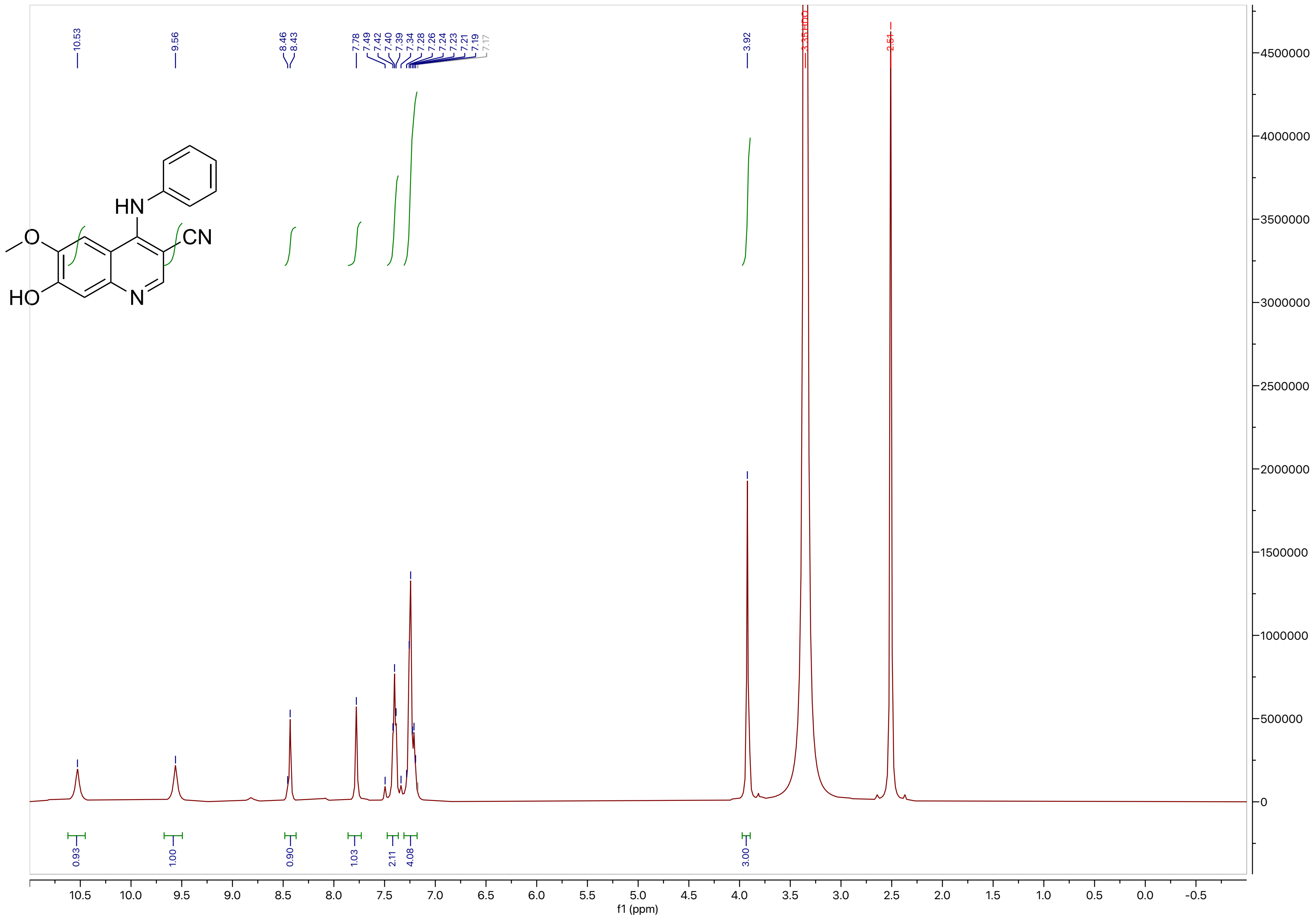

HN
CN
O
HO
N
10.53
9.56
8.46
8.43
7.78
7.49
7.42
7.40
7.39
7.34
7.28
7.26
7.24
7.23
7.21
7.19
7.17
3.92
3.35 HDO
2.51
0.93
1.00
0.90
1.03
2.11
4.08
3.00
f1 (ppm)

**Compound 18**

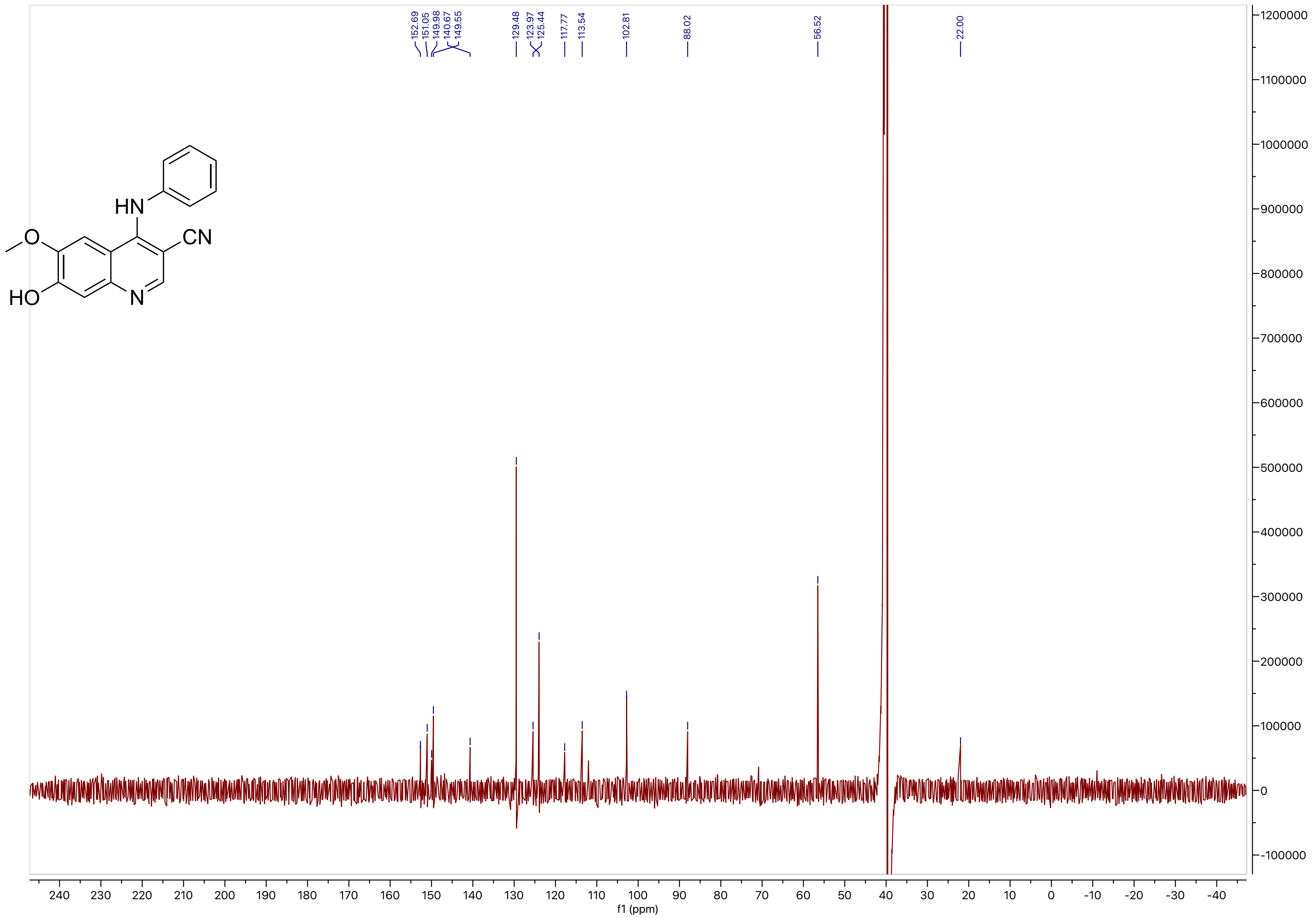

152.69
151.05
149.98
140.67
149.55
129.48
123.97
125.44
117.77
113.54
102.81
88.02
56.52
22.00
HN
CN
O
HO
N
f1 (ppm)

#### Compound 19

LCMS

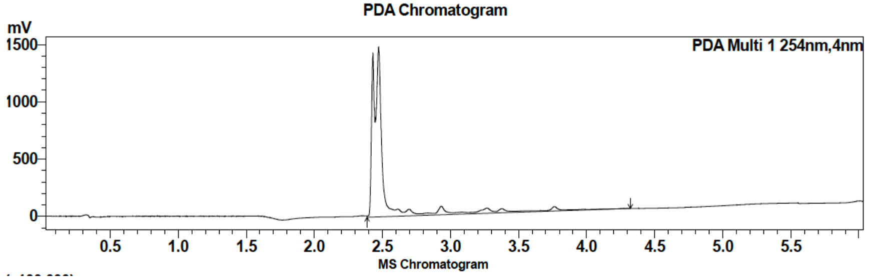

| Peak # | Ret. time | Area | Area % |
|---|---|---|---|
| 1 | 2.5 | 7130109 | 100.00 |

HRMS

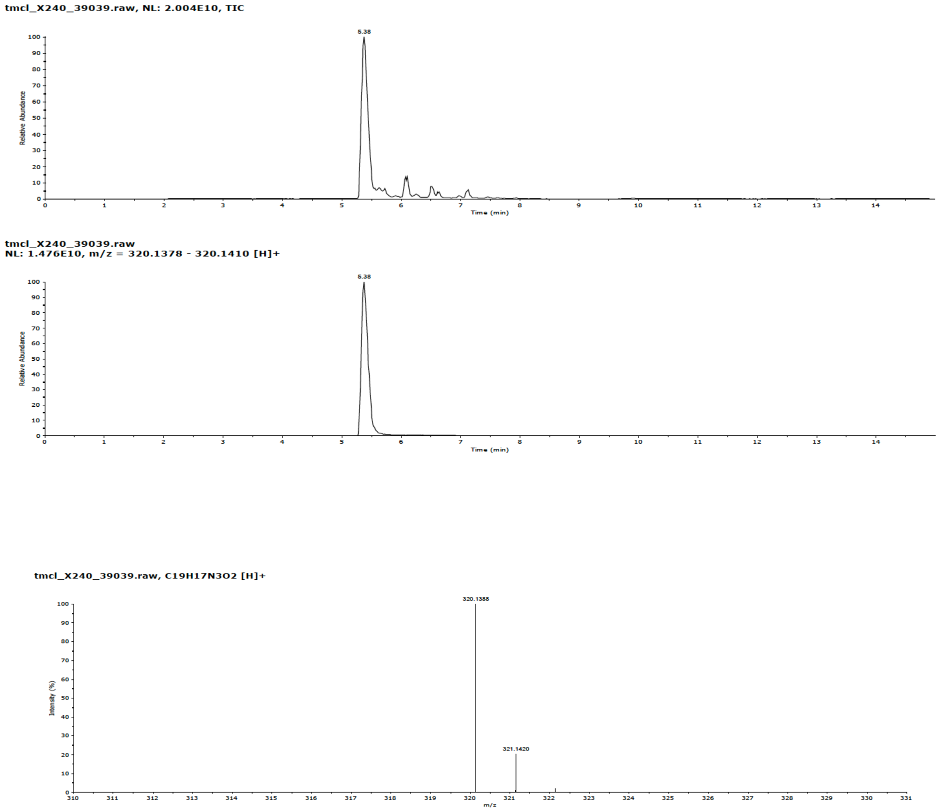

**Compound 19**

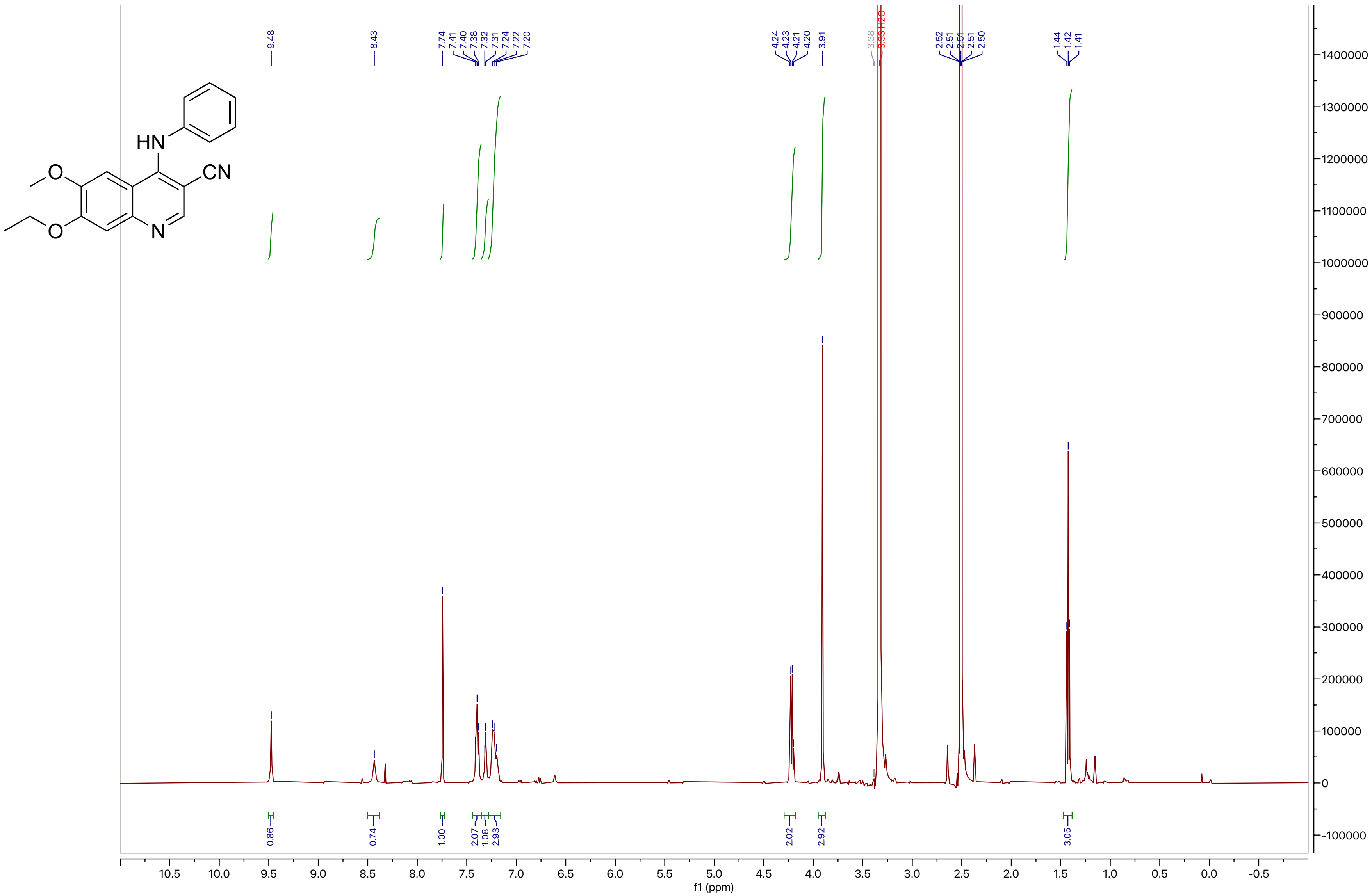
HN
CN
O
O
N
9.48
8.43
7.74
7.41
7.40
7.38
7.32
7.31
7.24
7.22
7.20
4.24
4.23
4.21
4.20
3.91
3.38
3.33 H2O
2.52
2.51
2.51
2.51
2.50
1.44
1.42
1.41
0.86
0.74
1.00
2.07
1.08
2.93
2.02
2.92
3.05
f1 (ppm)

#### Compound 19

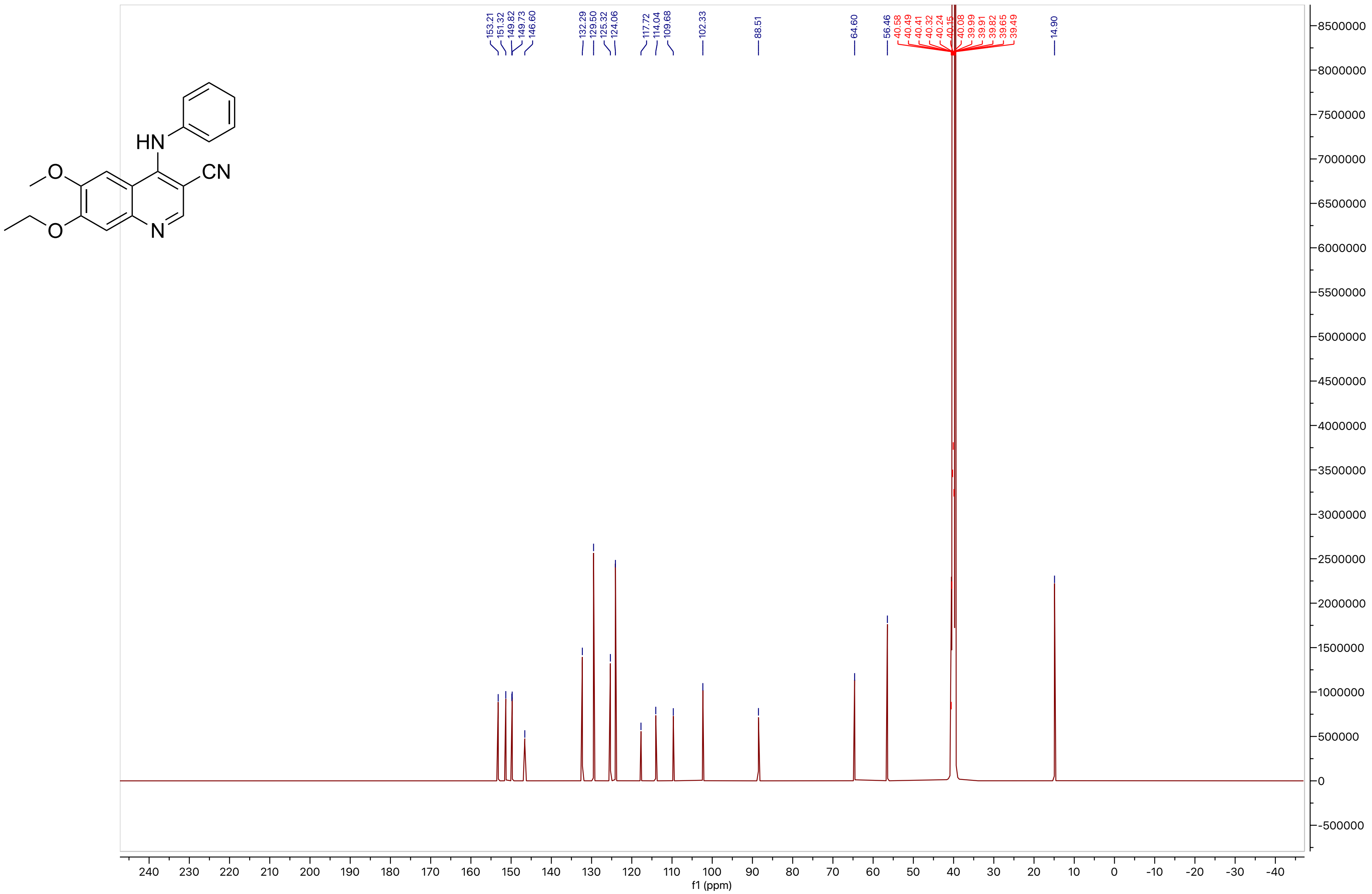
HN
CN
O
O
N
153.21
151.32
149.82
149.73
146.60
132.29
129.50
125.32
124.06
117.72
114.04
109.68
102.33
88.51
64.60
56.46
40.58
40.49
40.41
40.32
40.24
40.15
40.08
39.99
39.91
39.82
39.65
39.49
14.90
f1 (ppm)

**Compound 20**

LCMS

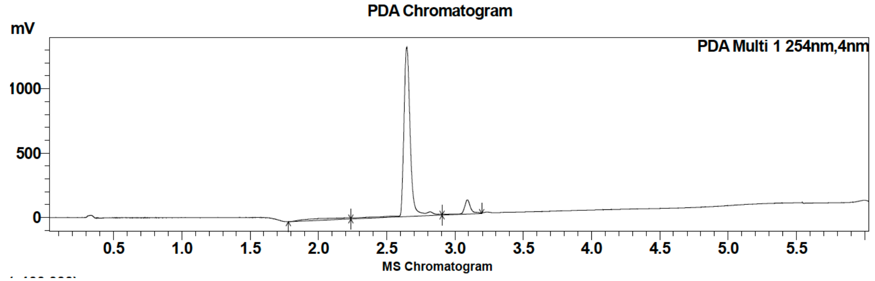

| Peak # | Ret. time | Area | Area % |
|---|---|---|---|
| 1 | 2.6 | 4365953 | 96.01 |
| 2 | 3.1 | 181519 | 3.99 |

HRMS

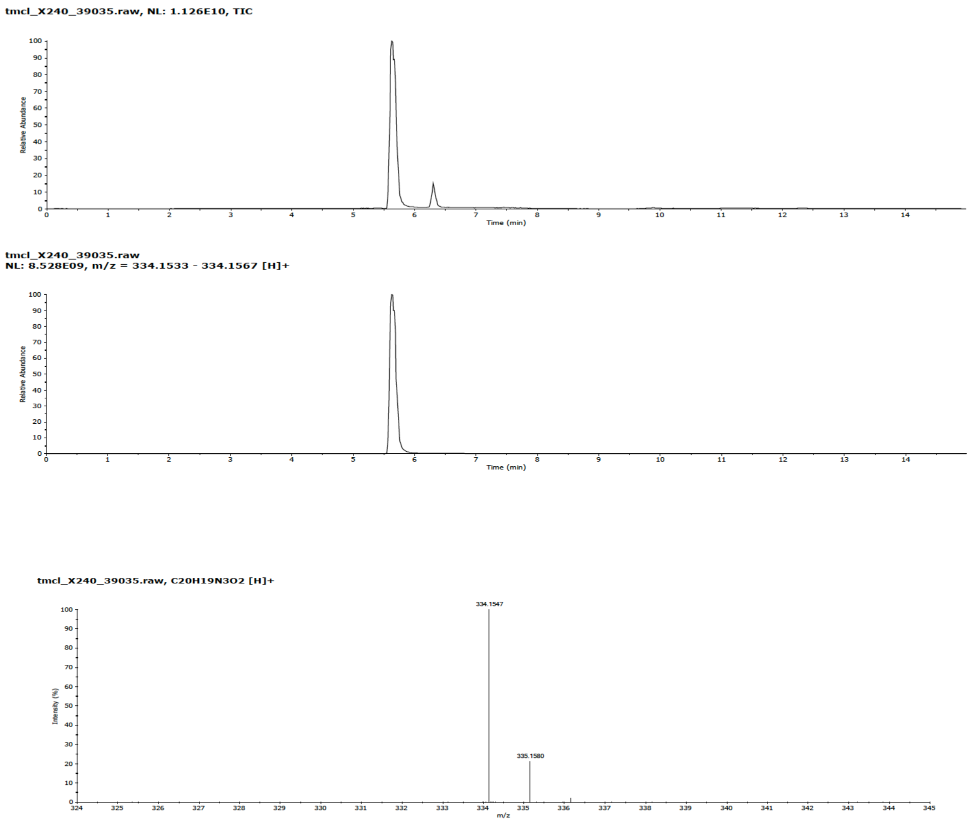

**Compound 20**

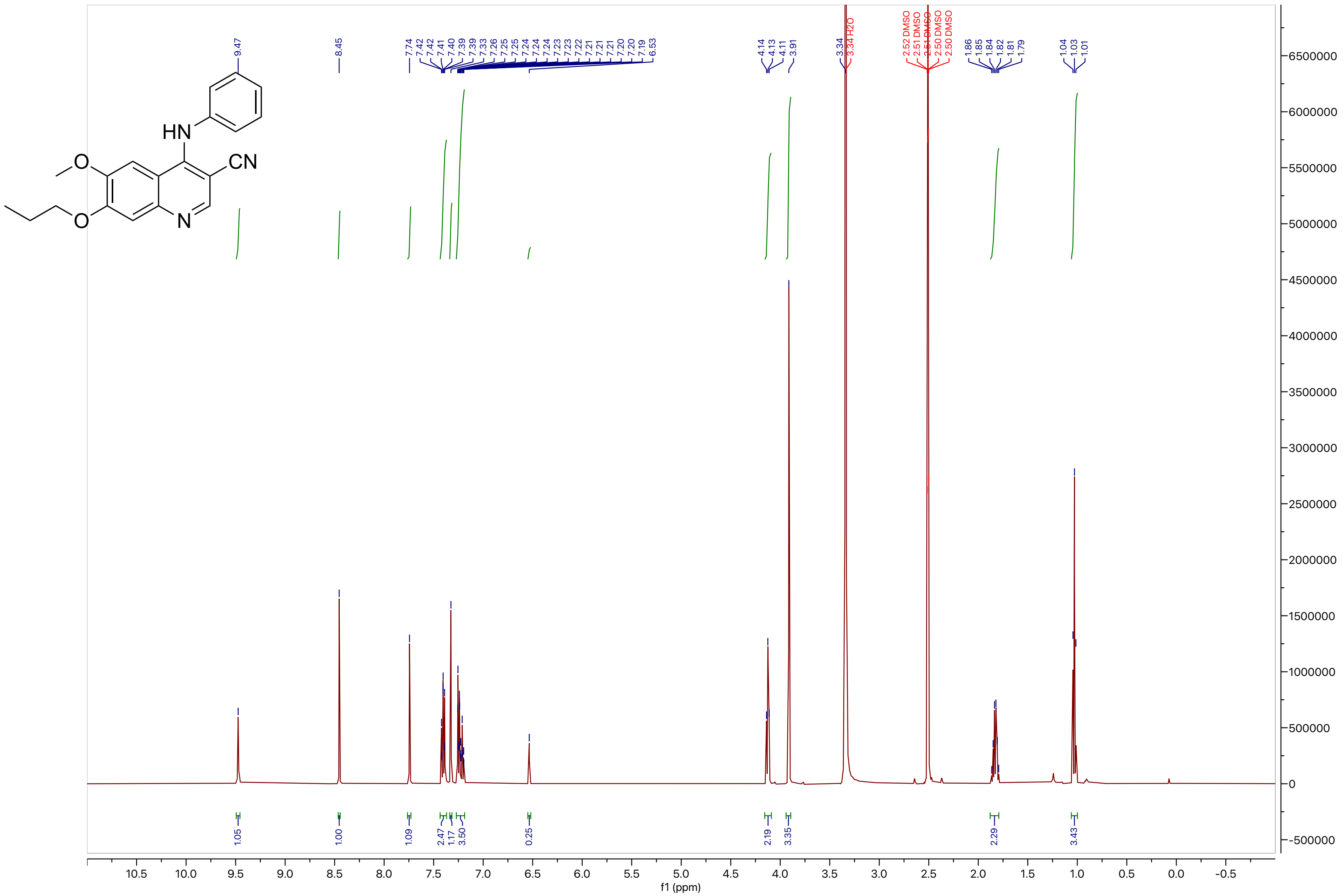

**Compound 20**

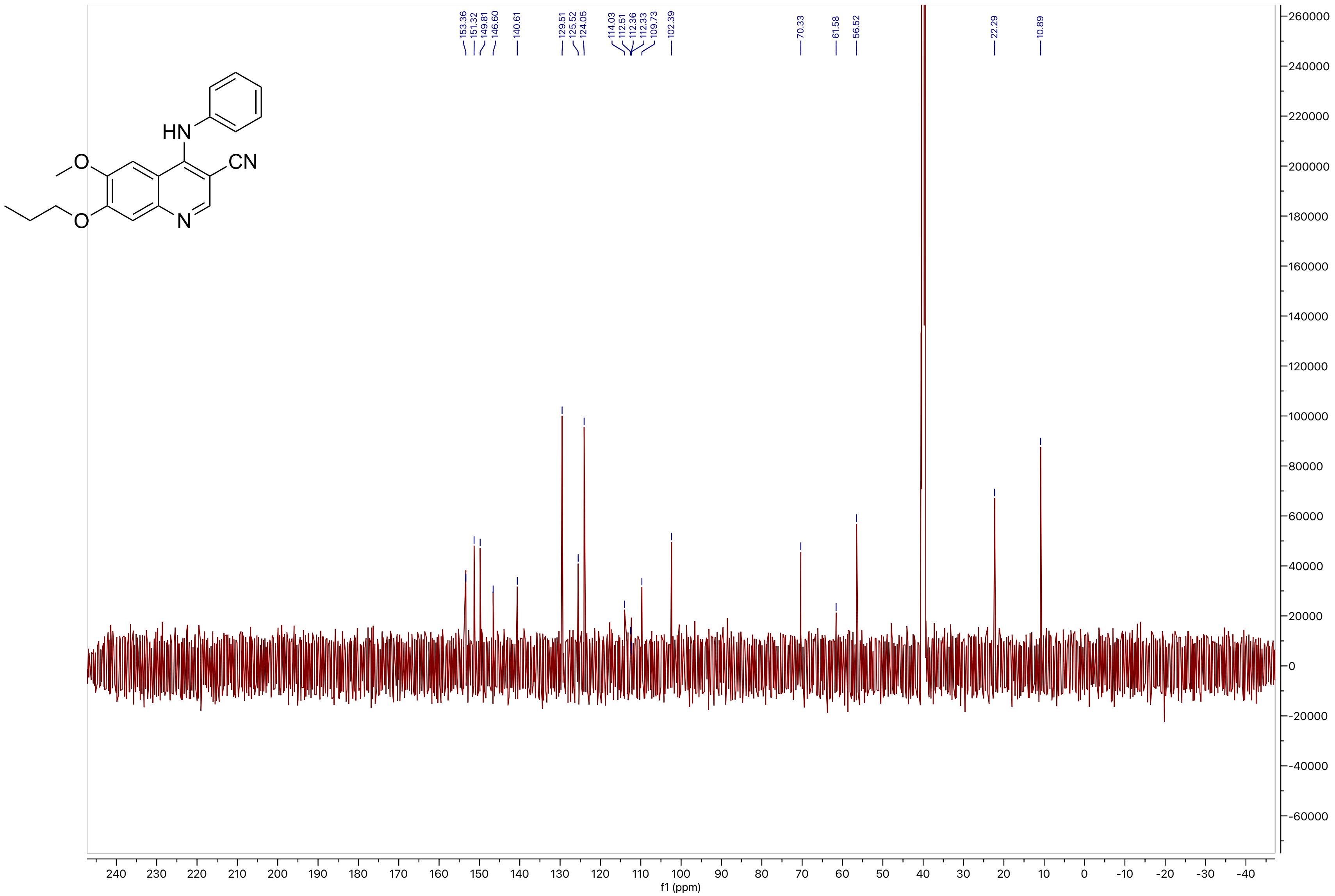
HN
O
CN
O
N
153.36
151.32
149.81
146.60
140.61
129.51
125.52
124.05
114.03
112.51
112.36
112.33
109.73
102.39
70.33
61.58
56.52
22.29
10.89
f1 (ppm)

**Compound 21**

LCMS

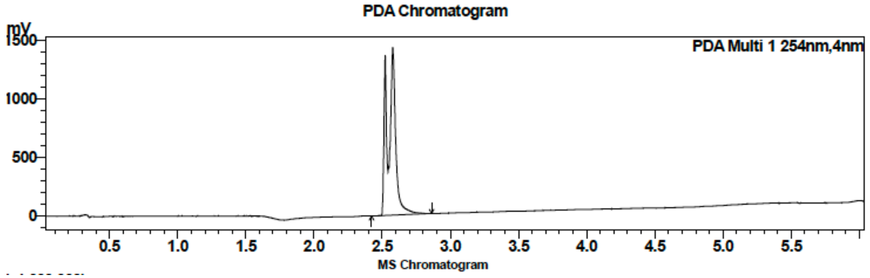

| Peak # | Ret. time | Area | Area % |
|---|---|---|---|
| 1 | 2.6 | 5741339 | 100.00 |

HRMS

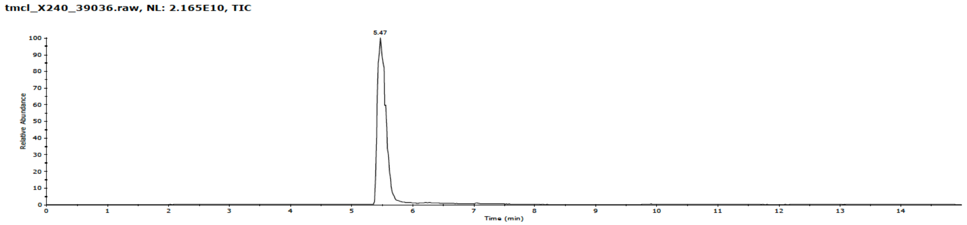

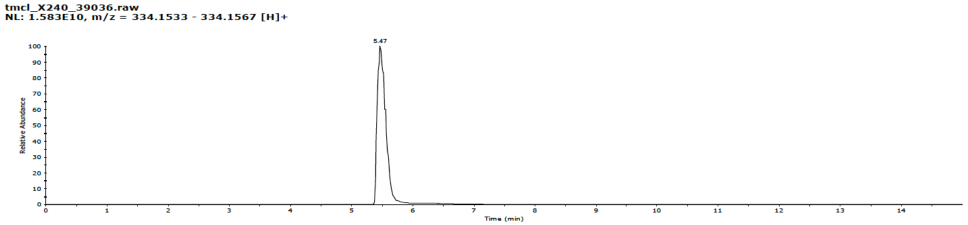

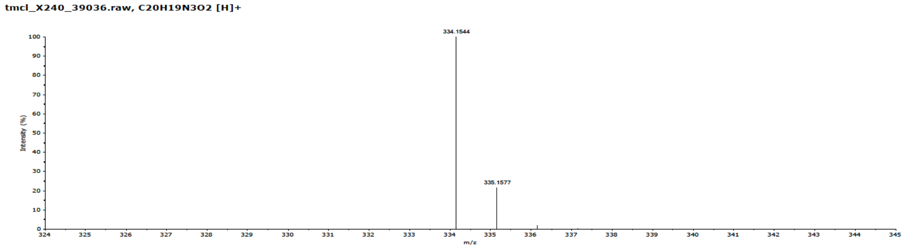

**Compound 21**

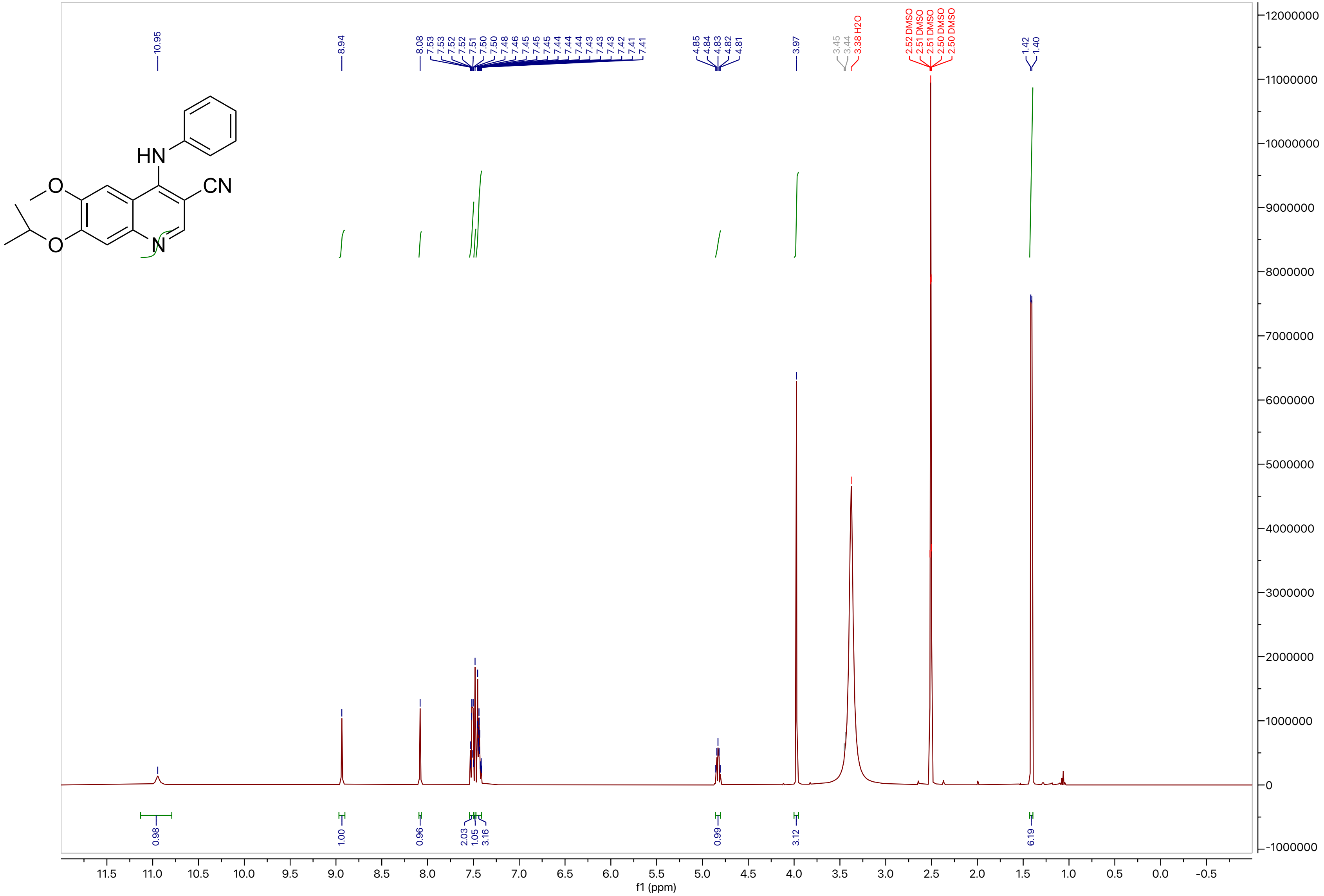

**Compound 21**

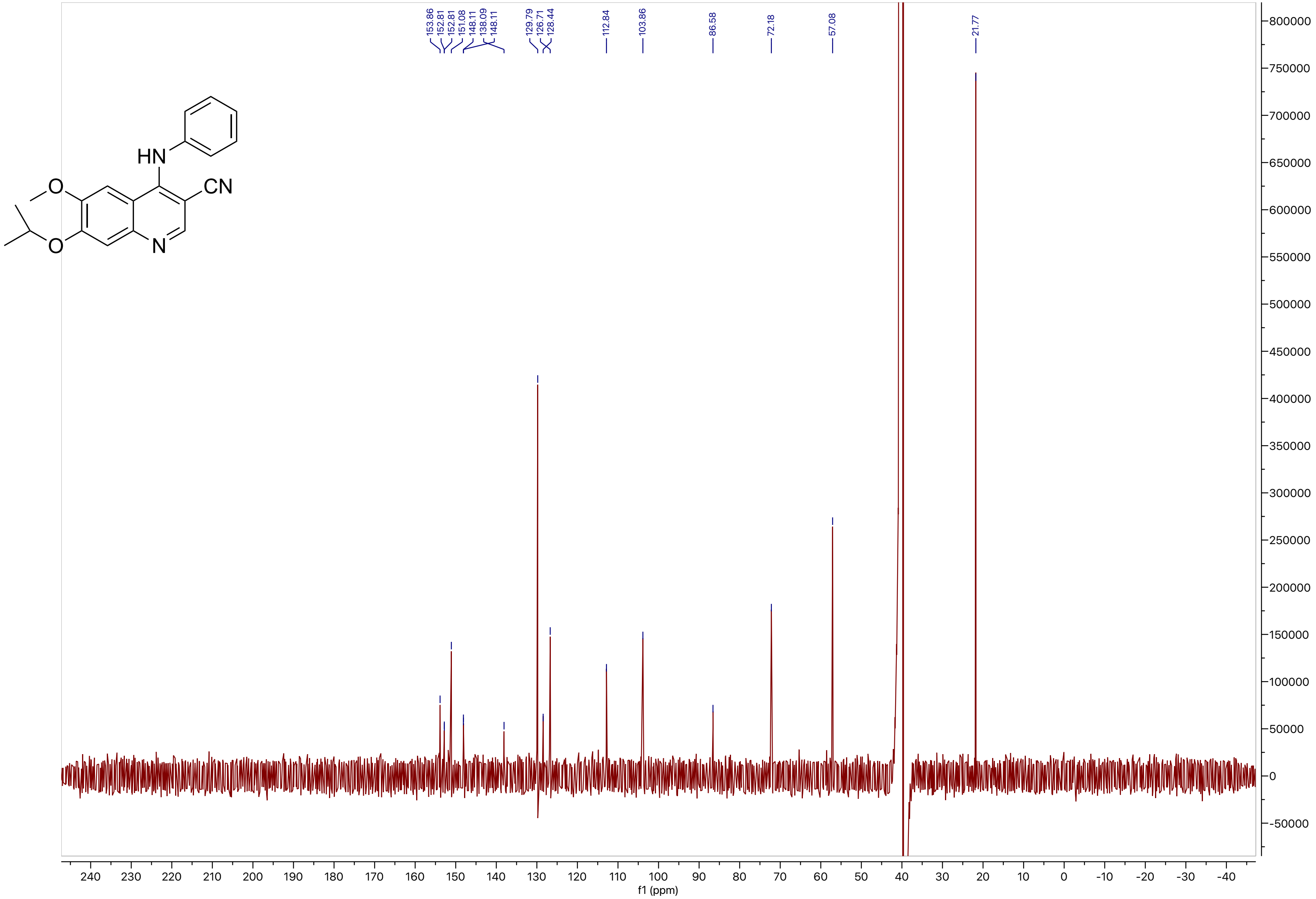

153.86
152.81
152.81
151.08
148.11
138.09
148.11
129.79
126.71
128.44
112.84
103.86
86.58
72.18
57.08
21.77
f1 (ppm)

**Compound 22**

LCMS

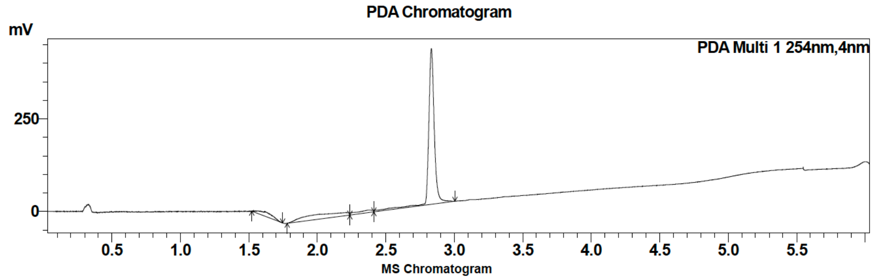

| Peak # | Ret. time | Area | Area % |
|---|---|---|---|
| 1 | 2.4 | 59903 | 4.74 |
| 2 | 2.8 | 1203682 | 95.26 |

HRMS

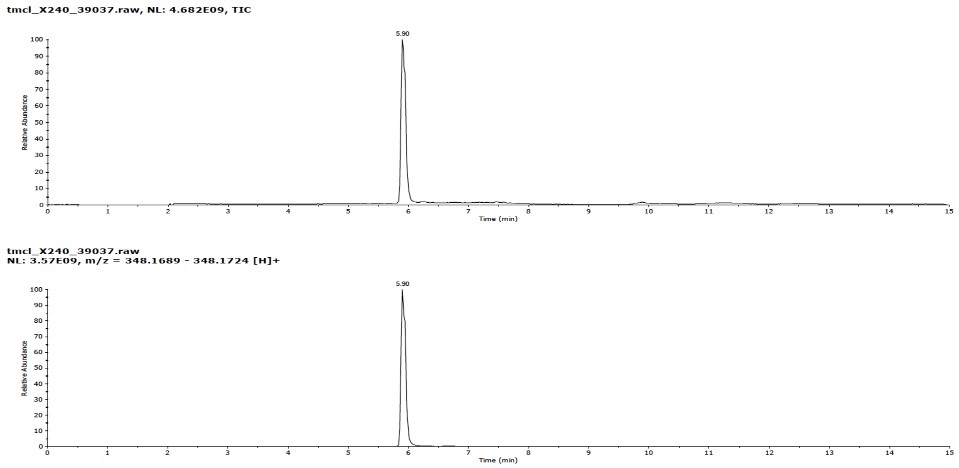

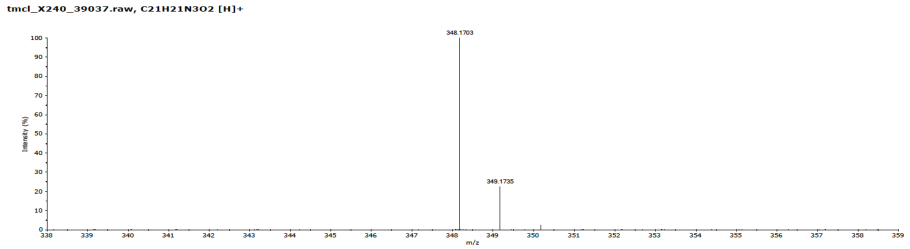

**Compound 22**

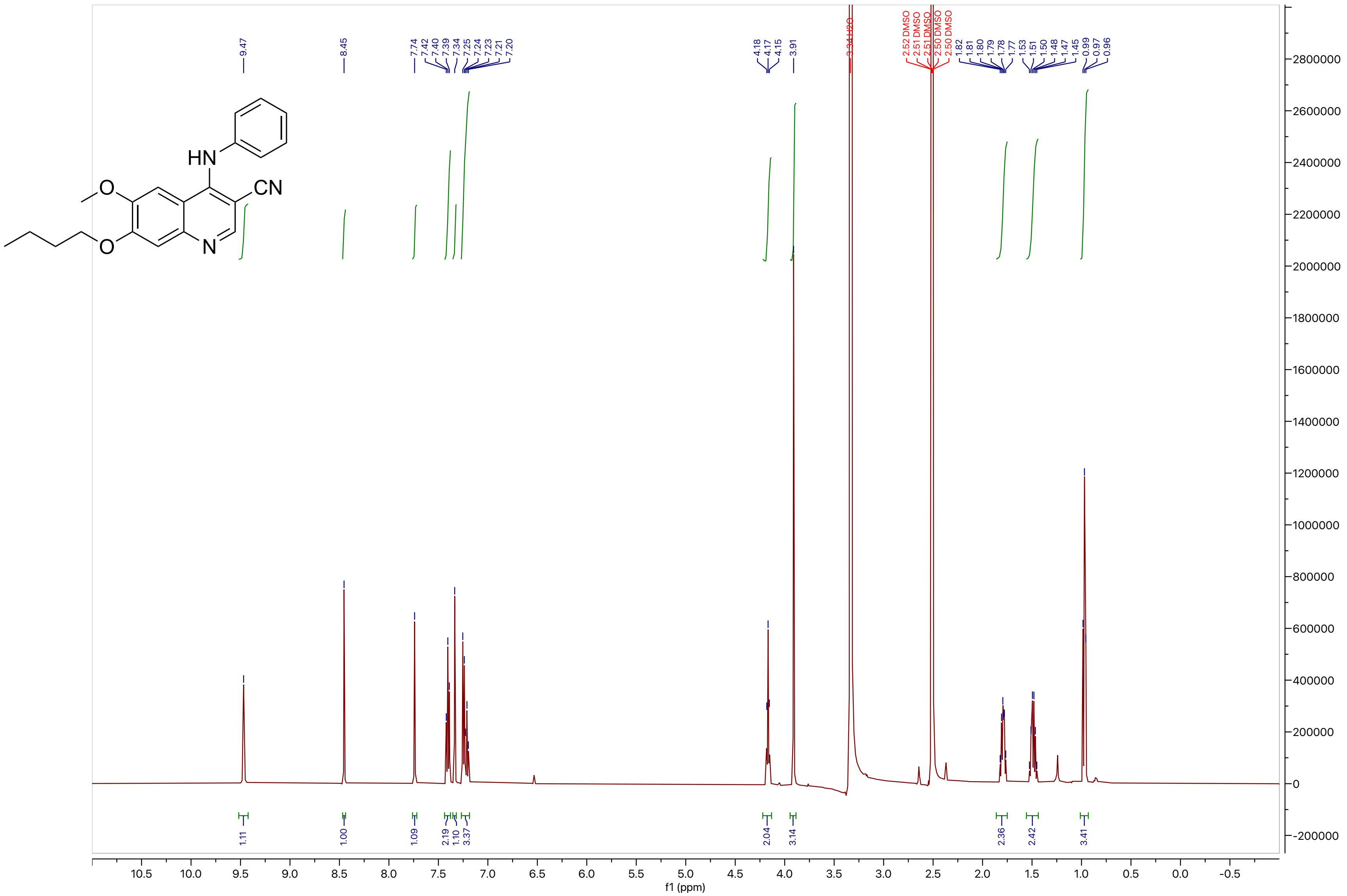

**Compound 22**

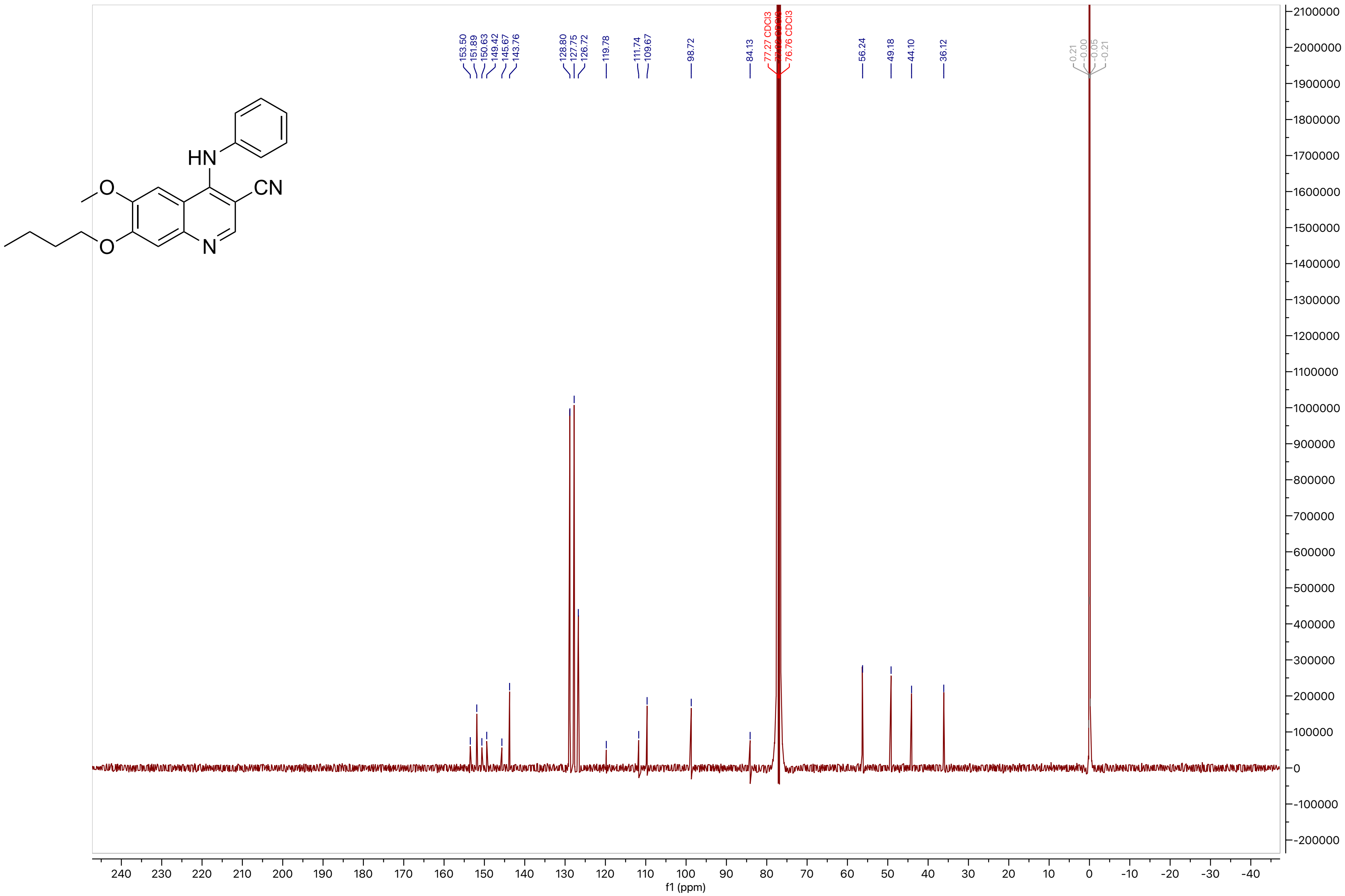

153.50
151.89
150.63
149.42
145.67
143.76
128.80
127.75
126.72
119.78
111.74
109.67
98.72
84.13
77.27 CDCl3
76.76 CDCl3
56.24
49.18
44.10
36.12
f1 (ppm)

**Compound 23**

LCMS

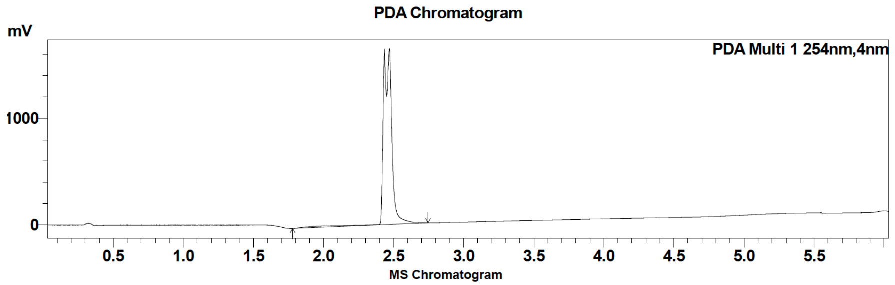

| Peak # | Ret. time | Area | Area % |
|---|---|---|---|
| 1 | 2.4 | 6865164 | 100.00 |

HRMS

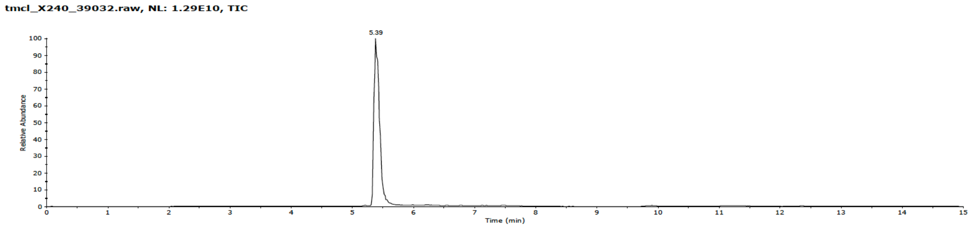

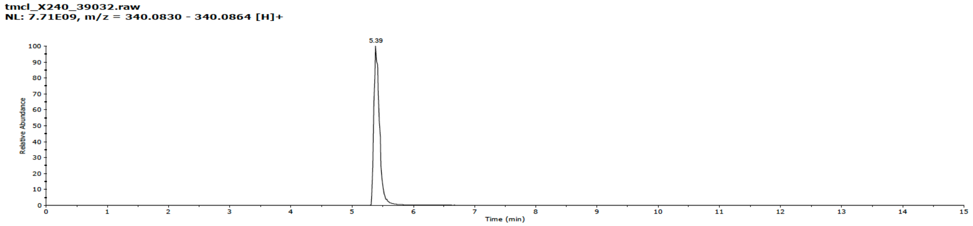

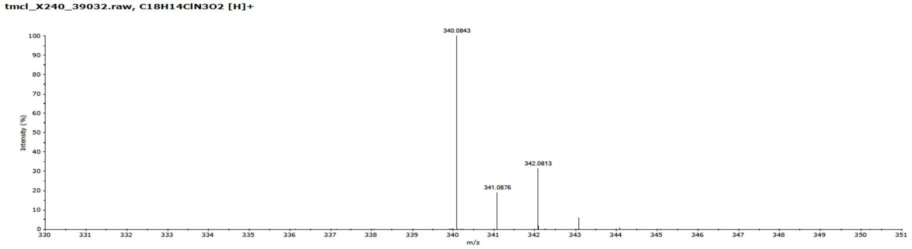

**Compound 23**

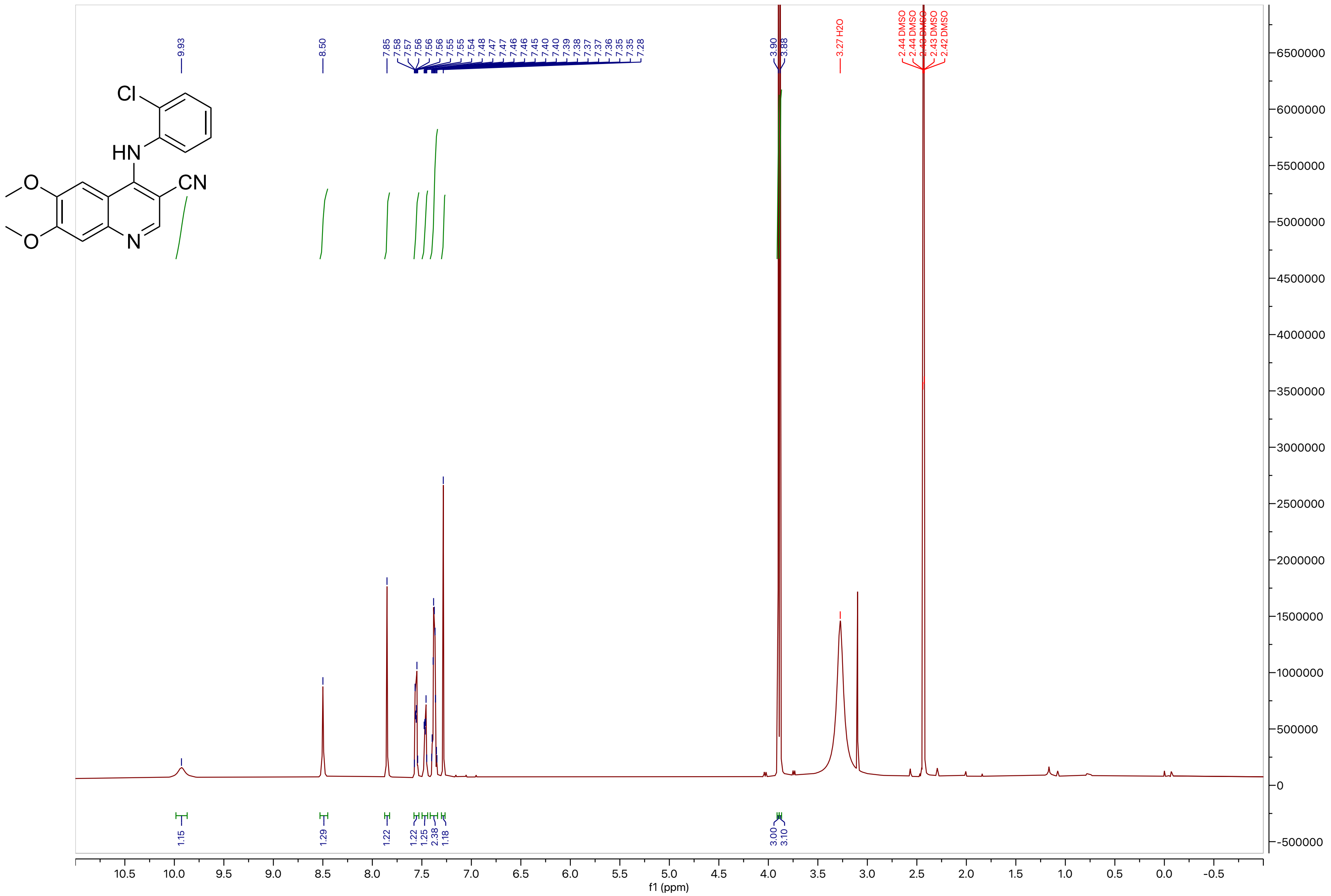

9.93
8.50
7.85
7.58
7.57
7.56
7.56
7.56
7.55
7.55
7.54
7.48
7.47
7.47
7.46
7.45
7.40
7.40
7.39
7.38
7.37
7.37
7.36
7.35
7.35
7.28
3.90
3.88
3.27 H2O
2.44 DMSO
2.44 DMSO
2.43 DMSO
2.43 DMSO
2.42 DMSO
Cl
HN
CN
O
O
N
1.15
1.29
1.22
1.22
1.25
2.38
1.18
3.00
3.10
f1 (ppm)

**Compound 23**

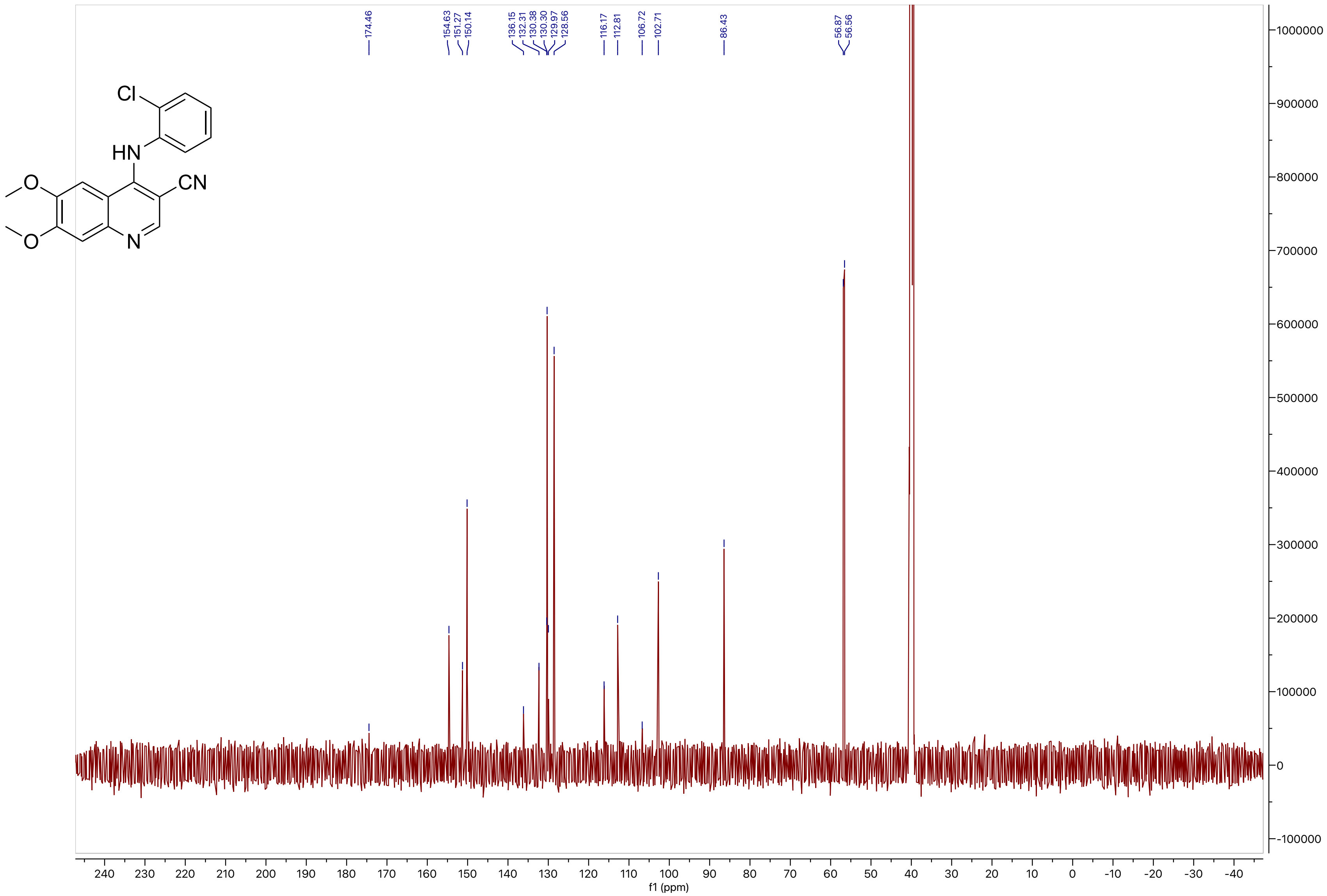
Cl
HN
CN
O
O
N
174.46
154.63
151.27
150.14
136.15
132.31
130.38
130.30
129.97
128.56
116.17
112.81
106.72
102.71
86.43
56.87
56.56
1000000
900000
800000
700000
600000
500000
400000
300000
200000
100000
0
-100000
240
230
220
210
200
190
180
170
160
150
140
130
120
110
100
90
80
70
60
50
40
30
20
10
0
-10
-20
-30
-40
f1 (ppm)

**Compound 24**

LCMS

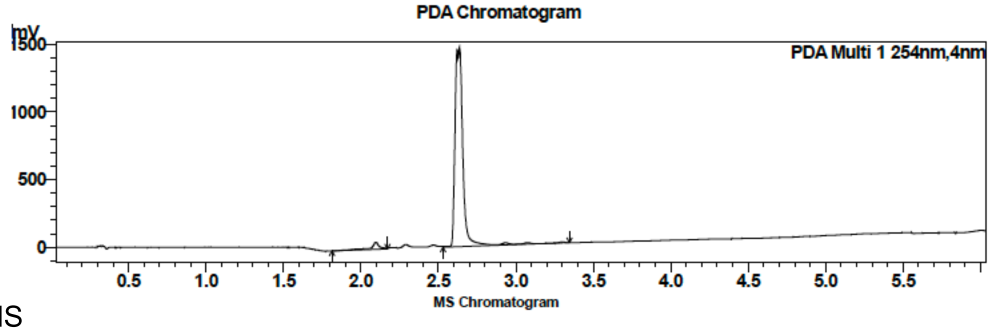

| Peak # | Ret. time | Area | Area % |
|---|---|---|---|
| 1 | 2.1 | 229555 | 4.28 |
| 2 | 2.6 | 5128433 | 95.72 |

HRMS

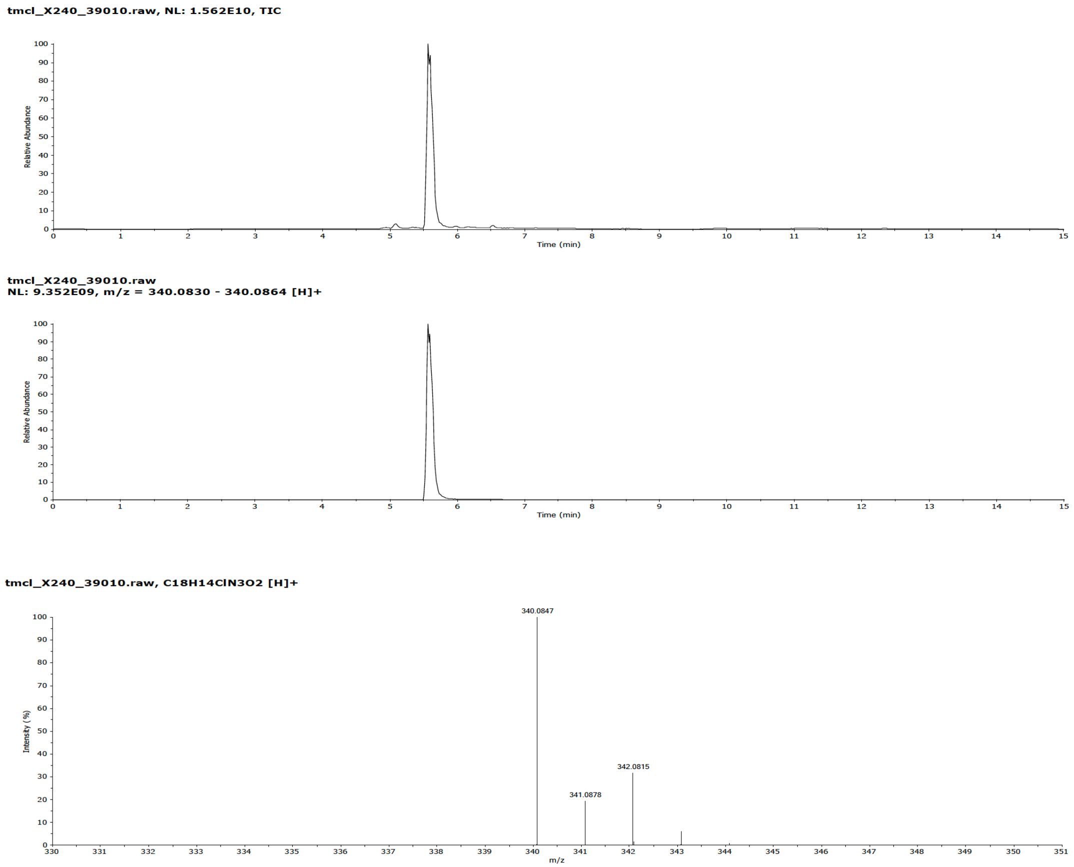

**Compound 24**

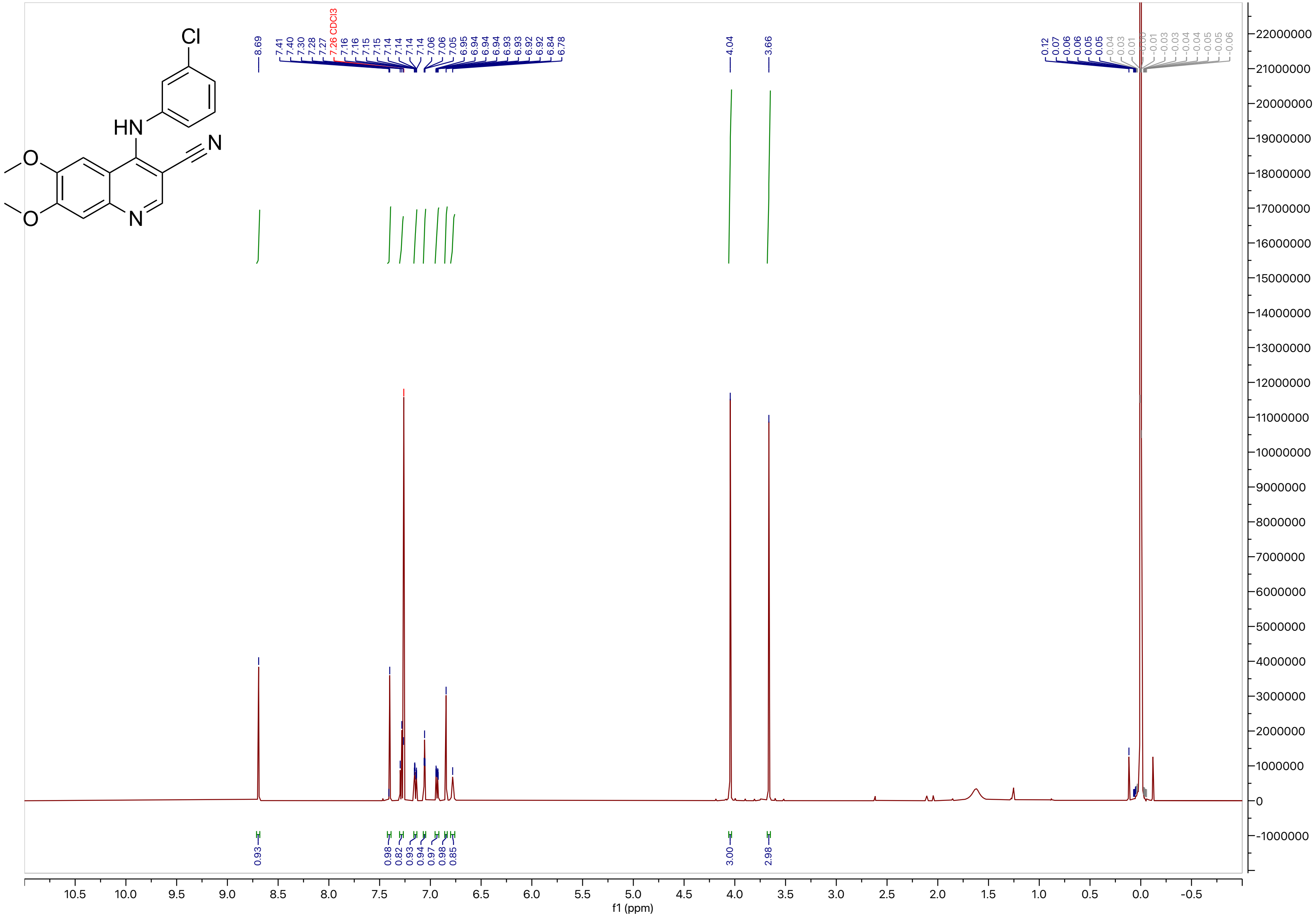

#### Compound 24

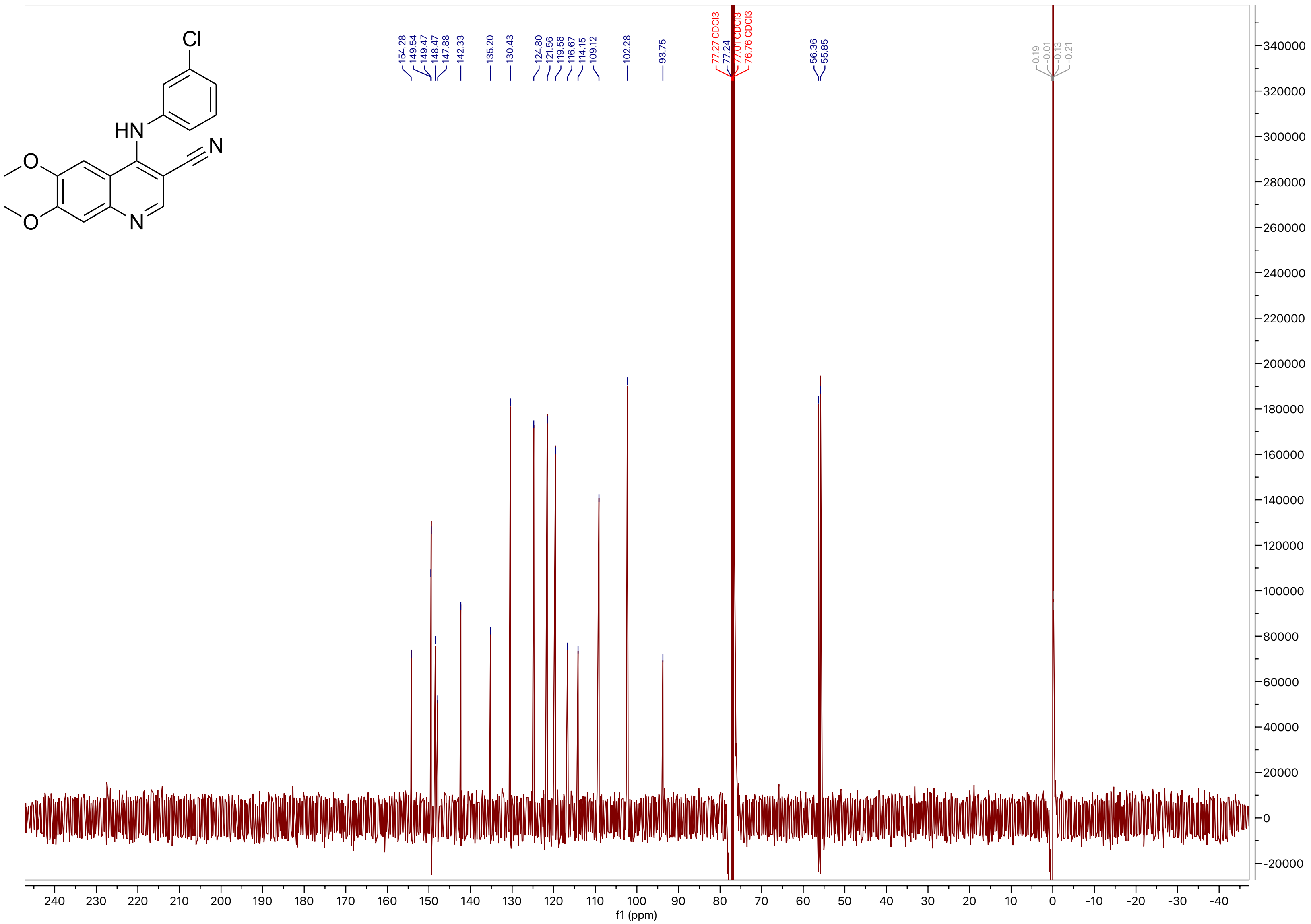
154.28
149.54
149.47
148.47
147.88
142.33
135.20
130.43
124.80
121.56
119.56
116.67
114.15
109.12
102.28
93.75
77.27 CDCl3
77.24
77.01 CDCl3
76.76 CDCl3
56.36
55.85
0.19
-0.01
-0.13
-0.21
f1 (ppm)

**Compound 25**

LCMS

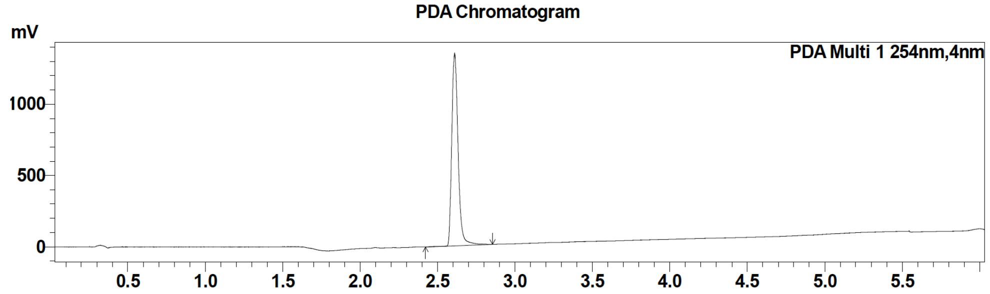

| Peak # | Ret. time | Area | Area % |
|---|---|---|---|
| 1 | 2.6 | 3845735 | 100 |

MS Chromatogram

HRMS

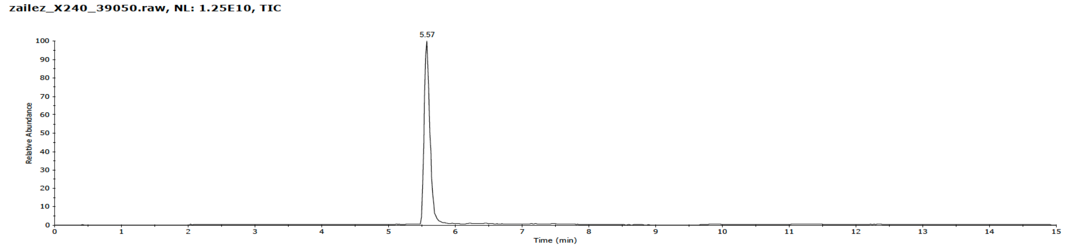

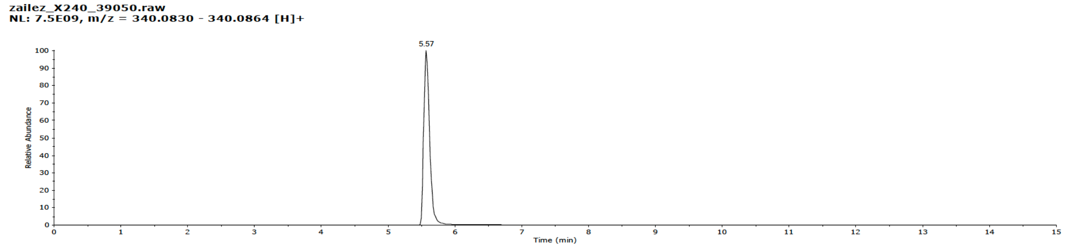

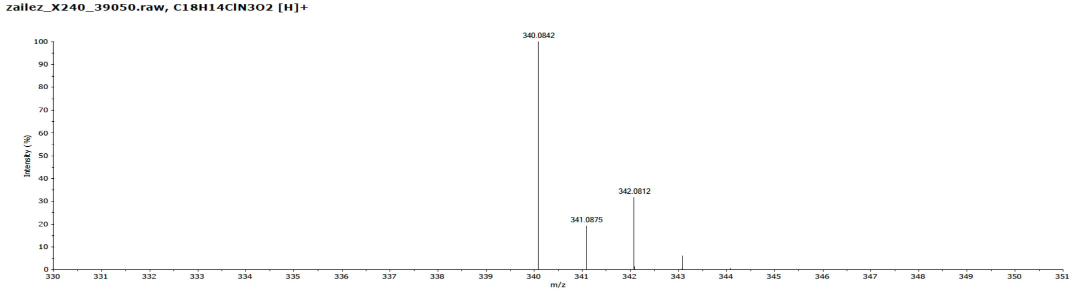

#### Compound 25

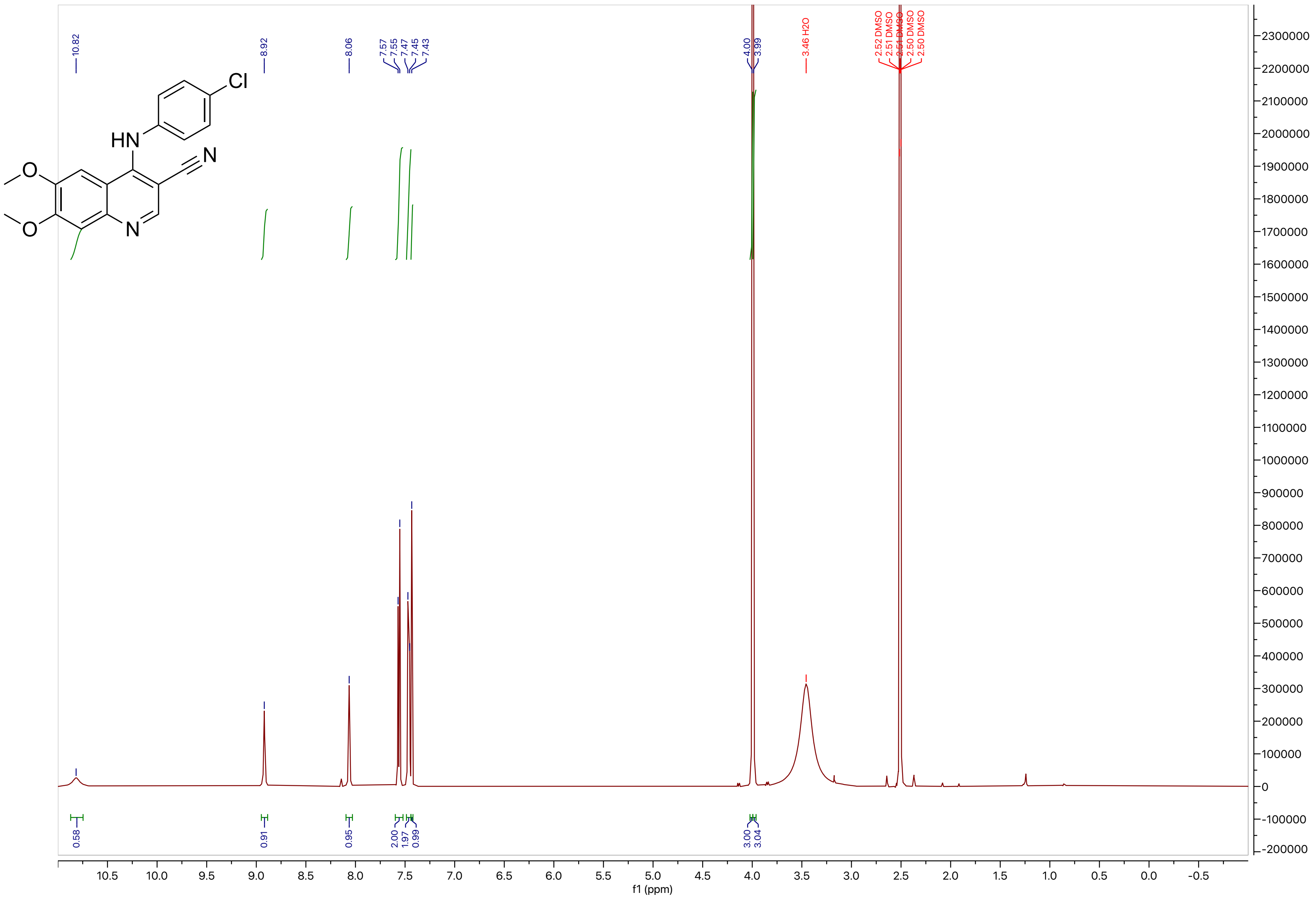
10.82
8.92
8.06
7.57
7.55
7.47
7.45
7.43
4.00
3.99
3.46 H2O
2.52 DMSO
2.51 DMSO
2.51 DMSO
2.50 DMSO
2.50 DMSO
0.58
0.91
0.95
2.00
1.97
0.99
3.00
3.04
f1 (ppm)
Cl
HN
N
O
O
N

### Compound 25

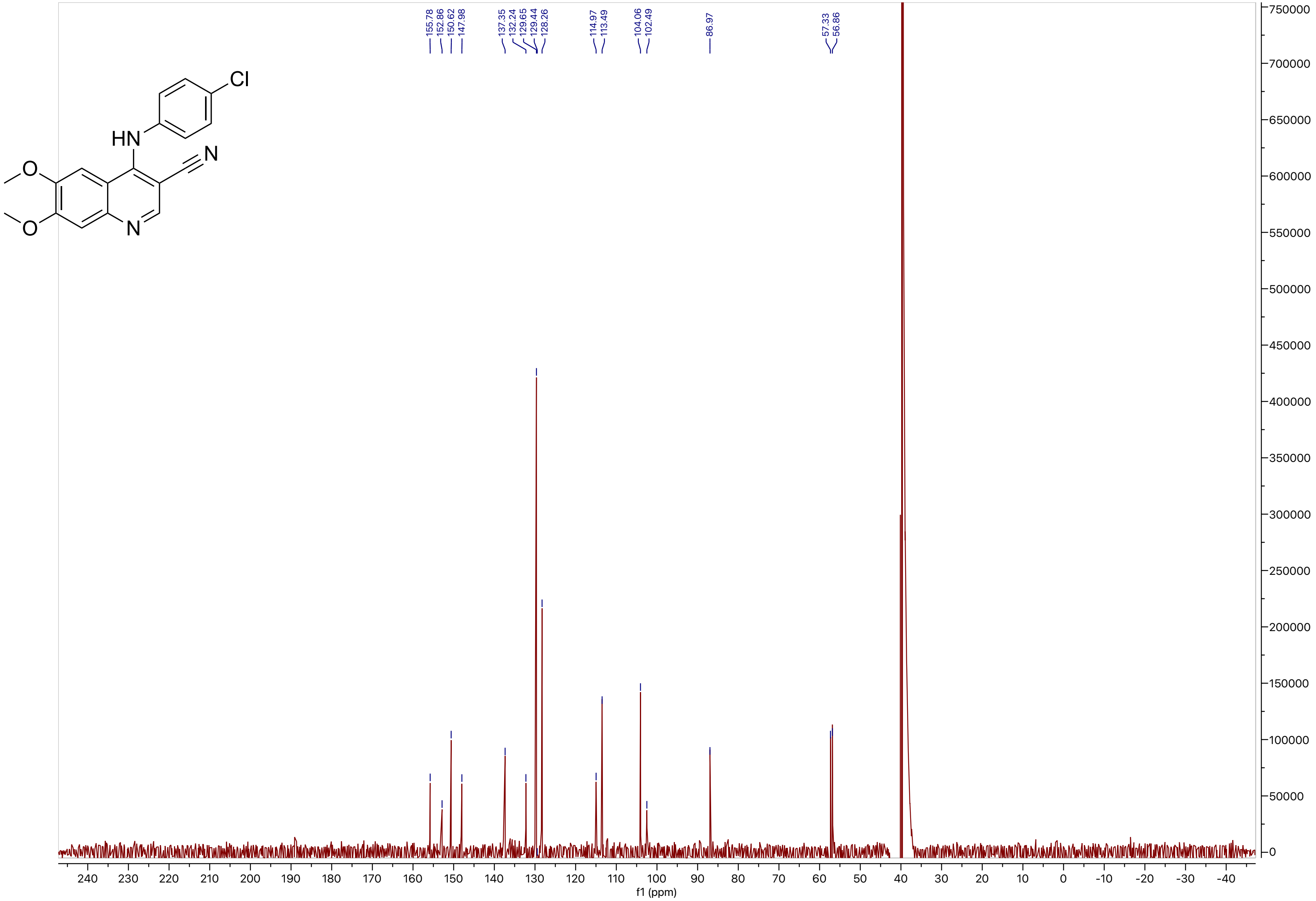
HN
Cl
N
O
O
N
155.78
152.86
150.62
147.98
137.35
132.24
129.65
129.44
128.26
114.97
113.49
104.06
102.49
86.97
57.33
56.86
f1 (ppm)

**Compound 26**

LCMS

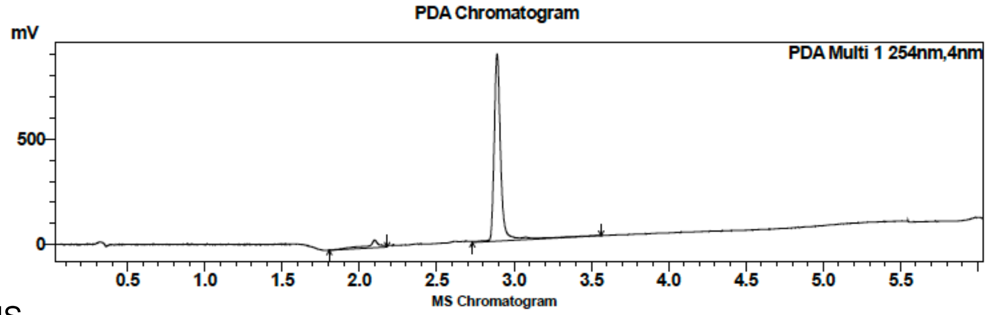

| Peak # | Ret. time | Area | Area % |
|---|---|---|---|
| 1 | 2.1 | 222394 | 7.77 |
| 2 | 2.4 | 2640708 | 92.23 |

HRMS

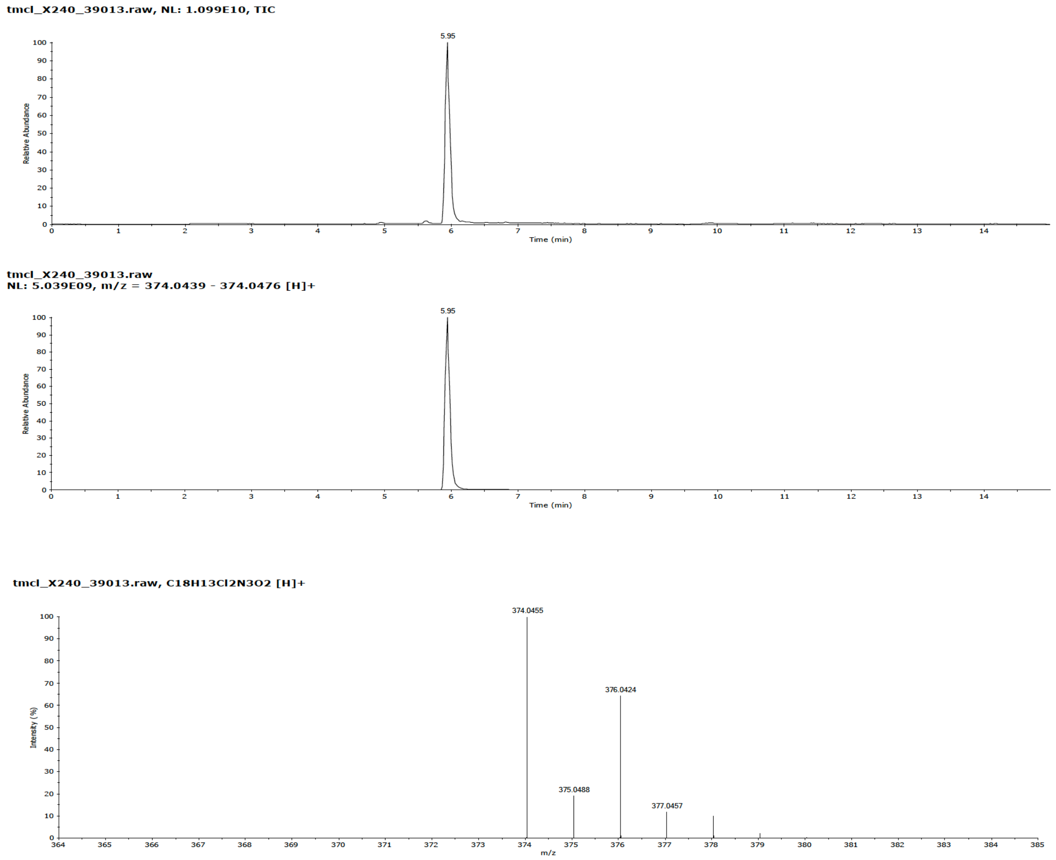

**Compound 26**

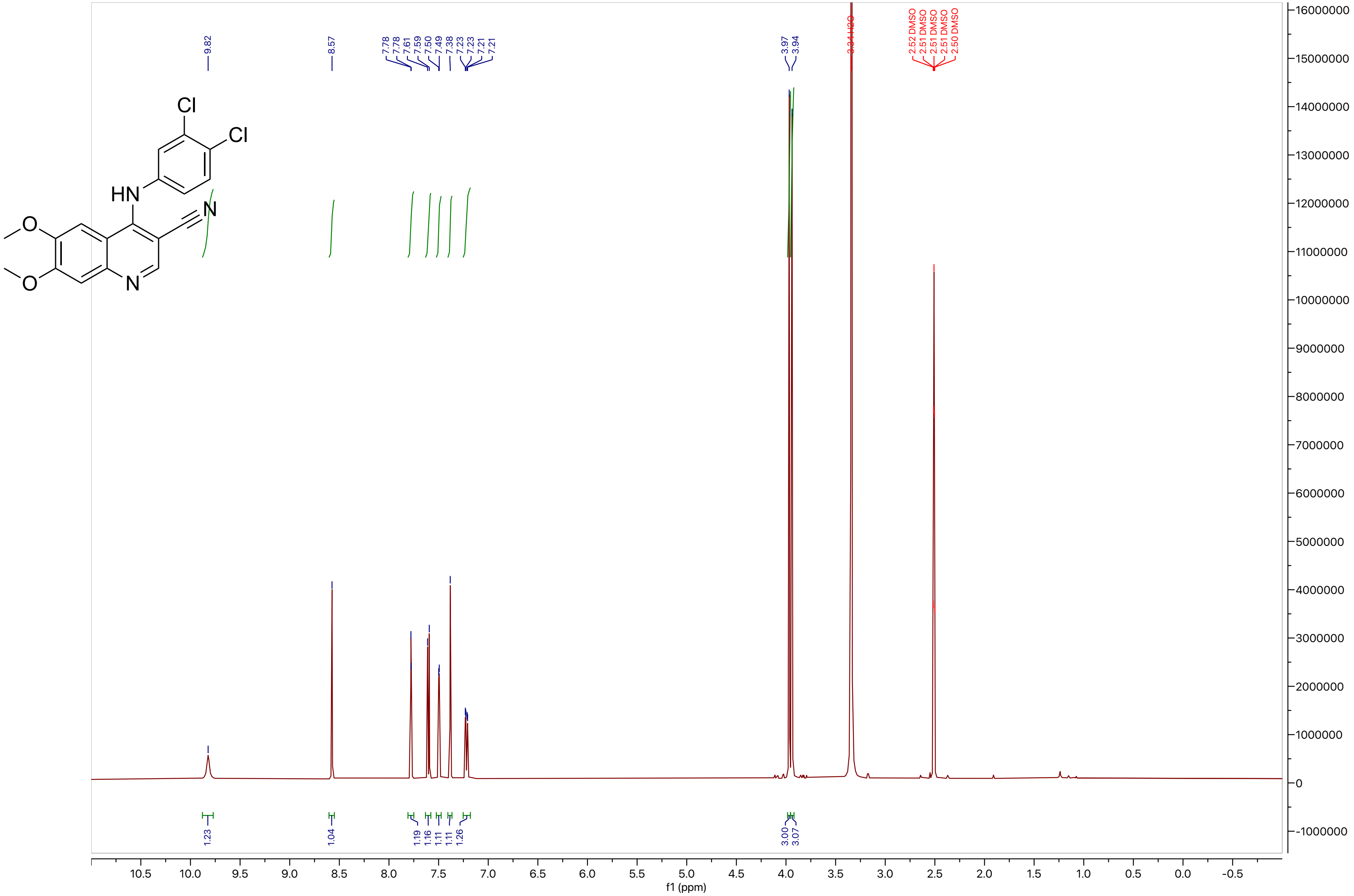

9.82
8.57
7.78
7.78
7.61
7.59
7.50
7.49
7.38
7.23
7.23
7.21
7.21
3.97
3.94
3.34 H2O
2.52 DMSO
2.51 DMSO
2.51 DMSO
2.51 DMSO
2.50 DMSO
Cl
Cl
HN
N
O
O
N
1.23
1.04
1.19
1.16
1.11
1.11
1.26
3.00
3.07
f1 (ppm)

#### Compound 26

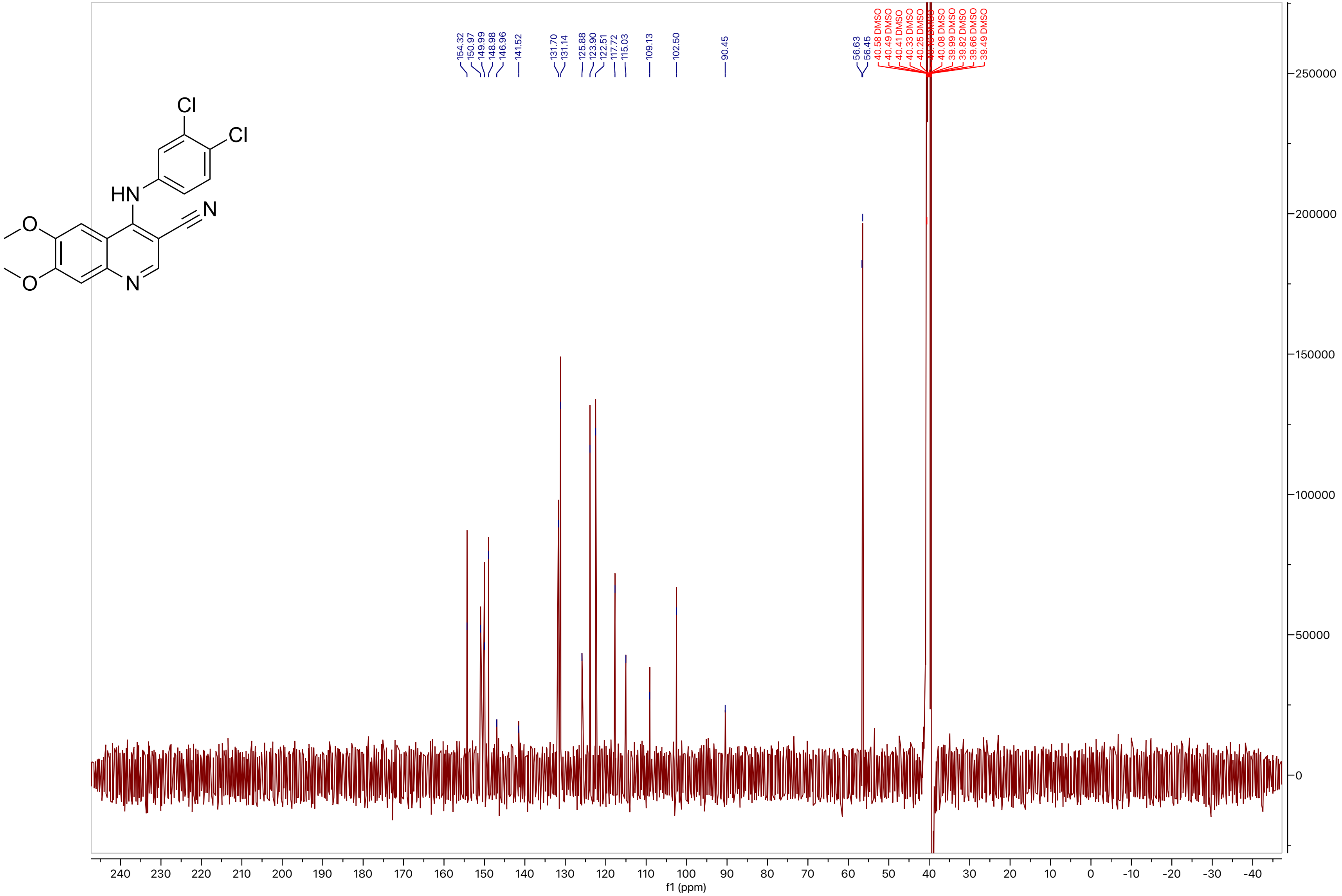

Cl
Cl
HN
N
O
O
N
154.32
150.97
149.99
148.98
146.96
141.52
131.70
131.14
125.88
123.90
122.51
117.72
115.03
109.13
102.50
90.45
56.63
56.45
40.58 DMSO
40.49 DMSO
40.41 DMSO
40.33 DMSO
40.25 DMSO
40.08 DMSO
39.99 DMSO
39.82 DMSO
39.66 DMSO
39.49 DMSO
250000
200000
150000
100000
50000
0
240
230
220
210
200
190
180
170
160
150
140
130
120
110
100
90
80
70
60
50
40
30
20
10
0
-10
-20
-30
-40
f1 (ppm)

**Compound 27**

LCMS

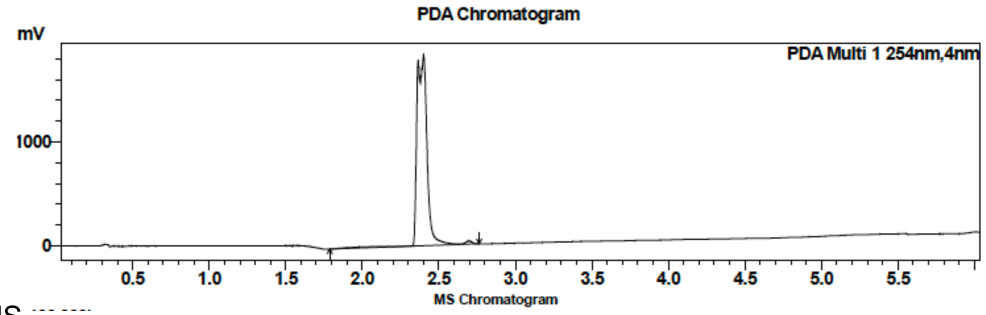

| Peak # | Ret. time | Area | Area % |
|---|---|---|---|
| 1 | 2.4 | 8450144 | 100.00 |

HRMS

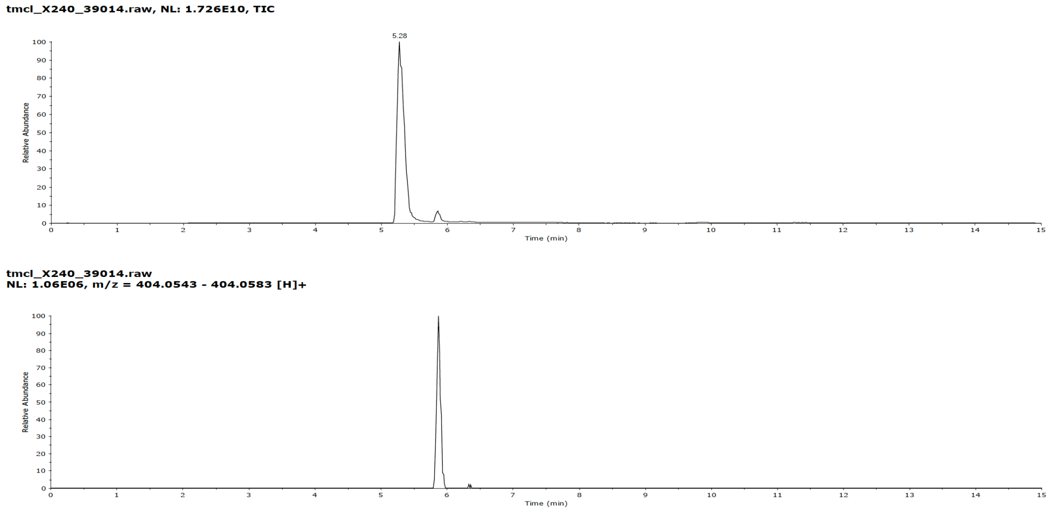

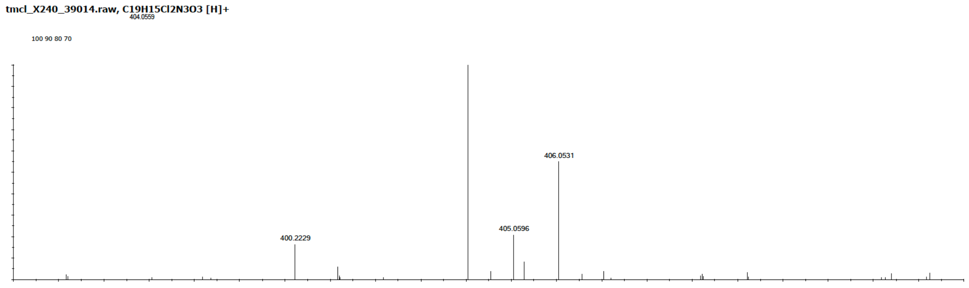

**Compound 27**

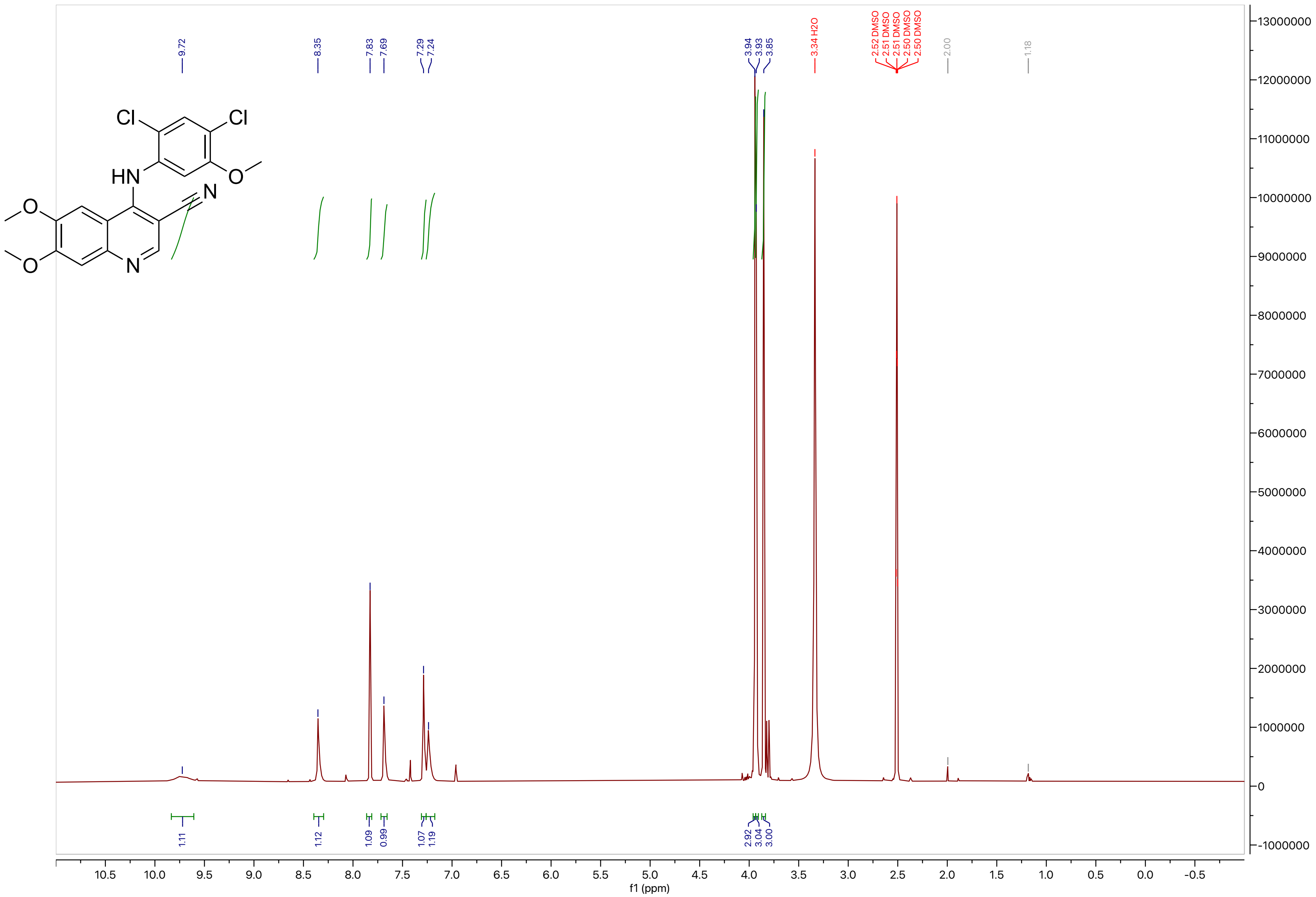
9.72
8.35
7.83
7.69
7.29
7.24
3.94
3.93
3.85
3.34 H2O
2.52 DMSO
2.51 DMSO
2.51 DMSO
2.50 DMSO
2.50 DMSO
2.00
1.18
1.11
1.12
1.09
0.99
1.07
1.19
2.92
3.04
3.00
f1 (ppm)

**Compound 27**

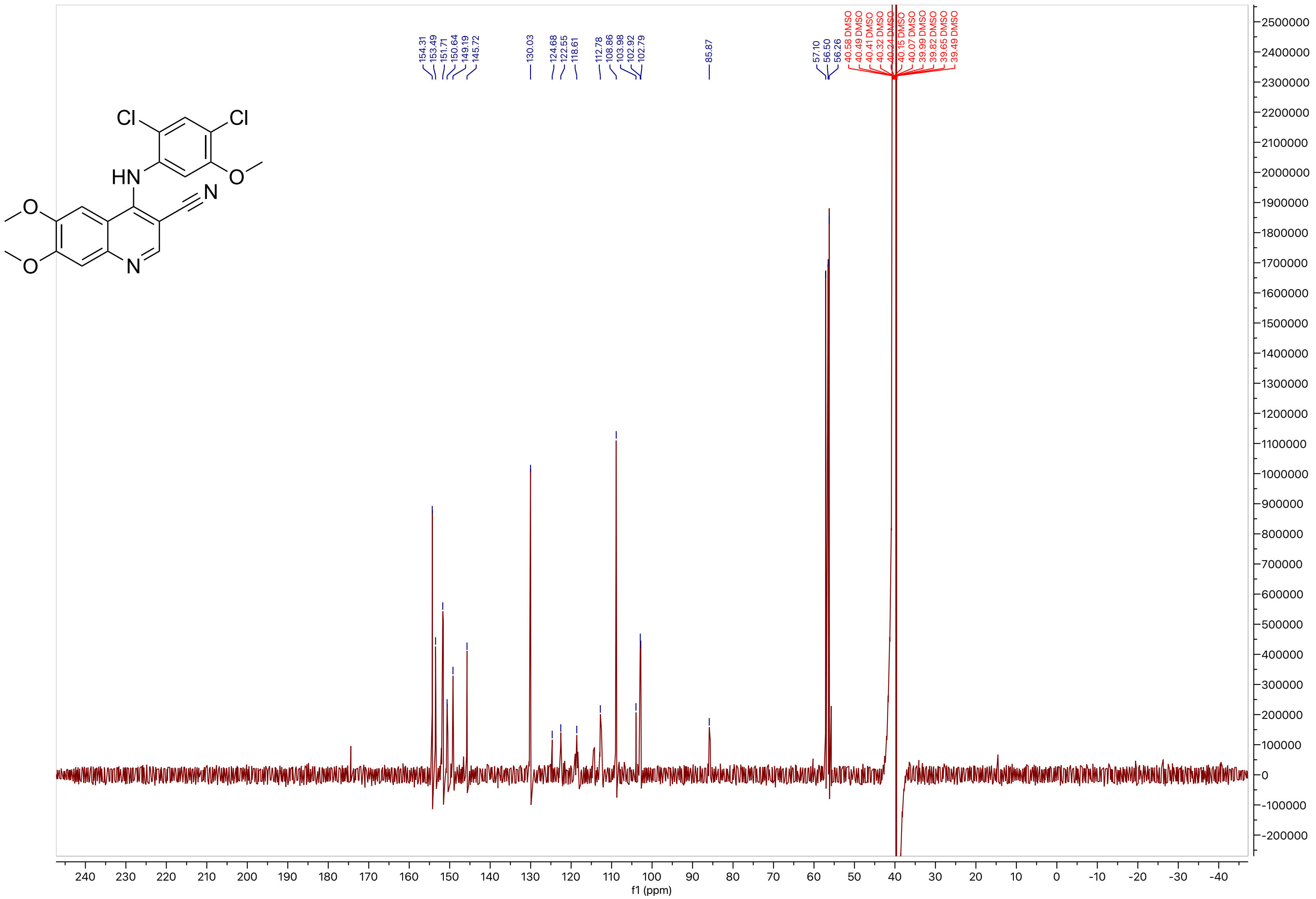
154.31
153.49
151.71
150.64
149.19
145.72
130.03
124.68
122.55
118.61
112.78
108.86
103.98
102.92
102.79
85.87
57.10
56.50
56.26
40.58 DMSO
40.49 DMSO
40.41 DMSO
40.32 DMSO
40.24 DMSO
40.15 DMSO
40.07 DMSO
39.99 DMSO
39.82 DMSO
39.65 DMSO
39.49 DMSO
f1 (ppm)

**Compound 28**

LCMS

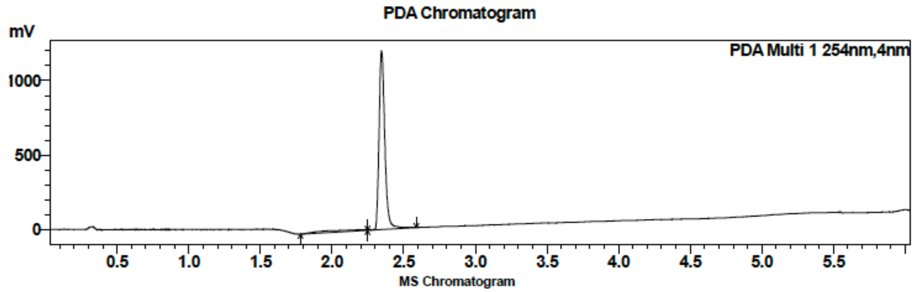

| Peak # | Ret. time | Area | Area % |
|---|---|---|---|
| 1 | 2.3 | 3356635 | 100.00 |

HRMS

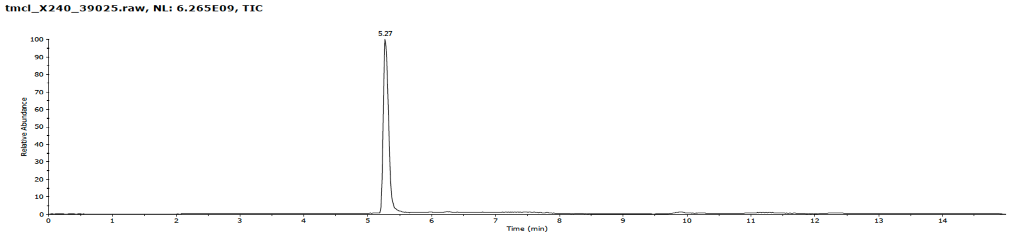

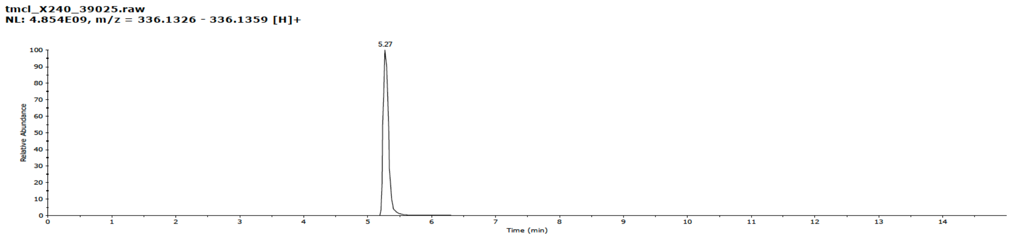

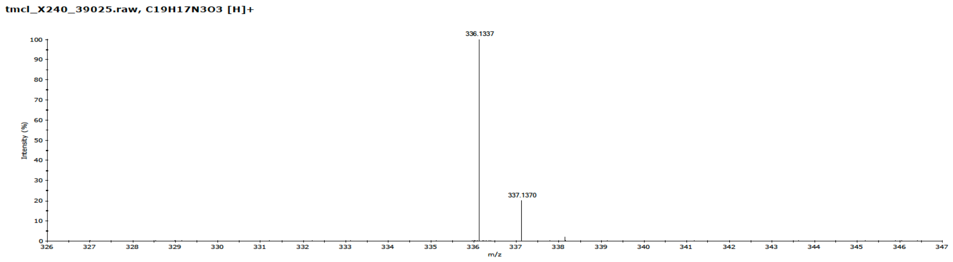

**Compound 28**

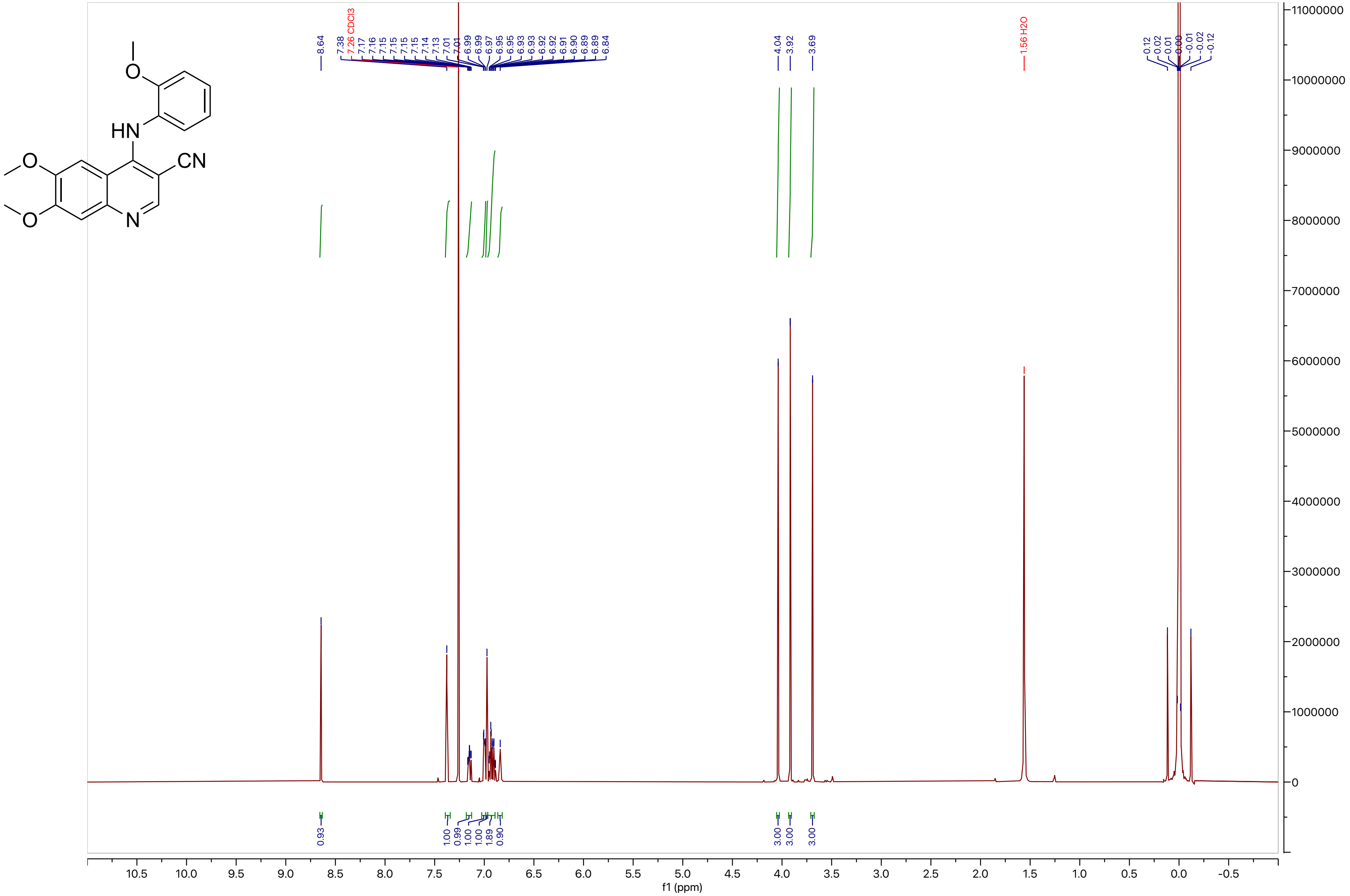

**Compound 28**

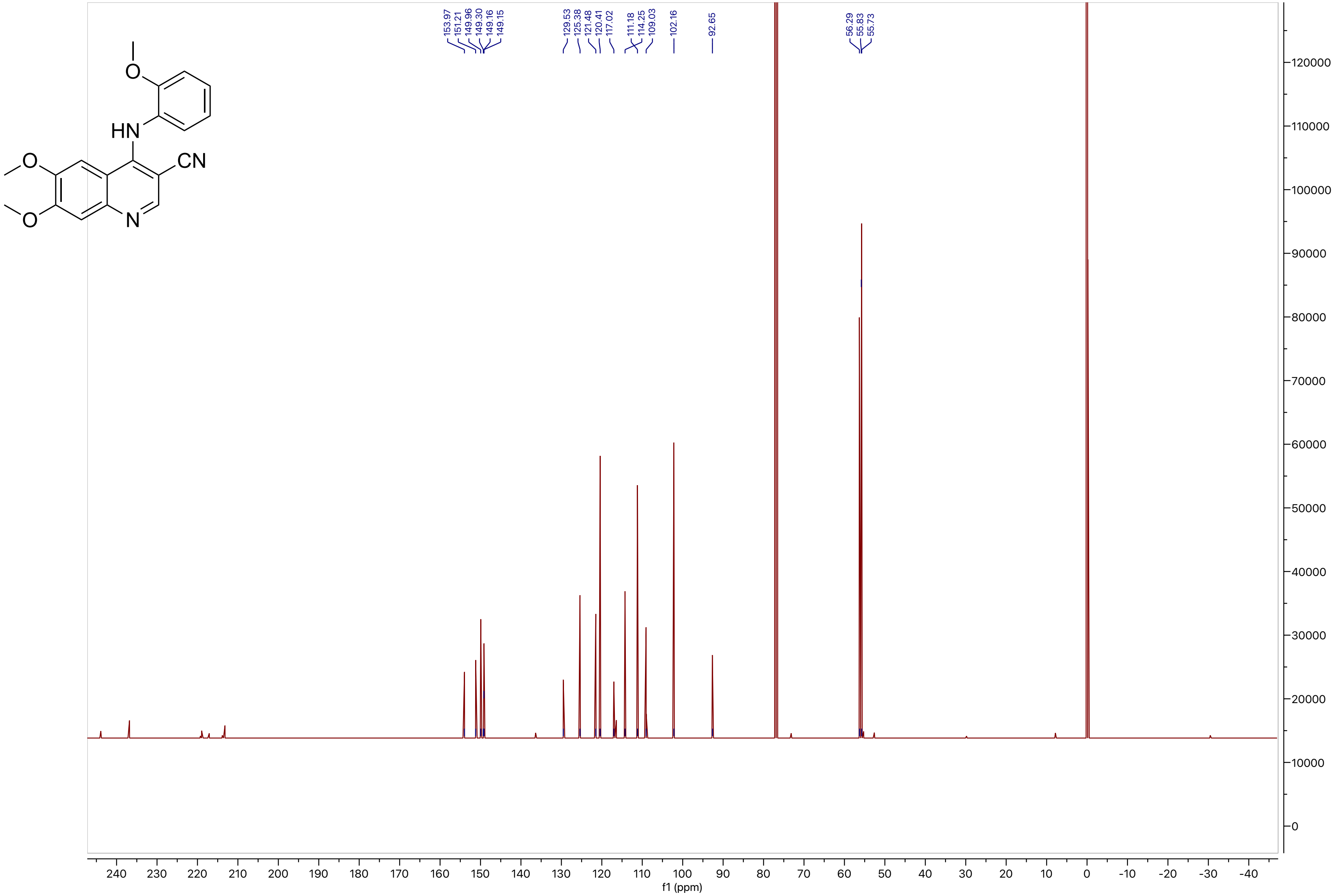
153.97
151.21
149.96
149.30
149.16
149.15
129.53
125.38
121.48
120.41
117.02
111.18
114.25
109.03
102.16
92.65
56.29
55.83
55.73
f1 (ppm)

**Compound 29**

LCMS

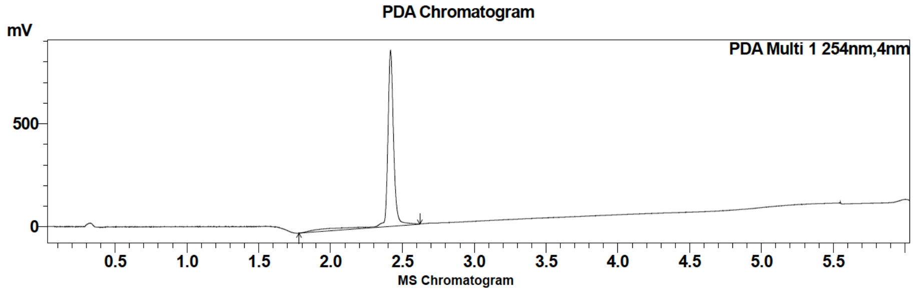

| Peak # | Ret. time | Area | Area % |
|---|---|---|---|
| 1 | 2.4 | 2610080 | 100.00 |

HRMS

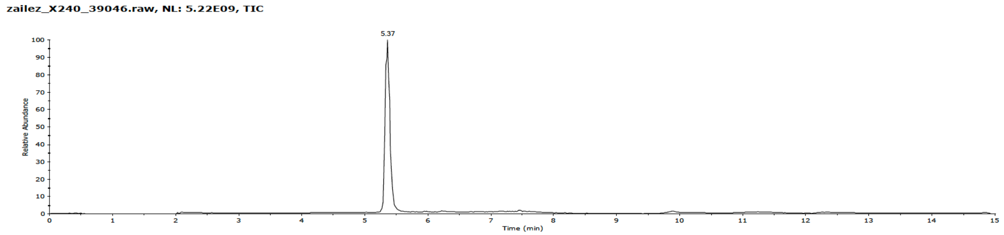

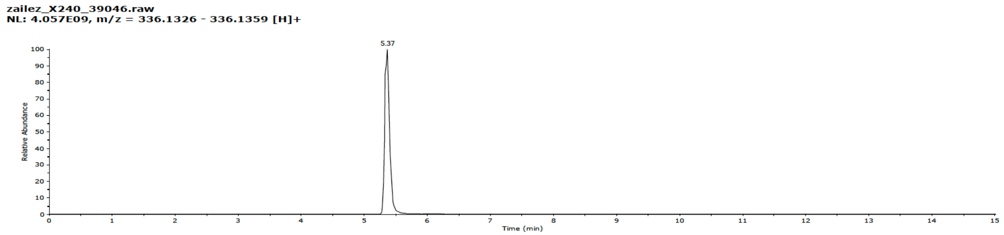

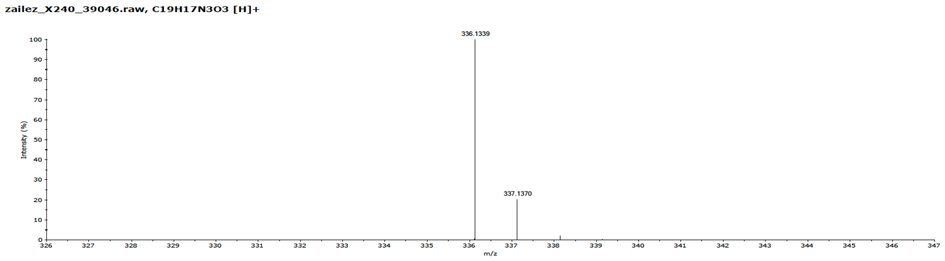

#### Compound 29

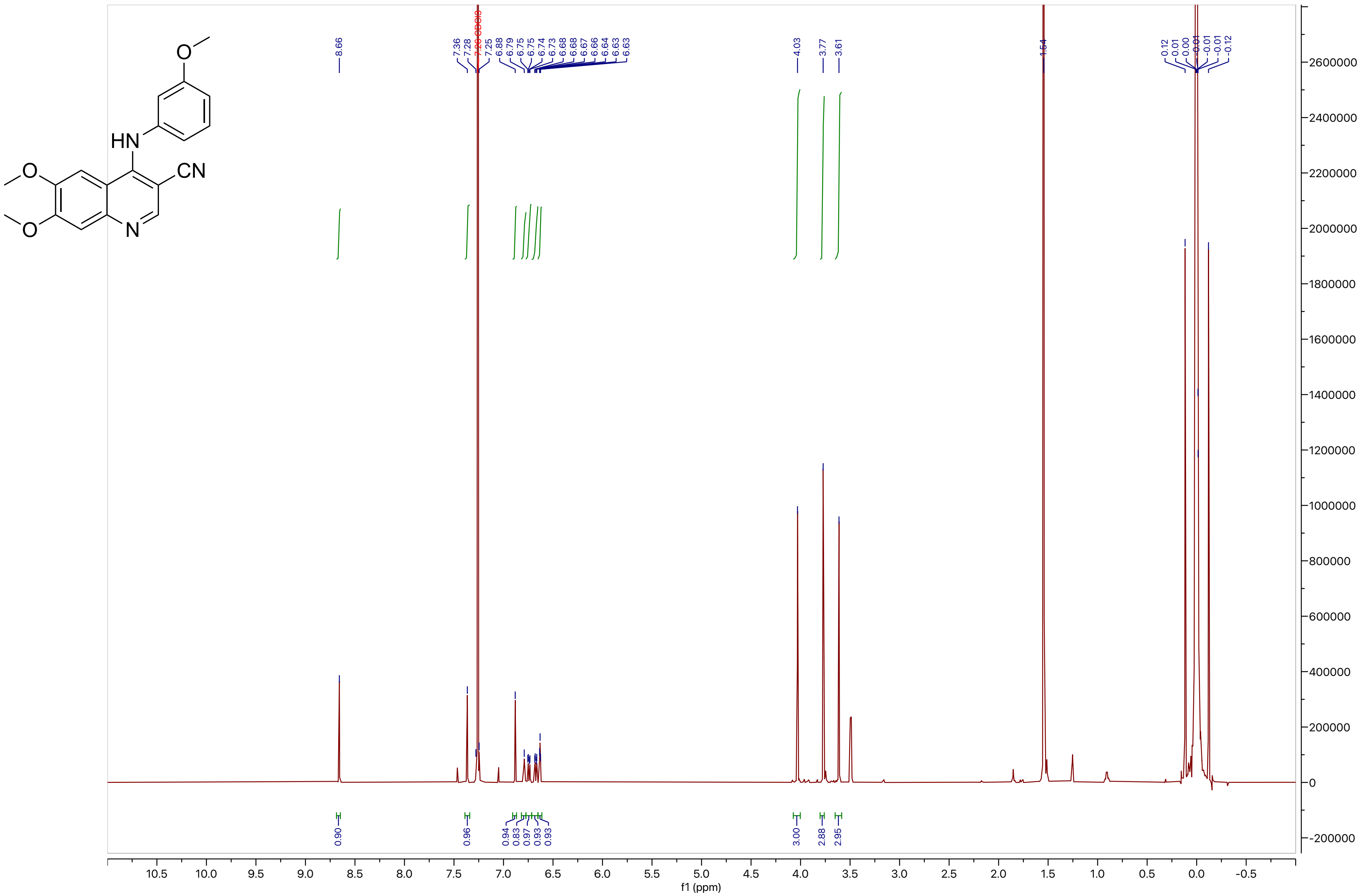
O
HN
CN
O
O
N
8.66
7.36
7.28
7.25
6.88
6.79
6.75
6.75
6.74
6.73
6.68
6.68
6.67
6.66
6.64
6.63
6.63
4.03
3.77
3.61
1.54
0.12
0.01
0.00
-0.01
-0.01
-0.01
-0.12
0.90
0.96
0.94
0.83
0.97
0.93
0.93
0.93
3.00
2.88
2.95
f1 (ppm)
10.5
10.0
9.5
9.0
8.5
8.0
7.5
7.0
6.5
6.0
5.5
5.0
4.5
4.0
3.5
3.0
2.5
2.0
1.5
1.0
0.5
0.0
-0.5
2600000
2400000
2200000
2000000
1800000
1600000
1400000
1200000
1000000
800000
600000
400000
200000
0
-200000

**Compound 29**

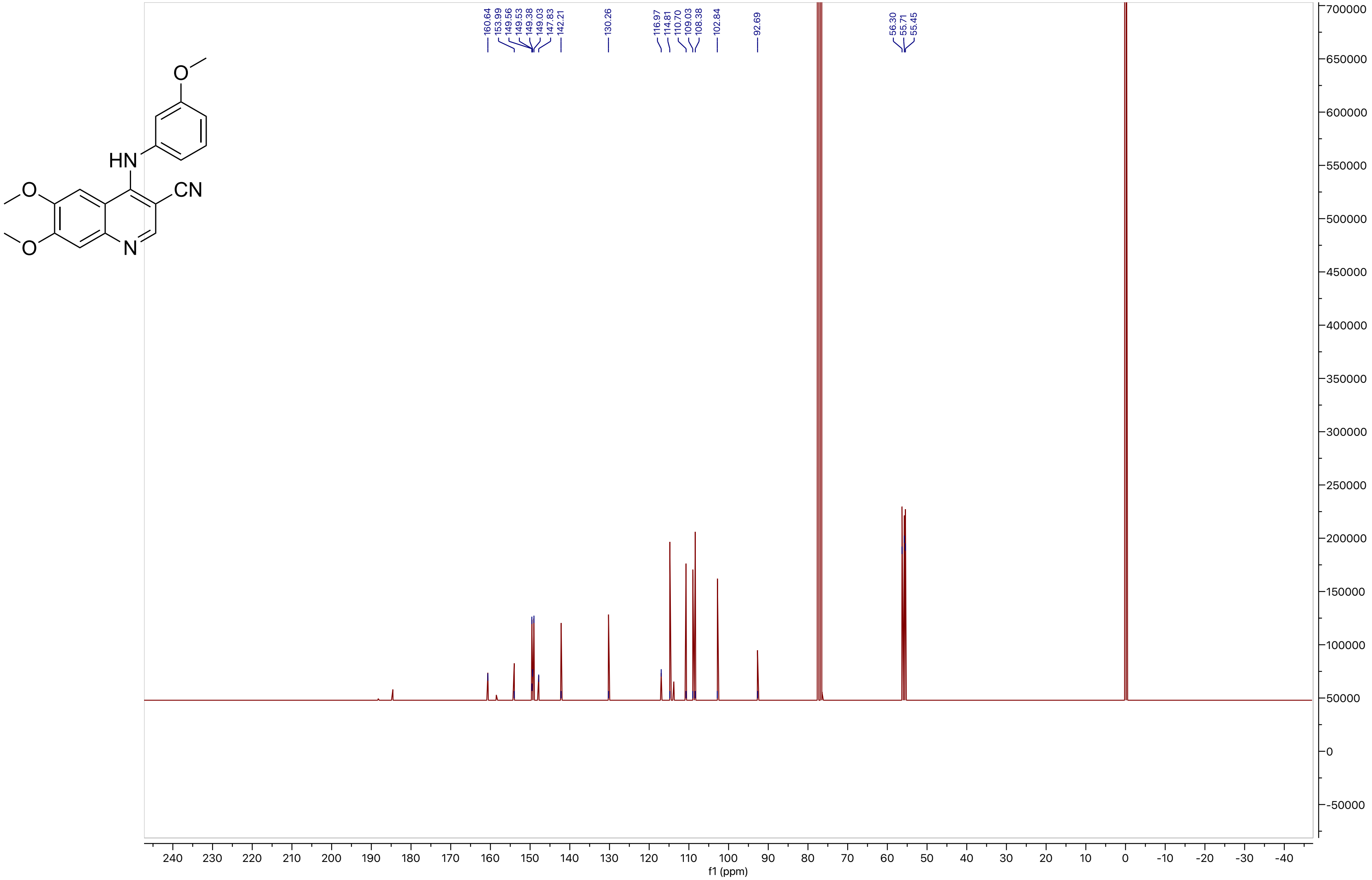

160.64
153.99
149.56
149.53
149.38
149.03
147.83
142.21
130.26
116.97
114.81
110.70
109.03
108.38
102.84
92.69
56.30
55.71
55.45
HN
CN
O
N
f1 (ppm)

**Compound 30**

LCMS

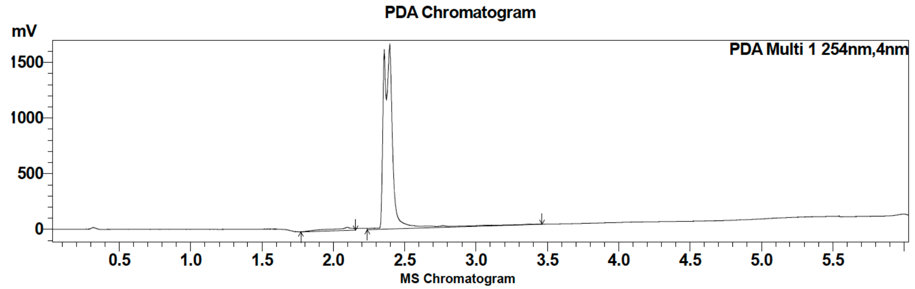

| Peak # | Ret. time | Area | Area % |
|---|---|---|---|
| 1 | 2.1 | 303815 | 4.05 |
| 2 | 2.4 | 7205709 | 95.95 |

HRMS

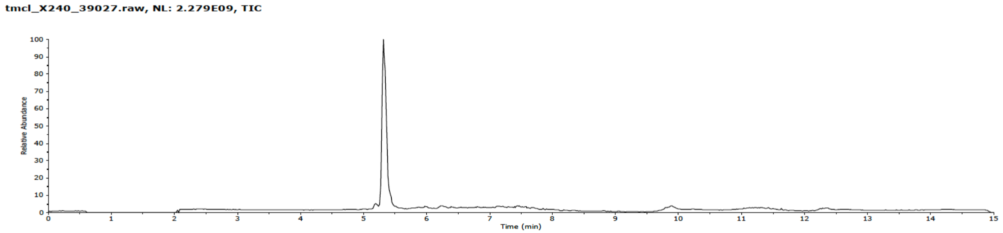

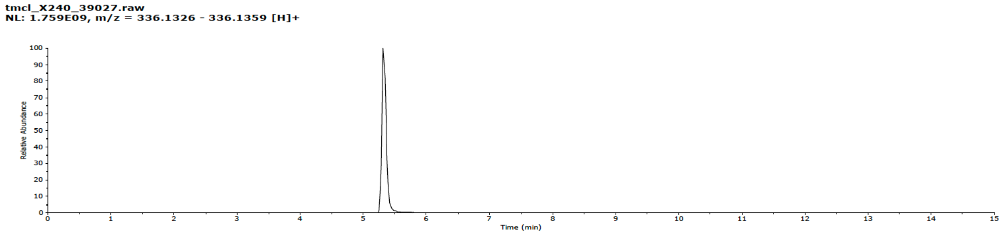

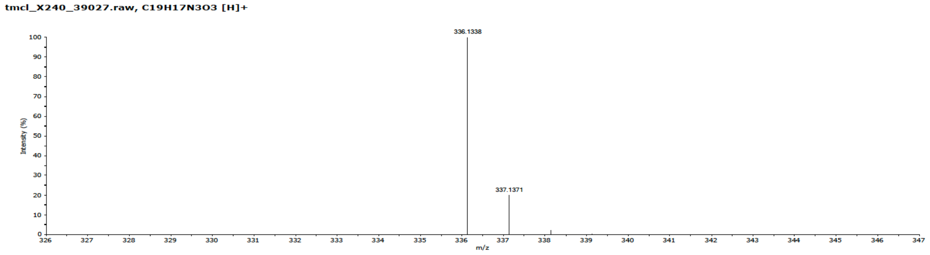

**Compound 30**

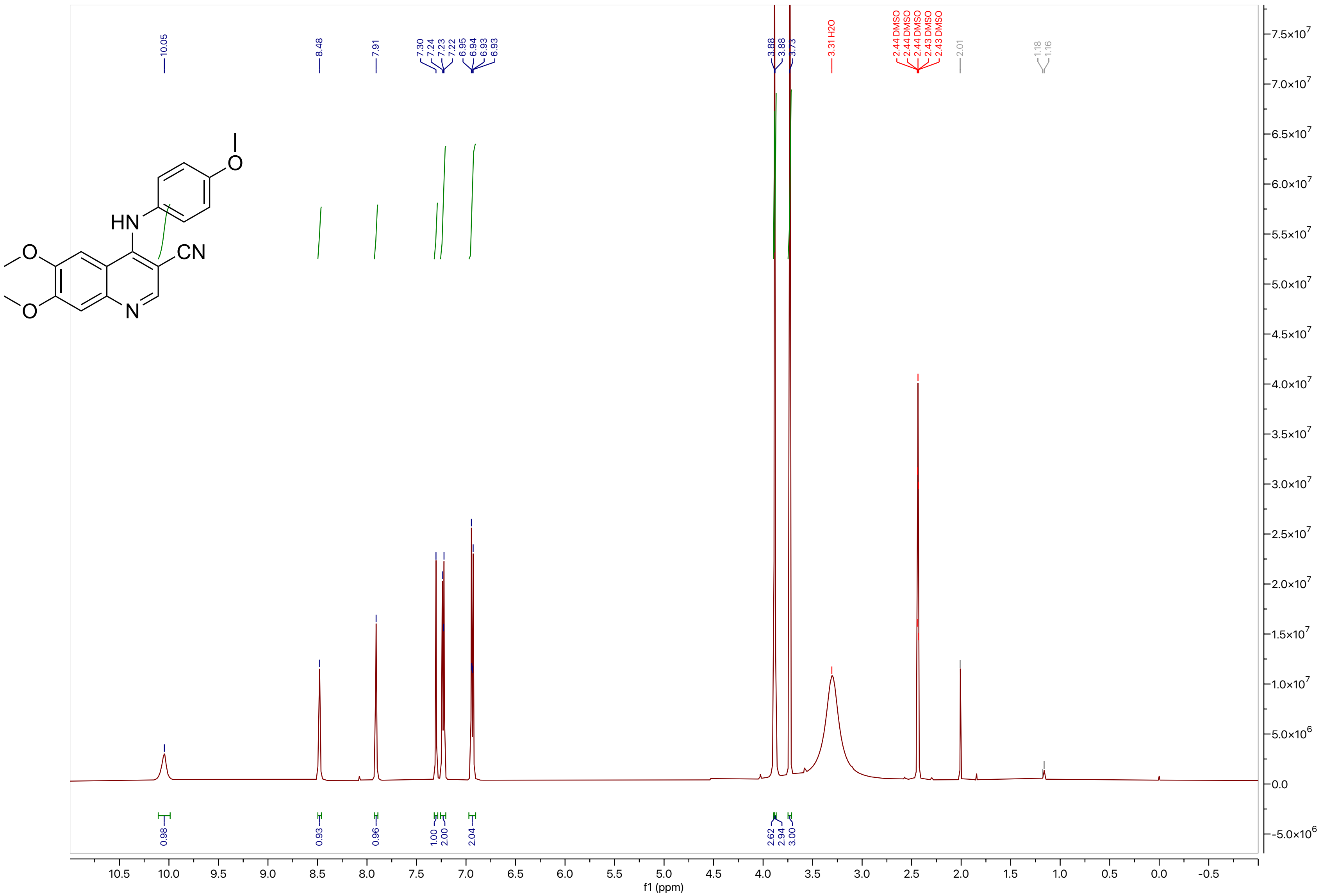
HN
CN
N
O
10.05
8.48
7.91
7.30
7.24
7.23
7.22
6.95
6.94
6.93
6.93
3.88
3.88
3.73
3.31 H2O
2.44 DMSO
2.44 DMSO
2.44 DMSO
2.43 DMSO
2.43 DMSO
2.01
1.18
1.16
0.98
0.93
0.96
1.00
2.00
2.04
2.62
2.94
3.00
f1 (ppm)

**Compound 30**

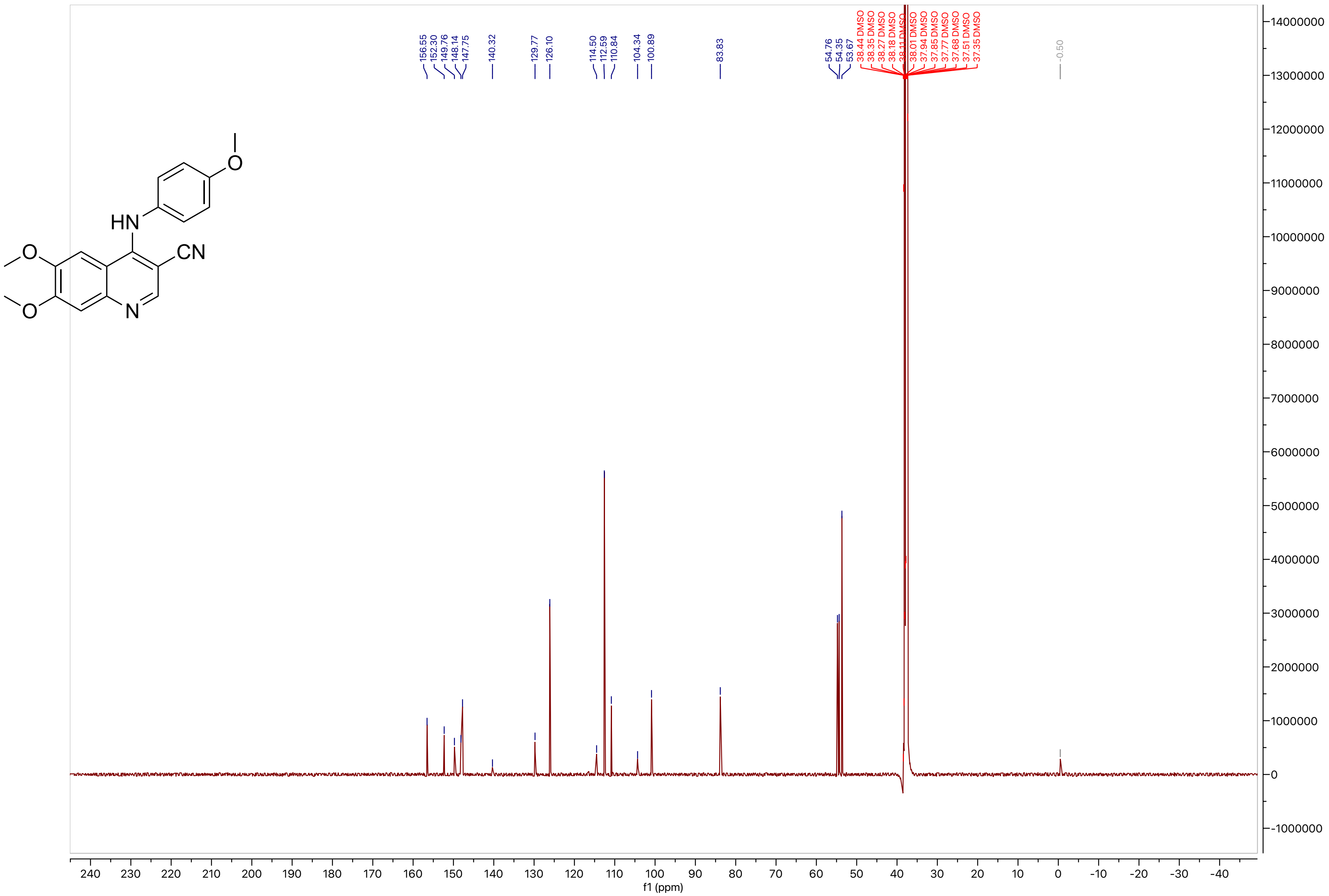

156.55
152.30
149.76
148.14
147.75
140.32
129.77
126.10
114.50
112.59
110.84
104.34
100.89
83.83
54.76
54.35
53.67
38.44 DMSO
38.35 DMSO
38.27 DMSO
38.18 DMSO
38.11 DMSO
38.01 DMSO
37.94 DMSO
37.85 DMSO
37.77 DMSO
37.68 DMSO
37.51 DMSO
37.35 DMSO
-0.50
f1 (ppm)

**Compound 31**

LCMS

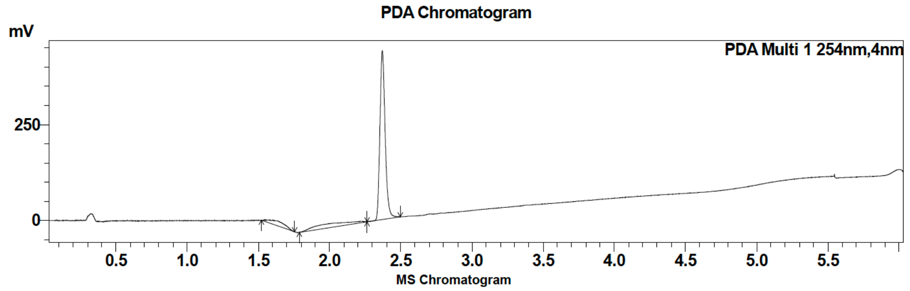

| Peak # | Ret. time | Area | Area % |
|---|---|---|---|
| 1 | 2.1 | 1125211 | 100.00 |

HRMS

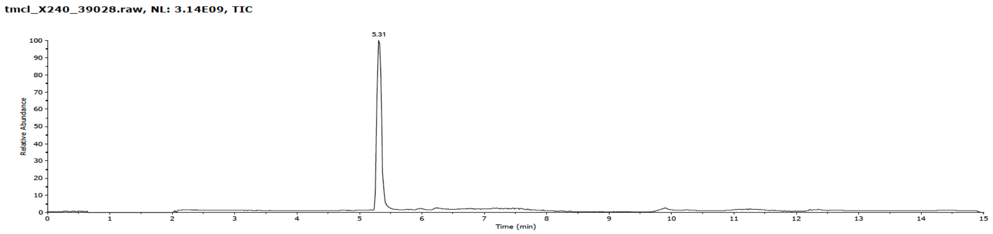

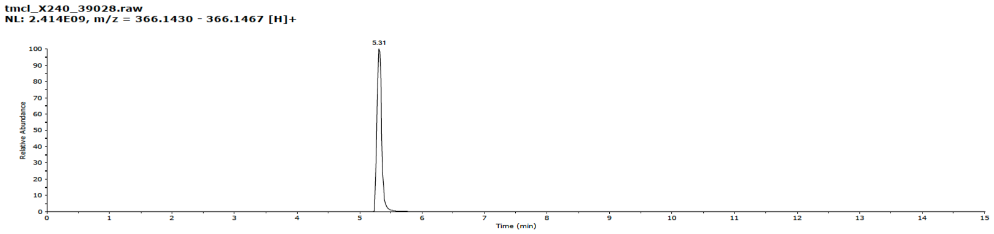

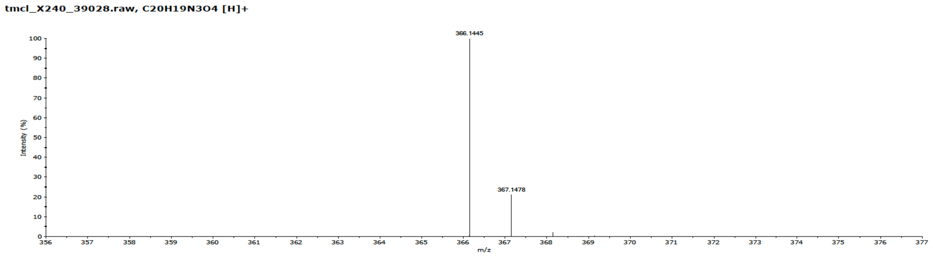

**Compound 31**

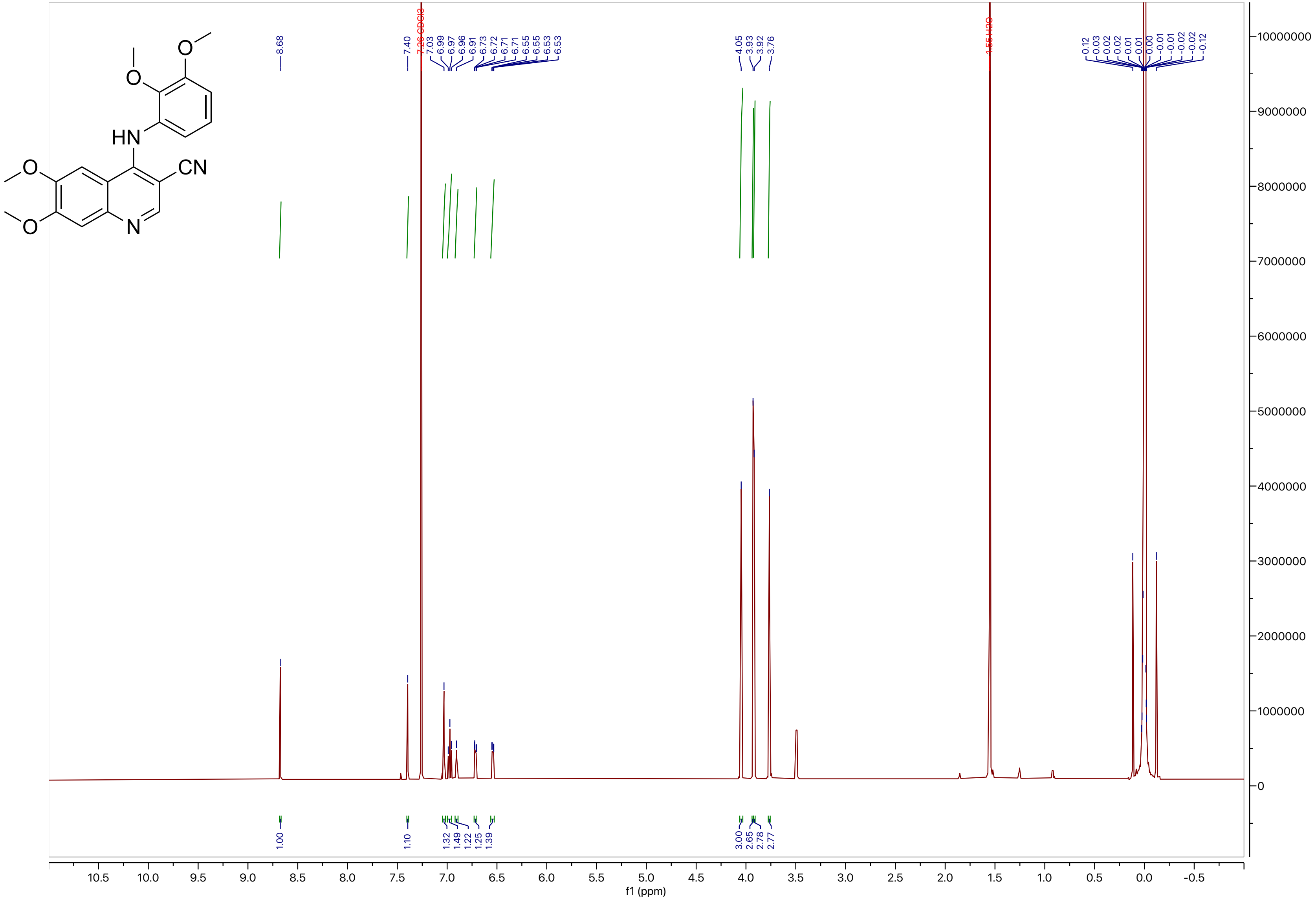

HN
CN
N
O
8.68
7.40
7.26 CDCl3
1.56 H2O
4.05
3.93
3.92
3.76
1.00
1.10
3.00
2.65
2.78
2.77
f1 (ppm)

**Compound 31**

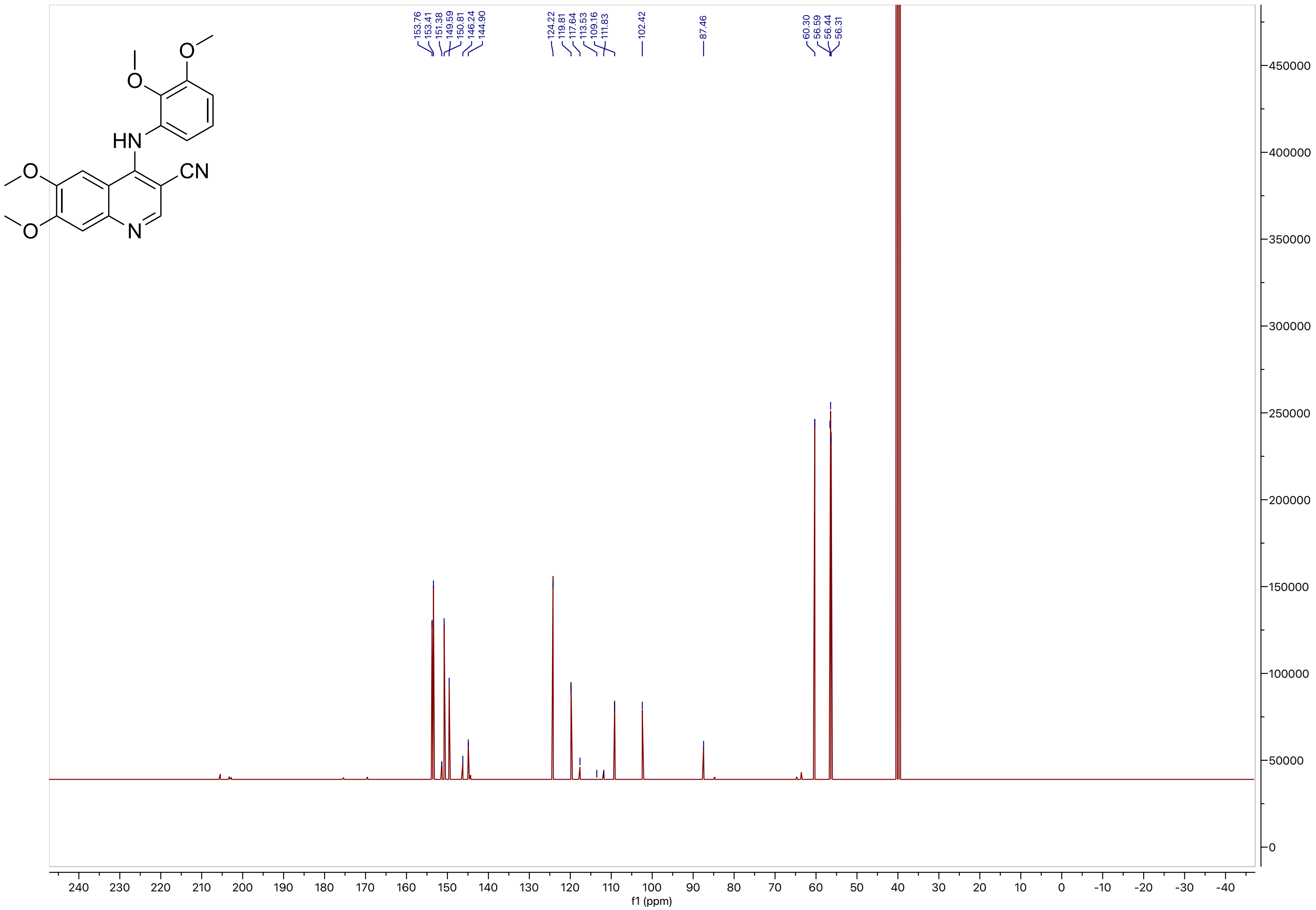
HN
CN
N
O
153.76
153.41
151.38
149.59
150.81
146.24
144.90
124.22
119.81
117.64
113.53
109.16
111.83
102.42
87.46
60.30
56.59
56.44
56.31
f1 (ppm)

**Compound 32**

LCMS

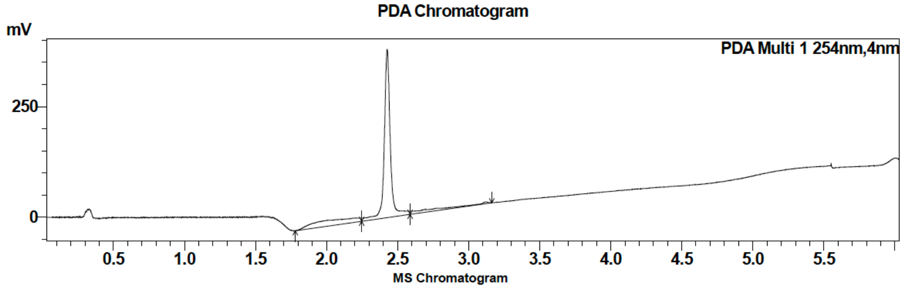

| Peak # | Ret. time | Area | Area % |
|---|---|---|---|
| 1 | 2.4 | 1136231 | 98.46 |
| 2 | 3.1 | 17824 | 1.54 |

HRMS

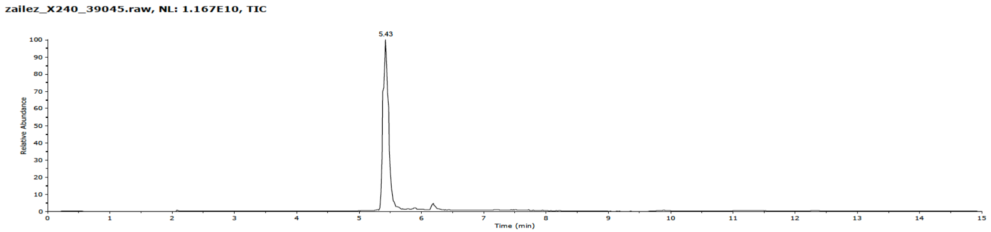

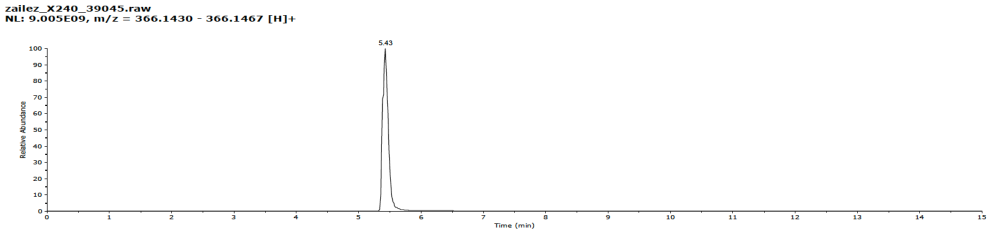

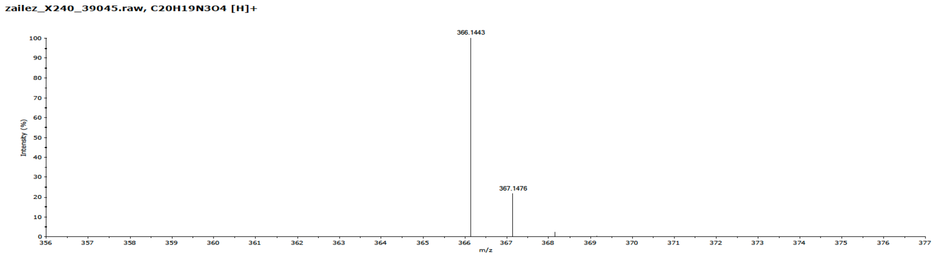

**Compound 32**

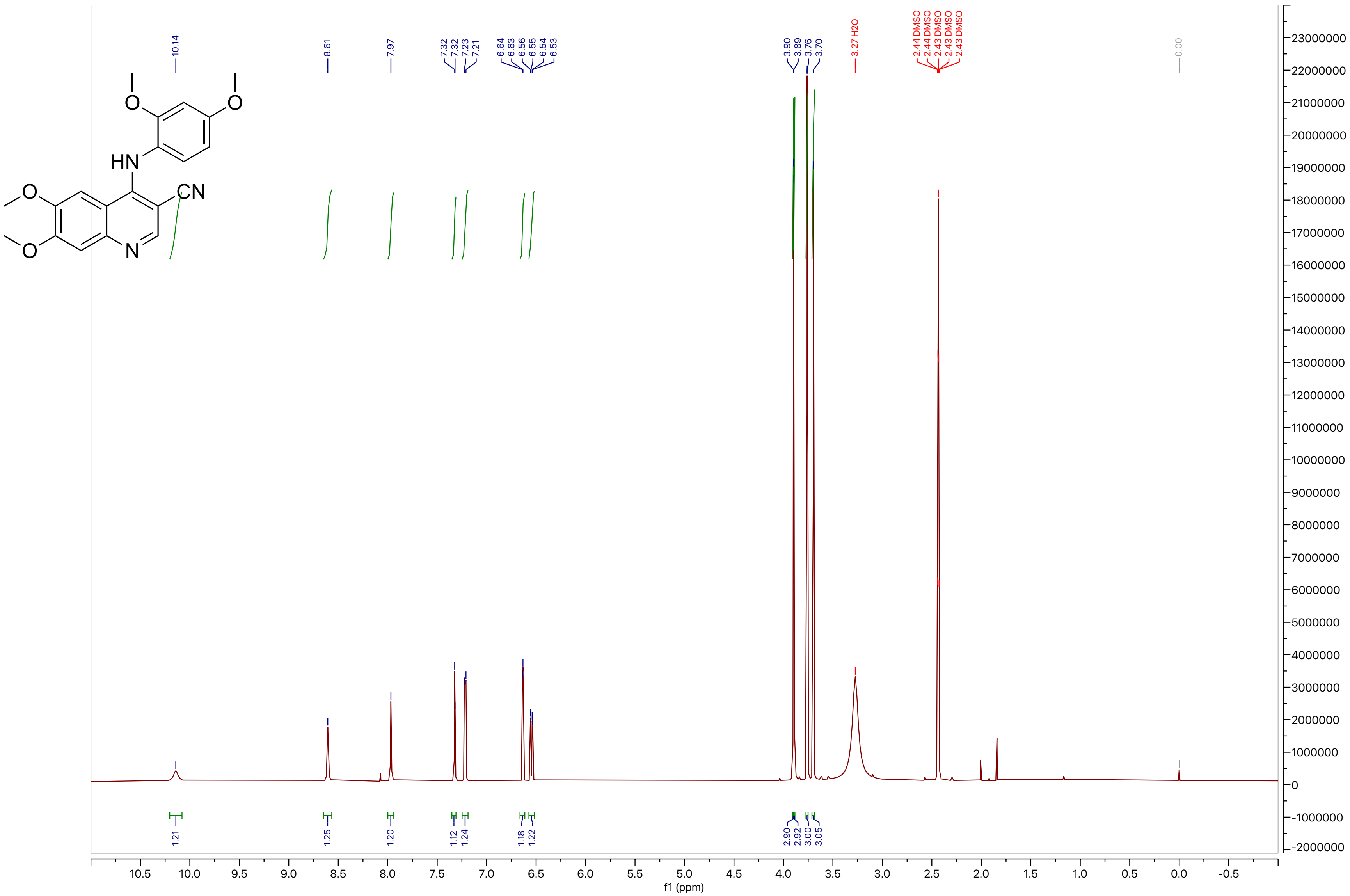

10.14
8.61
7.97
7.32
7.32
7.23
7.21
6.64
6.63
6.56
6.55
6.54
6.53
3.90
3.89
3.76
3.70
3.27 H2O
2.44 DMSO
2.44 DMSO
2.43 DMSO
2.43 DMSO
2.43 DMSO
0.00
1.21
1.25
1.20
1.12
1.24
1.18
1.22
2.90
2.92
3.00
3.05
f1 (ppm)

**Compound 32**

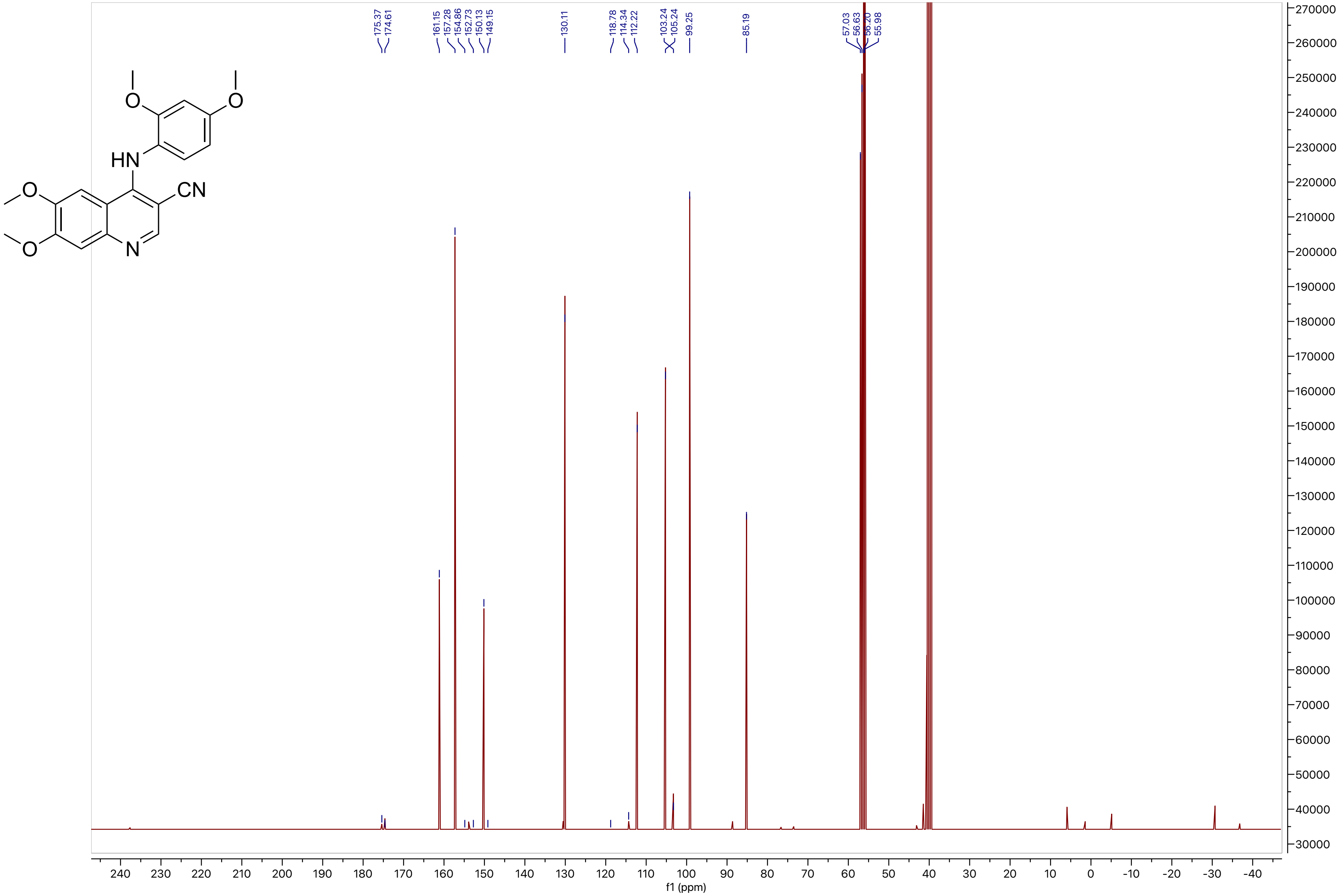
HN
CN
N
O
175.37
174.61
161.15
157.28
154.86
152.73
150.13
149.15
130.11
118.78
114.34
112.22
103.24
105.24
99.25
85.19
57.03
56.63
56.20
55.98
f1 (ppm)

**Compound 33**

LCMS

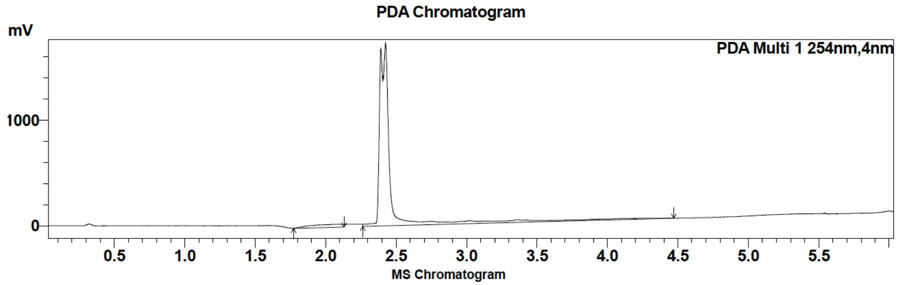

| Peak # | Ret. time | Area | Area % |
|---|---|---|---|
| 1 | 2.1 | 451294 | 4.90 |
| 2 | 2.4 | 8763122 | 95.10 |

HRMS

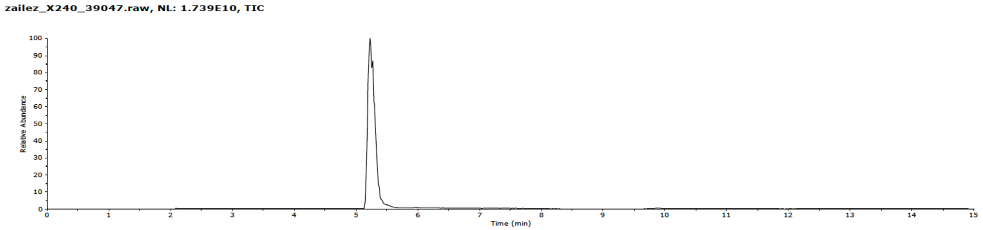

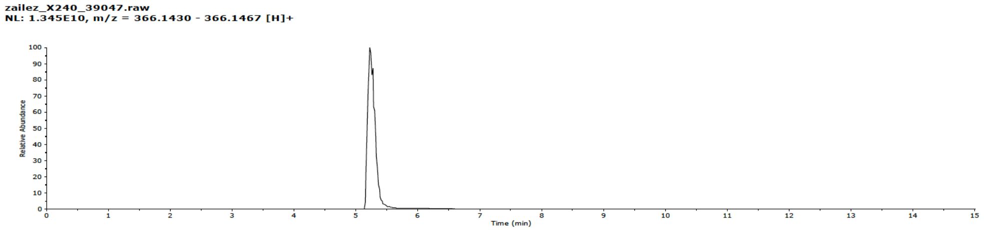

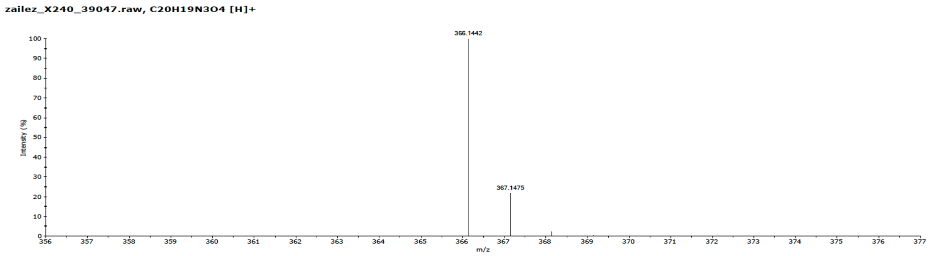

**Compound 33**

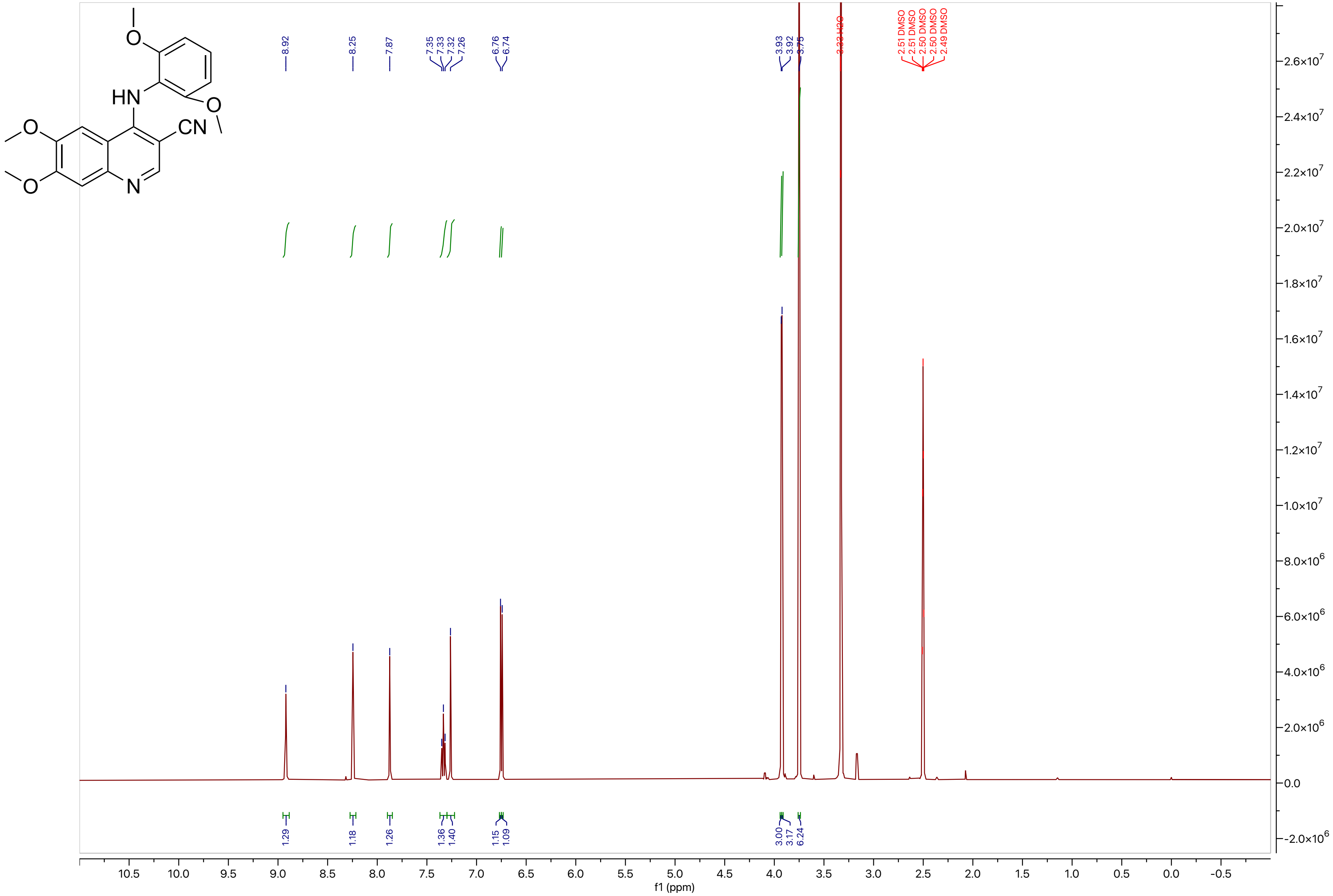
HN
CN
N
O
8.92
8.25
7.87
7.35
7.33
7.32
7.26
6.76
6.74
3.93
3.92
3.75
3.33 H2O
2.51 DMSO
2.51 DMSO
2.50 DMSO
2.50 DMSO
2.49 DMSO
1.29
1.18
1.26
1.36
1.40
1.15
1.09
3.00
3.17
6.24
f1 (ppm)

### Compound 33

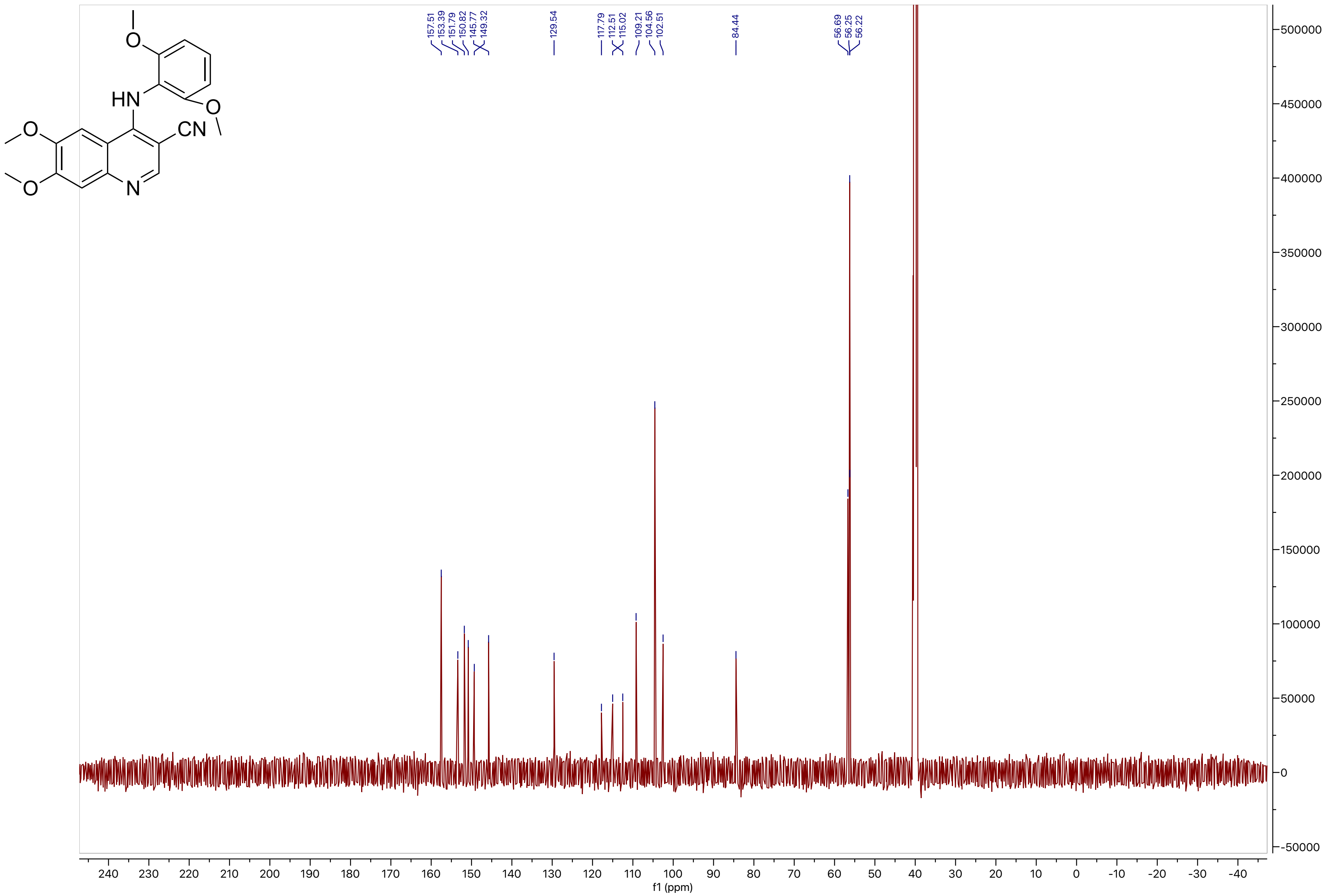
157.51
153.39
151.79
150.82
145.77
149.32
129.54
117.79
112.51
115.02
109.21
104.56
102.51
84.44
56.69
56.25
56.22
HN
CN
O
N
f1 (ppm)

**Compound 34**

LCMS

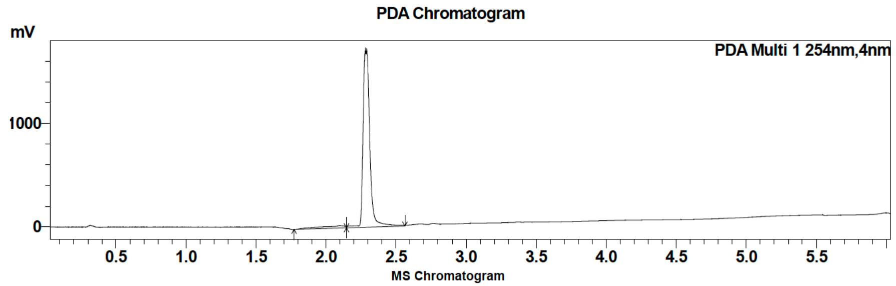

| Peak # | Ret. time | Area | Area % |
|---|---|---|---|
| 1 | 2.1 | 220885 | 3.68 |
| 2 | 2.4 | 5779494 | 96.32 |

HRMS

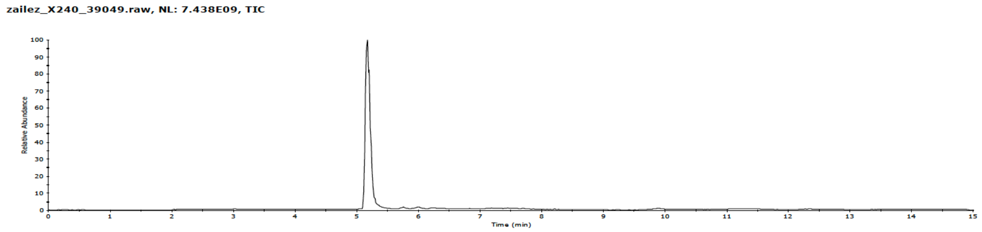

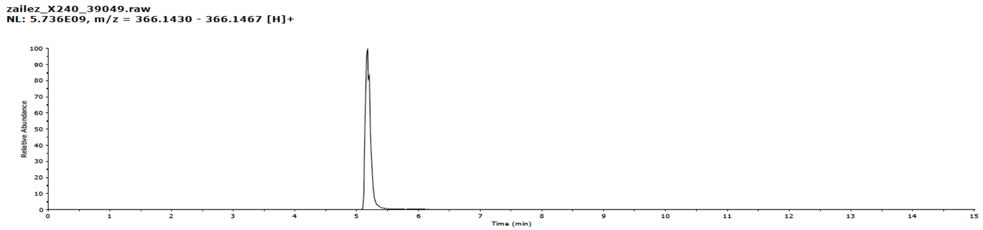

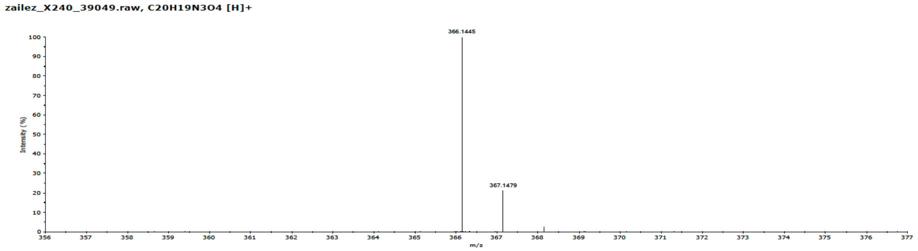

**Compound 34**

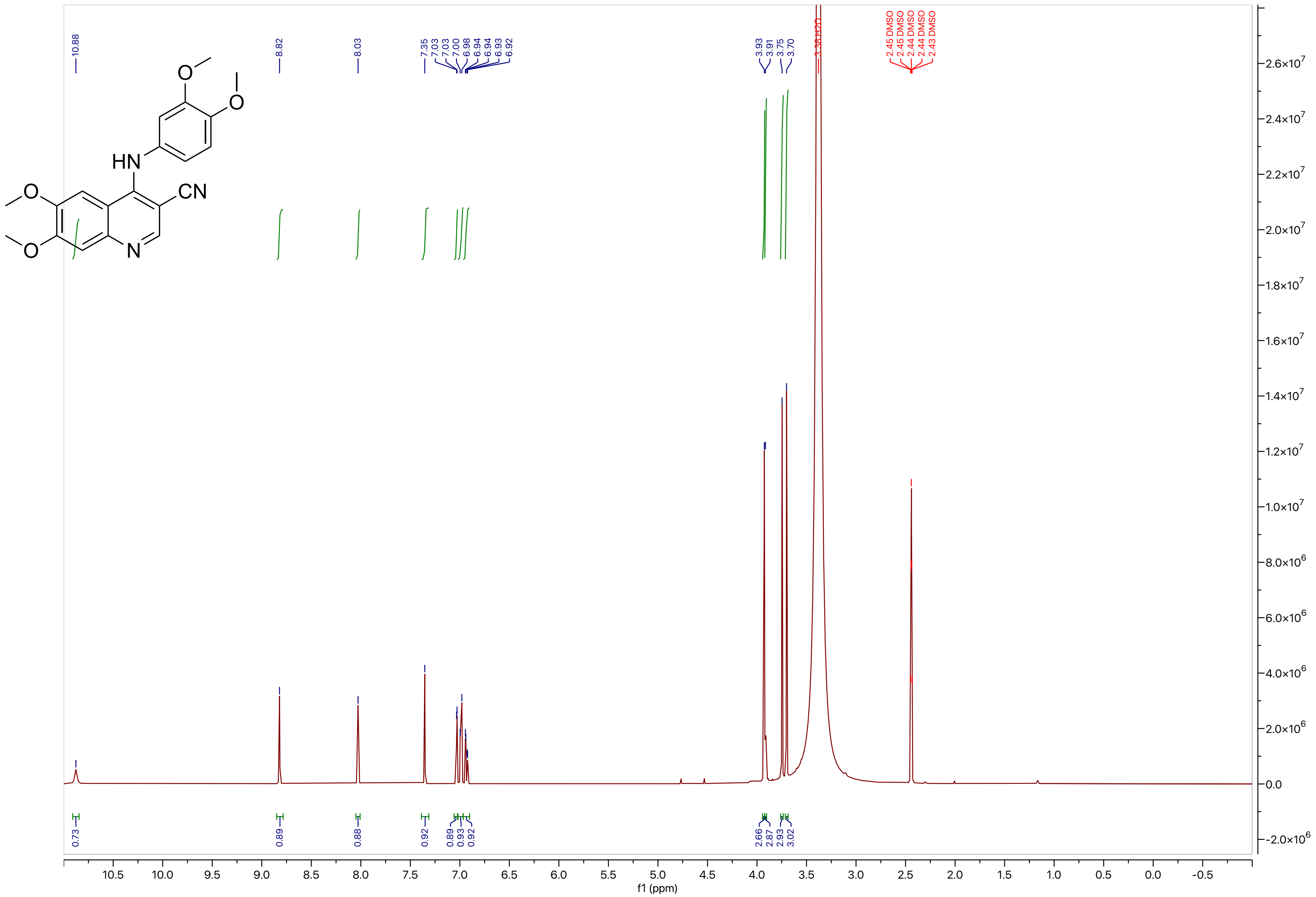
10.88
8.82
8.03
7.35
7.03
7.03
7.00
6.98
6.94
6.94
6.93
6.92
3.93
3.91
3.75
3.70
3.36 H2O
2.45 DMSO
2.45 DMSO
2.44 DMSO
2.44 DMSO
2.43 DMSO
0.73
0.89
0.88
0.92
0.89
0.93
0.92
2.66
2.87
2.93
3.02
f1 (ppm)

**Compound 34**

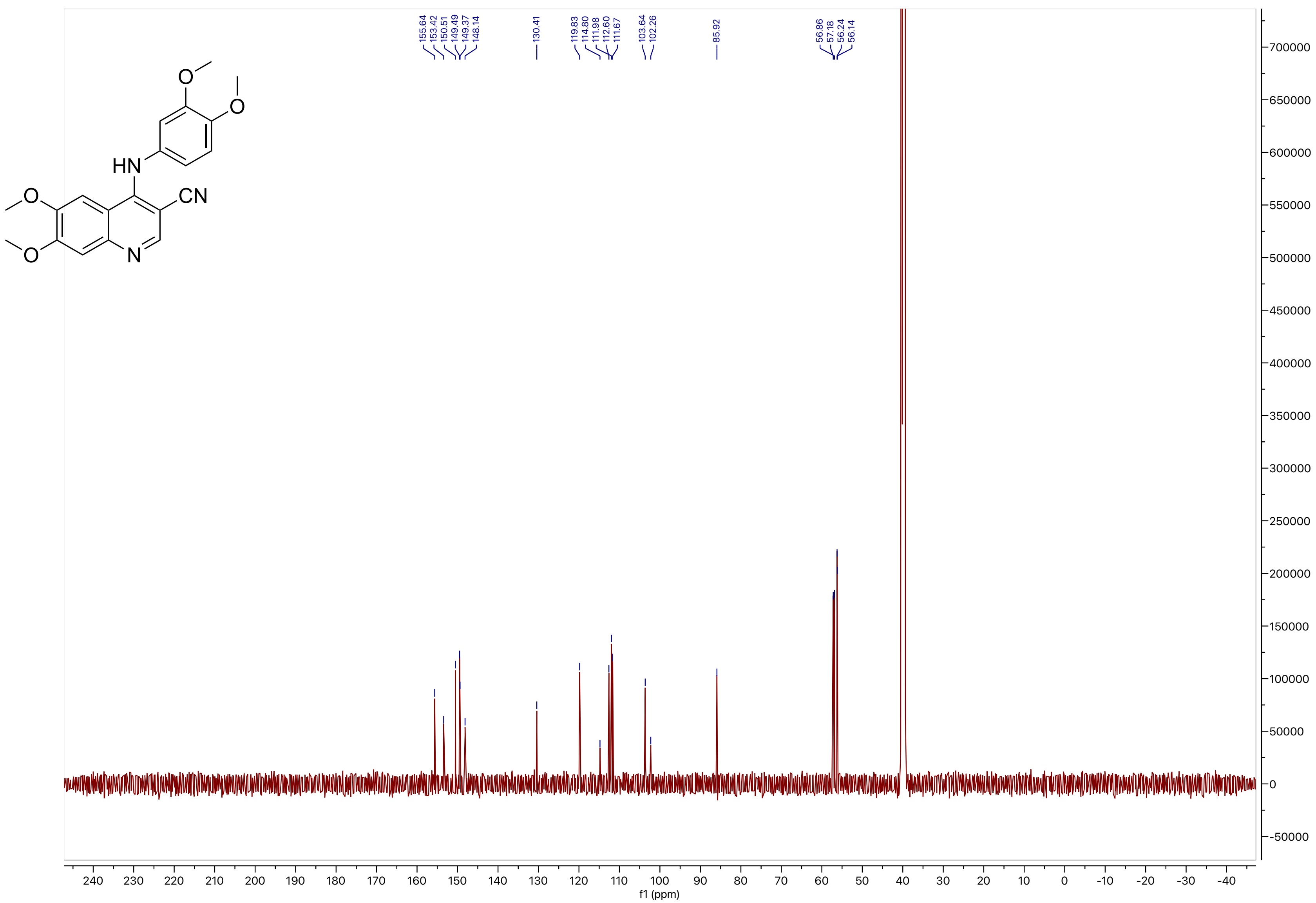
155.64
153.42
150.51
149.49
149.37
148.14
130.41
119.83
114.80
111.98
112.60
111.67
103.64
102.26
85.92
56.86
57.18
56.24
56.14
HN
CN
N
O
f1 (ppm)

**Compound 35**

LCMS

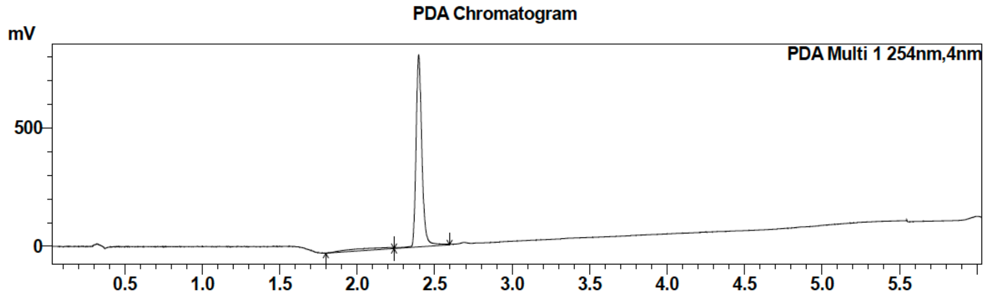

| Peak # | Ret. time | Area | Area % |
|---|---|---|---|
| 1 | 2.4 | 2143249 | 100 |

MS Chromatogram

HRMS

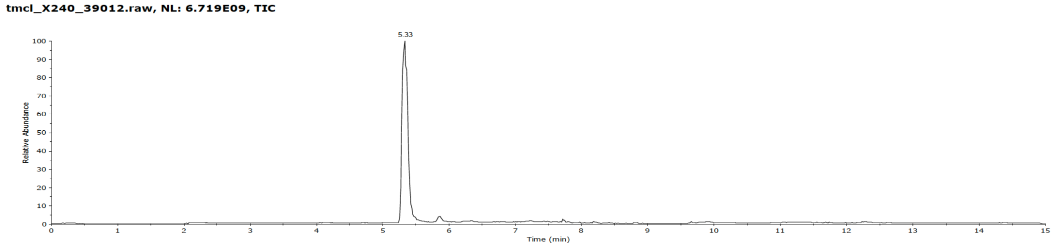

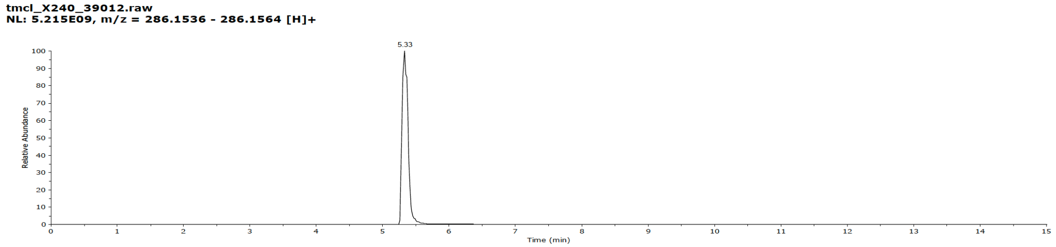

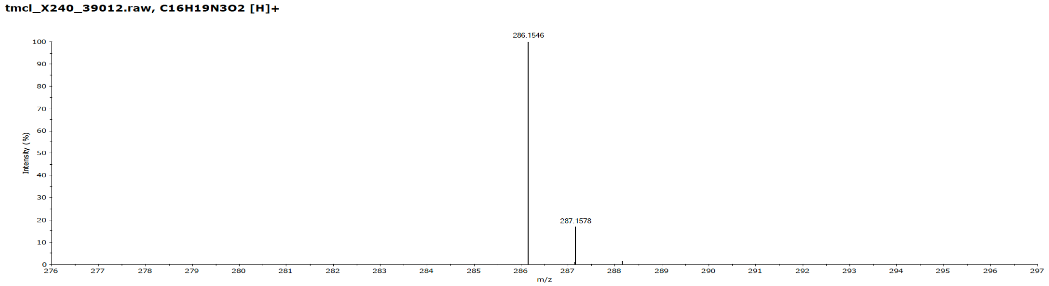

**Compound 35**

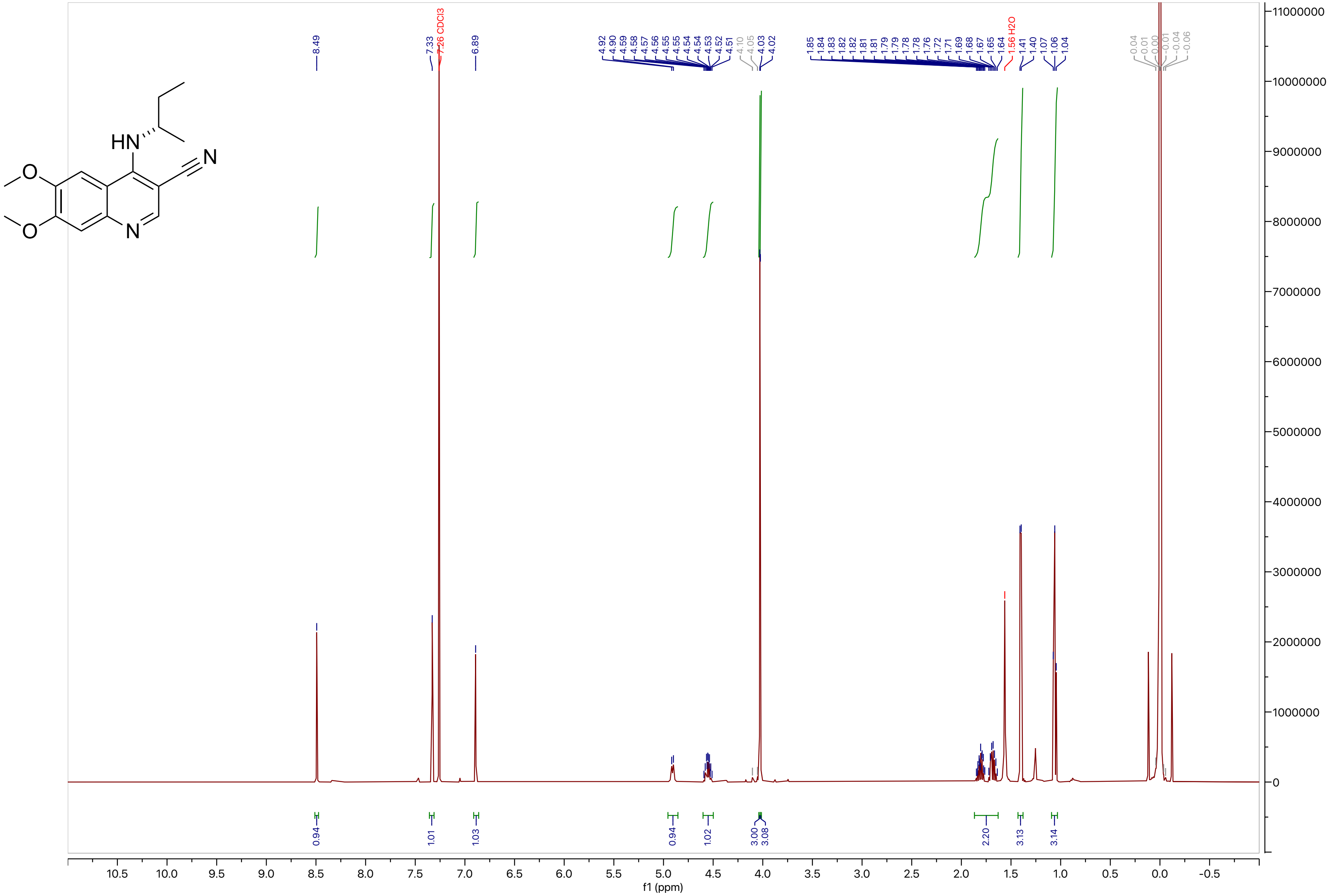

**Compound 35**

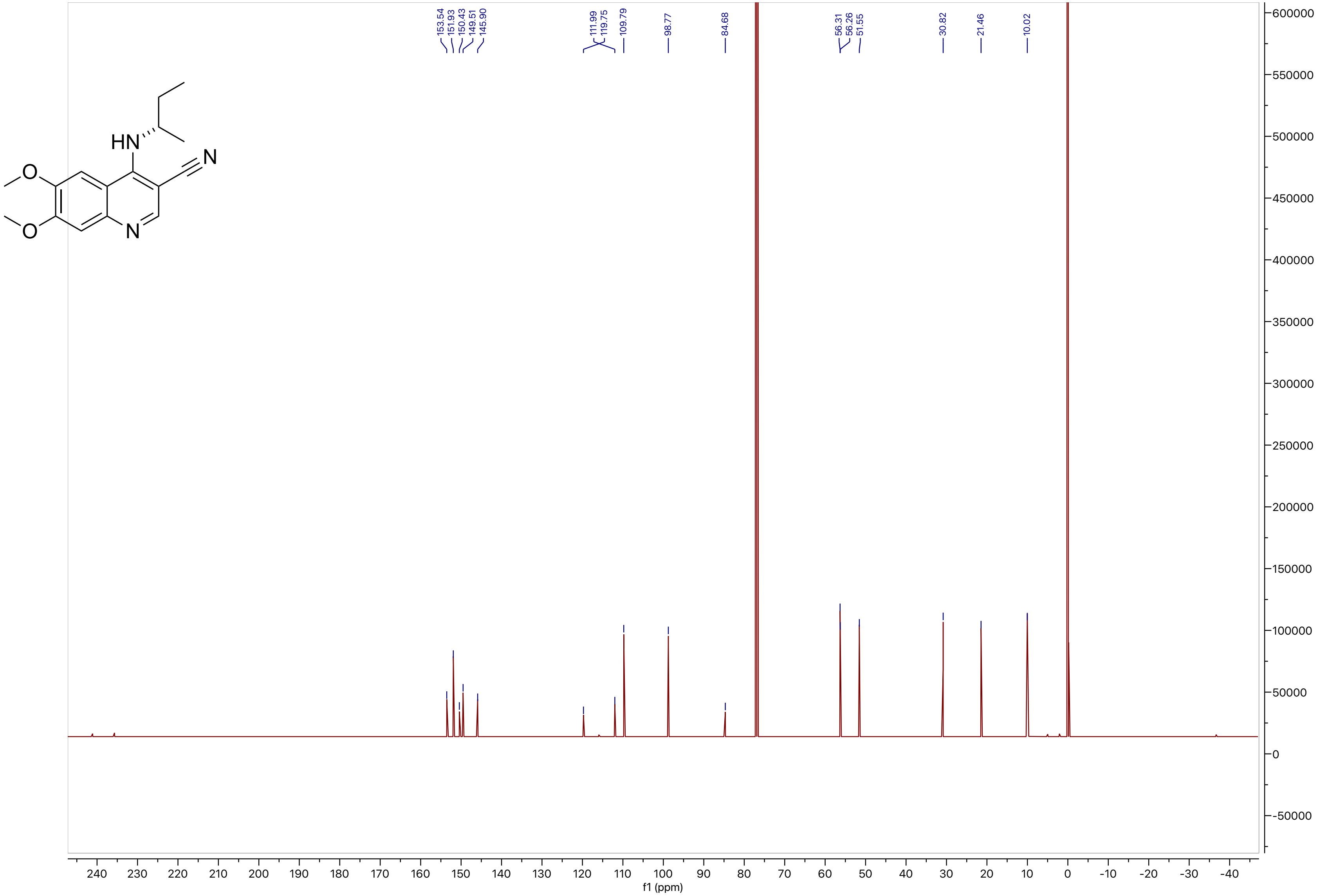
HN
N
O
O
N
153.54
151.93
150.43
149.51
145.90
111.99
119.75
109.79
98.77
84.68
56.31
56.26
51.55
30.82
21.46
10.02
f1 (ppm)

**Compound 36**

LCMS

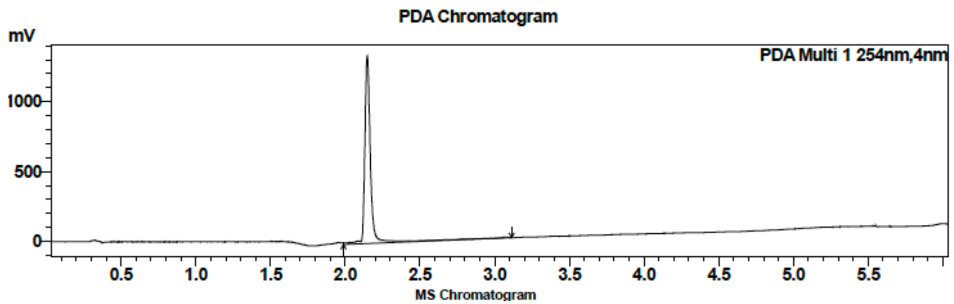

| Peak # | Ret. time | Area | Area % |
|---|---|---|---|
| 1 | 2.1 | 3622349 | 100.00 |

HRMS

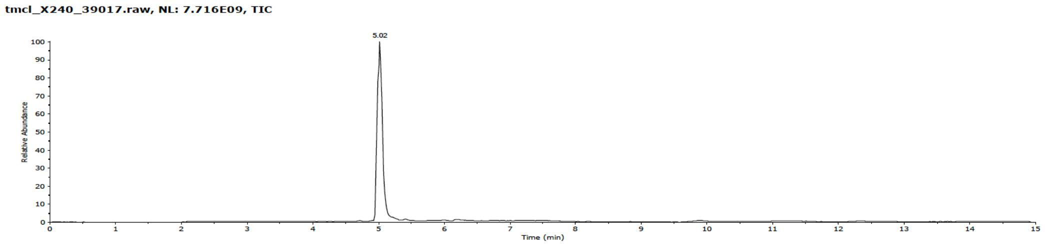

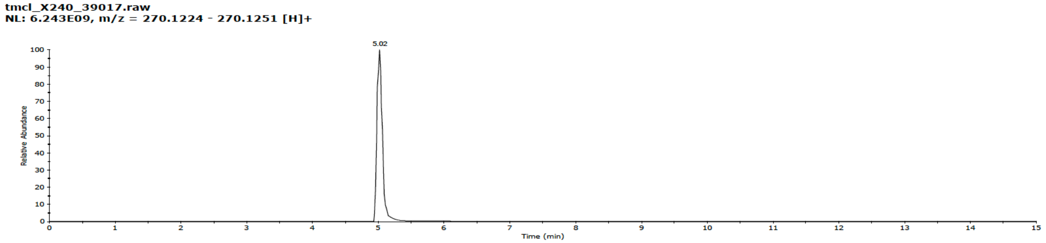

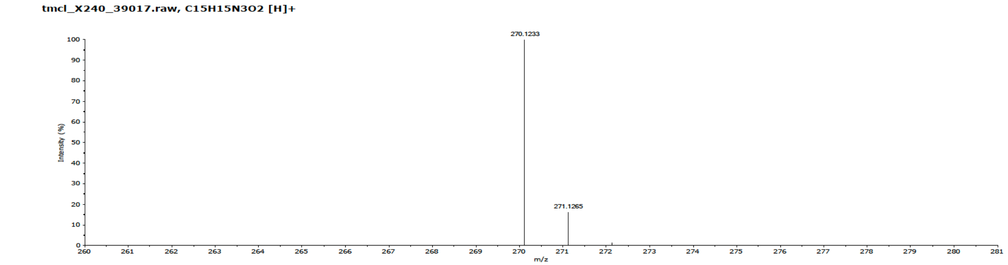

**Compound 36**

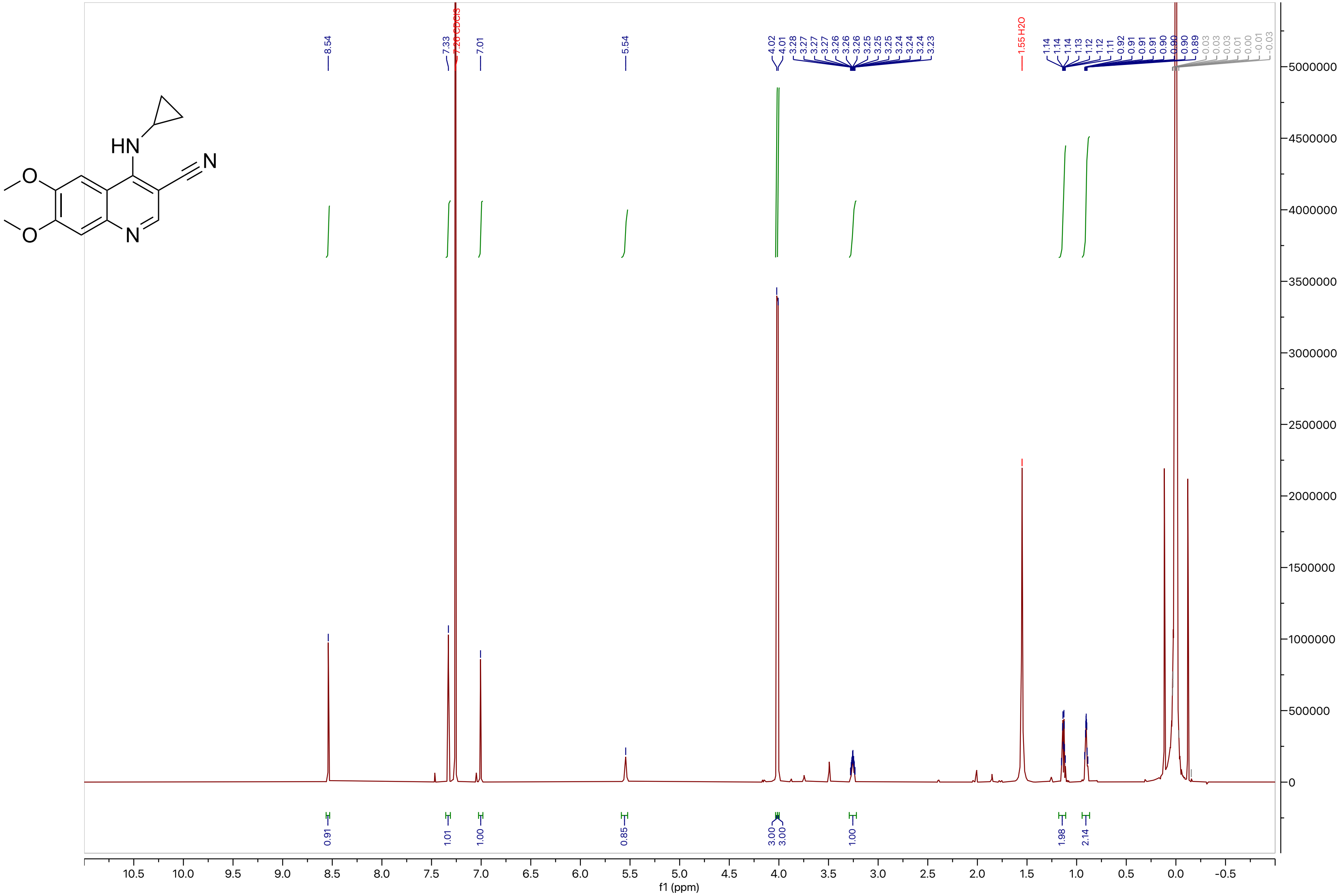

HN
N
O
O
N
8.54
7.33
7.26 CDCl3
7.01
5.54
4.02
4.01
3.28
3.27
3.27
3.27
3.26
3.26
3.26
3.25
3.25
3.25
3.24
3.24
3.24
3.23
1.55 H2O
1.14
1.14
1.14
1.13
1.12
1.12
1.11
0.92
0.91
0.91
0.91
0.90
0.90
0.90
0.89
0.91
1.01
1.00
0.85
3.00
3.00
1.00
1.98
2.14
10.5
10.0
9.5
9.0
8.5
8.0
7.5
7.0
6.5
6.0
5.5
5.0
4.5
4.0
3.5
3.0
2.5
2.0
1.5
1.0
0.5
0.0
-0.5
f1 (ppm)
5000000
4500000
4000000
3500000
3000000
2500000
2000000
1500000
1000000
500000
0

**Compound 36**

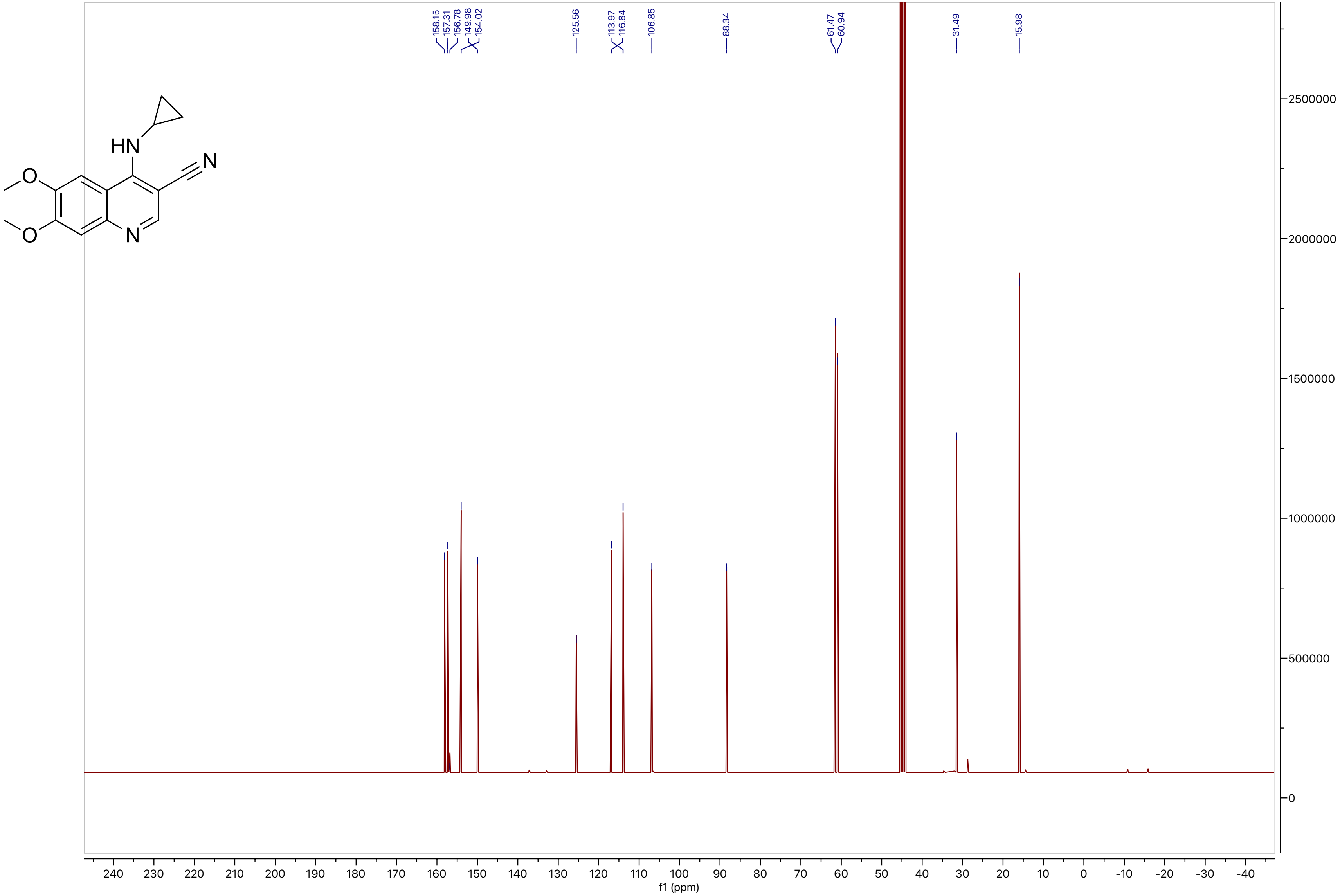
HN
N
O
O
N
158.15
157.31
156.78
149.98
154.02
125.56
113.97
116.84
106.85
88.34
61.47
60.94
31.49
15.98
2500000
2000000
1500000
1000000
500000
0
240 230 220 210 200 190 180 170 160 150 140 130 120 110 100 90 80 70 60 50 40 30 20 10 0 -10 -20 -30 -40
f1 (ppm)

**Compound 37**

LCMS

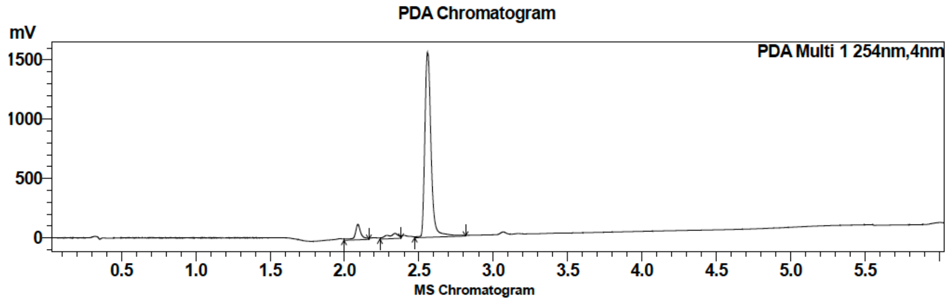

| Peak # | Ret. time | Area | Area % |
|---|---|---|---|
| 1 | 2.1 | 162065 | 3.32 |
| 2 | 2.3 | 116063 | 2.38 |
| 3 | 2.6 | 4597589 | 94.30 |

HRMS

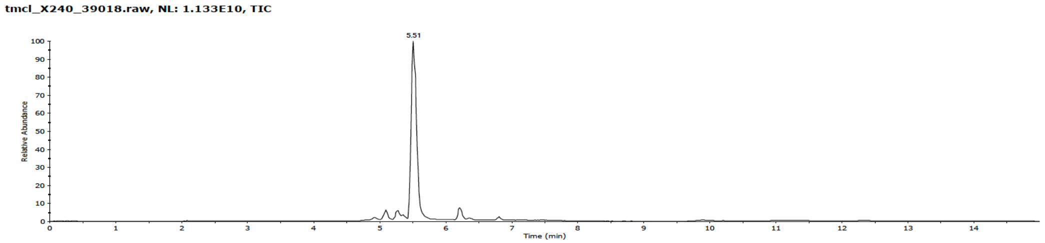

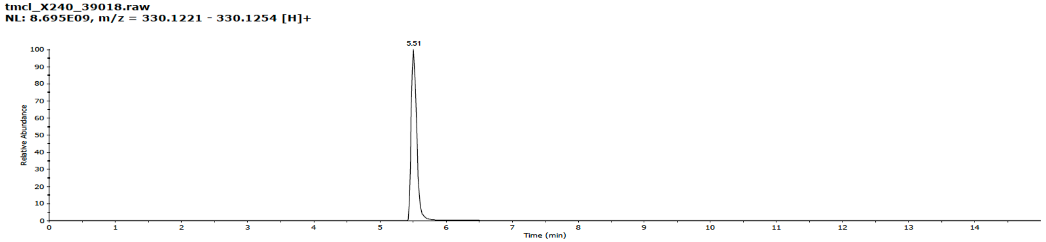

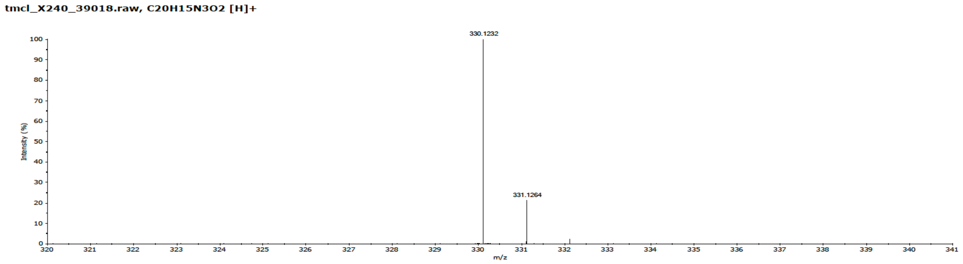

**Compound 37**

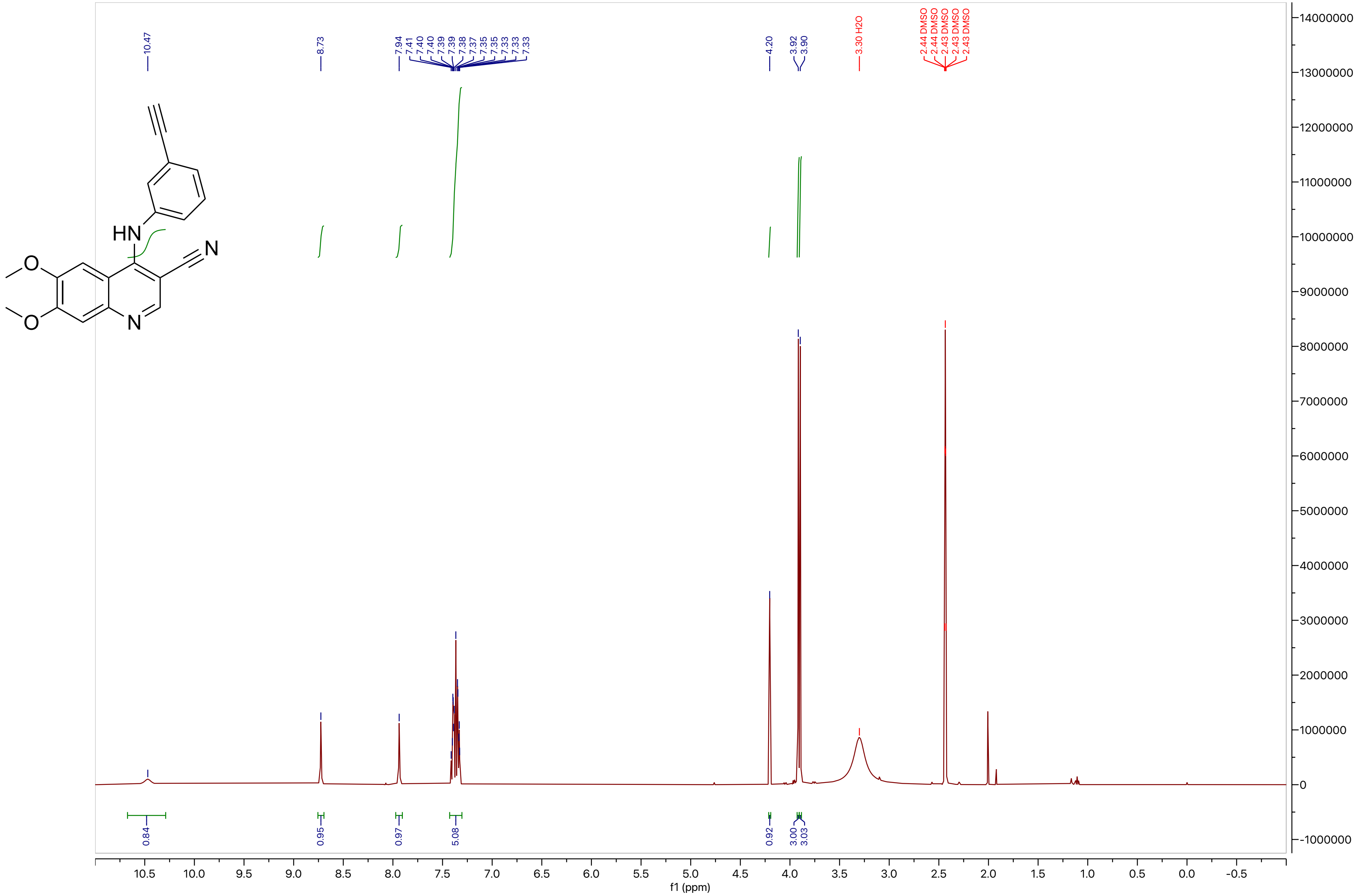

10.47
8.73
7.94
7.41
7.40
7.40
7.39
7.39
7.38
7.37
7.35
7.35
7.33
7.33
7.33
4.20
3.92
3.90
3.30 H2O
2.44 DMSO
2.44 DMSO
2.43 DMSO
2.43 DMSO
2.43 DMSO
HN
N
O
O
N
0.84
0.95
0.97
5.08
0.92
3.00
3.03
f1 (ppm)

**Compound 37**

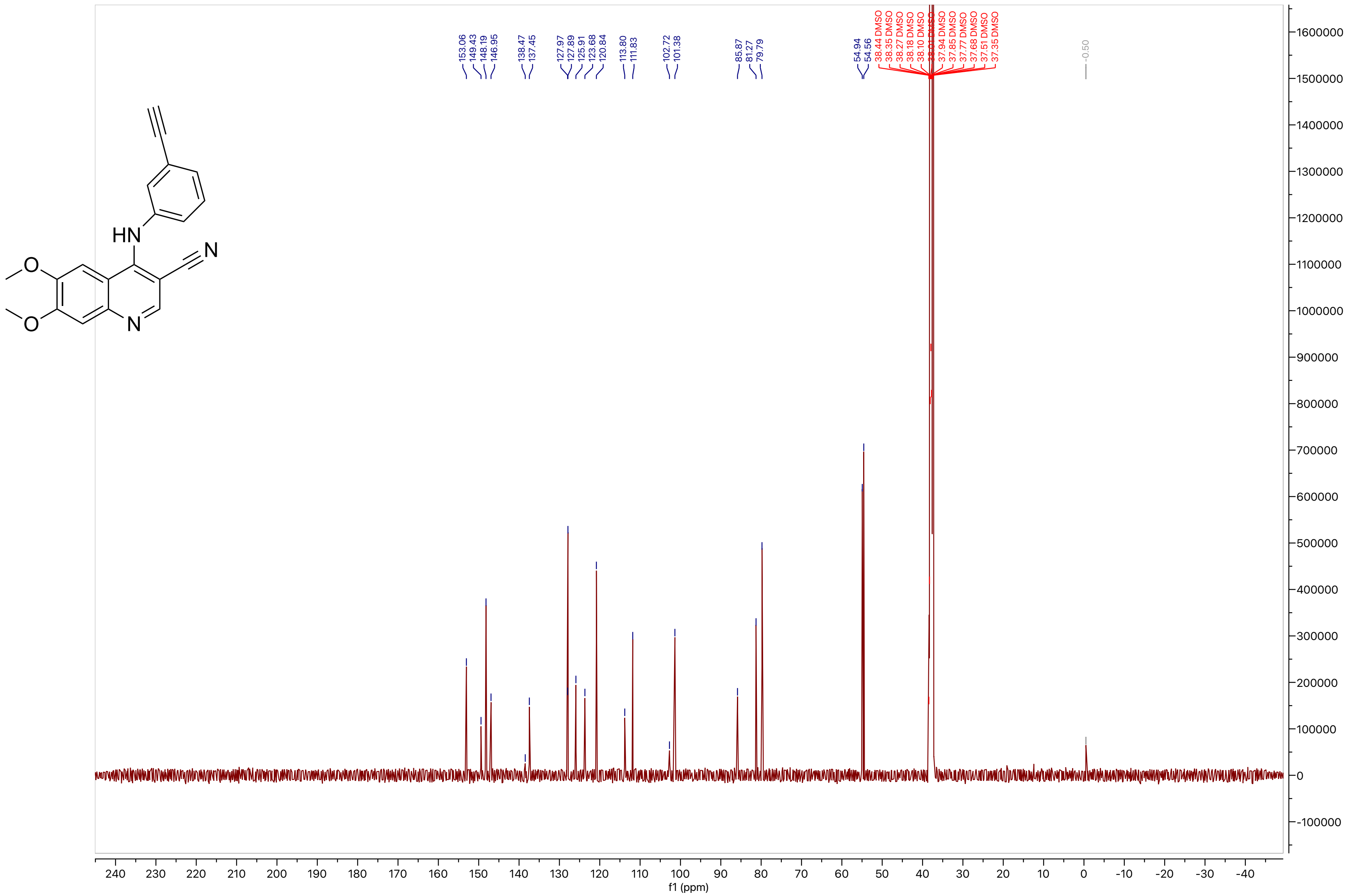
153.06
149.43
148.19
146.95
138.47
137.45
127.97
127.89
125.91
123.68
120.84
113.80
111.83
102.72
101.38
85.87
81.27
79.79
54.94
54.56
38.44 DMSO
38.35 DMSO
38.27 DMSO
38.18 DMSO
38.10 DMSO
37.94 DMSO
37.85 DMSO
37.77 DMSO
37.68 DMSO
37.51 DMSO
37.35 DMSO
-0.50
f1 (ppm)
HN
N
O
O
N

**Compound 38**

LCMS

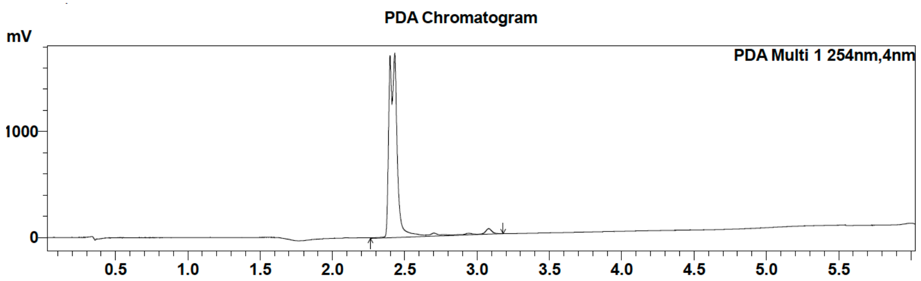

| Peak # | Ret. time | Area | Area % |
|---|---|---|---|
| 1 | 2.4 | 6769479 | 100.00 |

MS Chromatogram

HRMS

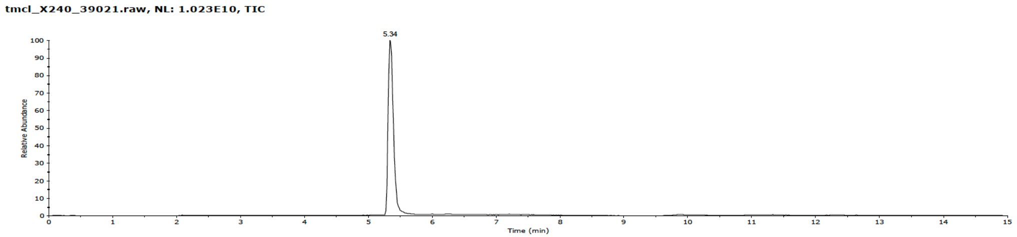

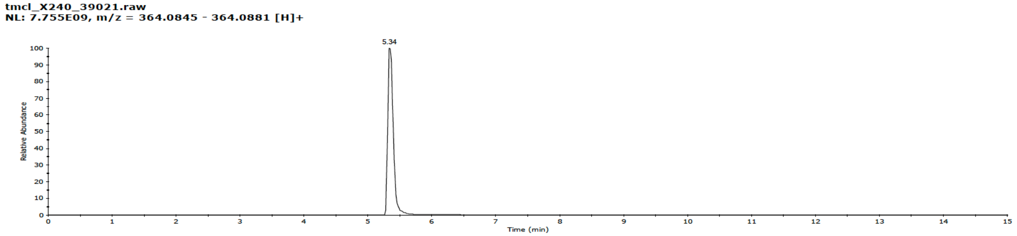

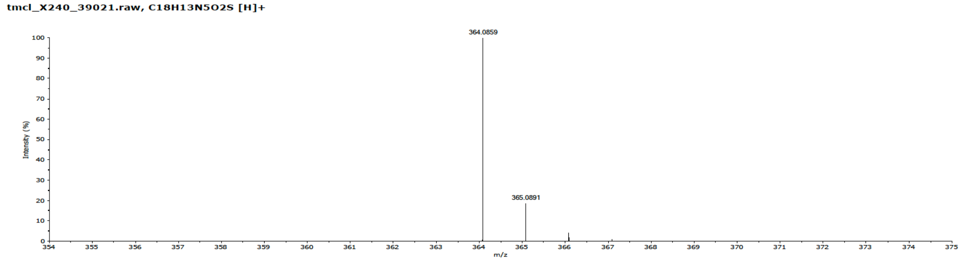

**Compound 38**

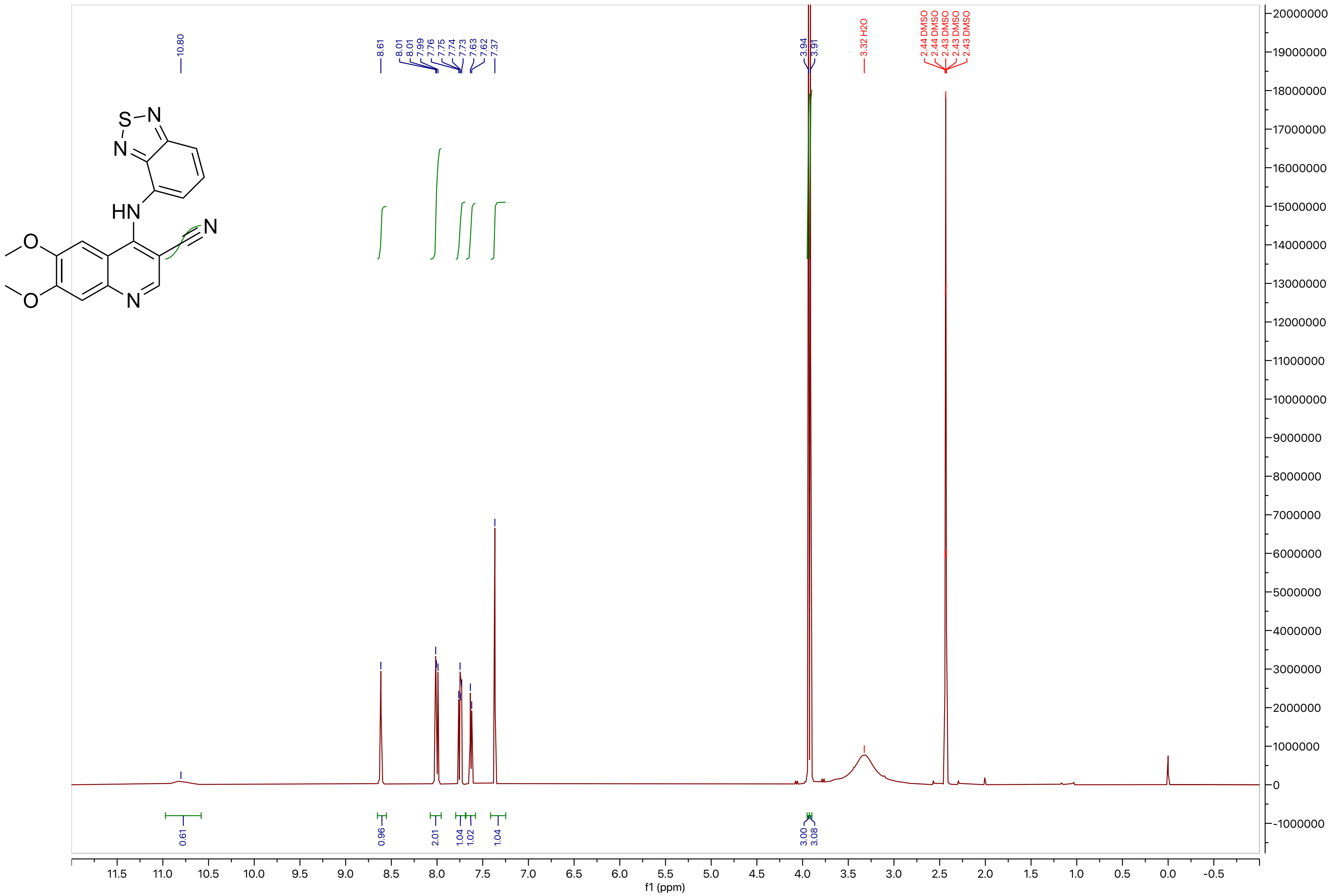
10.80
8.61
8.01
8.01
7.99
7.76
7.75
7.74
7.73
7.63
7.62
7.37
3.94
3.91
3.32 H2O
2.44 DMSO
2.44 DMSO
2.43 DMSO
2.43 DMSO
2.43 DMSO
0.61
0.96
2.01
1.04
1.02
1.04
3.00
3.08
f1 (ppm)

**Compound 38**

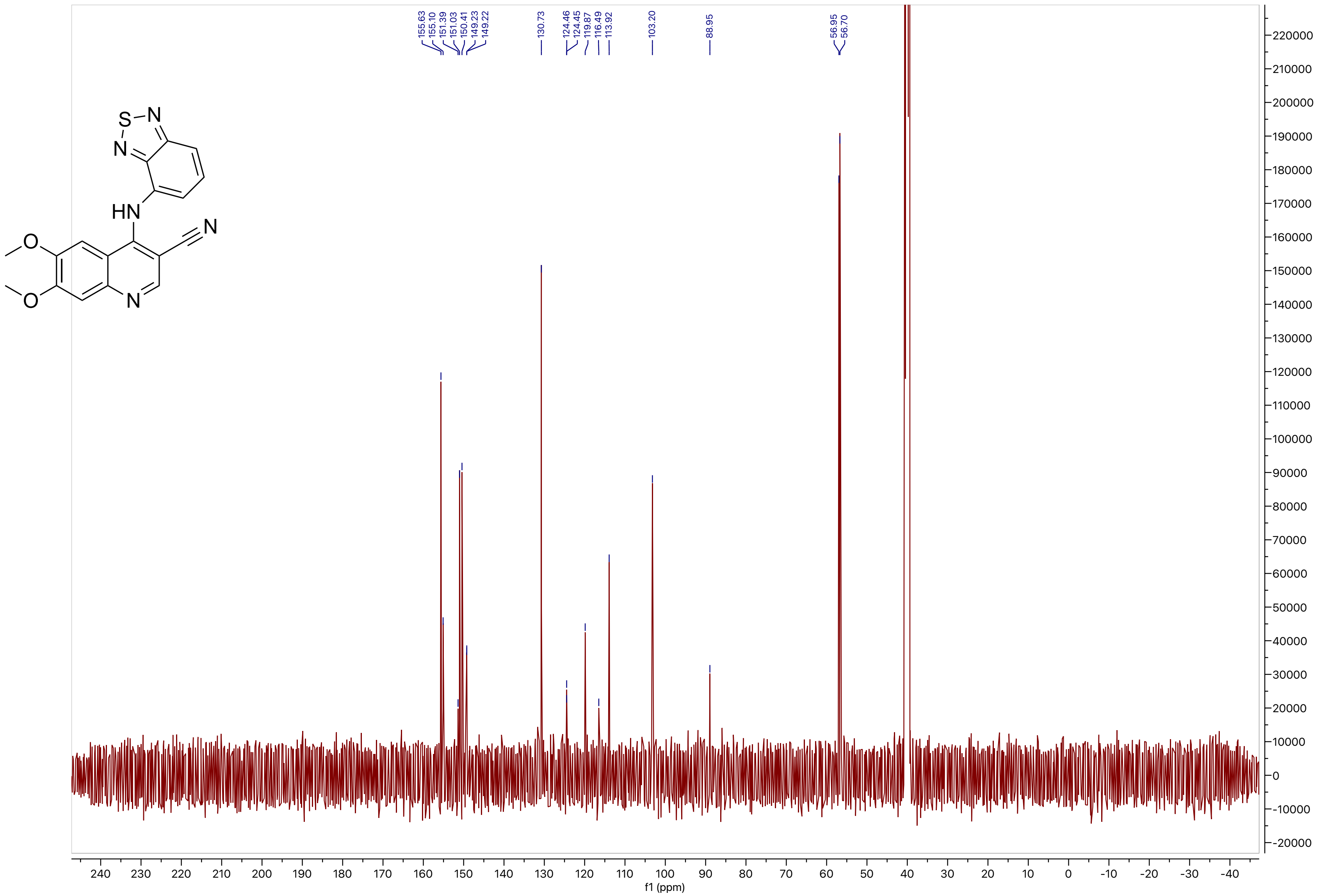

155.63
155.10
151.39
151.03
150.41
149.23
149.22
130.73
124.46
124.45
119.87
116.49
113.92
103.20
88.95
56.95
56.70
S
N
HN
O
f1 (ppm)

**Compound 39**

LCMS

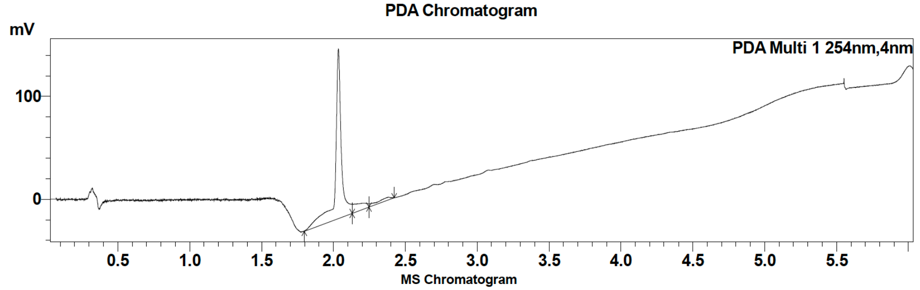

| Peak # | Ret. time | Area | Area % |
|---|---|---|---|
| 1 | 2.0 | 479088 | 95.42 |
| 2 | 2.4 | 23007 | 4.58 |

HRMS

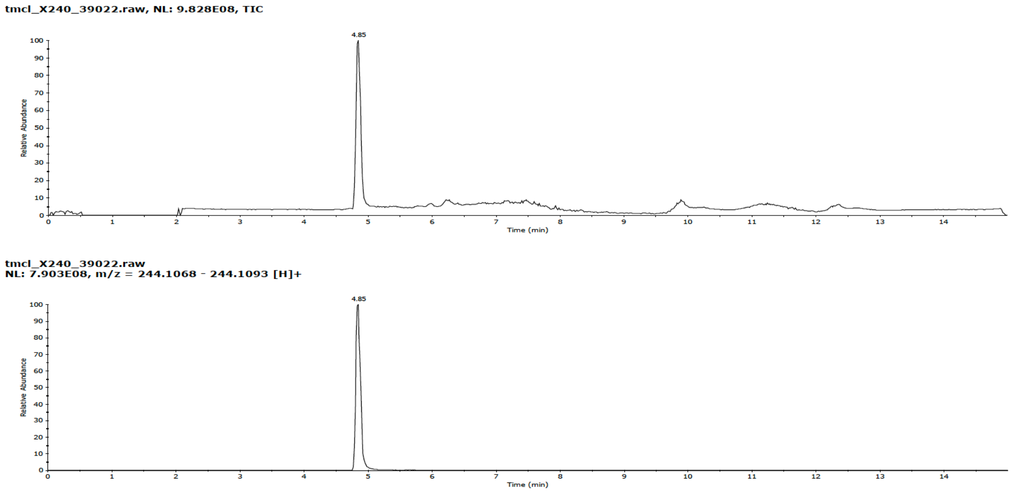

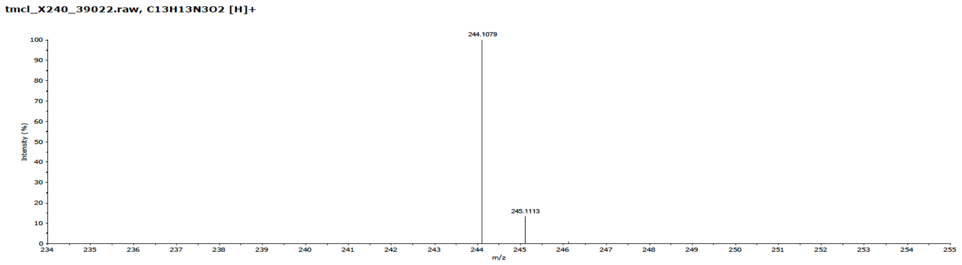

**Compound 39**

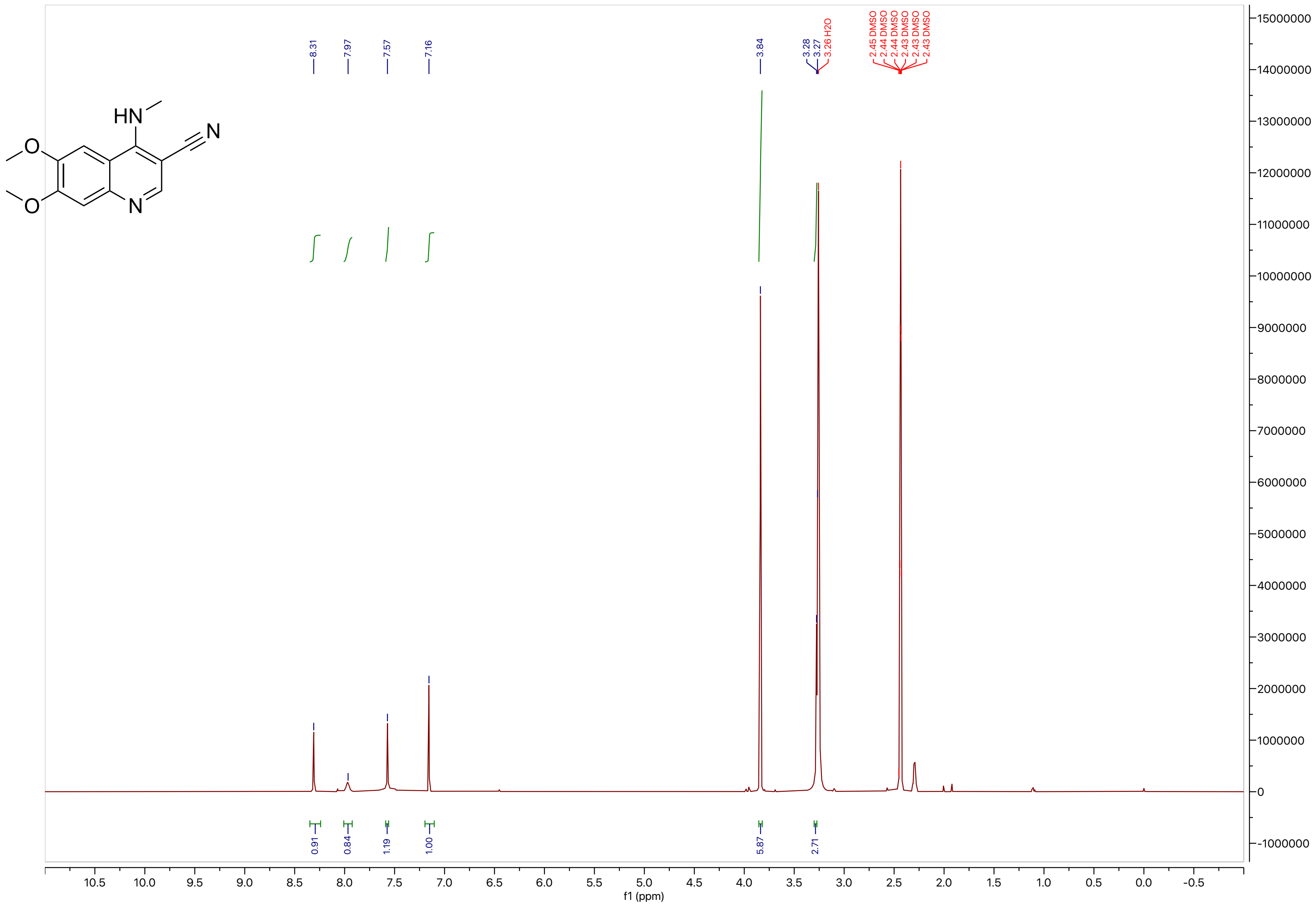

HN
O
O
N
N
8.31
7.97
7.57
7.16
3.84
3.28
3.27
3.26 H2O
2.45 DMSO
2.44 DMSO
2.44 DMSO
2.43 DMSO
2.43 DMSO
2.43 DMSO
0.91
0.84
1.19
1.00
5.87
2.71
f1 (ppm)

**Compound 39**

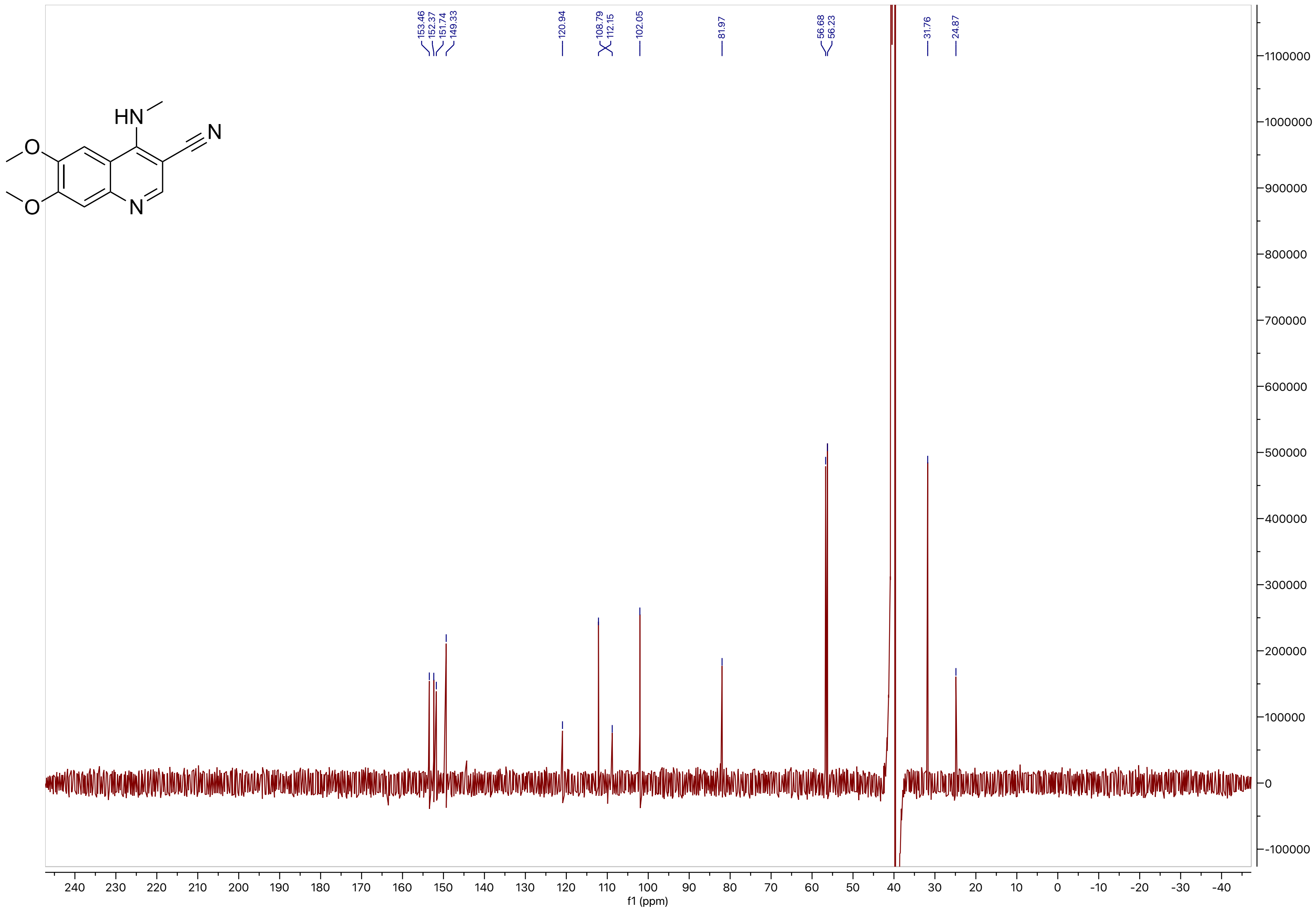

153.46
152.37
151.74
149.33
120.94
108.79
112.15
102.05
81.97
56.68
56.23
31.76
24.87
f1 (ppm)

**Compound 40**

LCMS

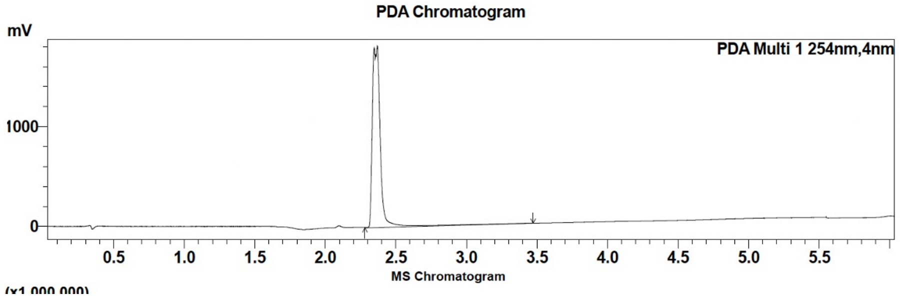

| Peak # | Ret. time | Area | Area % |
|---|---|---|---|
| 1 | 2.4 | 7147307 | 100.00 |

HRMS

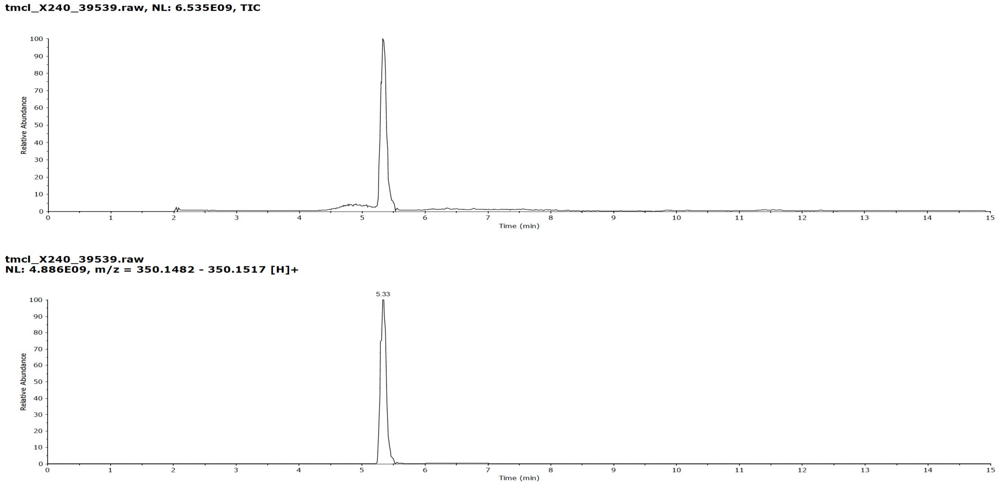

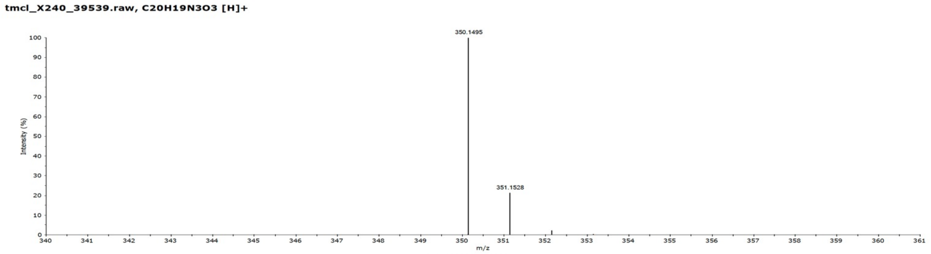

**Compound 40**

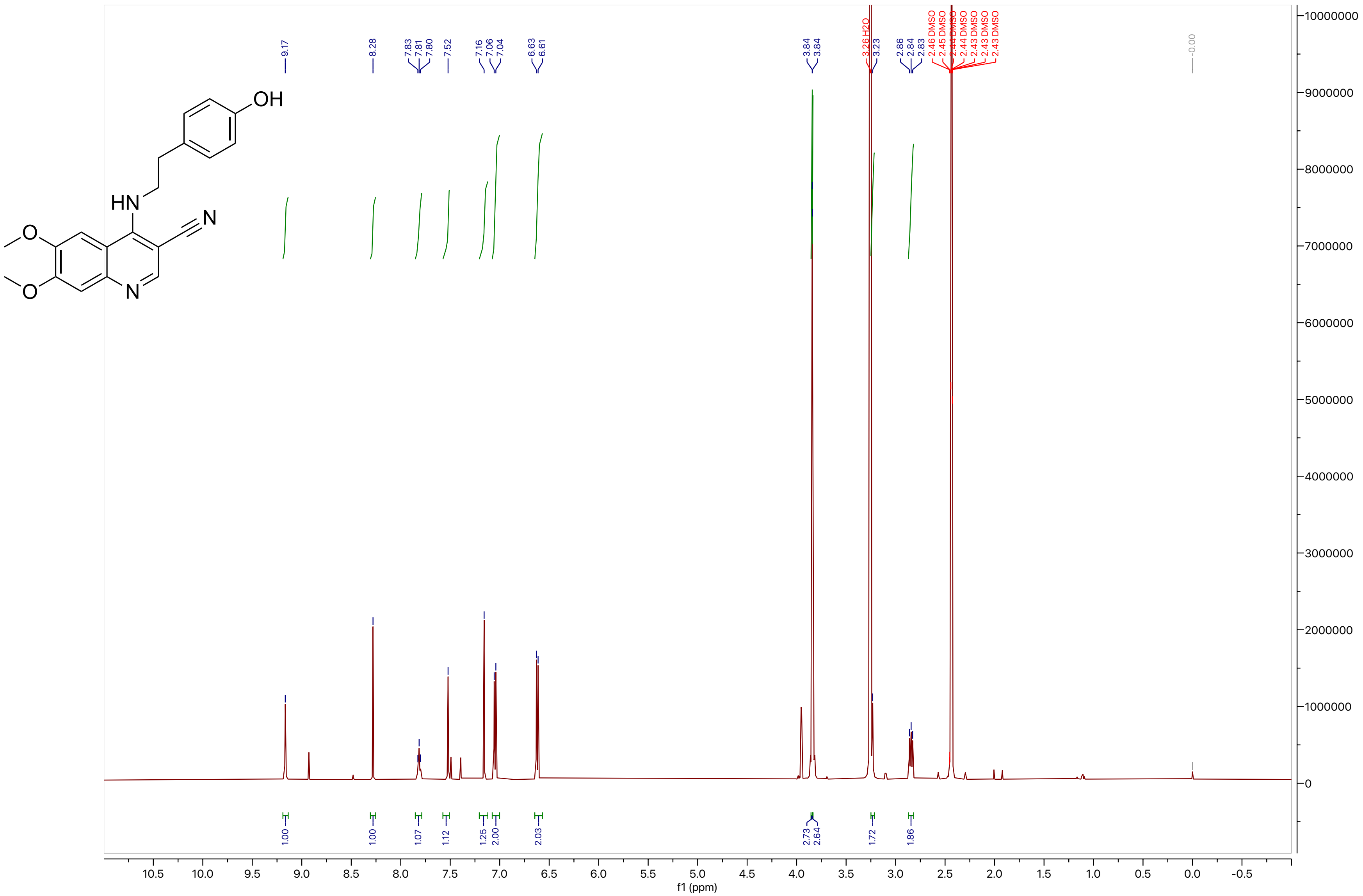
OH
HN
N
O
O
N
9.17
8.28
7.83
7.81
7.80
7.52
7.16
7.06
7.04
6.63
6.61
3.84
3.84
3.26 H2O
3.23
2.86
2.84
2.83
2.46 DMSO
2.45 DMSO
2.44 DMSO
2.44 DMSO
2.43 DMSO
2.43 DMSO
2.43 DMSO
-0.00
1.00
1.00
1.07
1.12
1.25
2.00
2.03
2.73
2.64
1.72
1.86
f1 (ppm)

### Compound 40

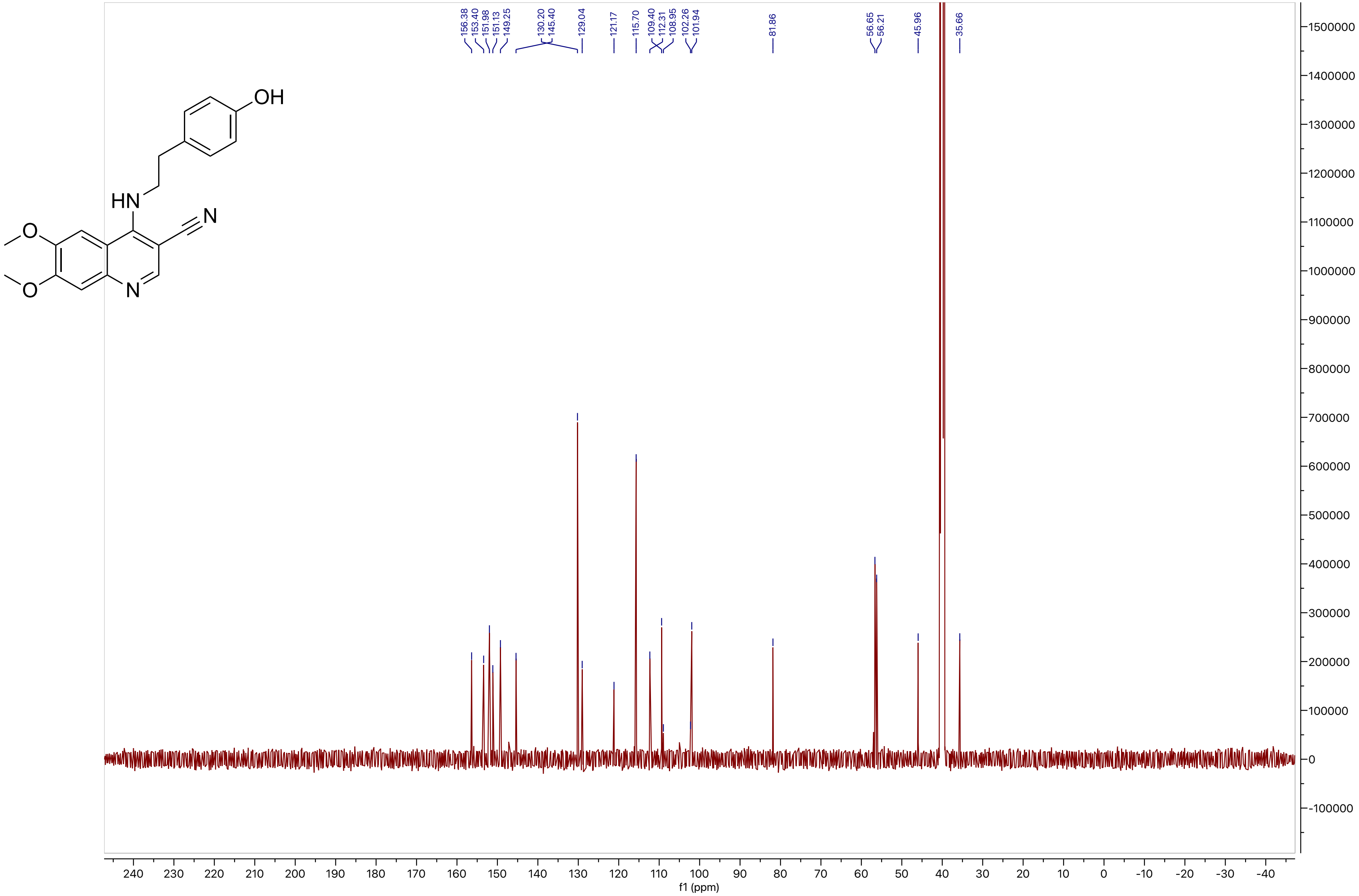

OH
HN
N
O
O
N
156.38
153.40
151.98
151.13
149.25
130.20
145.40
129.04
121.17
115.70
109.40
112.31
108.95
102.26
101.94
81.86
56.65
56.21
45.96
35.66
f1 (ppm)

#### Compound 41

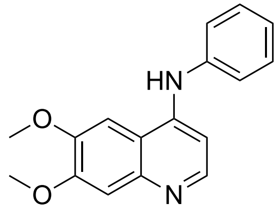

LCMS

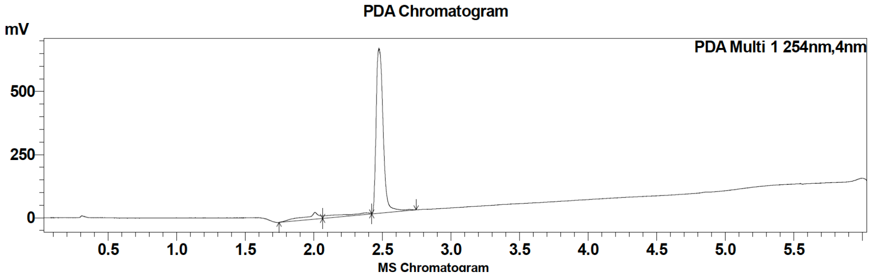

| Peak # | Ret. time | Area | Area % |
|---|---|---|---|
| 1 | 2.4 | 49268 | 2.08 |
| 2 | 2.5 | 2318126 | 97.91 |

HRMS

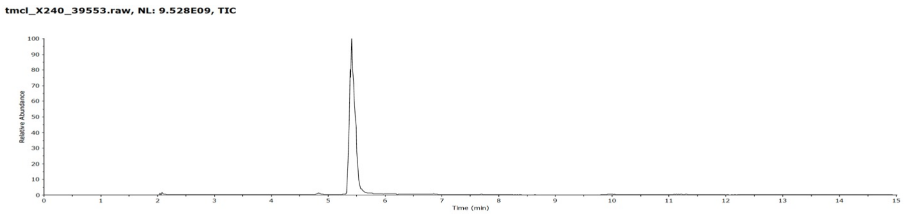

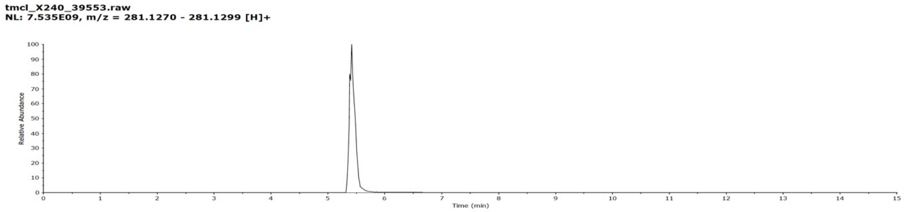

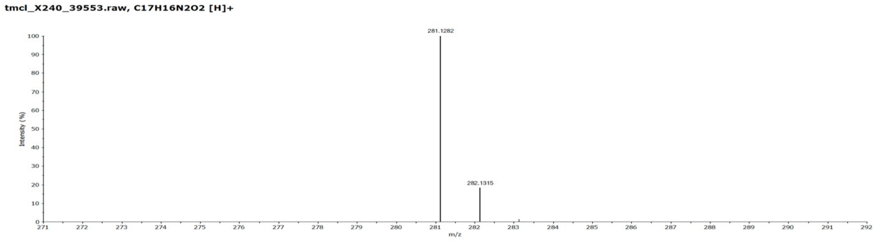

**Compound 41**

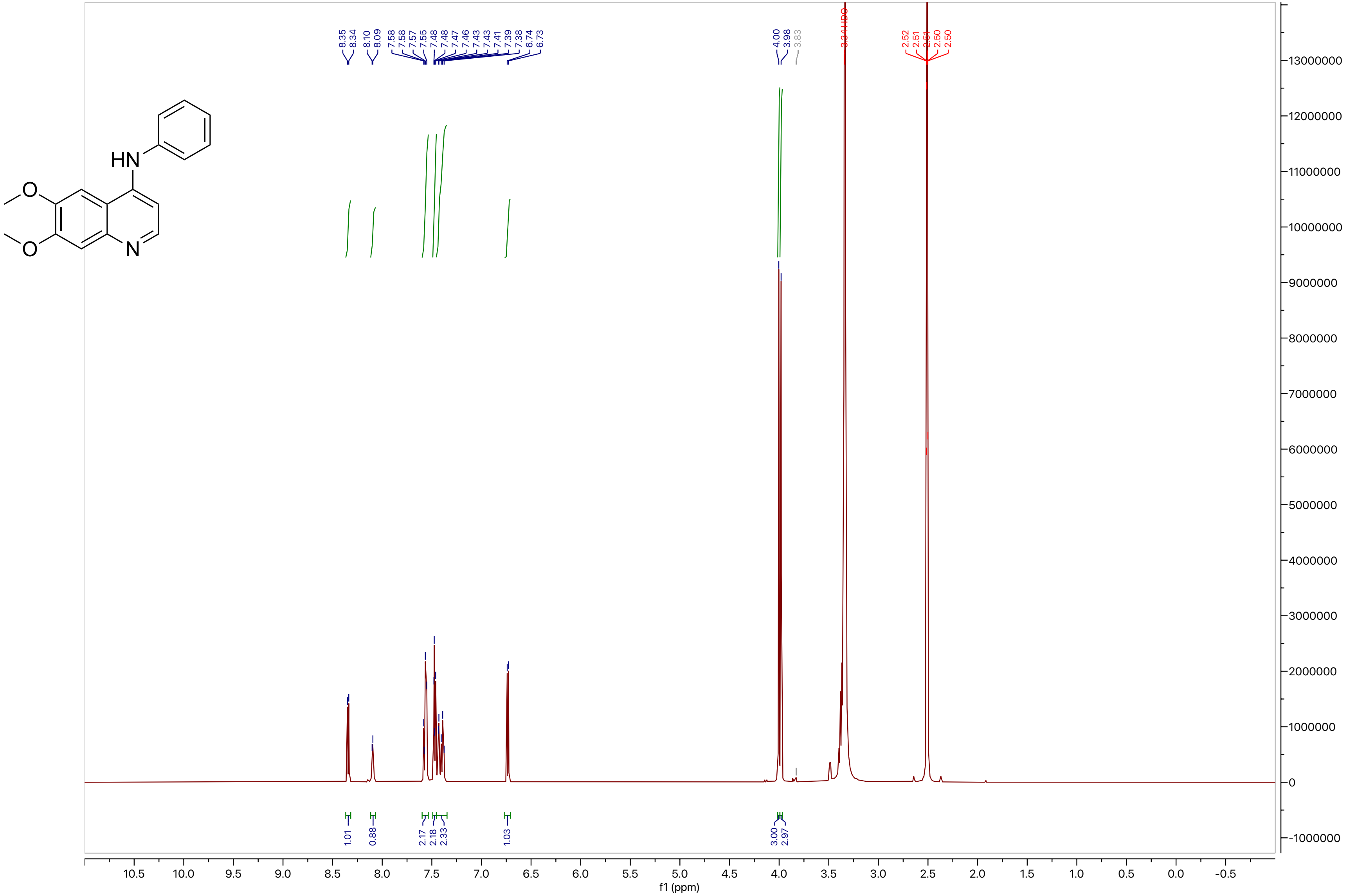

**Compound 41**

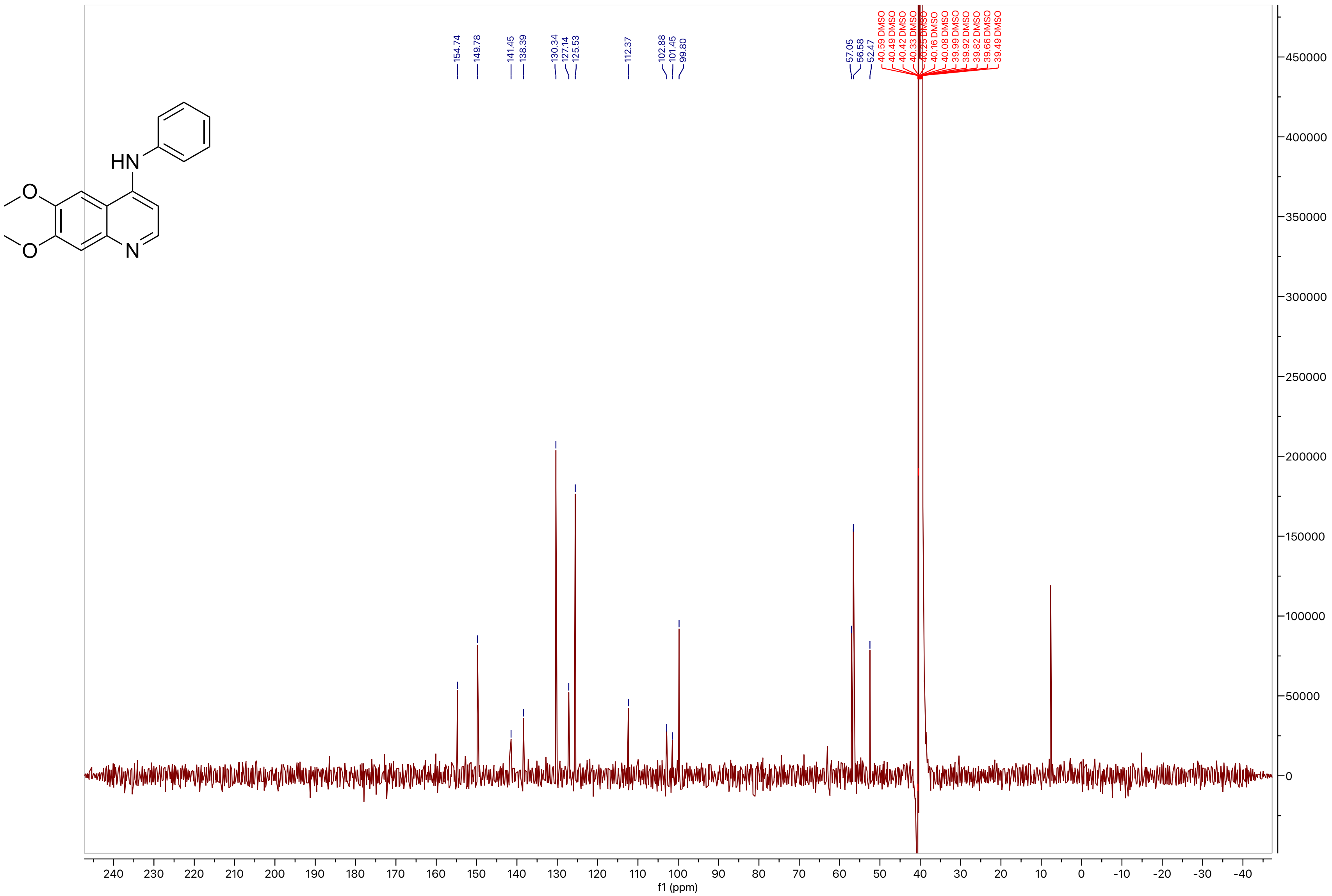
HN
O
O
N
154.74
149.78
141.45
138.39
130.34
127.14
125.53
112.37
102.88
101.45
99.80
57.05
56.58
52.47
40.59 DMSO
40.49 DMSO
40.42 DMSO
40.33 DMSO
40.25 DMSO
40.16 DMSO
40.08 DMSO
39.99 DMSO
39.92 DMSO
39.82 DMSO
39.66 DMSO
39.49 DMSO
f1 (ppm)

#### Compound 42

LCMS

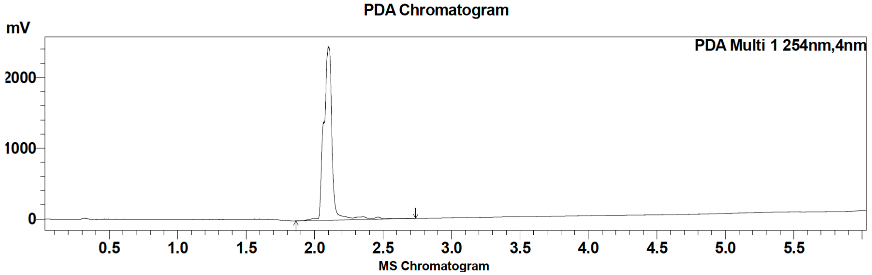

| Peak # | Ret. time | Area | Area % |
|---|---|---|---|
| 1 | 2.1 | 2438959 | 100.00 |

HRMS

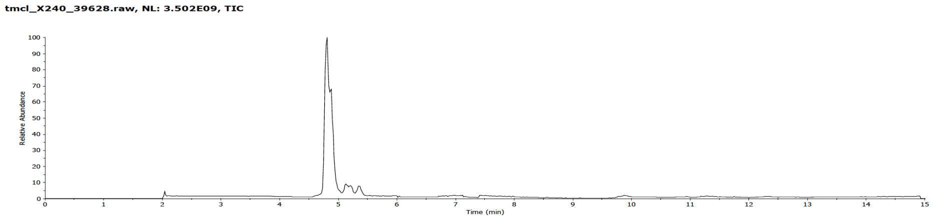

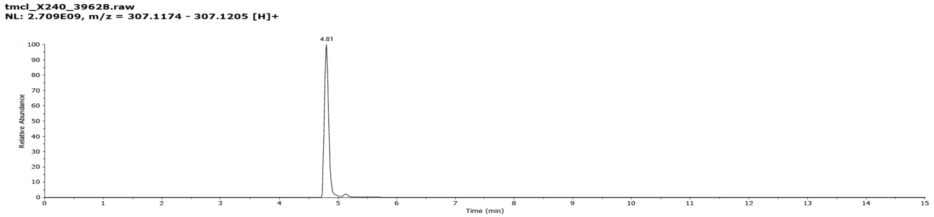

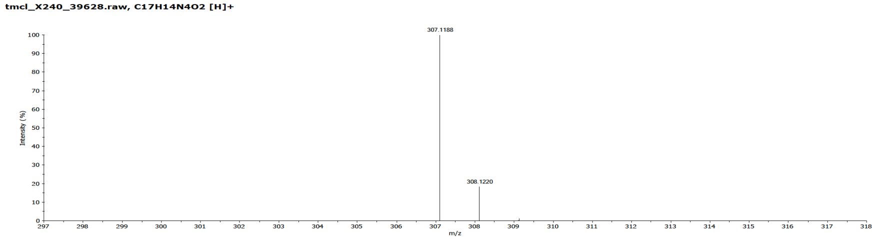

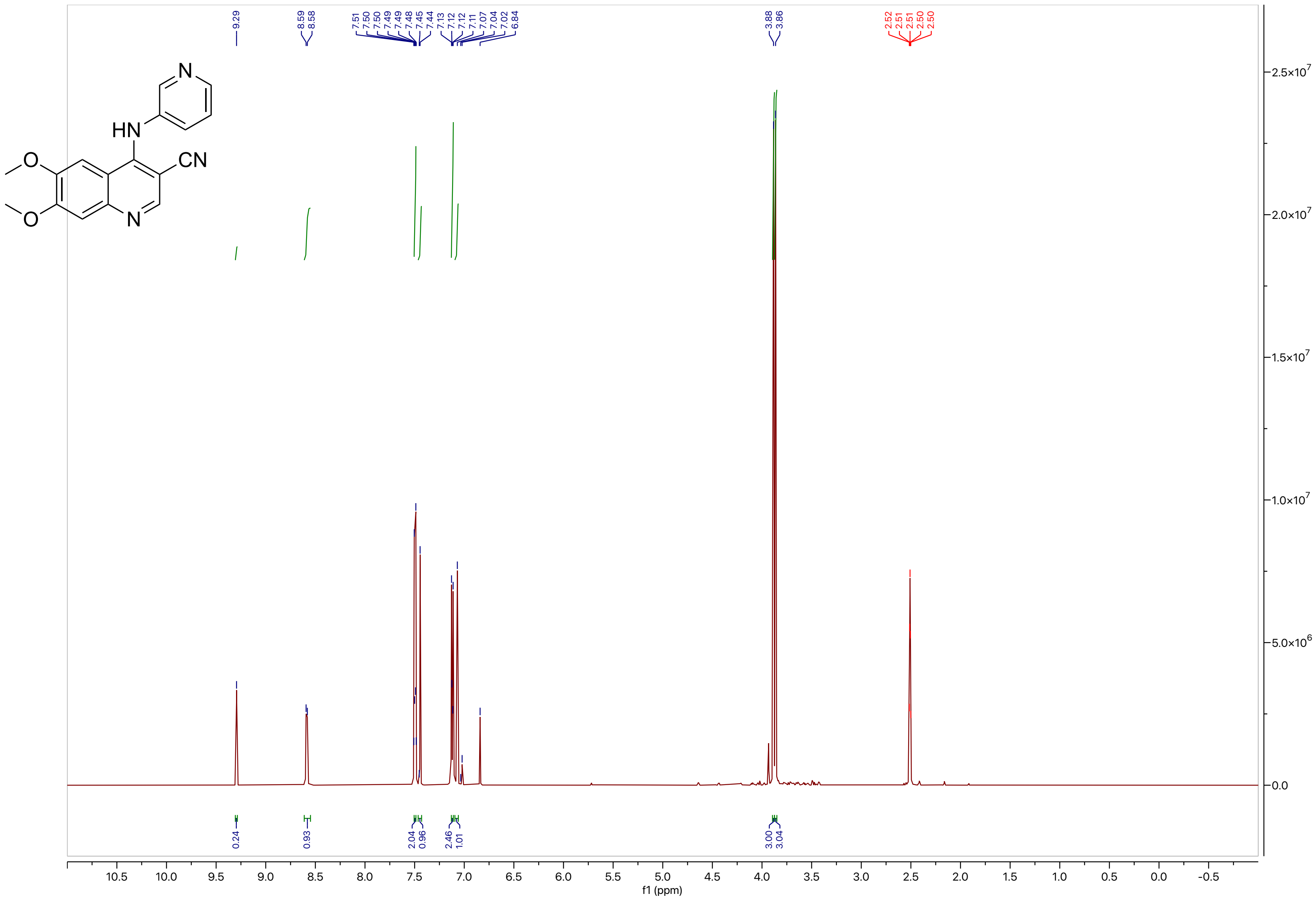
9.29
8.59
8.58
7.51 7.50 7.50 7.49 7.49 7.48 7.45 7.44 7.13 7.12 7.12 7.11 7.07 7.04 7.02 6.84
3.88
3.86
2.52 2.51 2.51 2.50 2.50
0.24
0.93
2.04
0.96
2.46
1.01
3.00
3.04
f1 (ppm)

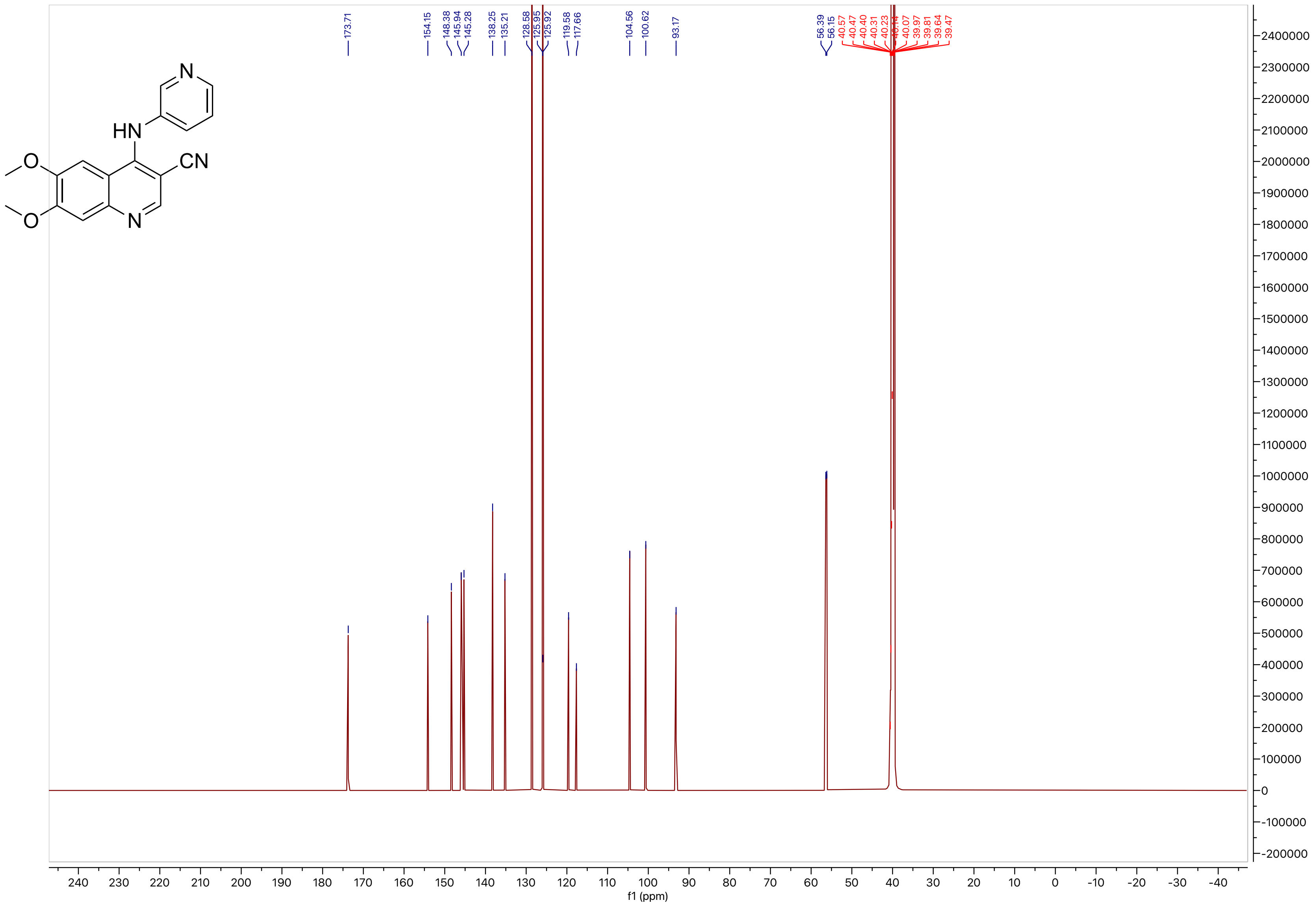
HN
CN
N
O
O
173.71
154.15
148.38
145.94
145.28
138.25
135.21
128.58
125.95
125.92
119.58
117.66
104.56
100.62
93.17
56.39
56.15
40.57
40.47
40.40
40.31
40.23
40.14
40.07
39.97
39.81
39.64
39.47
f1 (ppm)

#### Compound 43

LCMS

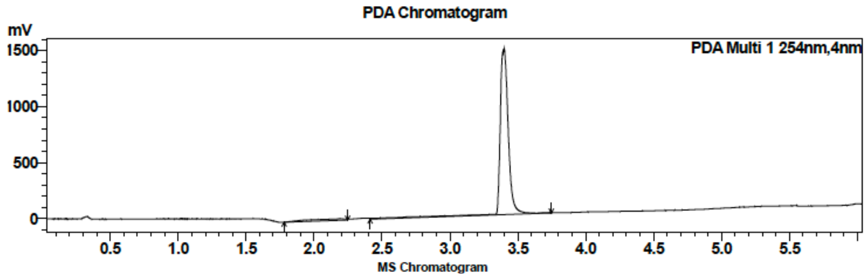

| Peak # | Ret. time | Area | Area % |
|---|---|---|---|
| 1 | 3.4 | 6314337 | 100.00 |

HRMS

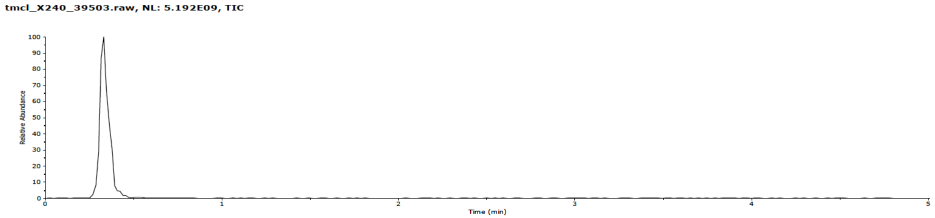

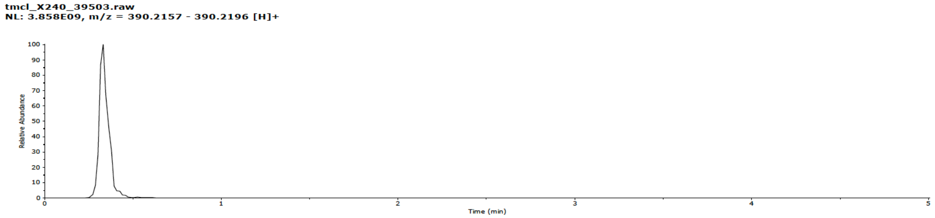

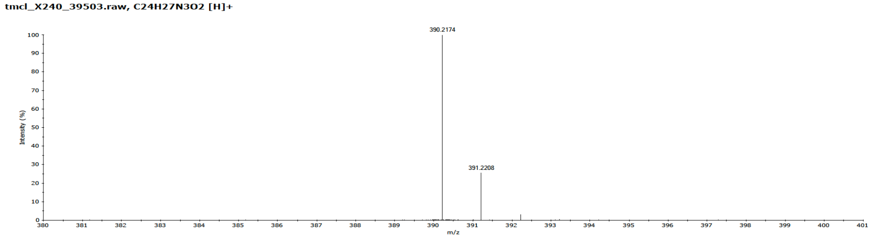

#### Compound 43

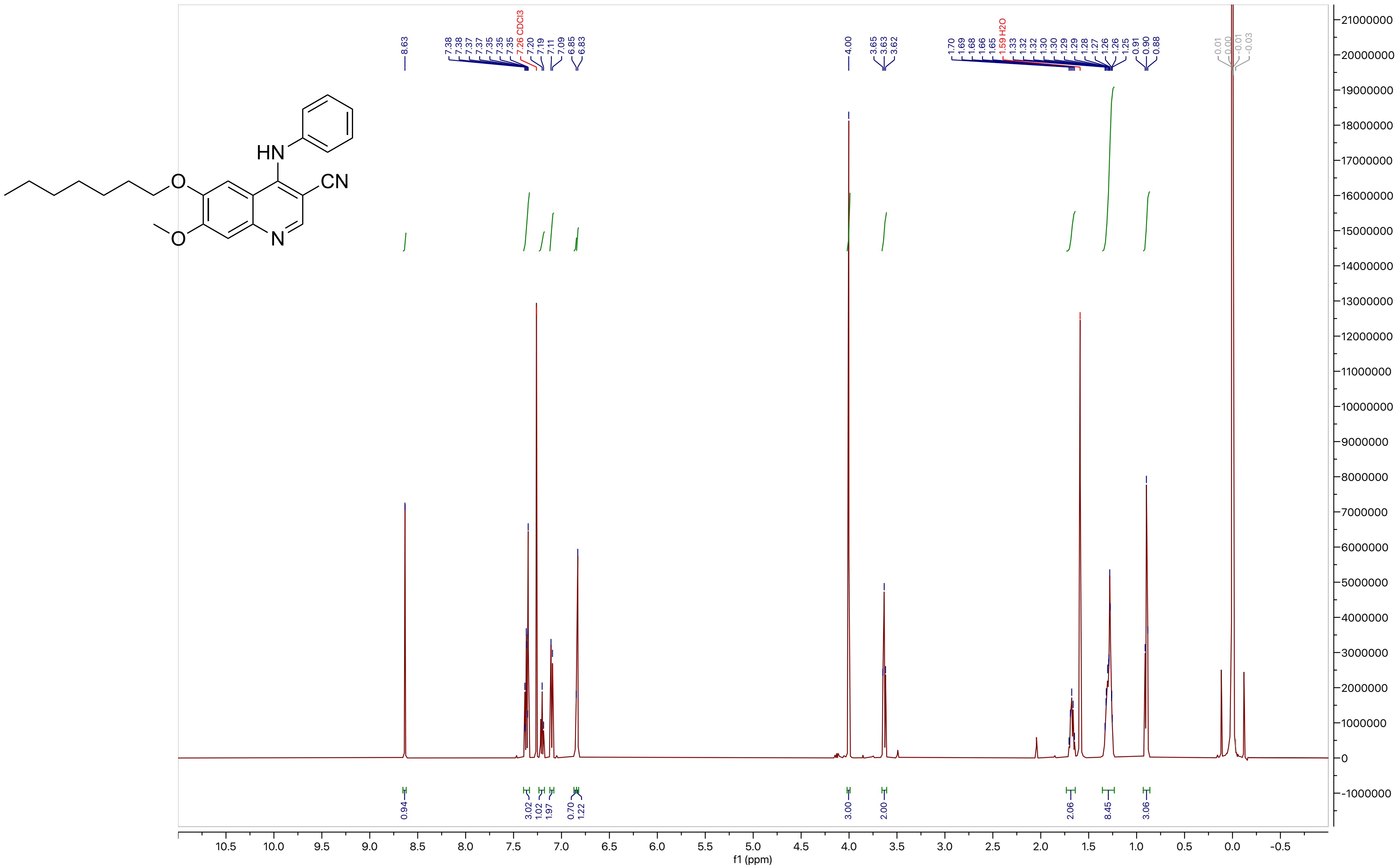

**Compound 43**

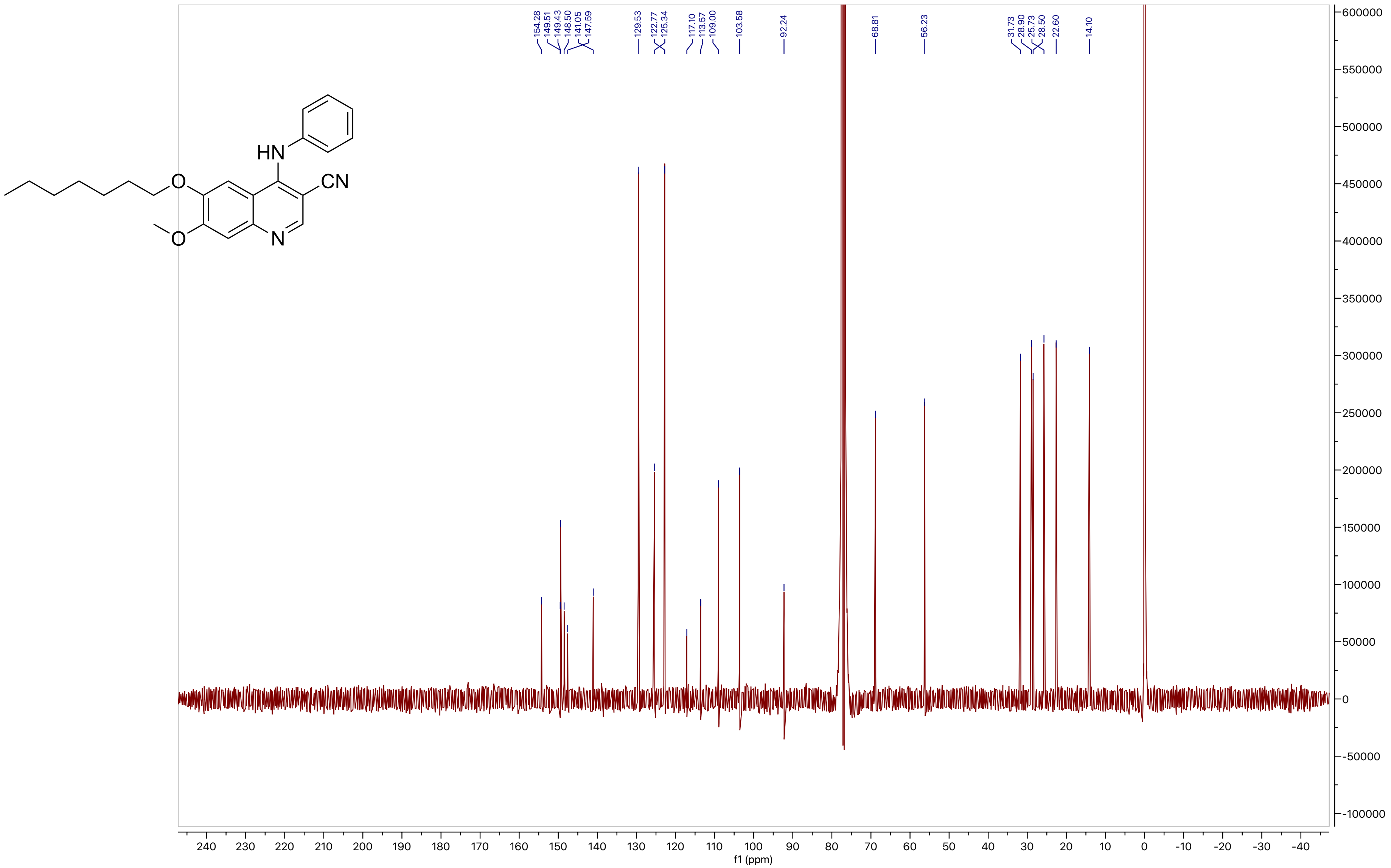
154.28
149.51
149.43
148.50
141.05
147.59
129.53
122.77
125.34
117.10
113.57
109.00
103.58
92.24
68.81
56.23
31.73
28.90
25.73
28.50
22.60
14.10
f1 (ppm)

**Compound 44**

LCMS

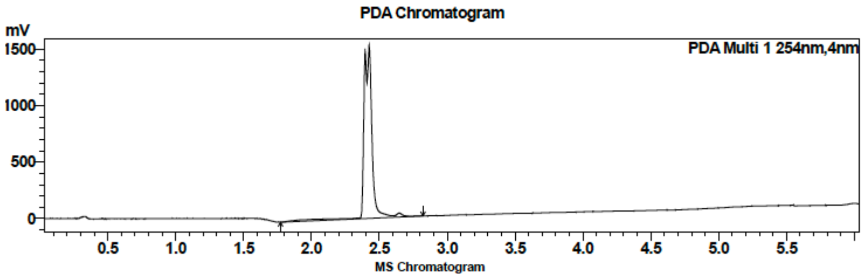

| Peak # | Ret. time | Area | Area % |
|---|---|---|---|
| 1 | 2.4 | 6269126 | 100.00 |

HRMS

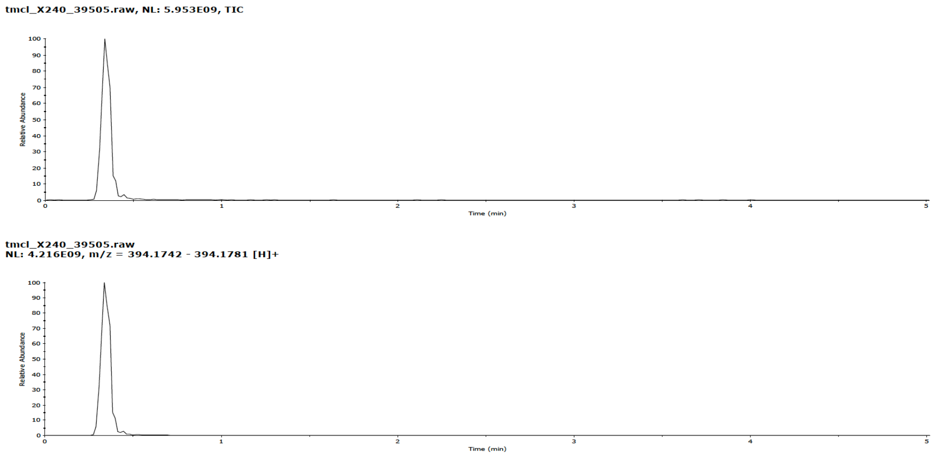

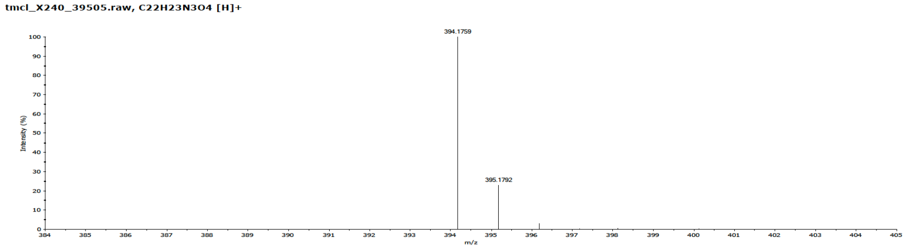

**Compound 44**

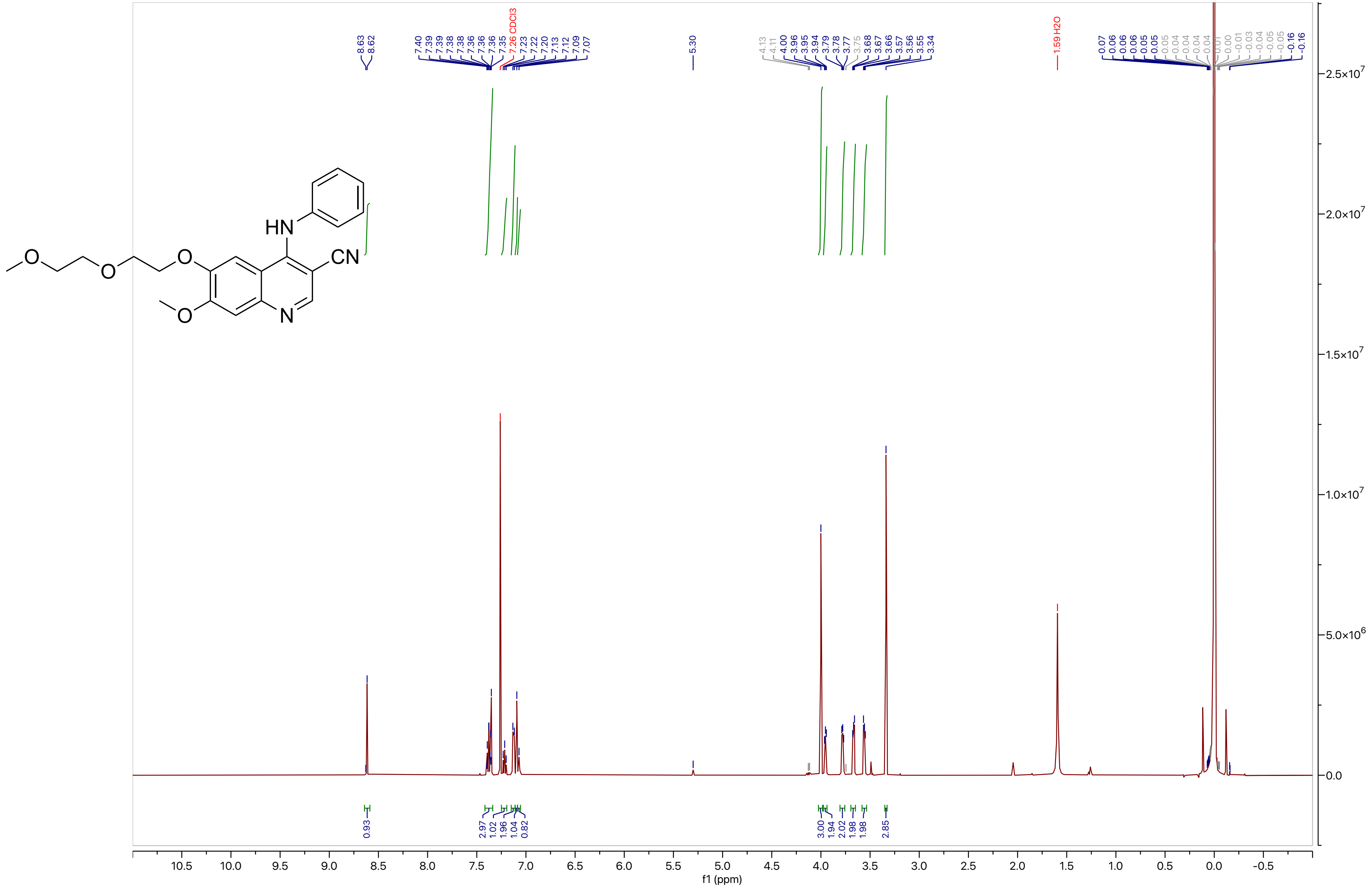

**Compound 44**

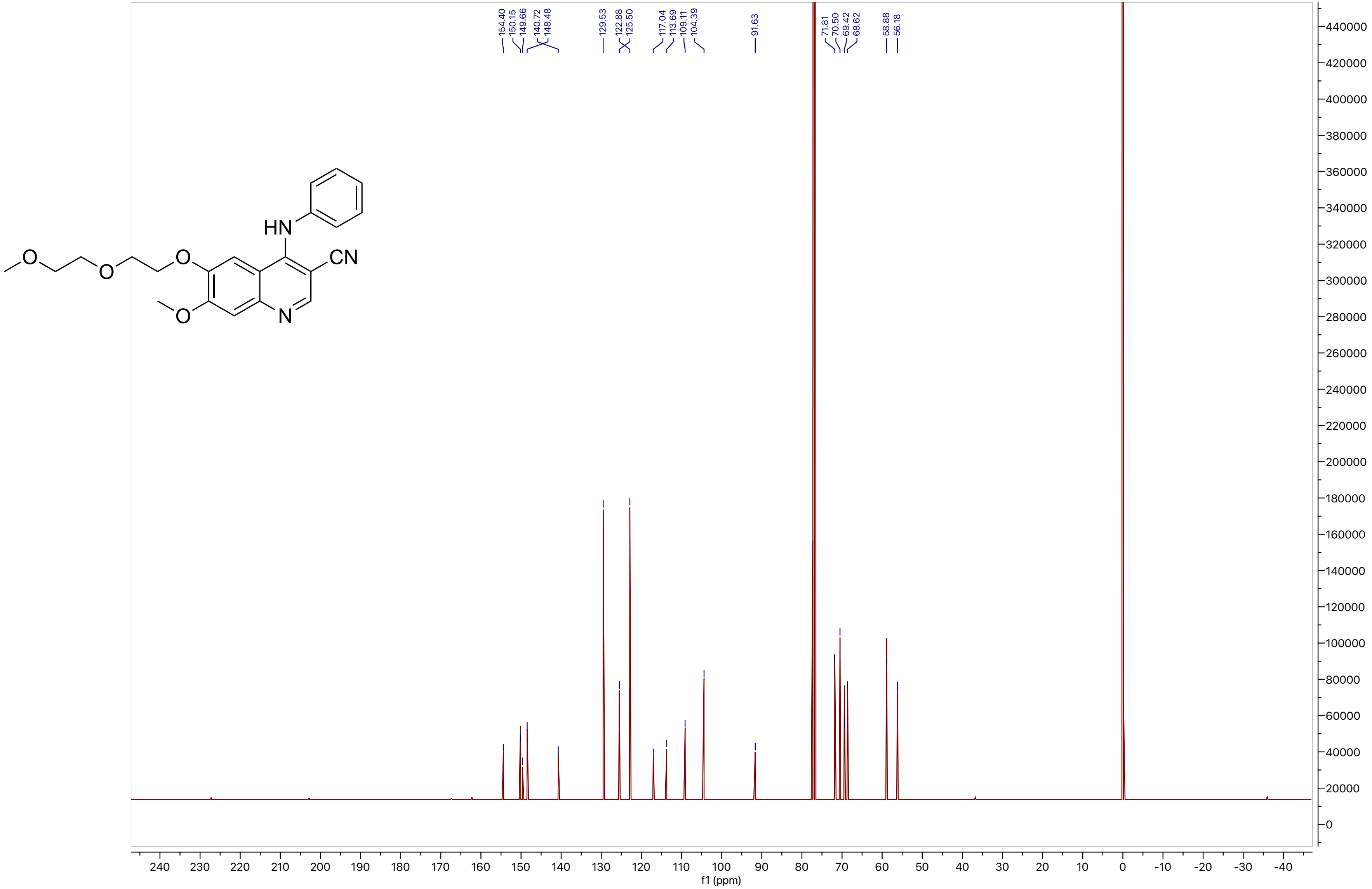

154.40
150.15
149.66
140.72
148.48
129.53
122.88
125.50
117.04
113.69
109.11
104.39
91.63
71.81
70.50
69.42
68.62
58.88
56.18
HN
O
O
O
O
CN
N
f1 (ppm)

**Compound 45**

LCMS

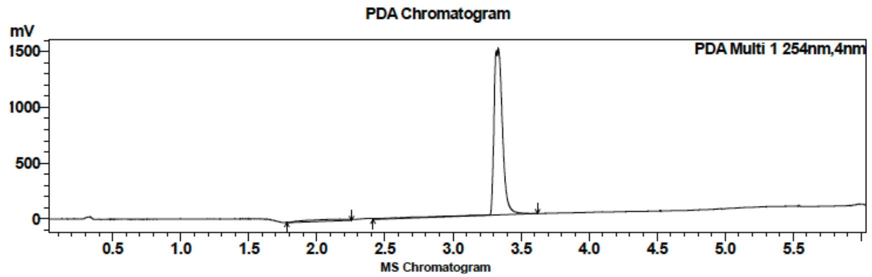

| Peak # | Ret. time | Area | Area % |
|---|---|---|---|
| 1 | 2.2 | 322343 | 4.62 |
| 2 | 3.3 | 6658261 | 95.38 |

HRMS

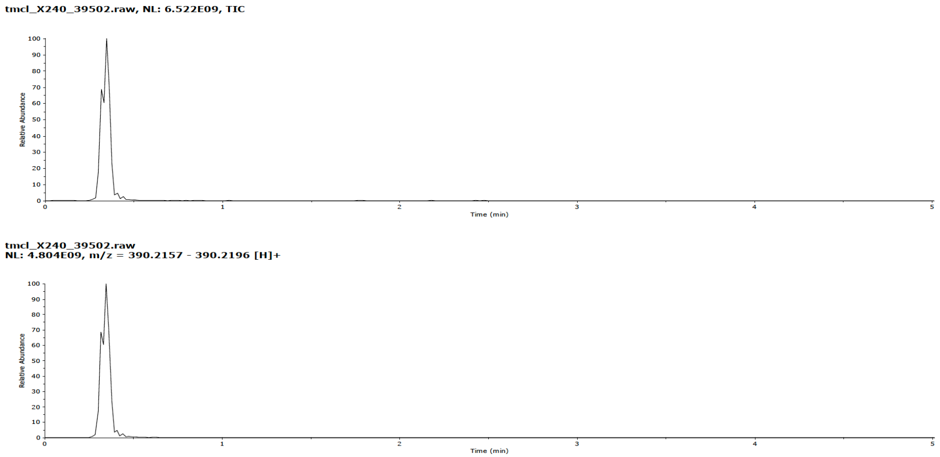

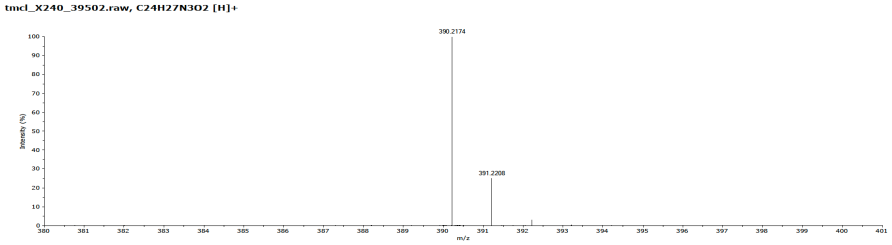

### Compound 45

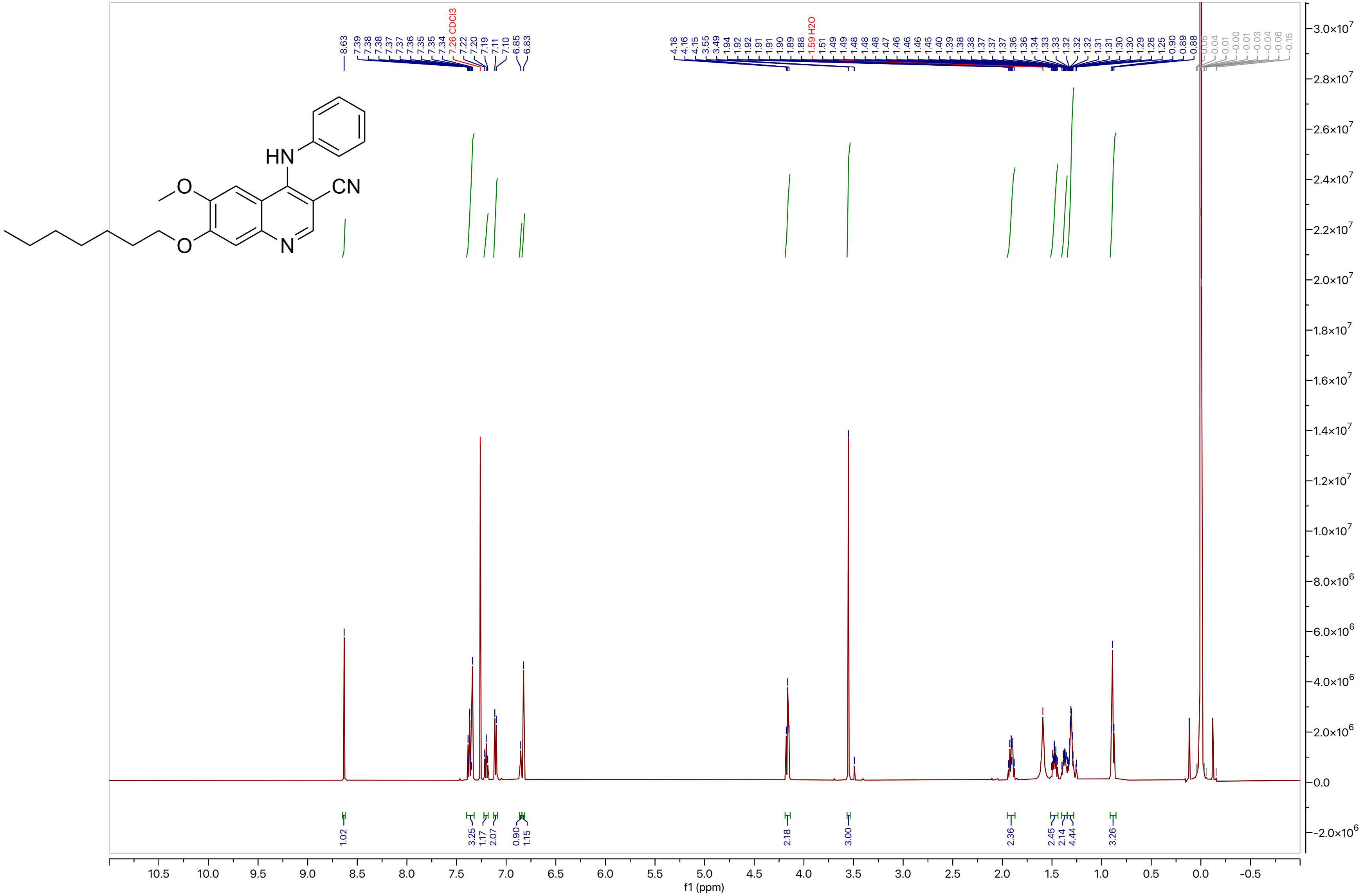

### Compound 45

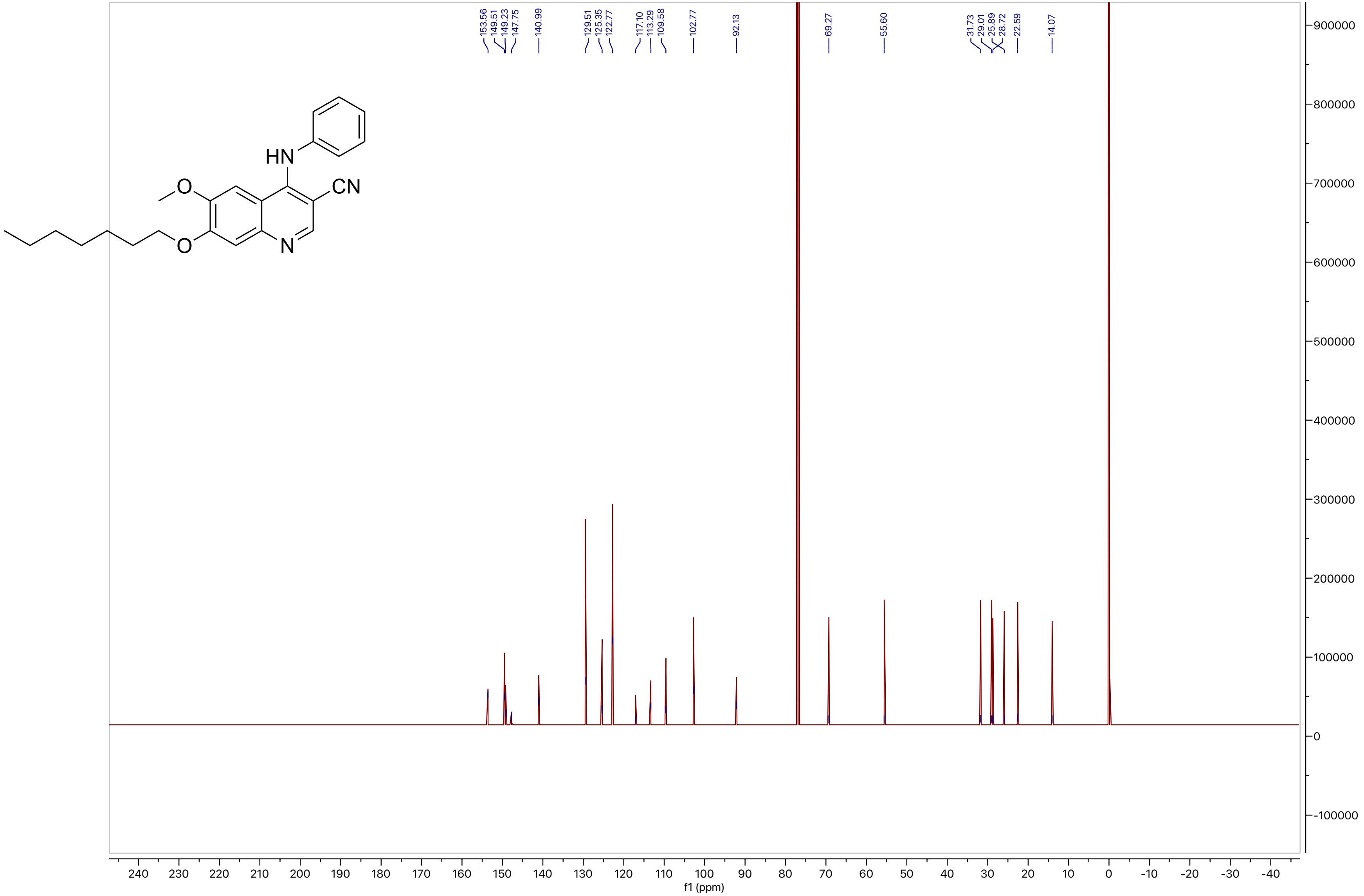
153.56
149.51
149.23
147.75
140.99
129.51
125.35
122.77
117.10
113.29
109.58
102.77
92.13
69.27
55.60
31.73
29.01
25.89
28.72
22.59
14.07
HN
O
CN
O
N
f1 (ppm)

**Compound 46**

LCMS

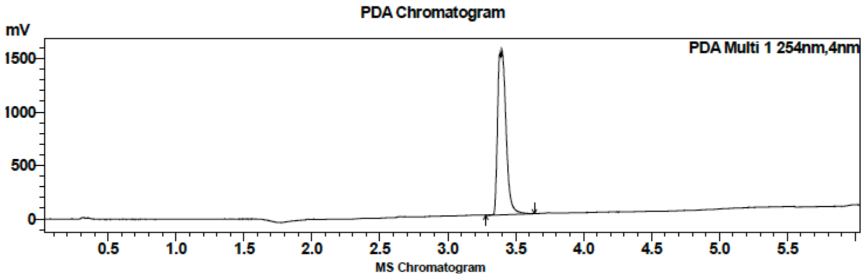

| Peak # | Ret. time | Area | Area % |
|---|---|---|---|
| 1 | 3.4 | 6642847 | 100.00 |

HRMS

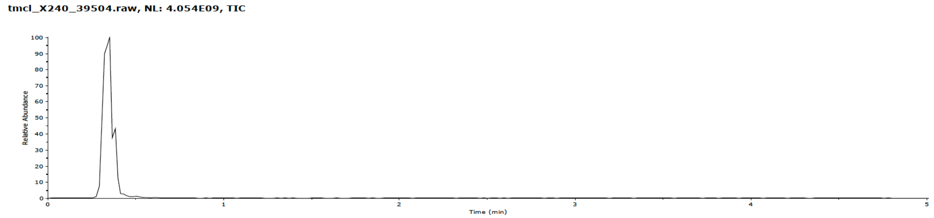

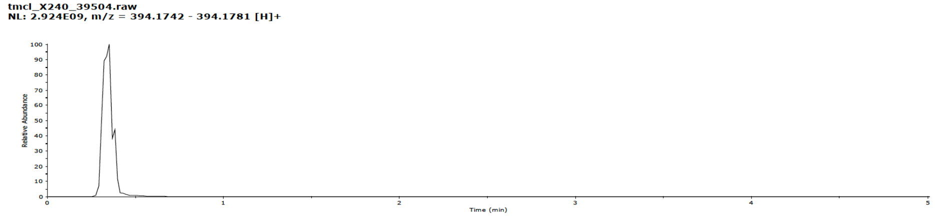

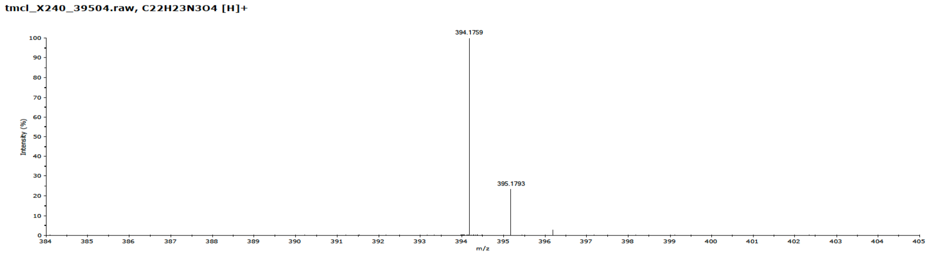

**Compound 46**

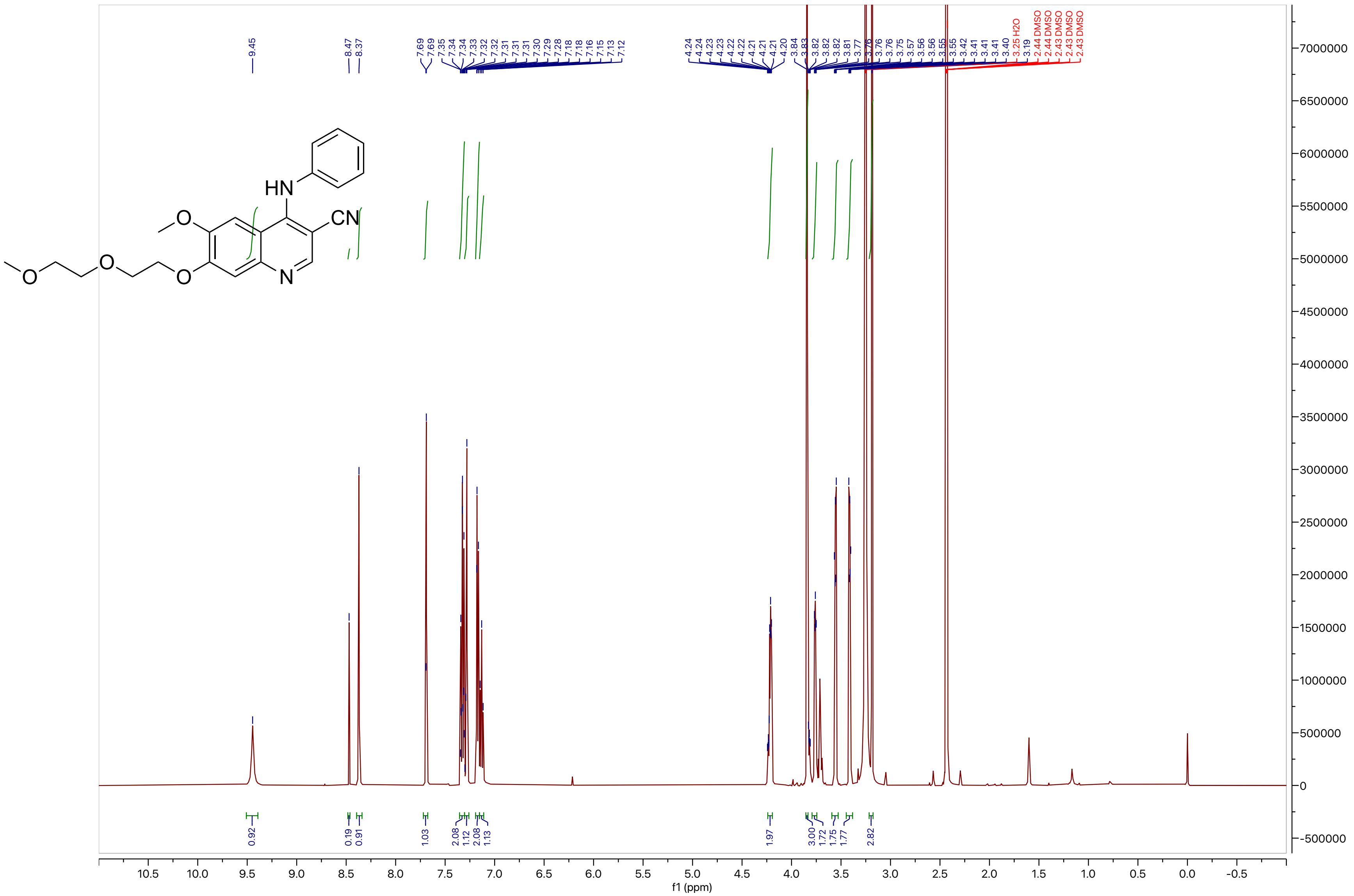
HN
CN
O
O
O
O
N
9.45
8.47
8.37
7.69
7.69
7.35
7.34
7.34
7.33
7.32
7.32
7.31
7.31
7.31
7.30
7.29
7.28
7.18
7.16
7.15
7.13
7.12
4.24
4.24
4.23
4.23
4.22
4.22
4.21
4.21
4.21
4.20
3.84
3.83
3.82
3.82
3.82
3.81
3.77
3.76
3.76
3.75
3.57
3.56
3.56
3.55
3.55
3.42
3.41
3.41
3.41
3.40
3.25 H2O
3.19
2.44 DMSO
2.44 DMSO
2.43 DMSO
2.43 DMSO
2.43 DMSO
0.92
0.19
0.91
1.03
2.08
1.12
2.08
1.13
1.97
3.00
1.72
1.75
1.77
2.82
f1 (ppm)

**Compound 46**

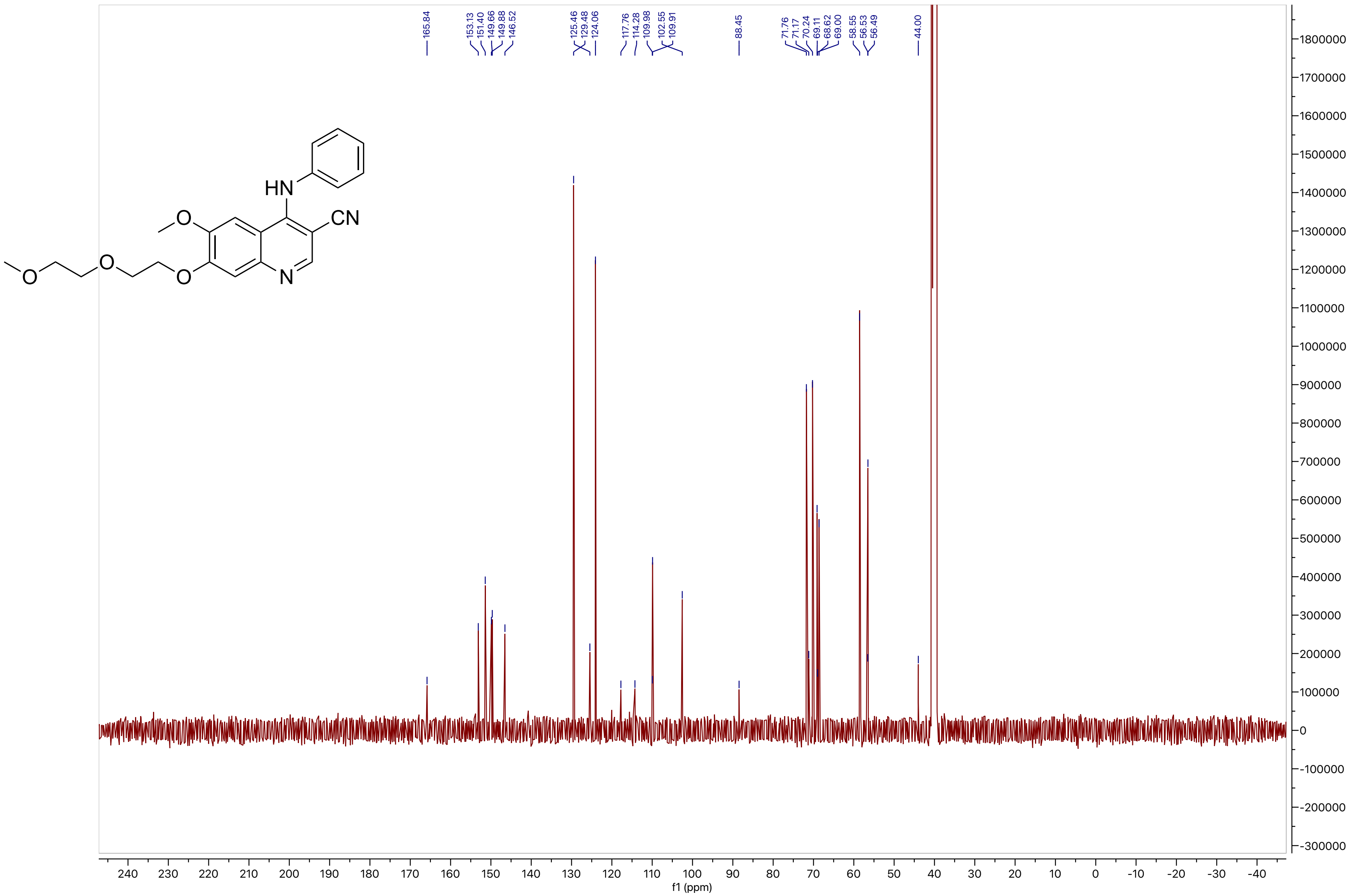

165.84
153.13
151.40
149.66
149.88
146.52
125.46
129.48
124.06
117.76
114.28
109.98
102.55
109.91
88.45
71.76
71.17
70.24
69.11
68.62
69.00
58.55
56.53
56.49
44.00
HN
O
CN
O
O
O
N
f1 (ppm)

**Compound 47**

LCMS

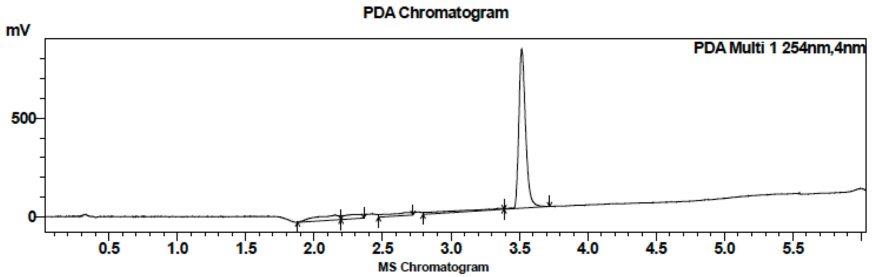

| Peak # | Ret. time | Area | Area % |
|---|---|---|---|
| 1 | 2.2 | 115136 | 3.98 |
| 2 | 3.5 | 2775010 | 96.01 |

HRMS

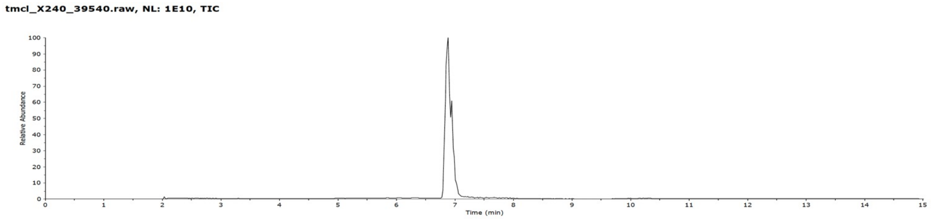

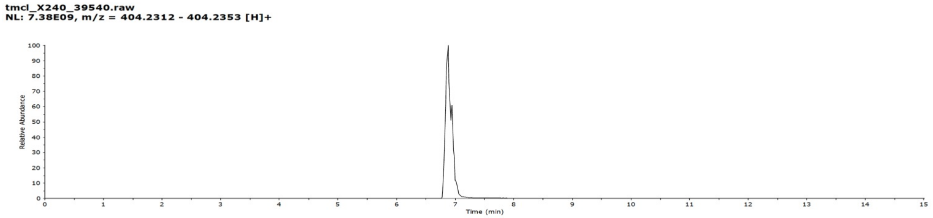

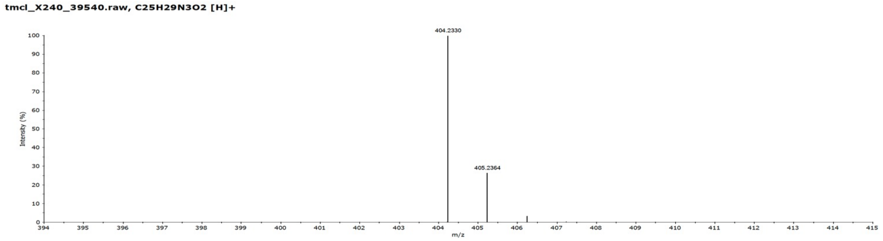

**Compound 47**

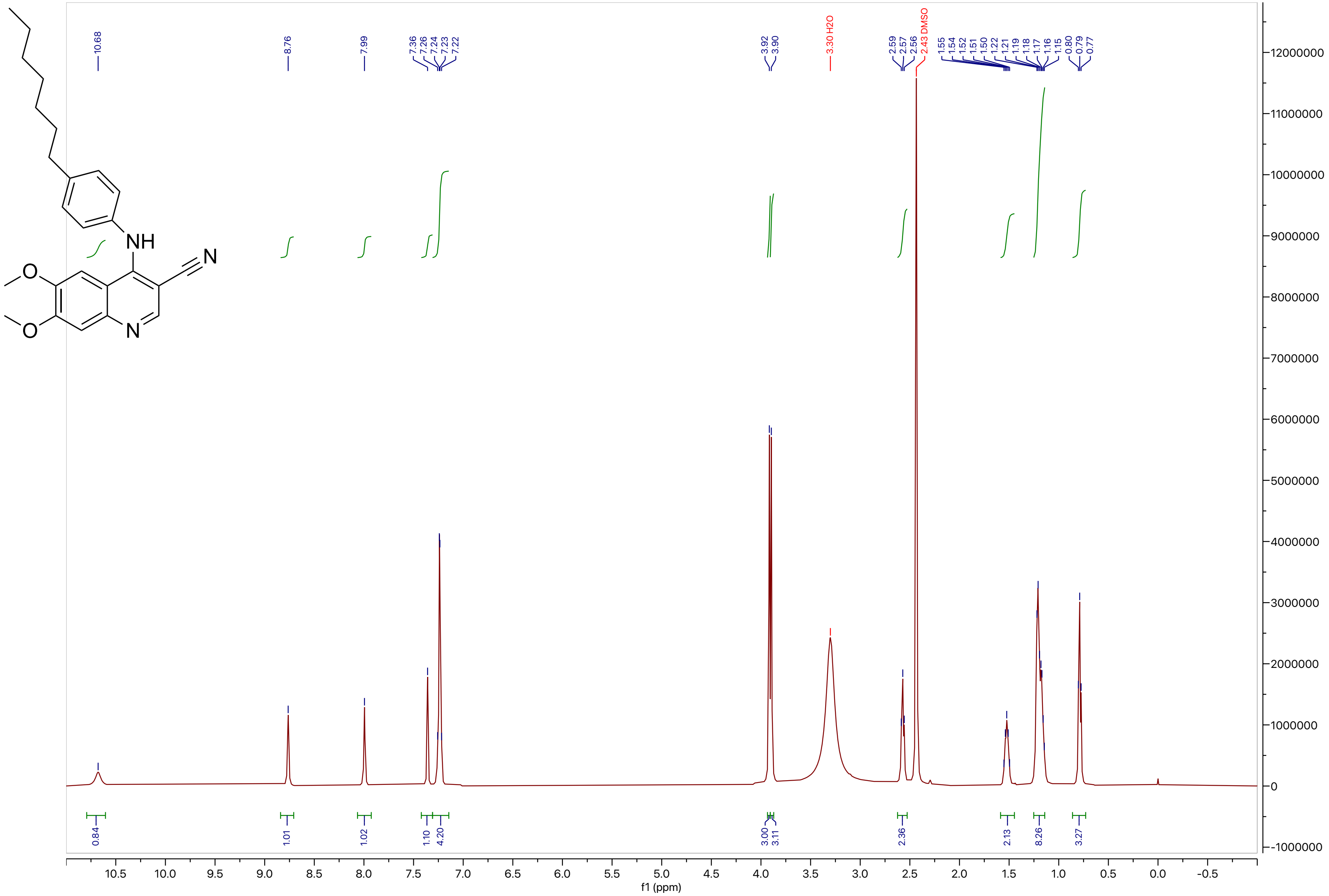

NH
N
O
O
N
10.68
8.76
7.99
7.36
7.26
7.24
7.23
7.22
3.92
3.90
3.30 H2O
2.59
2.57
2.56
2.43 DMSO
1.55
1.54
1.52
1.51
1.50
1.22
1.21
1.19
1.18
1.17
1.16
1.15
0.80
0.79
0.77
0.84
1.01
1.02
1.10
4.20
3.00
3.11
2.36
2.13
8.26
3.27
f1 (ppm)

**Compound 47**

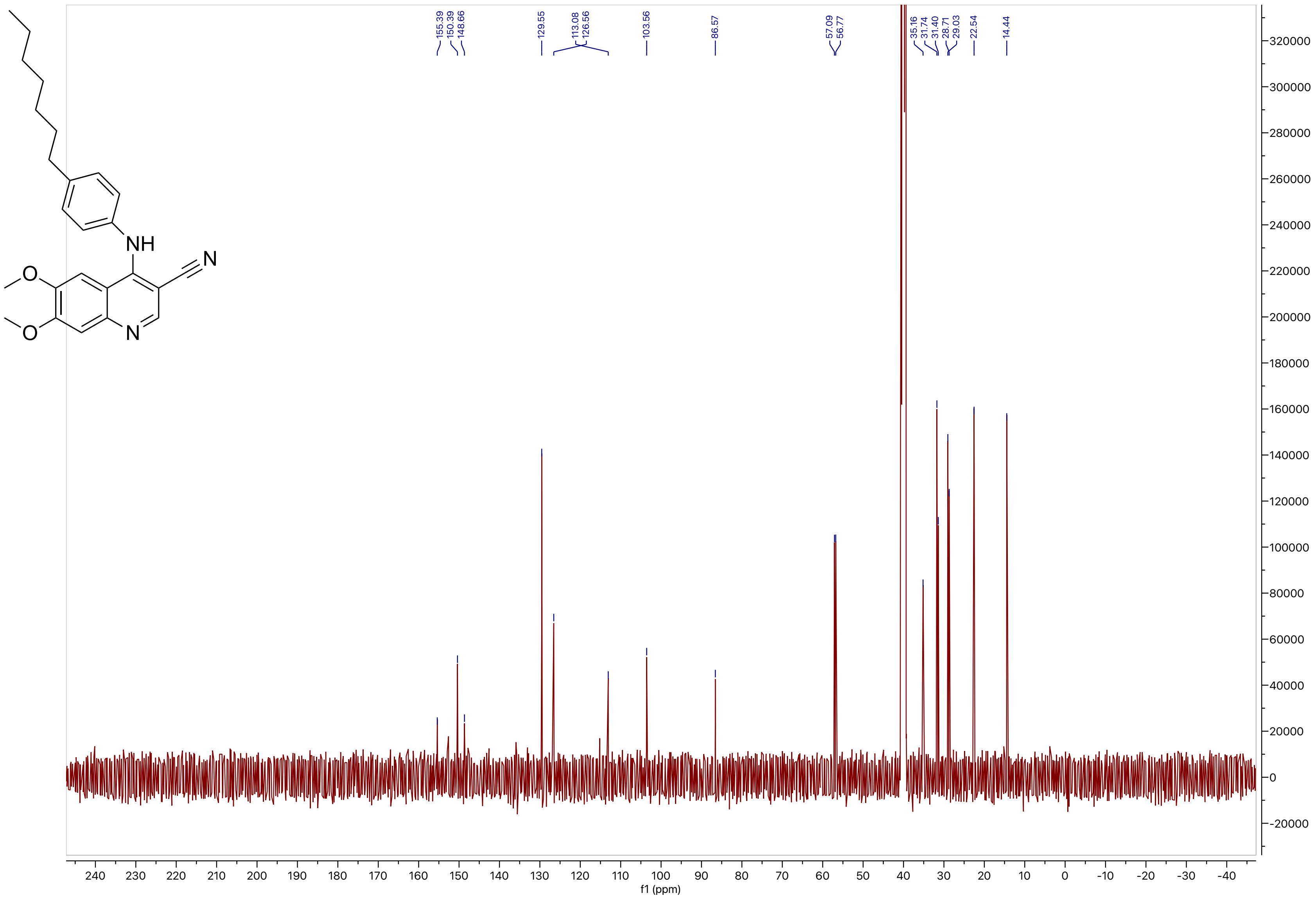
155.39
150.39
148.66
129.55
113.08
126.56
103.56
86.57
57.09
56.77
35.16
31.74
31.40
28.71
29.03
22.54
14.44
f1 (ppm)

#### Compound 48

LCMS

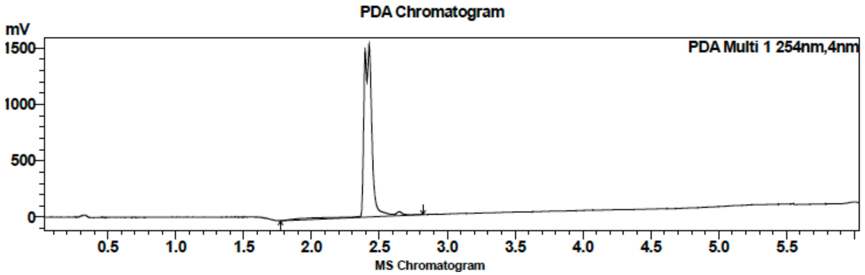

| Peak # | Ret. time | Area | Area % |
|---|---|---|---|
| 1 | 2.4 | 6269126 | 100.00 |

HRMS

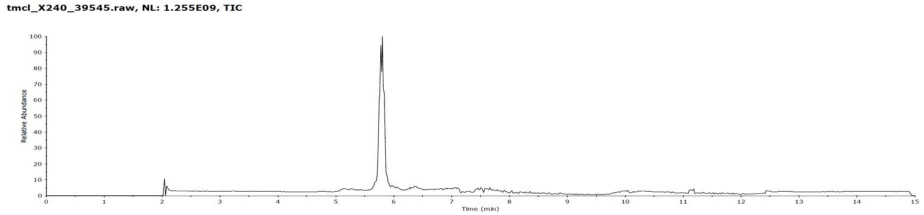

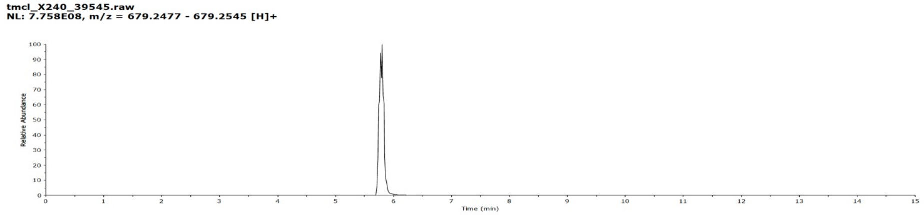

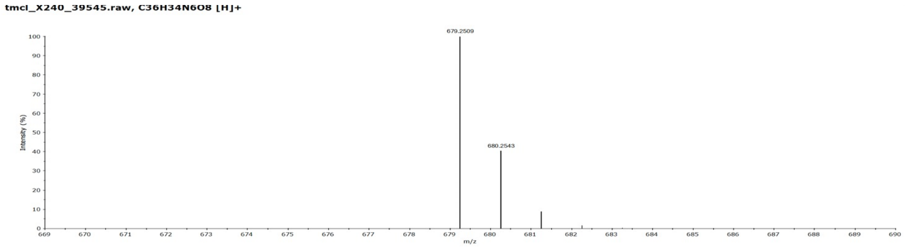

**Compound 48**

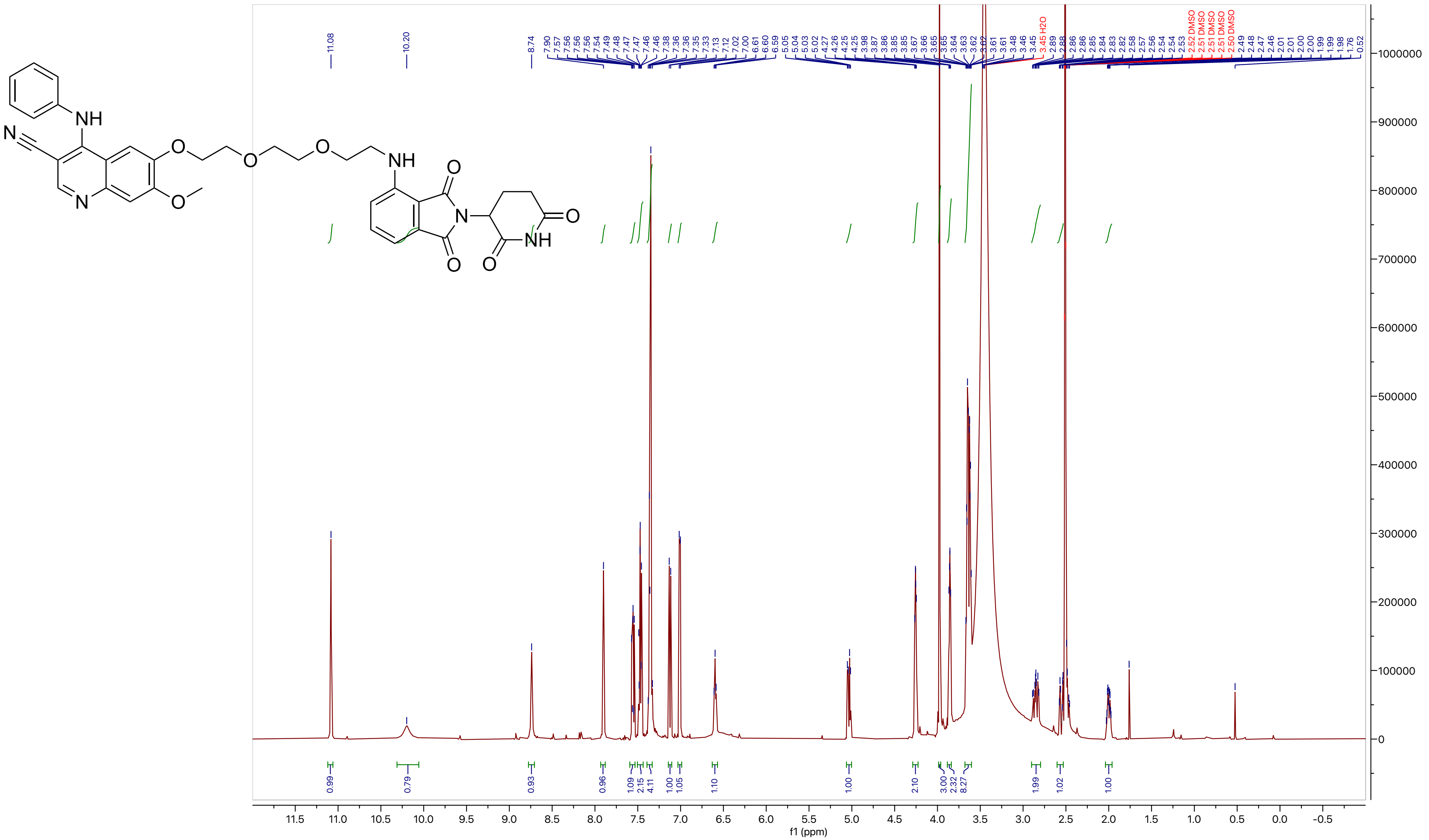

**Compound 48**

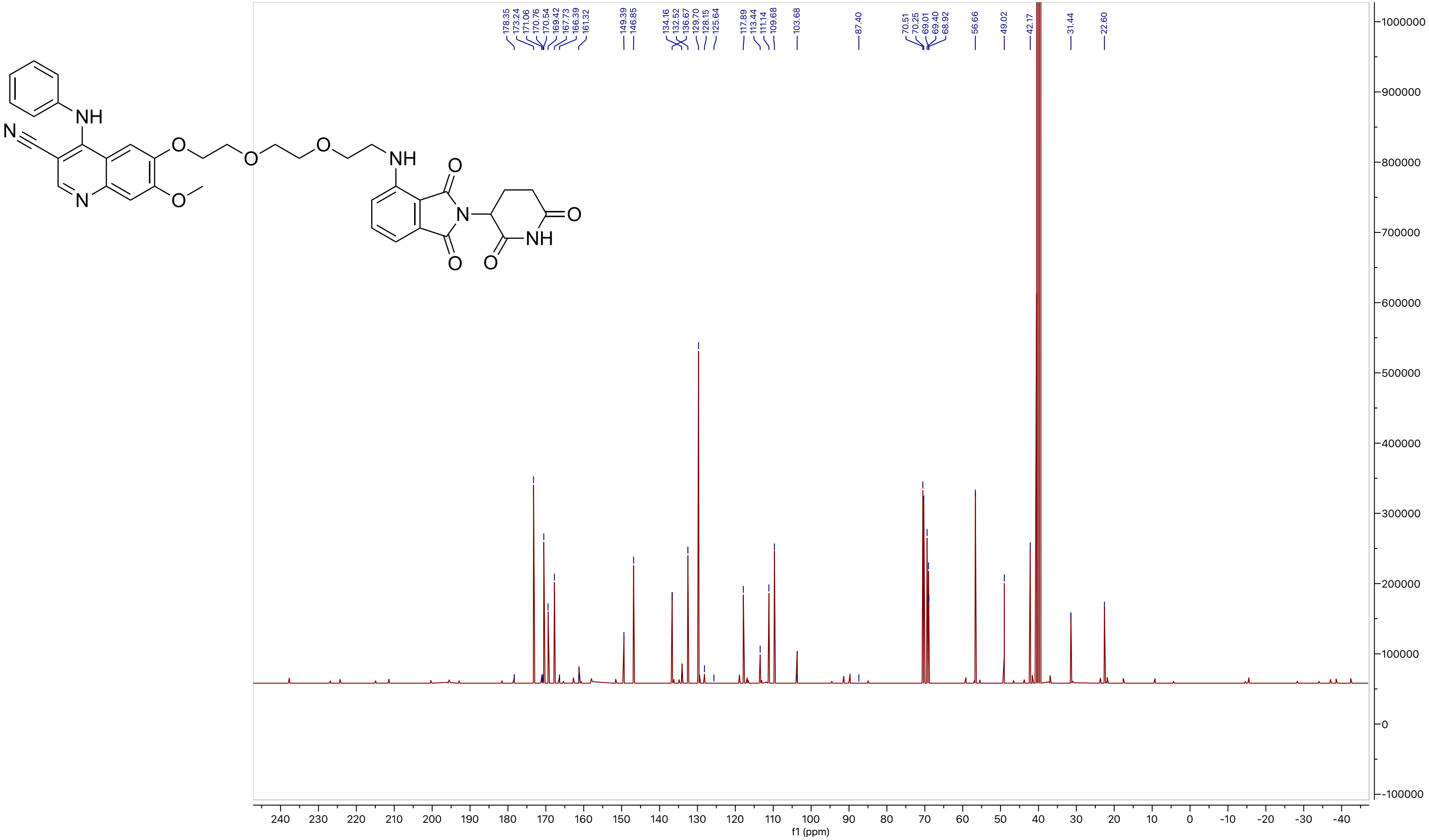

178.35
173.24
171.06
170.76
170.54
169.42
167.73
166.39
161.32
149.39
146.85
134.16
132.52
136.67
129.70
128.15
125.64
117.89
113.44
111.14
109.68
103.68
87.40
70.51
70.25
69.01
69.40
68.92
56.66
49.02
42.17
31.44
22.60
f1 (ppm)

**Compound 49**

LCMS

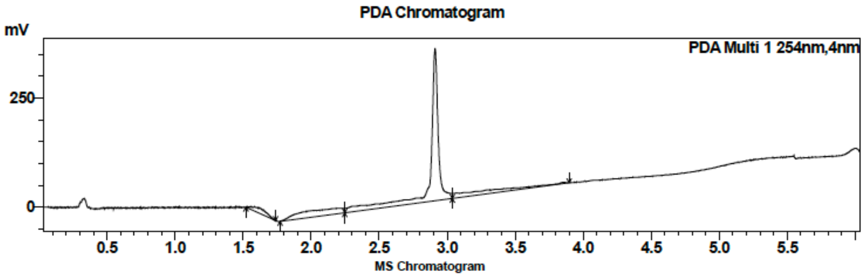

| Peak # | Ret. time | Area | Area % |
|---|---|---|---|
| 1 | 2.9 | 1432057 | 100.00 |

HRMS

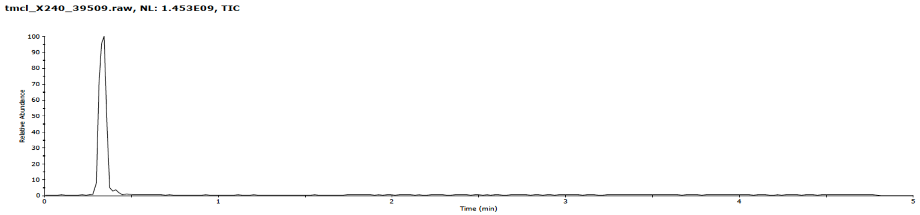

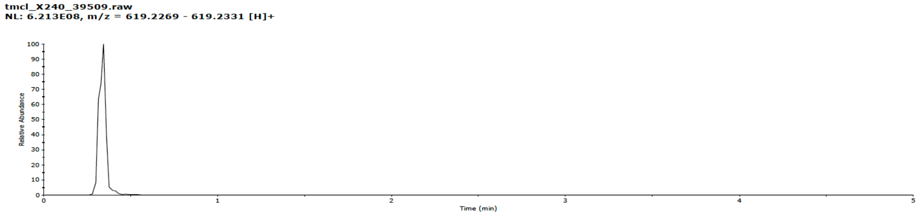

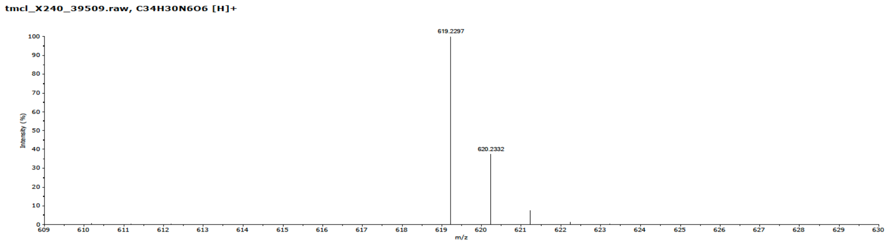

**Compound 49**

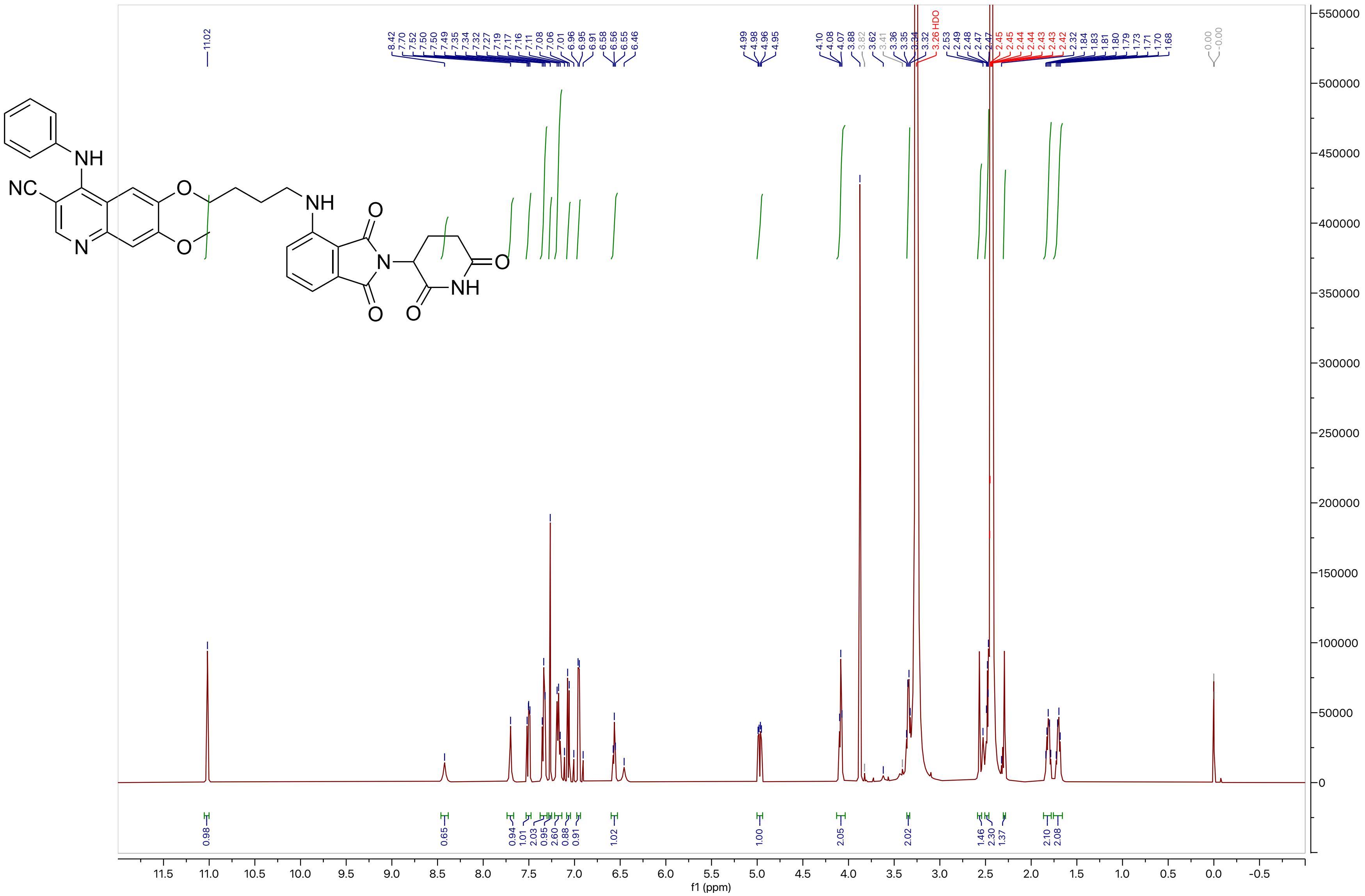

**Compound 49**

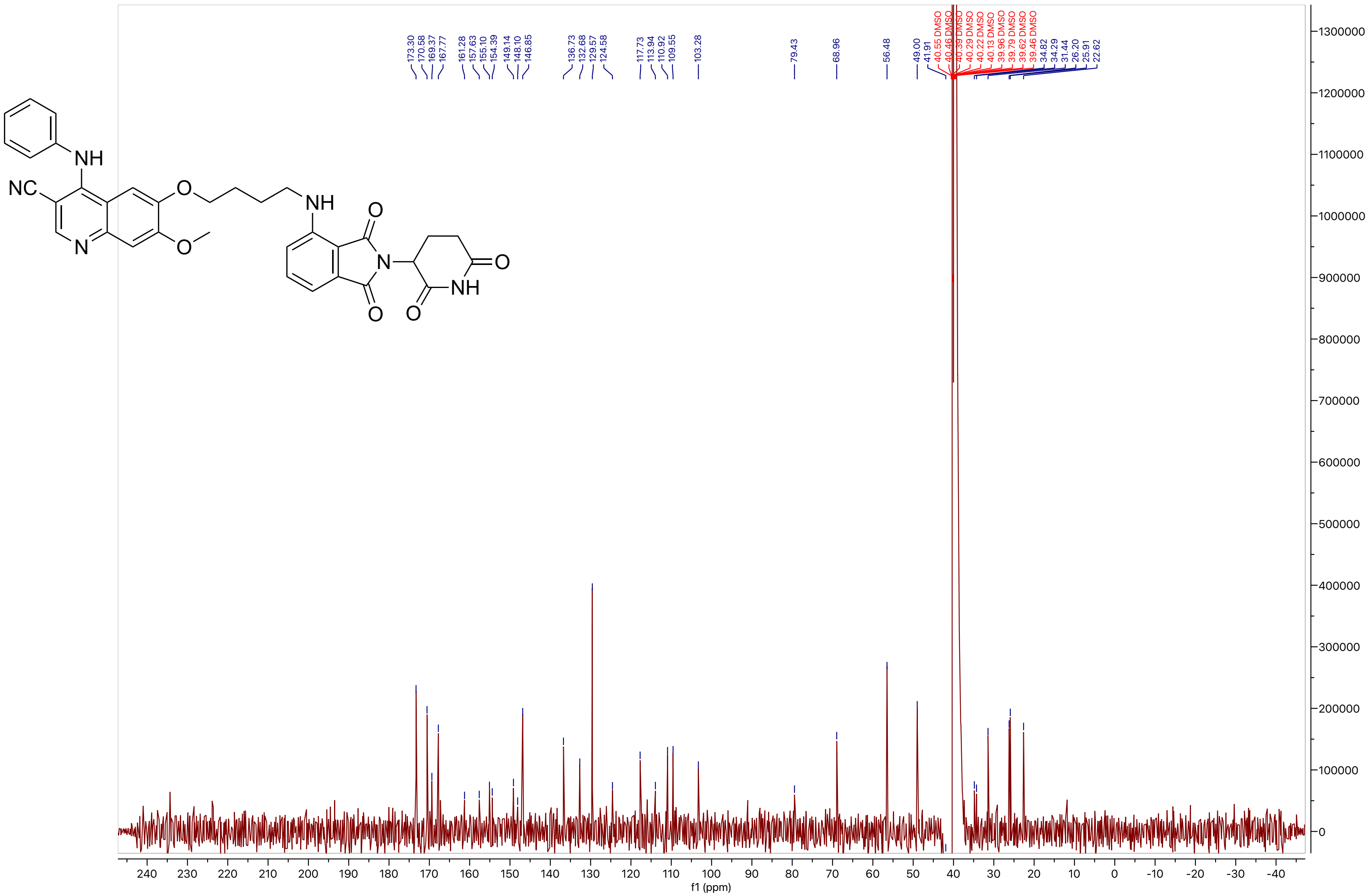

**Compound 50**

LCMS

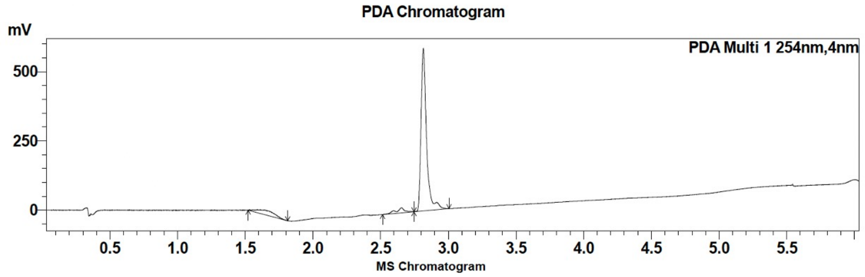

| Peak # | Ret. time | Area | Area % |
|---|---|---|---|
| 1 | 2.7 | 81994 | 4.73 |
| 2 | 2.8 | 1652456 | 95.27 |

HRMS

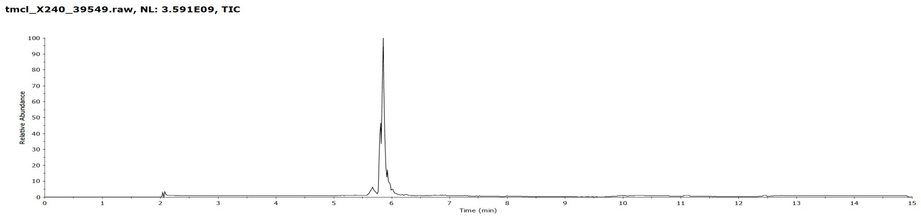

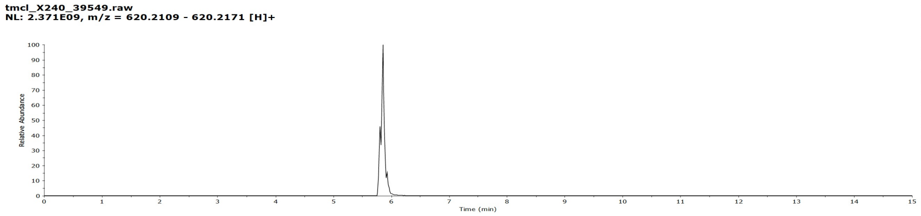

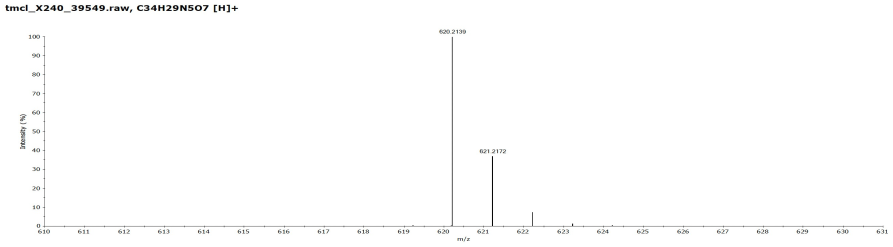

**Compound 50**

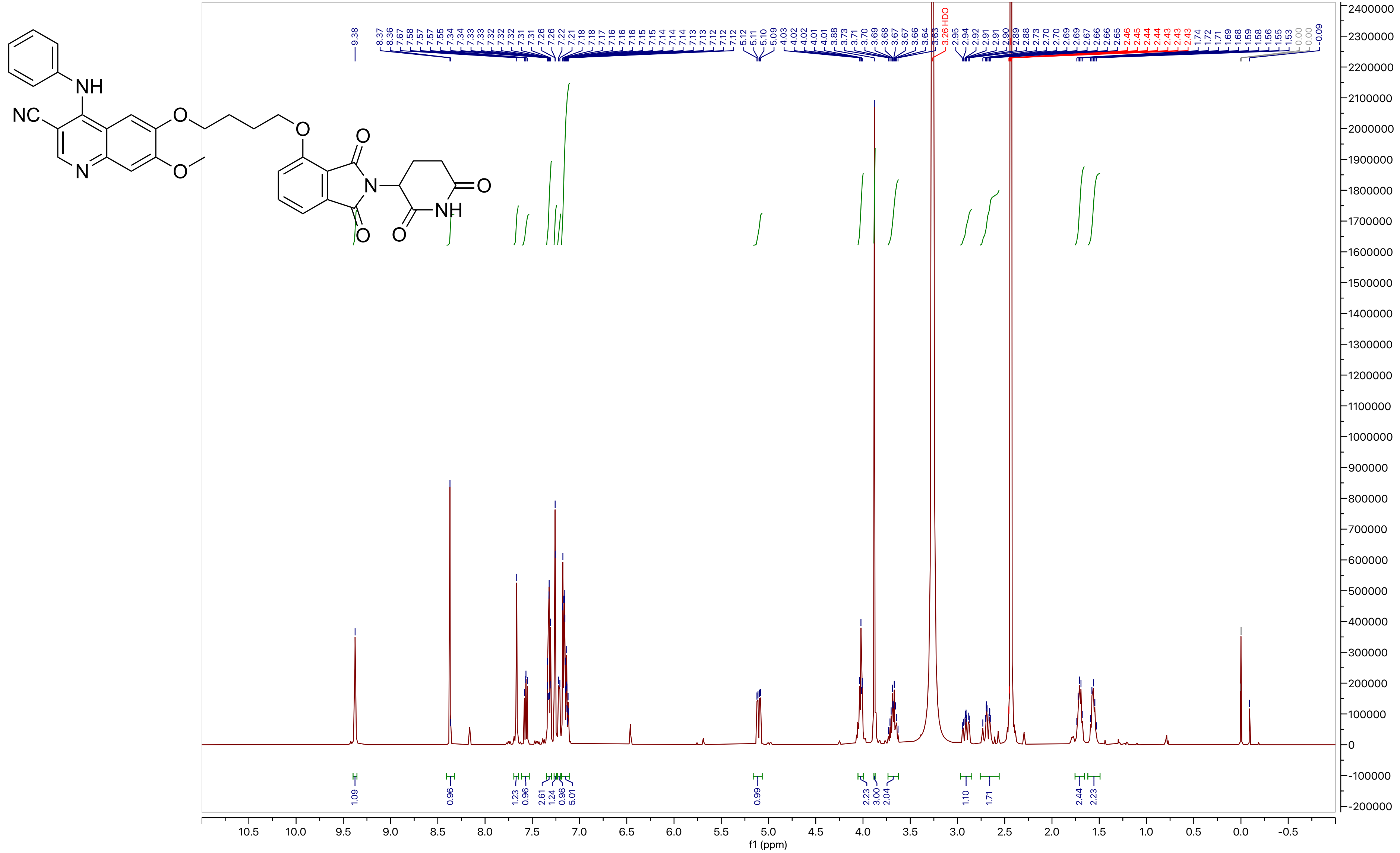

**Compound 50**

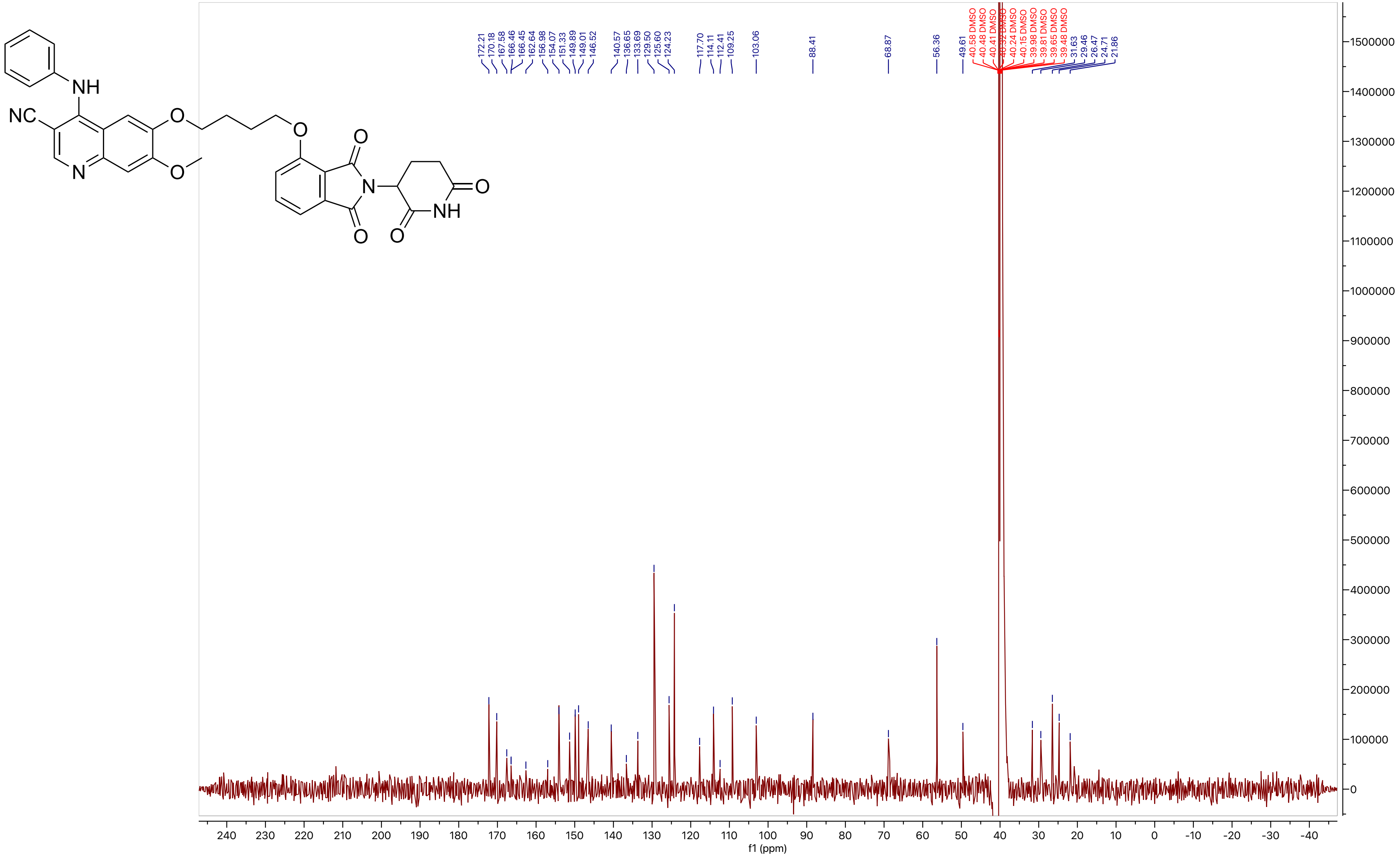

172.21
170.18
167.58
166.46
166.45
162.64
156.98
154.07
151.33
149.89
149.01
146.52
140.57
136.65
133.69
129.50
125.60
124.23
117.70
114.11
112.41
109.25
103.06
88.41
68.87
56.36
49.61
40.58 DMSO
40.48 DMSO
40.41 DMSO
40.32 DMSO
40.24 DMSO
40.15 DMSO
39.98 DMSO
39.81 DMSO
39.65 DMSO
39.48 DMSO
31.63
29.46
26.47
24.71
21.86
f1 (ppm)

#### Compound 51

LCMS

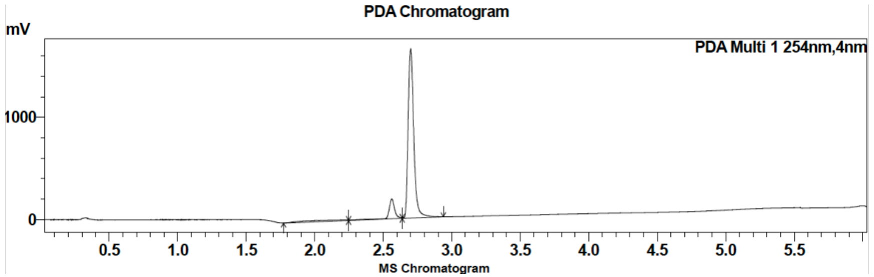

| Peak # | Ret. time | Area | Area % |
|---|---|---|---|
| 1 | 2.6 | 178912 | 3.63 |
| 2 | 2.7 | 4746190 | 96.37 |

HRMS

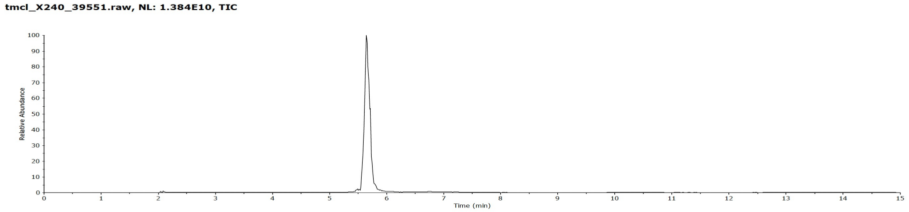

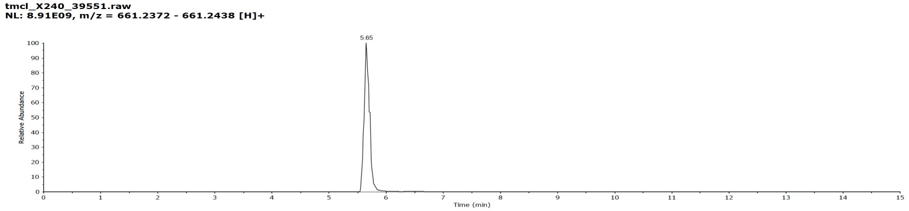

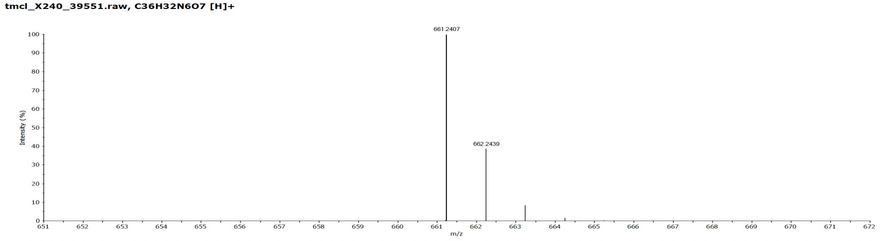

### Compound 51

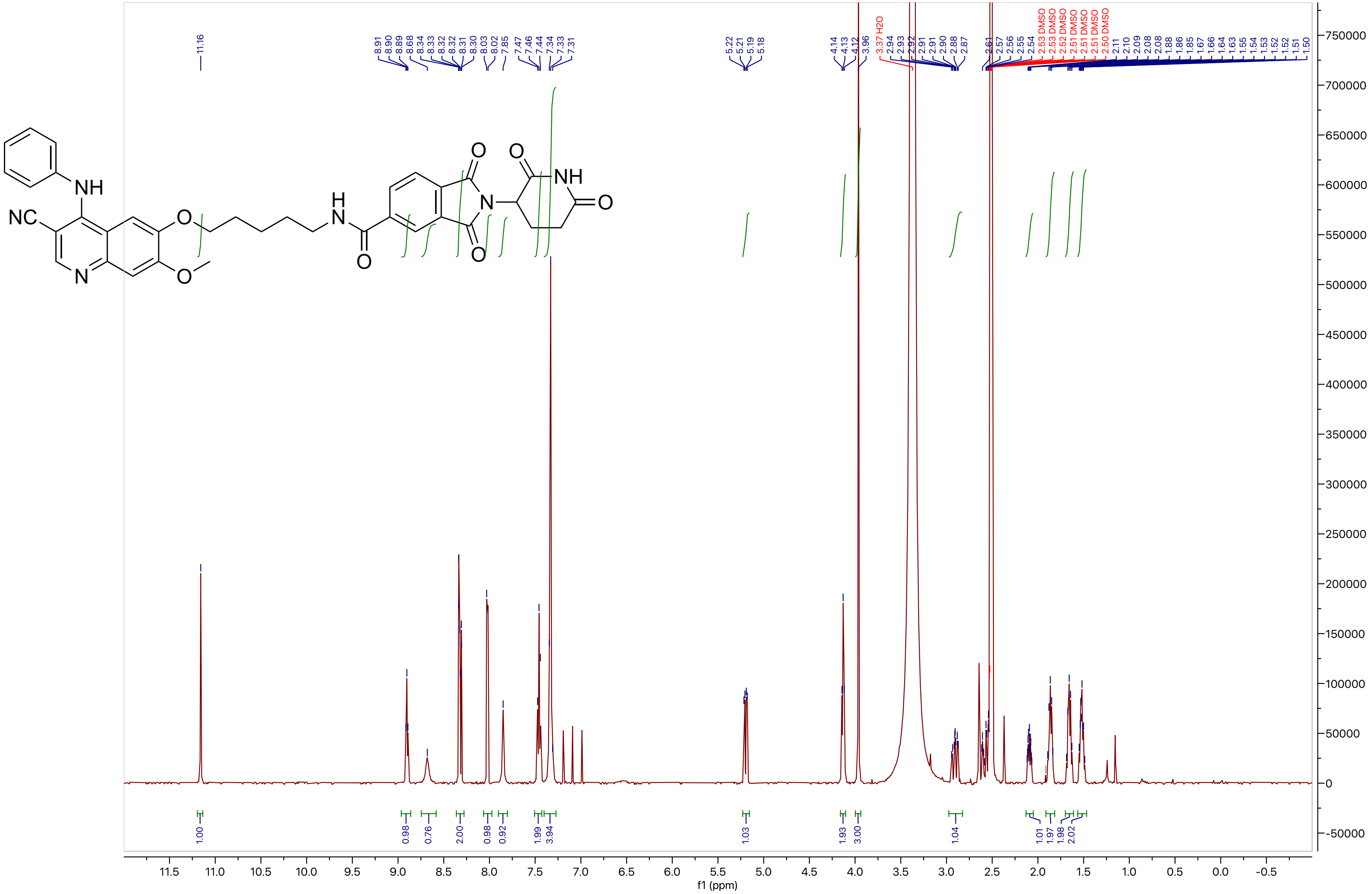

NH
NC
N
O
O
H
N
O
O
N
O
O
NH
O
11.16
8.91 8.90 8.89 8.68 8.34 8.33 8.32 8.32 8.31 8.30 8.03 8.02 7.85 7.47 7.46 7.44 7.34 7.33 7.31
5.22 5.21 5.19 5.18
4.14 4.13 4.12 3.96
3.37 H2O
2.94 2.93 2.92 2.91 2.91 2.90 2.88 2.87 2.61 2.57 2.56 2.55 2.54
2.53 DMSO 2.53 DMSO 2.52 DMSO 2.51 DMSO 2.51 DMSO 2.51 DMSO 2.50 DMSO
2.11 2.10 2.09 2.08 2.08 1.88 1.86 1.85 1.67 1.66 1.64 1.63 1.55 1.54 1.53 1.52 1.52 1.51 1.50
1.00 0.98 0.76 2.00 0.98 0.92 1.99 3.94 1.03 1.93 3.00 1.04 1.01 1.97 1.98 2.02
11.5 11.0 10.5 10.0 9.5 9.0 8.5 8.0 7.5 7.0 6.5 6.0 5.5 5.0 4.5 4.0 3.5 3.0 2.5 2.0 1.5 1.0 0.5 0.0 -0.5
f1 (ppm)
750000 700000 650000 600000 550000 500000 450000 400000 350000 300000 250000 200000 150000 100000 50000 0 -50000

**Compound 51**

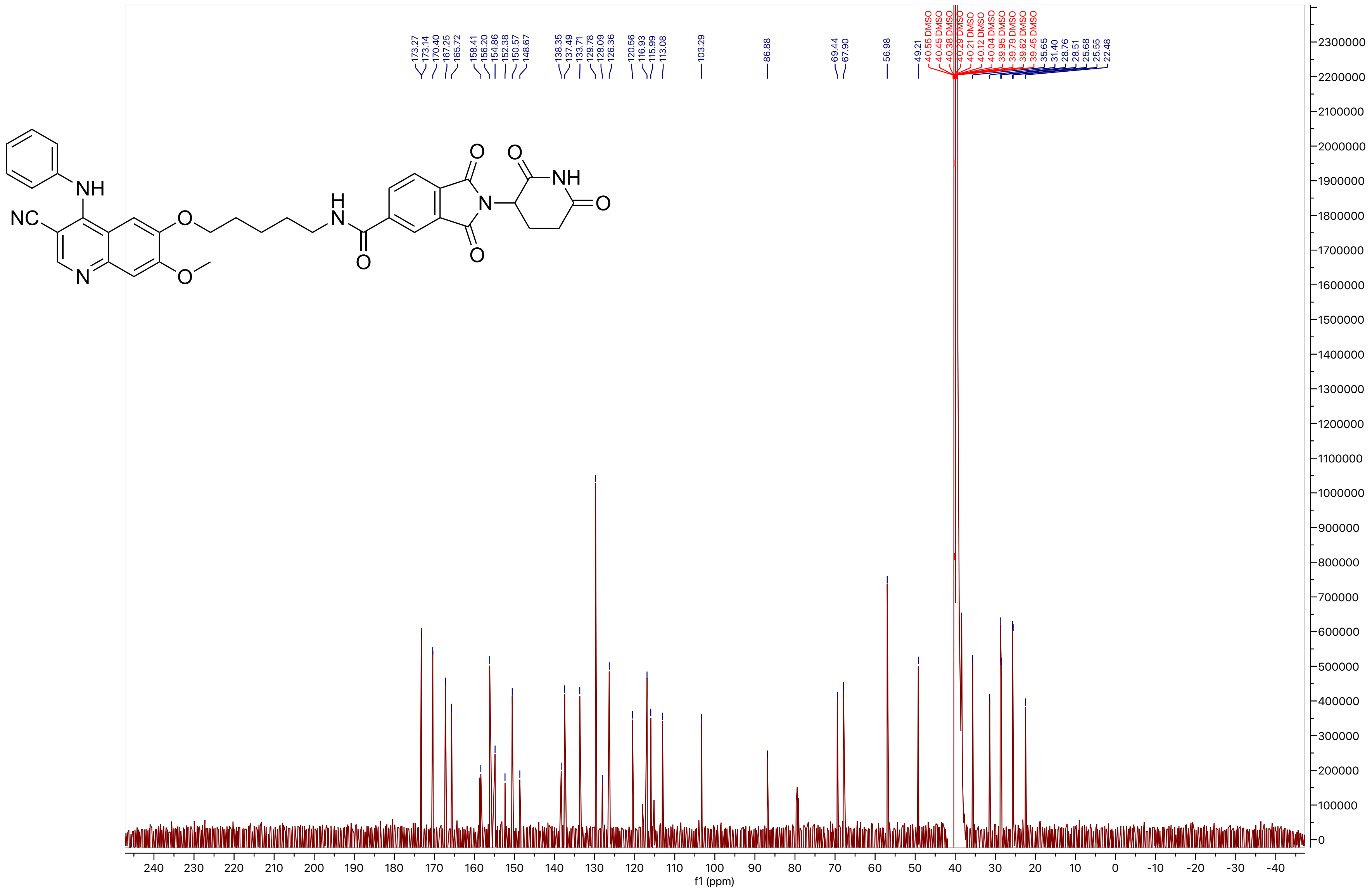

**Compound 52**

LCMS

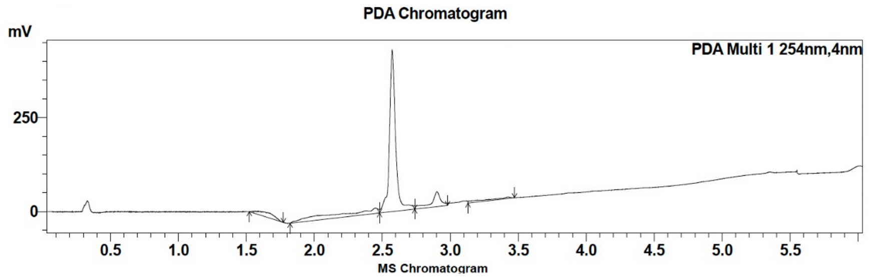

| Peak # | Ret. time | Area | Area % |
|---|---|---|---|
| 1 | 2.6 | 1360649 | 95.83 |
| 2 | 2.9 | 59220 | 4.17 |

HRMS

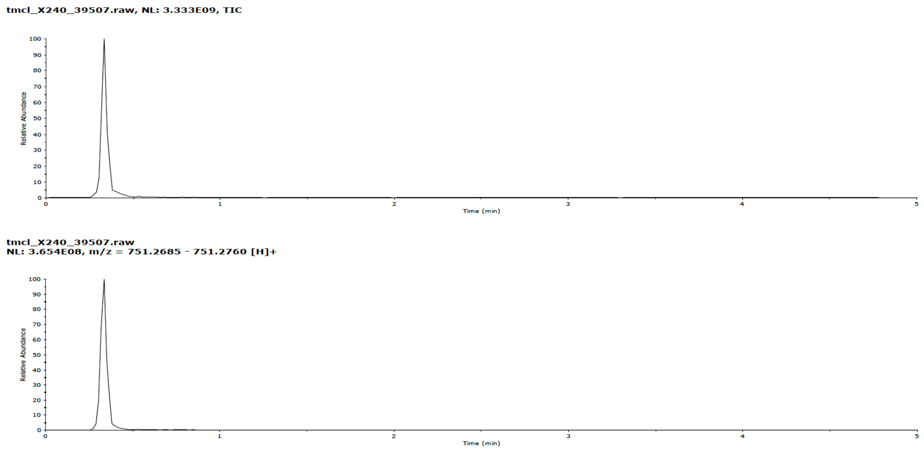

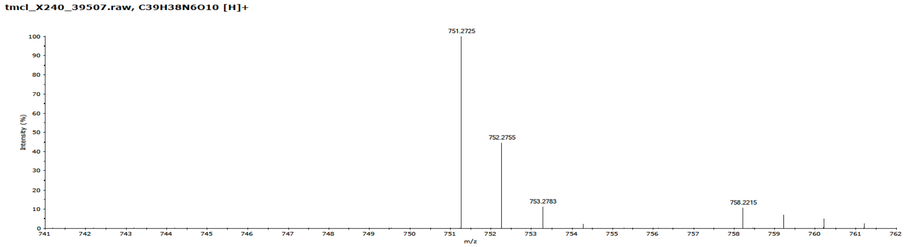

**Compound 52**

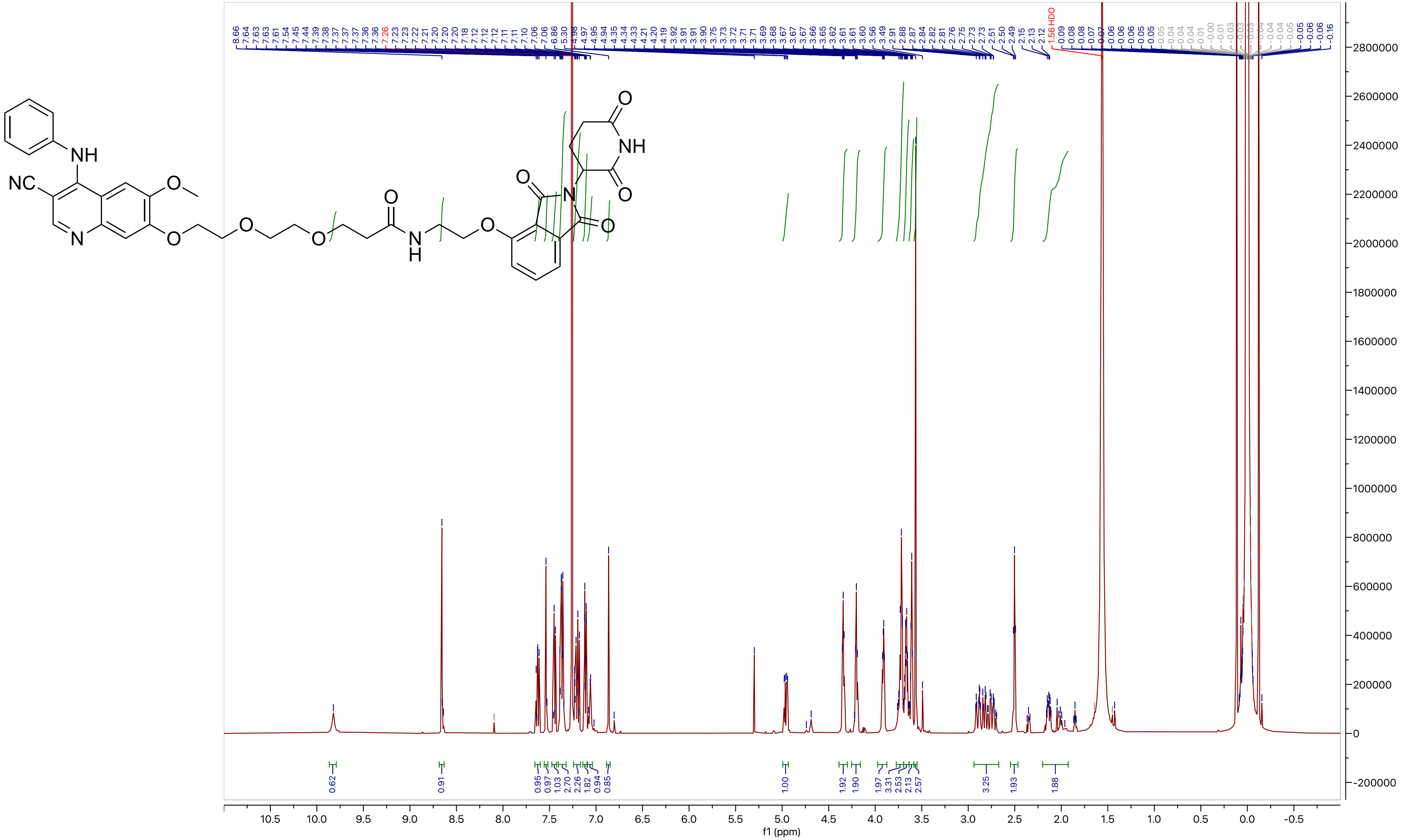

**Compound 52**

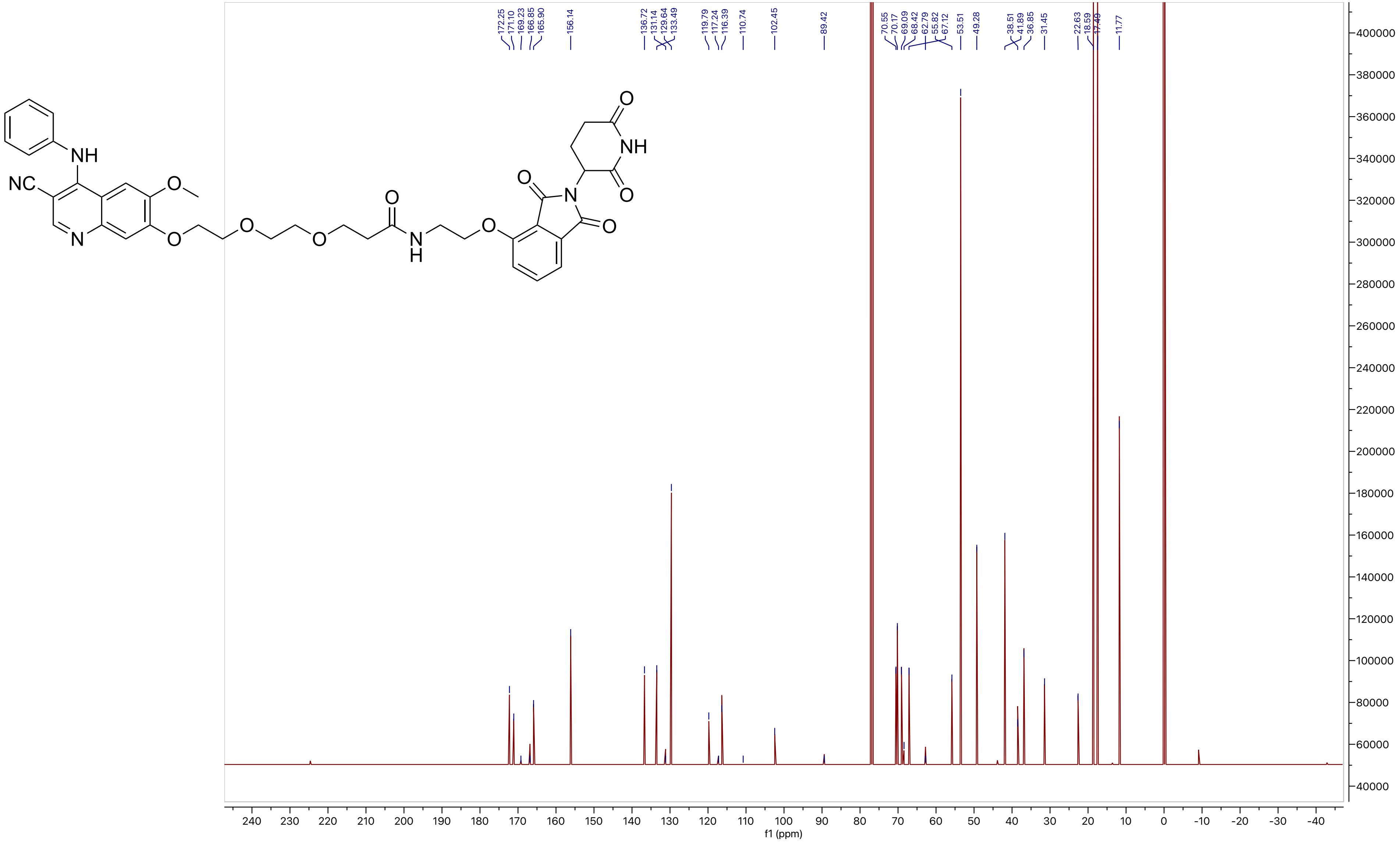
172.25
171.10
169.23
166.85
165.90
156.14
136.72
131.14
129.64
133.49
119.79
117.24
116.39
110.74
102.45
89.42
70.55
70.17
69.09
68.42
62.79
55.82
67.12
53.51
49.28
38.51
41.89
36.85
31.45
22.63
18.59
17.49
11.77
f1 (ppm)

**Compound 53**

LCMS

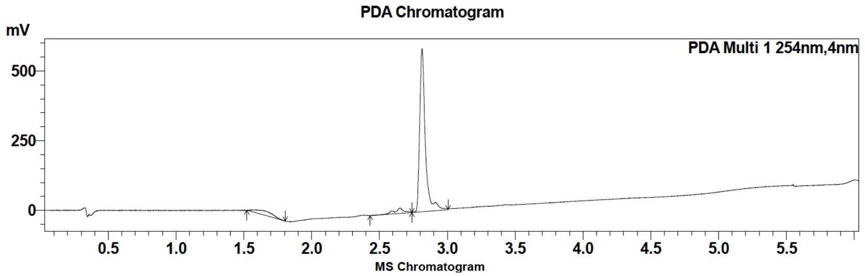

| Peak # | Ret. time | Area | Area % |
|---|---|---|---|
| 1 | 2.7 | 81787 | 4.62 |
| 2 | 2.8 | 1688794 | 95.38 |

HRMS

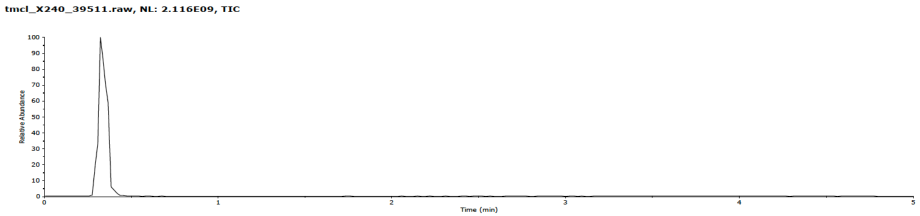

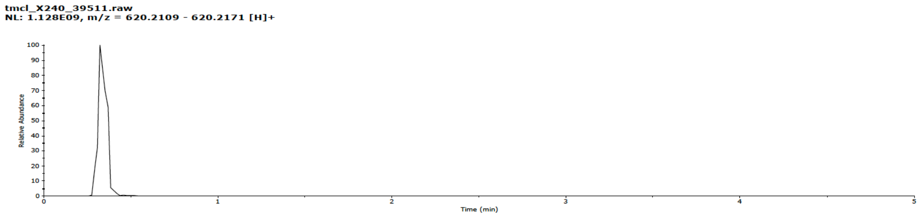

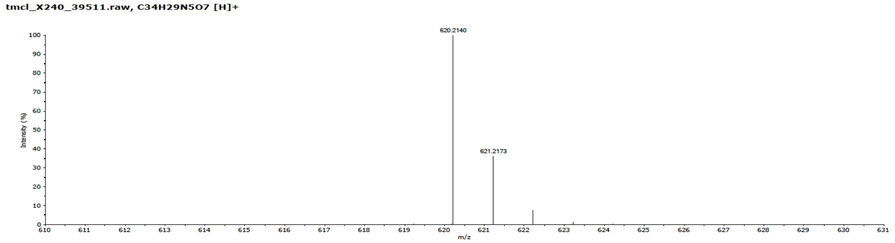

#### Compound 53

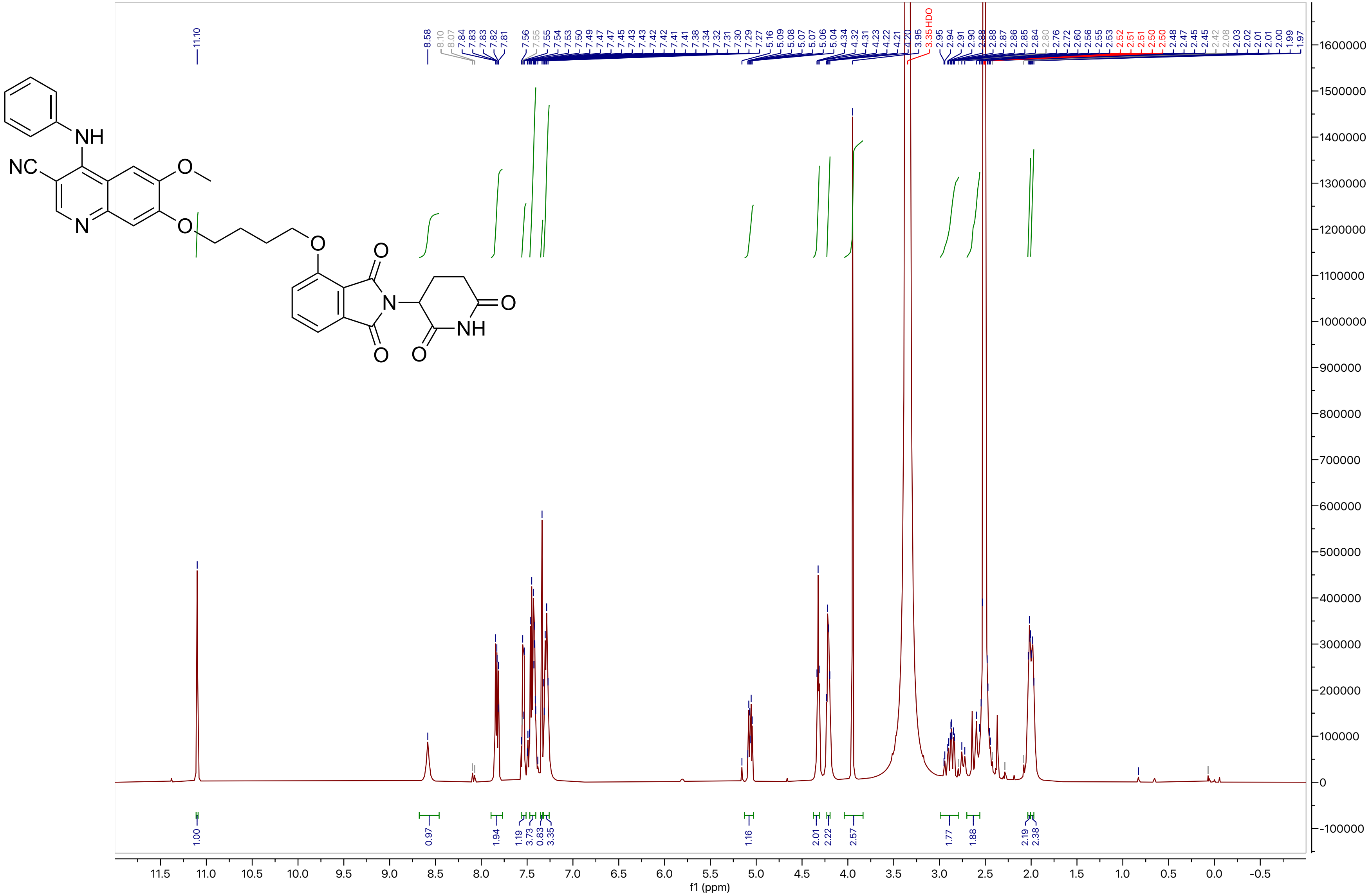

**Compound 53**

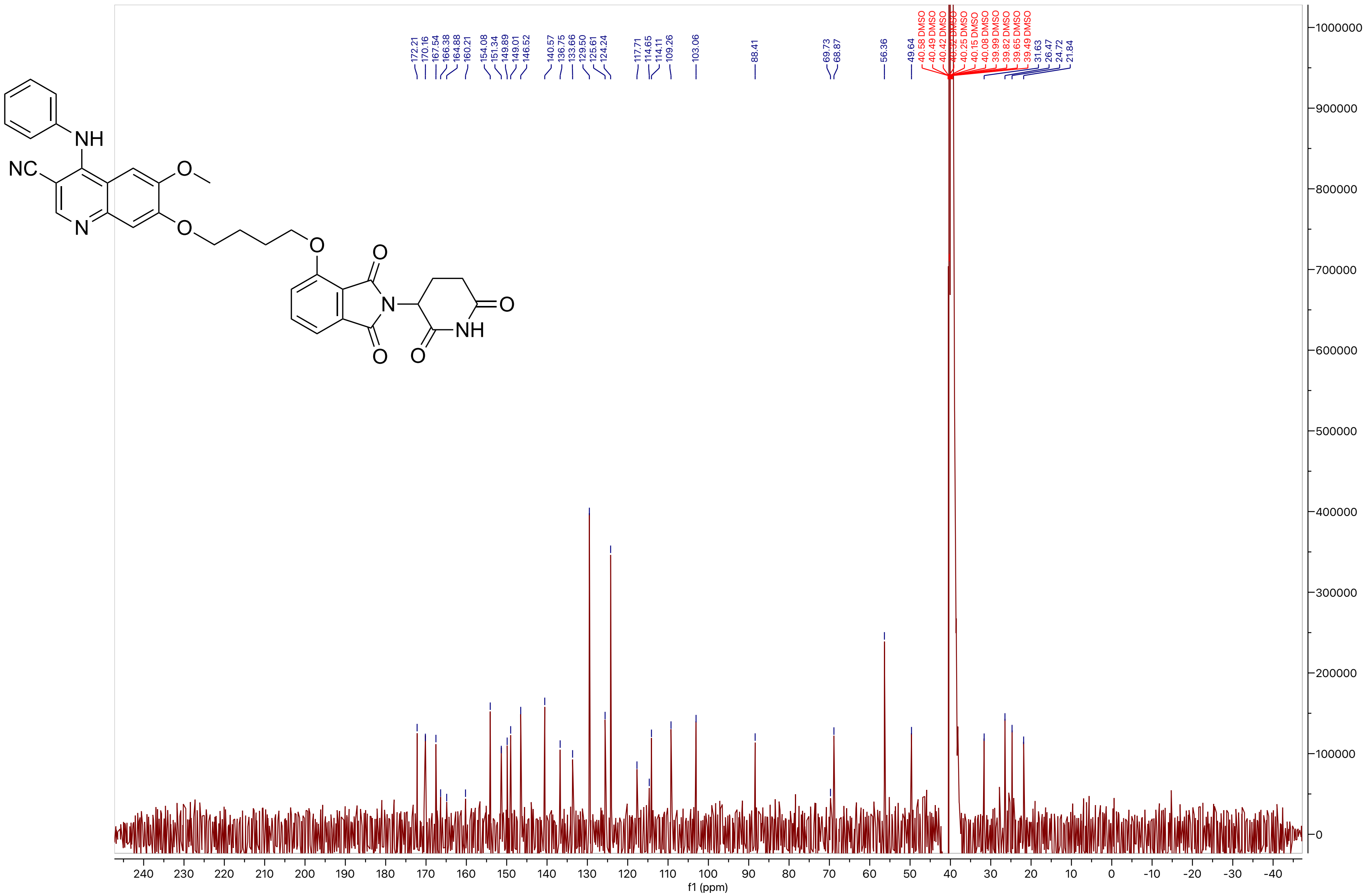

**Compound 54**

LCMS

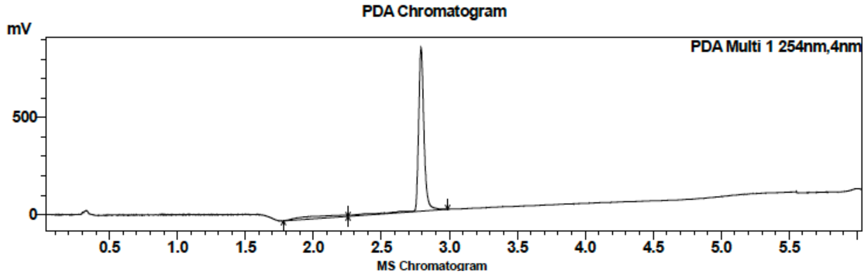

| Peak # | Ret. time | Area | Area % |
|---|---|---|---|
| 1 | 2.8 | 2399455 | 100.00 |

HRMS

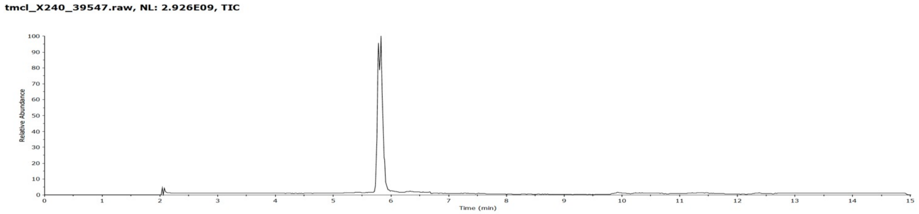

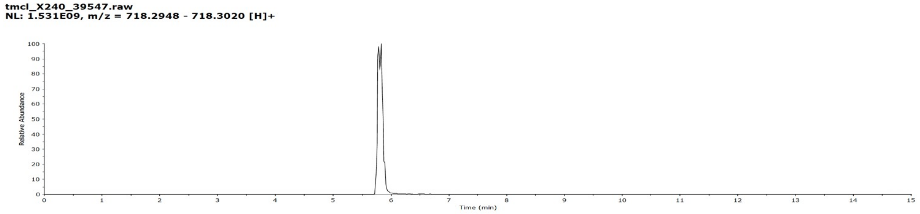

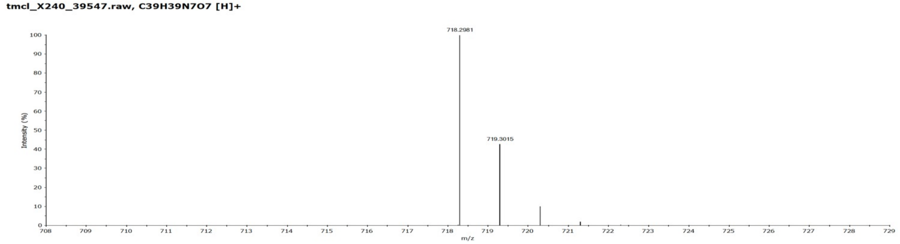

**Compound 54**

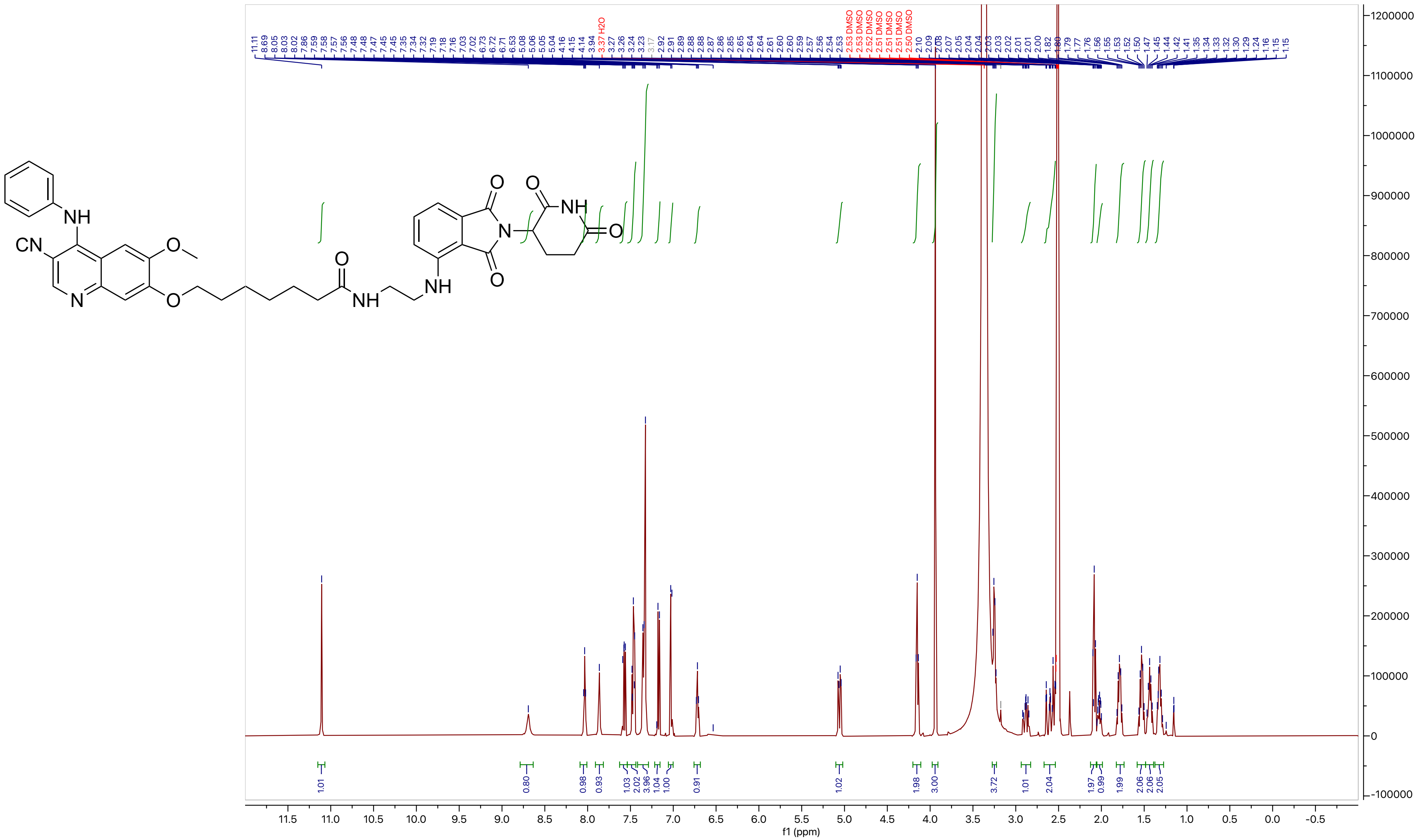

**Compound 54**

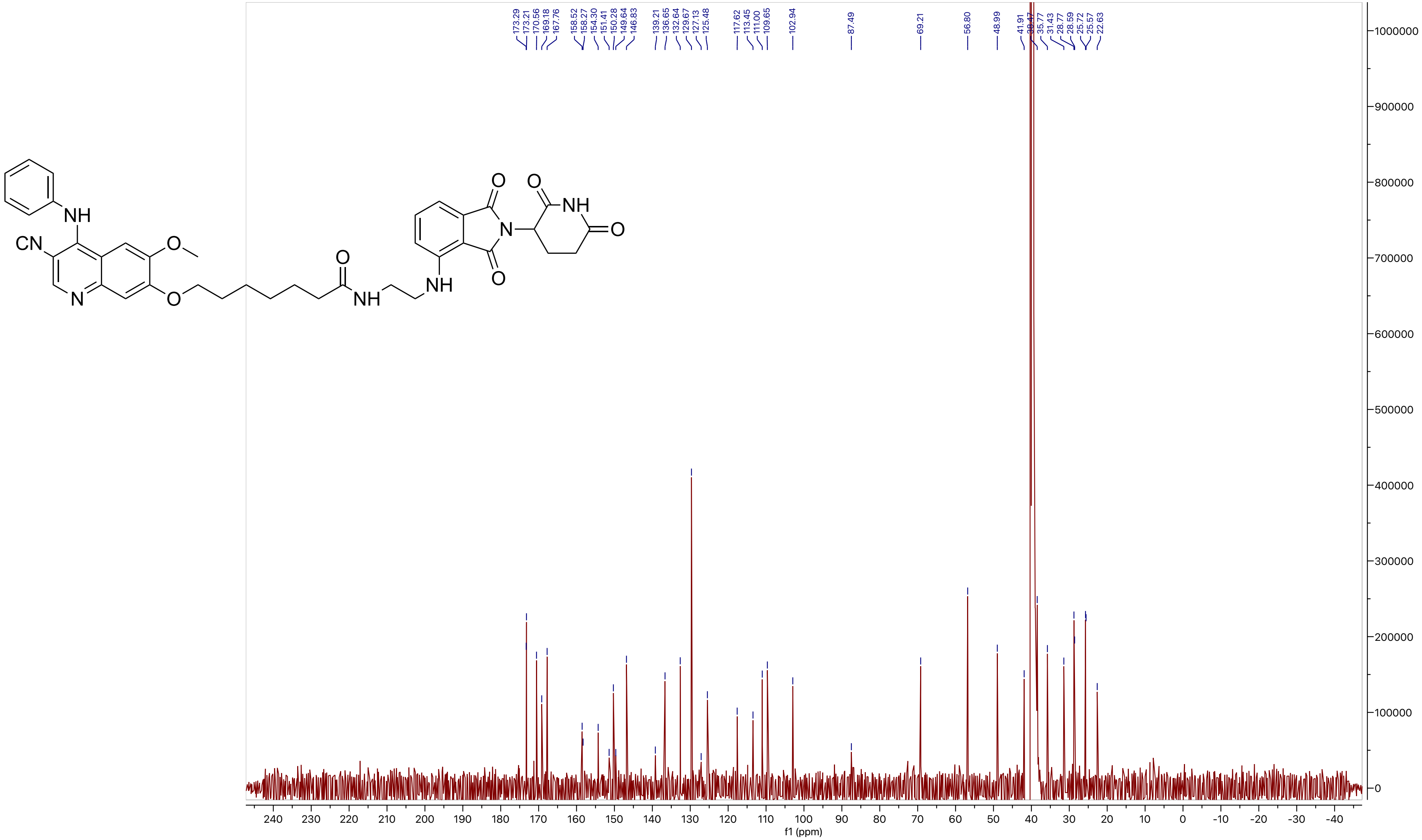
NH
CN
O
N
O
O
NH
NH
O
N
O
O
NH
O
173.29
173.21
170.56
169.18
167.76
158.52
158.27
154.30
151.41
150.28
149.64
146.83
139.21
136.65
132.64
129.67
127.13
125.48
117.62
113.45
111.00
109.65
102.94
87.49
69.21
56.80
48.99
41.91
35.77
31.43
28.77
28.59
25.72
25.57
22.63
f1 (ppm)

#### Compound 55

LCMS

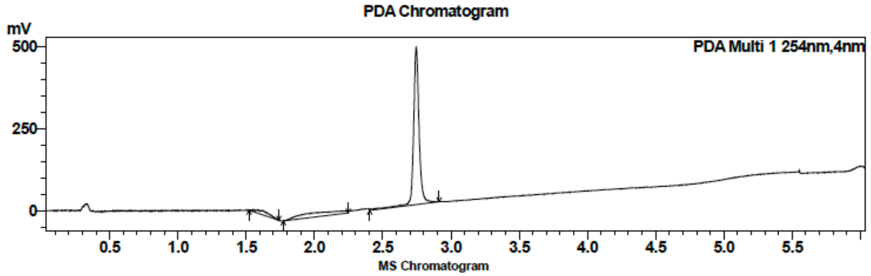

| Peak # | Ret. time | Area | Area % |
|---|---|---|---|
| 1 | 2.7 | 1346782 | 100.00 |

HRMS

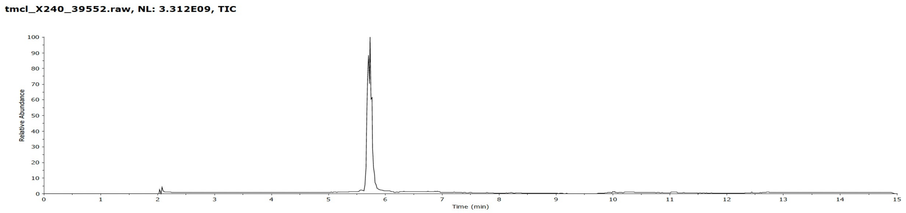

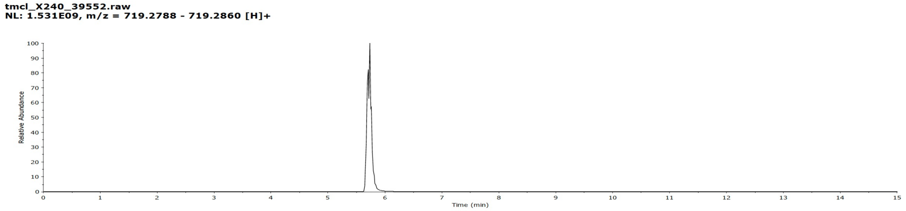

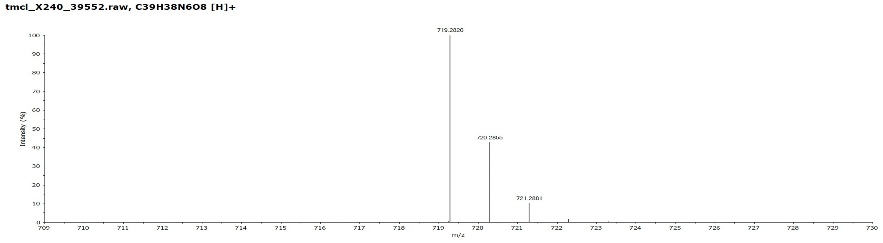

**Compound 55**

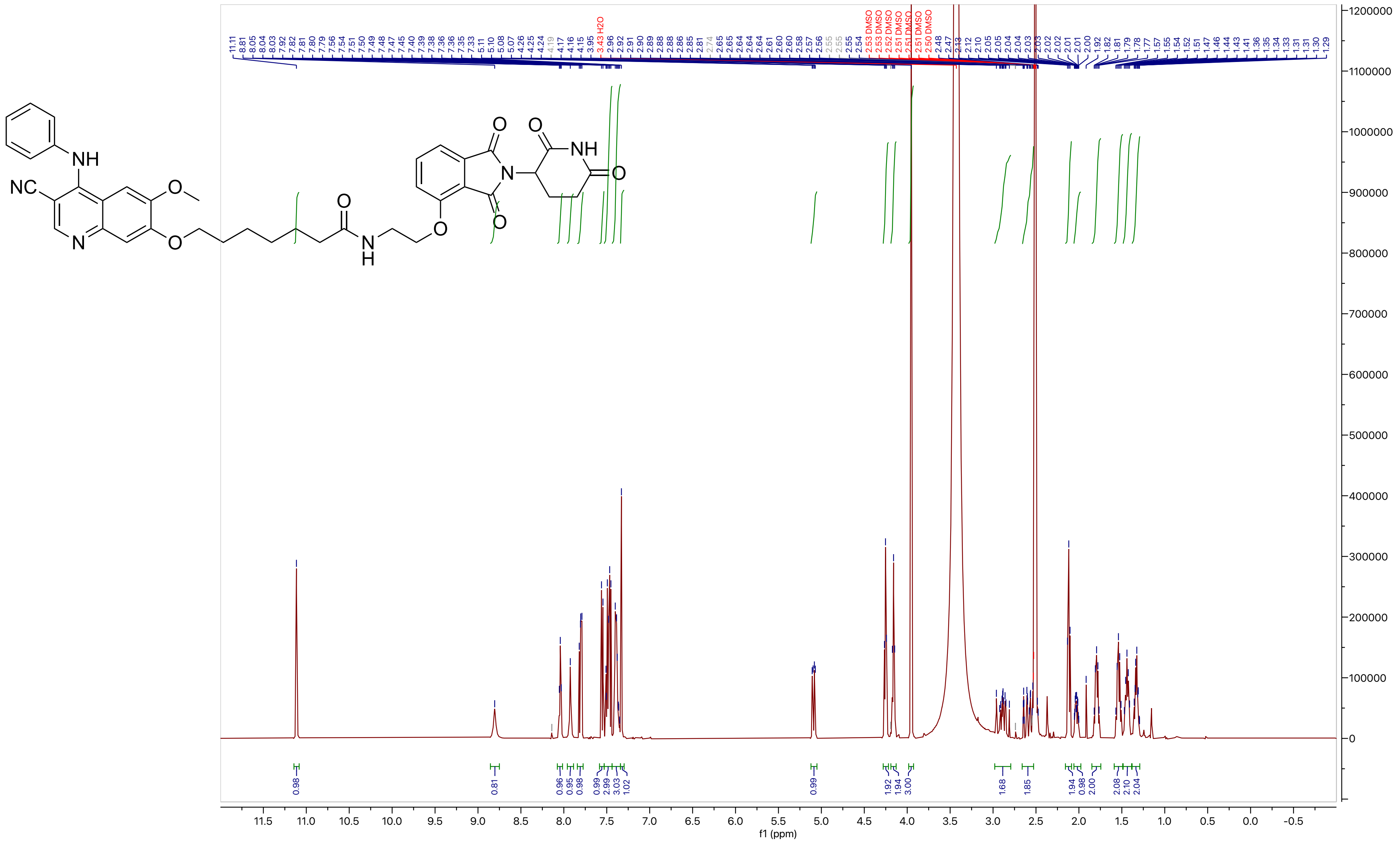

**Compound 55**

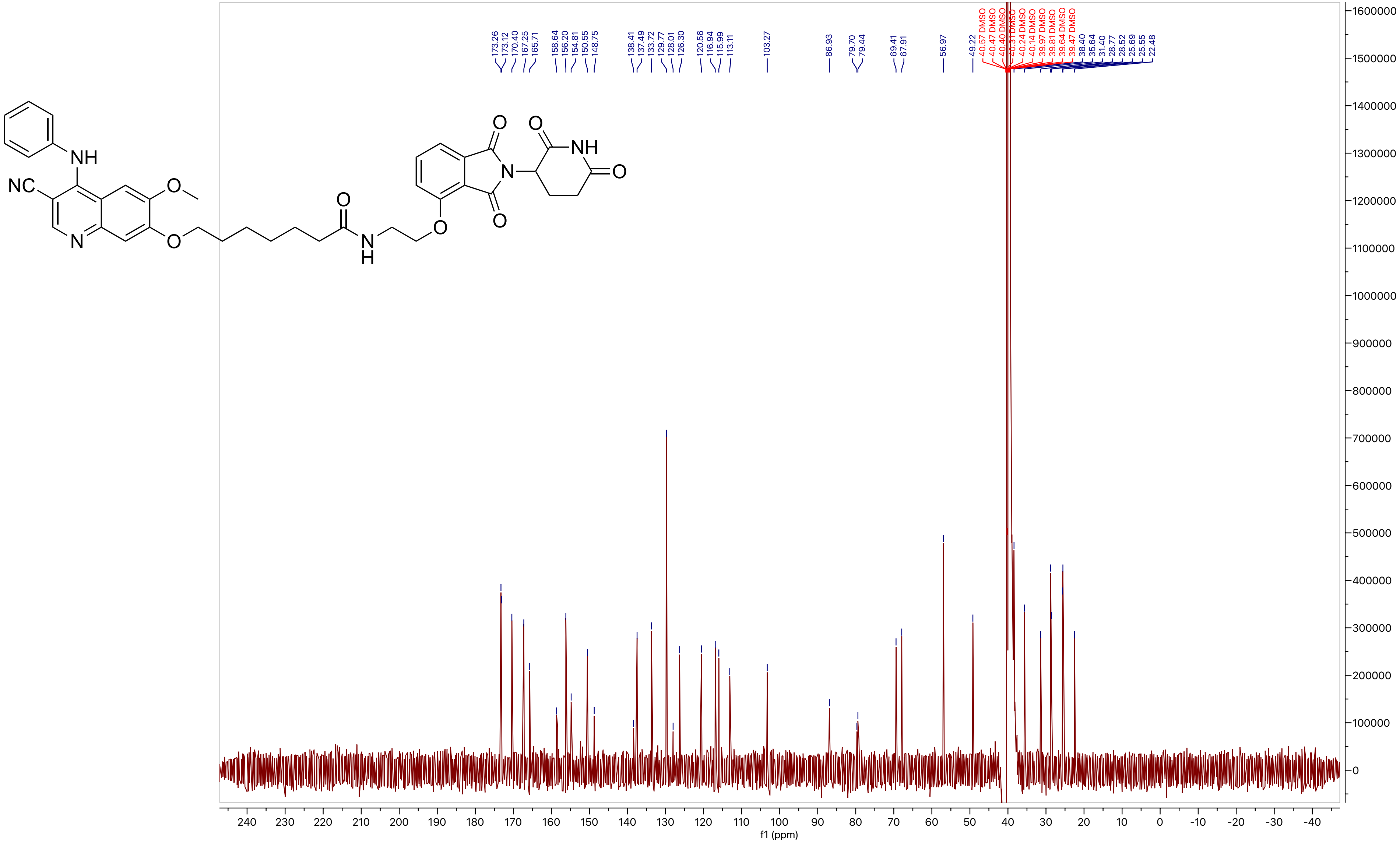
